# Selective Editing and Functionalization of the Mammalian Lipidome

**DOI:** 10.64898/2026.04.24.720406

**Authors:** Binyou Wang, Lukas Lüthy, Logan Tenney, Leo Qi, Takeshi Harayama, Kim Ekroos, Johannes Morstein

**Affiliations:** Division of Chemistry and Chemical Engineering, California Institute of Technology, Pasadena, California 91125, USA; Institut de Pharmacologie Moléculaire et Cellulaire, Université Côte d’Azur - CNRS UMR7275 - Inserm U1323, Valbonne, France; Lipidomics Consulting Ltd., Esbo, Finland

## Abstract

Lipids exhibit extraordinary molecular diversity, yet tools to selectively manipulate defined lipid classes in living cells are lacking. Here we show that lipid tail structure biases metabolic fate, enabling the design of synthetic lipid analogs with programmable metabolic selectivity. This approach enables selective cellular production of distinct lipid species or subclasses, including types of neutral lipids, phospholipids, sphingolipids, and ether lipids, without genetic or enzymatic perturbation. We further couple metabolic selectivity to chemical functionalization using bifunctional lipids, in which one modification directs metabolic flux and a second enables bioorthogonal tagging. Using this strategy, we achieve selective *in situ* labeling of different lipid pools in living cells. Together, our work establishes a chemical biology strategy that enables unprecedented precision in modulating, functionalizing, and rewiring the mammalian lipidome.

## Introduction

Lipids are central to cellular organization, metabolism, and signaling, yet they remain one of the least tractable classes of biomolecules for systematic functional interrogation.^1–3^ Lipid databases such as SwissLipids^4^ and LIPID MAPS^5^ now catalog over 10^6^ distinct lipid species, underscoring the vast chemical diversity of the lipidome. Despite this complexity, tools to selectively manipulate defined lipid classes or species in living cells are limited, restricting our ability to establish causal links between lipid identity and cellular function.^6–8^ Current approaches to perturb lipid biology rely largely on genetic or pharmacological modulation of lipid-metabolizing enzymes, which often produce broad and pleiotropic effects.

Recent advances in lipid chemical biology have significantly expanded our ability to interrogate lipid function in living systems.^9^ Enzyme-based membrane editors enable targeted remodeling of lipid headgroups at defined cellular locations with temporal control.^10,11^ Bifunctional phospholipids can be directly inserted into cellular membranes and subsequently fixed and labeled, providing a powerful approach to visualize lipid trafficking and distribution in living cells.^12,13^ Broad metabolic labeling strategies combined with organelle- or leaflet-specific reporters further enable spatially resolved analysis of lipid incorporation and turnover.^14,15^ Despite these advances, existing approaches primarily enable the observation or localized modification of lipids and do not provide a general strategy to control how lipids are routed through endogenous metabolic networks. Emerging evidence indicates that lipid tail structure is a key determinant of metabolic fate.^16^ We therefore reasoned that systematic chemical modification of lipid substrates could be used to bias their routing through metabolic networks and produce selective outcomes. In contrast to classical analog-sensitive chemical genetics,^17^ which relies on protein engineering to create orthogonal enzyme-substrate pairs, we hypothesized that native lipid metabolic pathways can discriminate neosubstrates through differential shape sensing in lipid binding pockets.

Here, we systematically define the structural dependence of lipid metabolism and test whether lipid tails can be engineered to control metabolic routing. Across diverse lipid classes, we show that lipid tail modifications enable selective metabolic routing and functionalization, establishing a general chemical biology strategy to edit and functionalize the mammalian lipidome. This approach creates new opportunities to interrogate lipid biology and to reprogram lipid flux in metabolic, neurological, and other diseases linked to dysregulated lipid metabolism.^18^

## Results

Fatty acids are central lipid metabolites and are incorporated into diverse lipid classes, including phospholipids, neutral lipids, lysolipids, and sphingolipids through the action of various lipid trafficking and metabolizing enzymes (Figure 1A). In humans, some fatty acids are exclusively sourced exogenously and dietary interventions with certain fatty acids have shown benefits in several clinical trials.^19,20^ To systematically map the structural dependence of lipid metabolism, we established an untargeted chemical lipidomics platform in which we treat cells with labeled lipids for a defined time, perform lipid extraction with MTBE,^21^ and conduct lipidomic analysis^22^ capturing both endogenous and labeled lipid metabolites (Figure 1B). Comparing four common fatty acids with distinct chemical structures (Figure 1C), we observed pronounced structural dependence in their metabolic incorporation across lipid classes (Figure 1D). Palmitic acid (d5-PA) showed reduced incorporation into phosphatidylethanolamine (PE), phosphatidylinositol (PI), and phosphatidylserine (PS), whereas arachidonic acid (d11-AA) was poorly incorporated into lysolipids and acylcarnitine (CAR). Saturated fatty acids were incorporated more effectively into phosphatidylglycerol (PG) and oleic acid showed the highest incorporation into triacyclglycerols (TAG) consistent with its role as a potent lipid droplet inducer.^23,24^ To assess cell-type dependence, we extended this analysis across HeLa, A549, Caco-2, and HepG2 cells and found that relative incorporation trends were conserved (Figure 1E). Different lipid classes exhibited distinct incorporation kinetics (Figure S1A-F) and the overall incorporation selectivity patterns were robust and not effected by variables such as cellular confluence (Figure S1G). MS/MS analysis of d7-SA revealed minimal elongation or truncation after 4 h, with limited desaturation primarily funneled into phosphatidylcholine (PC) (Figure S1H). All detected metabolites, including elongated, truncated, and desaturated species, were included in the analysis to capture overall metabolic selectivity. To determine whether fatty acid supplementation alters total lipid output, we quantified lipid overproduction by comparing the sum of endogenous and labeled lipids to untreated controls (Figure 1F). PC, PE, PI, PS, DAG, and TAG showed clear increases, whereas total PG levels remained constant, suggesting tighter homeostatic regulation of PG and compensation via reduction of endogenous species within this class.

**Figure 1.**
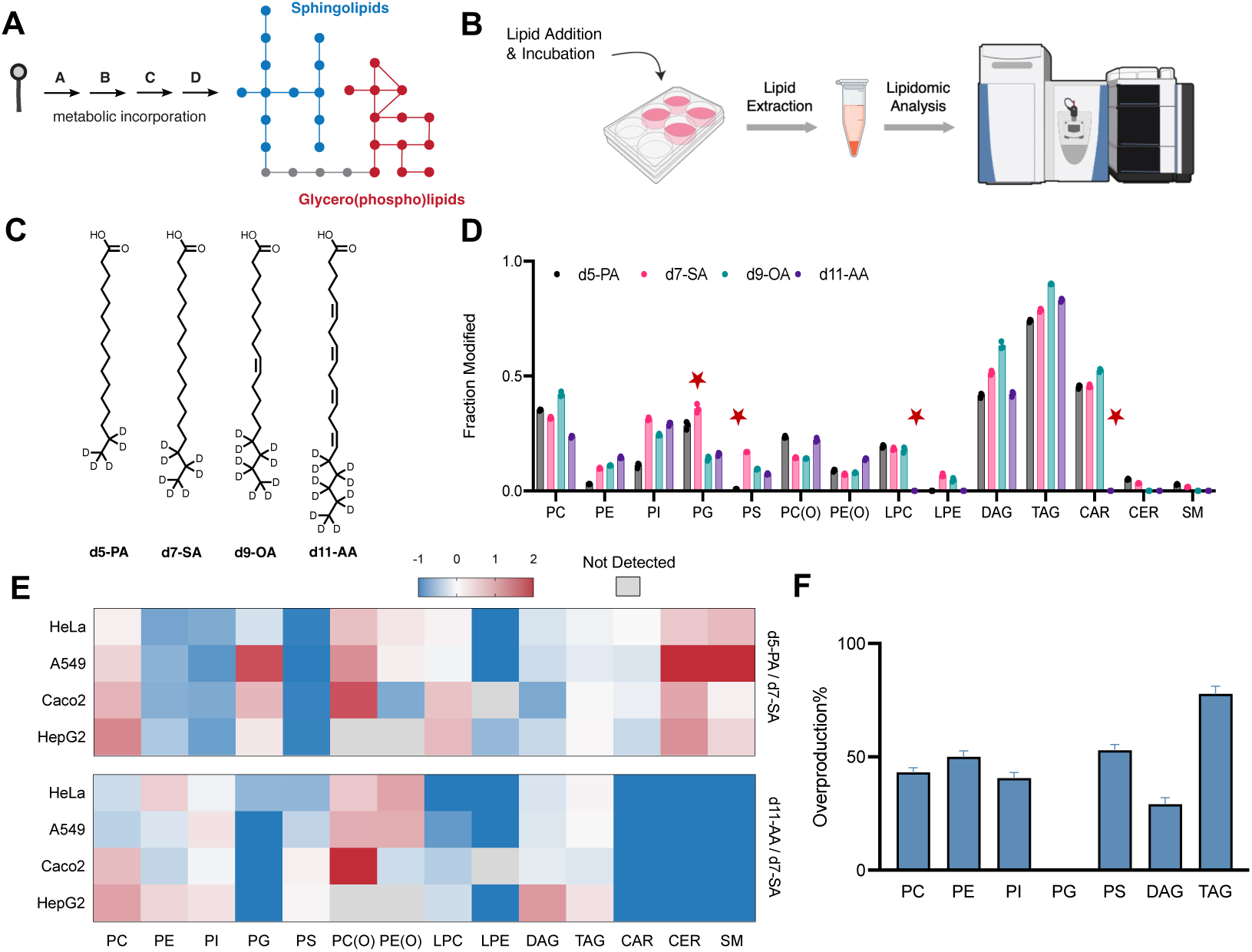
Lipid tail structure directs metabolic fate across the mammalian lipidome. **(A)** Schematic illustrating incorporation of exogenous lipid into cellular lipidome. **(B)** Schematic of chemical lipidomics workflow. Cells were treated with defined lipid species, followed by incubation, lipid extraction, and quantitative lipidomics analysis. **(C)** Chemical structures of isotopically labeled fatty acids: d5-palmitic acid (d5-PA), d7-stearic acid (d7-SA), d9-oleic acid (d9-OA), and d11-arachidonic acid (d11-AA). **(D)** Lipidomics analysis showing incorporation of labeled fatty acids (50 μM for 4 h in HeLa cells) across major lipid classes, shown as the fraction of modified lipids relative to endogenous levels (n = 3, mean ± s.d.). **(E)** Heatmap comparing relative incorporation efficiencies, expressed as ratios of d5-PA to d7-SA and d11-AA to d7-SA, across multiple cell lines (HeLa, A549, Caco-2, and HepG2). **(F)** Relative overproduction of lipid classes upon treatment with d7-SA compared to untreated controls (n = 6, mean ± s.d.).

While our data demonstrates dependence of metabolic fate on the structure of a fatty acid tail, overall incorporation of endogenous lipids remains relatively broad (Figure 2A). We reasoned that further chemical diversification could generate novel and more selective incorporation patterns, enabling targeted modulation of the lipidome (Figure 2B). To explore the feasibility of this hypothesis, we profiled branched fatty acids bearing methyl groups at defined positions, as well as a cyclopropyl analog (Figure 2C). These non-native lipids exhibit minimal endogenous background and are readily distinguishable in mass and retention time, enabling direct tracking without incorporation of isotopic labels. Although such branched fatty acids are not produced in mammalian cells, they occur in microorganisms and may influence host metabolism through the gut microbiome.^29^ Notably, we found striking differences in the metabolism and more selective incorporation patterns compared to the previously profiled dietary fatty acids (Figure 2D,E). **2-MeSA** was selectively routed into neutral lipids and acylcarnitine, with minimal phospholipid incorporation. In contrast, **10-MeSA** efficiently entered phospholipids with enrichment in PI and PG, whereas **13-MeMA** showed the opposite preference, favoring PC and (O)-PC over PI and PG. **CP-SA** exhibited a profile similar to **10-MeSA**, consistent with the shared position of modification. We next examined aromatic fatty acids with bulkier substitutions along the tail (Figure 2F). Similar to **2-MeSA**, an modification near the headgroup in **π-FA4** afforded selective incorporation into neutral lipids, with no detectable phospholipids or acylcarnitines (Figure 2G,H). In contrast, distal modifications (**π-FA1** and **π-FA2**) exhibited incorporation into both neutral lipids and phospholipids, albeit with distinct class distribution. Together, these results demonstrate that lipid tail modifications can program metabolic selectivity, enabling targeted enrichment of neutral lipids or defined phospholipid subsets. These patterns likely reflect structural constraints within lipid metabolic enzymes that encode differential substrate recognition. To test this hypothesis, we mapped the lipid binding pockets in three key metabolic enzymes, DGAT1, GPAT1, and CPT2 using PyVOL^28^, and found substantial variation in pocket size and shape. DGAT1 exhibits the largest and most diffuse lipid binding pocket, consistent with its ability to accommodate bulky modifications and route substrates into neutral lipid metabolism (Figure 2I). In contrast, GPAT1 contains a smaller, more restrictive pocket (Figure 2J), while CPT2 displays an intermediate architecture (Figure 2K), consistent with selective conversion of certain fatty acids into acylcarnitines and exclusion of others. Finally, we asked whether selective lipid metabolism translates into lipid-dependent cellular phenotypes by examining lipid droplet formation. **π-FA1** was efficiently converted into TAGs, whereas **π-FA3** showed minimal TAG formation. Consistent with these metabolic differences, **π-FA1** robustly induced lipid droplet formation, while **π-FA3** had little effect (Figure 2L,M).

**Figure 2.**
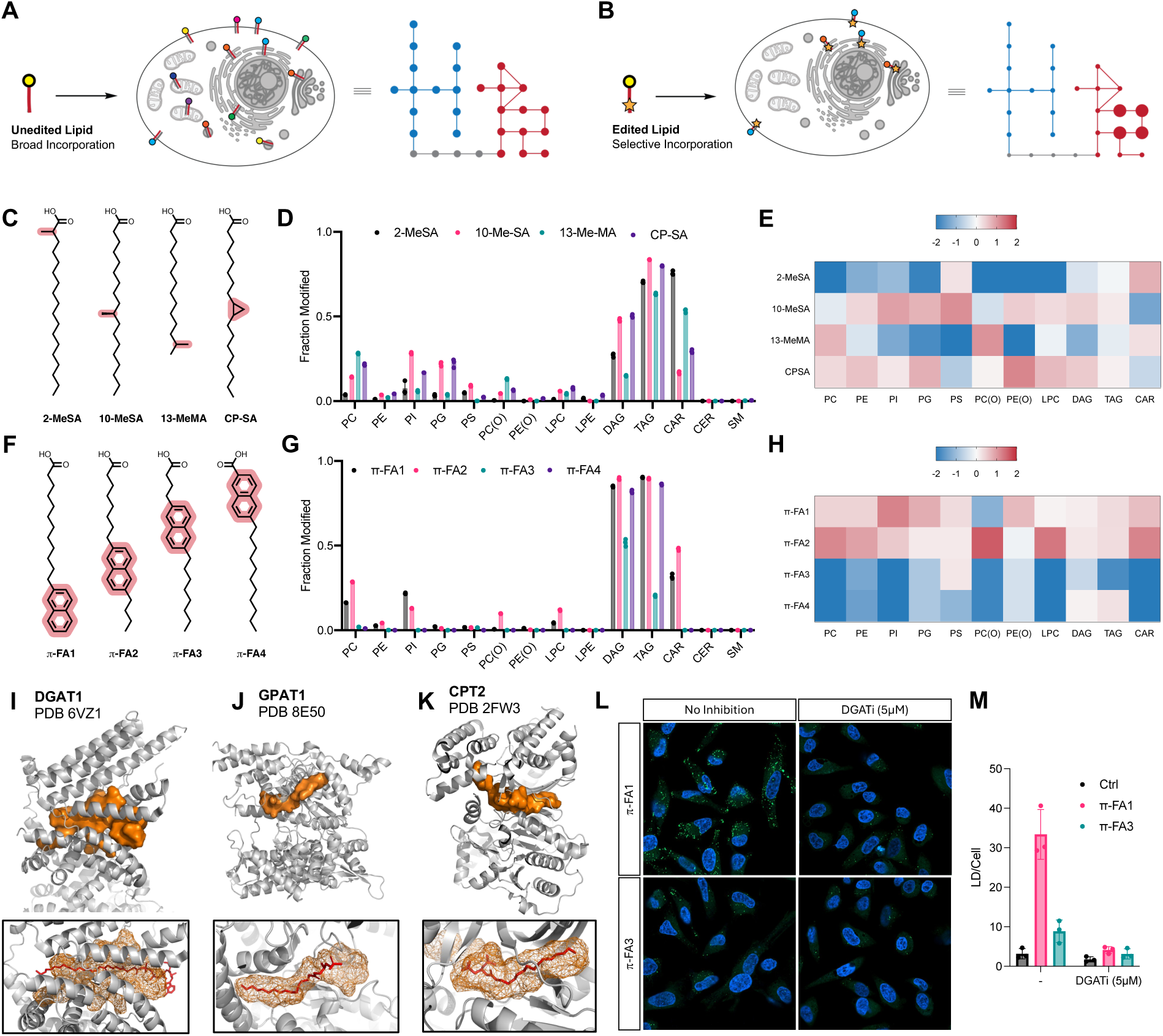
Chemical Editing of the Glycero(phospho)lipidome. **(A)** Schematic of broad lipidome-wide incorporation of endogenous fatty acids. **(B)** Schematic of selective incorporation of a chemically edited fatty acid. **(C)** Chemical structures of branched fatty acids. **(D)** Lipidomics analysis showing incorporation of branched fatty acids (50 μM for 4 h in HeLa cells) across major lipid classes, shown as the fraction of modified lipids relative to endogenous levels (n = 3, mean ± s.d.). **(E)** Heat map showing the relative change in branched fatty acid incorporation (log_2_-fold). **(F)** Chemical structures of aromatic fatty acids. **(G)** Lipidomics analysis showing incorporation of aromatic fatty acids (50 μM for 4 h in HeLa cells) across major lipid classes, shown as the fraction of modified lipids relative to endogenous levels (n = 3, mean ± s.d.). **(H)** Heat map showing the relative change in aromatic fatty acid incorporation (log_2_-fold). **(I–K)** Map of lipid binding pocket in lipid metabolic enzymes DGAT1 (PDB 6VZ1^25^), GPAT1 (PDB 8E50^26^), and CPT2 (PDB 2FW3^27^) generated using PyVOL.^28^ **(L)** Lipid Droplet Imaging in HeLa cells treated with 50 µM aromatic fatty acid for 4 h in the presence and absence of DGAT1 (15 μM A-922500) inhibitors. Lipid Droplets were stained with BODIPY 493/503 and nuclei with Hoechst 33342. **(M)** Quantification of lipid droplet content per cell (n = 3, mean ± s.d.).

We next asked whether combining chemical modifications with distinct metabolic entry points could further expand control over lipid incorporation. Alkylglycerols are thought to be selectively routed into ether lipids (Figure 3A).^30^ To test this, we synthesized deuterated and aromatic analogs (Figure 3B) and observed highly selective incorporation into ether lipids (Figure 3C; Figure S2A-H). Beyond class-level selectivity, these modifications enabled tuning of specific ether lipid species within a class, affording a level of control not readily achievable with genetic or pharmacological approaches (Figure 3D-F; Figure S2I-N). Supplementation with ether lipids further induced secondary changes to endogenous lipid pools (Figure 3G-I), including remodeled ceramides, consistent with reported crosstalk between ether lipids and sphingolipids.^31^ Notably, acylcarnitines were strongly upregulated following alkylglycerol treatment, suggesting a previously unrecognized link between ether lipid metabolism and acylcarnitine-associated energy metabolism.

**Figure 3.**
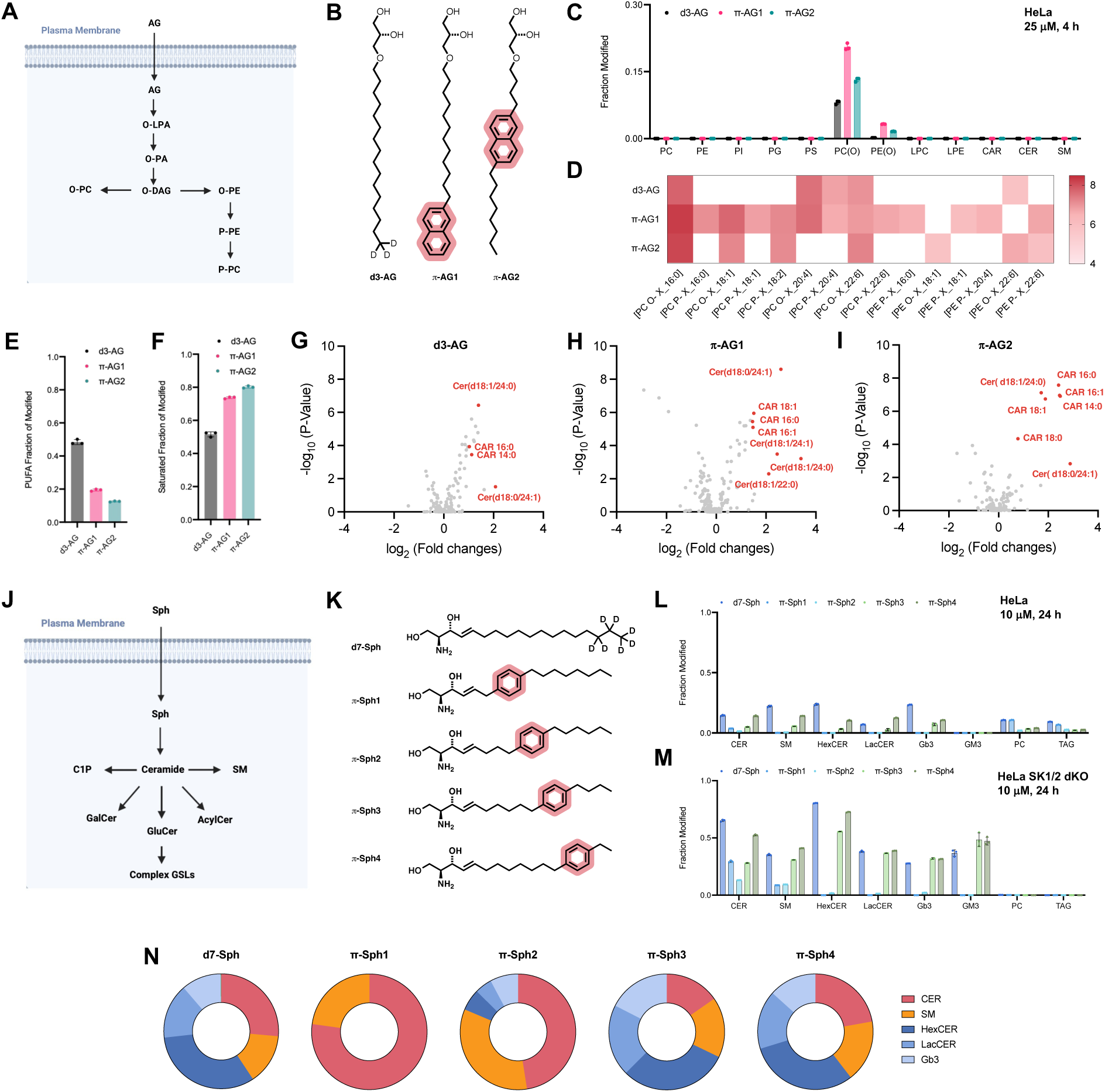
Chemical editing of the ether- and sphingolipidome. **(A)** Schematic of alkylglycerol (AG) metabolism. **(B)** Chemical structures of modified alkylglycerols. **(C)** Lipidomics analysis showing incorporation of alkylglycerols (50 μM for 4 h in HeLa cells) across major lipid classes, shown as the fraction of modified lipids relative to endogenous levels (n = 3, mean ± s.d.). **(D)** Heat map showing the relative levels (log_10_) of ether lipids formed from modified alkylglycerols (50 μM for 4 h). **(E)** Fraction of polyunsaturated lipid tails (PUFAs) in modified ether lipids. **(F)** Fraction of saturated lipid tails in modified ether lipids. **(G–I)** Volcano plots showing changes in endogenous lipids upon alkylglycerol treatments. **(J)** Schematic of sphingosine (Sph) metabolism. **(K)** Chemical structures of aromatic sphingosine analogues. **(L)** Lipidomics analysis showing incorporation of sphingosines (10 μM for 24 h in HeLa cells) across different lipid classes, shown as the fraction of modified lipids relative to endogenous levels (n = 3, mean ± s.d.). **(M)** Lipidomics analysis showing incorporation of sphingosines (10 μM for 24 h in HeLa SK1/2 dKO cells) across different lipid classes, shown as the fraction of modified lipids relative to endogenous levels (n = 3, mean ± s.d.). **(N)** Distribution of modified lipid fractions of **d7-Sph** and **π-Sph1-4** after 24 treatment of HeLa SK1/2 dKO cells in **(M)**.

To extend this approach to sphingolipids, we focused on sphingosine as a central metabolic precursor. Ceramides are the central branch point of sphingolipid metabolism, but their poor cell permeability limits their direct use.^32,33^ In contrast, sphingosine is readily taken up and converted into ceramides by ceramide synthases^34^, followed by metabolism into major sphingolipid classes, including sphingomyelin and complex glycosphingolipids (Figure 3J).^35,36^ We synthesized four aromatic sphingosine derivatives (Figure 3K) and found distinct selectivity patterns: **π-Sph1** and **π-Sph2** were preferentially converted into sphingomyelin, whereas **π-Sph3** and **π-Sph4** were incorporated into sphingomyelin and complex glycosphingolipids (Figure 3L; Figure S3A,B To prevent conversion of sphingosine (Sph) into fatty acids via S1P and thereby enhance flux into complex sphingolipids, we employed a double sphingosine kinase knockout HeLa cell line, which resulted in increased sphingolipid production (Figure 3M,N; Figure S3C-F). These results demonstrate that sphingolipid class distribution can be selectively tuned through substrate design, with potential relevance for diseases characterized by aberrant glycosphingolipid accumulation, such as Gb3 buildup in Fabry’s disease.^37^

Chemically modified lipids with selective metabolic routing enable targeted overproduction of defined lipid classes, providing a new level of control over lipid metabolism. We reasoned that this selectivity could be extended to enable class-specific functionalization. To test this, we designed bifunctional fatty acid analogs comprising a chemical modification that directs metabolic routing and an azide handle for bioorthogonal functionalization via strain-promoted azide-alkyne cycloaddition (SPAAC^39^; Figure 4A). We hypothesized that **cl-2MeSA** would preferentially incorporate into neutral lipids, while **cl-10MeSA** would target phospholipids with a distinct incorporation profile relative to **cl-SA** (Figure 4B). Lipidomic analysis demonstrated preferential neutral lipid incorporation of **cl-2MeSA** and increased phospholipid labeling with **cl-SA** and **cl-10MeSA** (Figure 4C,D). While bifunctional analogs retained selectivity, it was less pronounced than for **2-MeSA** and **10-MeSA** (Figure 2), indicating that the azide modification influences metabolic incorporation. Consistent with this, all three clickable analogs showed increased β-oxidation, likely due to the extended tail length introduced by the azide, which reduces selective incorporation patterns (Figure S4A-H). Further structural optimization is therefore likely to enhance selectivity. Despite this, we observed pronounced differences in the pools of functionalized lipids by live-cell imaging following SPAAC labeling with a low-background strained-alkyne fluorophore (CO-1^38^; Figure 4E). **cl-SA** and **cl-10MeSA** preferentially labeled phospholipids, resulting in strong fluorescence at the endoplasmic reticulum and associated membranes, consistent with sites of phospholipid synthesis and remodeling (Figure 4F). In contrast, under conditions that promote lipid droplet formation, **cl-2MeSA** selectively labeled neutral lipid pools and lipid droplets, whereas **cl-SA** and **cl-10MeSA** remained enriched in phospholipids (Figure 4F,G; Figure S4I,J). Together, these results establish a strategy for *in situ* functionalization of distinct lipid pools, enabling selective visualization of newly synthesized lipids and, more broadly, chemical targeting of defined lipid classes.

**Figure 4.**
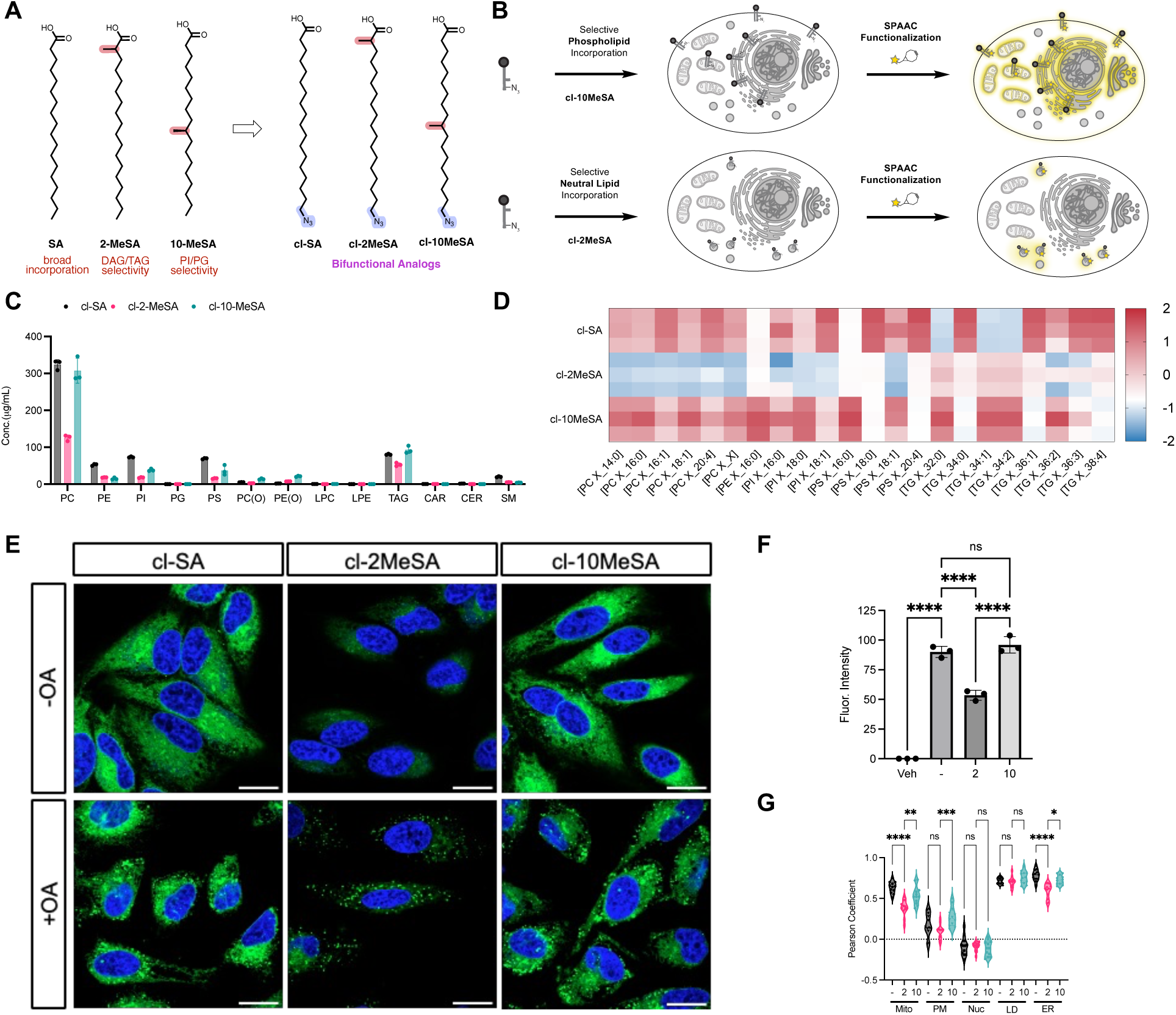
Selective *in situ* lipid functionalization. **(A)** Design and chemical structures of clickable branched fatty acid analogues. **(B)** Schematic illustrating the selective metabolic incorporation of bifunctional fatty acid analogues and functionalization using click chemistry. **(C)** Lipidomics analysis of functionalized probes (50 µM for 4 h in HeLa) showing incorporation across major lipid classes. Absolute concentrations (µg/mL) were quantified using class-specific deuterated internal standards (n = 3; mean ± s.d.). **(D)** Heat map comparing the relative incorporation of each probe into lipid species, normalized by z-score across treatments. **(E)** Confocal imaging of bifunctional fatty acid analogues in live cells after 4 hour incubation and subsequent SPAAC labeling with BODIPY-BCN (CO-1^38^). Images were taken with and without co-treatment of oleic acid to induce lipid droplet formation. Scale bar represents 20 μm. **(F)** Fluorescence intensities after labeling of bifunctional lipids (n = 3; mean ± s.d.). **(G)** Pearson correlation coefficients between the fluorescent signal of each probe and markers for different cellular organelles (n = 12; mean ± s.d.; *p<0.05, **p<0.01, ***p<0.001, ****p<0.0001).

## Conclusion

In this work, we show that lipid tail structure is a programmable determinant of metabolic fate and that this principle can be leveraged to selectively route substrates through endogenous lipid metabolic networks. By systematically modifying central lipid metabolites and quantifying their downstream incorporation, we demonstrate that chemical variation of fatty acids, alkylglycerols, and sphingosines modulates metabolic flux and generates distinct lipid profiles. Fatty acid tail architecture strongly influenced incorporation into phospholipids, neutral lipids, lysolipids, and acylcarnitines, and modified branched and aromatic fatty acids produced strikingly selective routing patterns, enabling preferential enrichment of neutral lipids or defined phospholipid subsets. Modified alkylglycerols enabled selective routing into ether lipids and chemical modifications lead to distinct species distributions. Chemically modified sphingosines provided selective access to distinct sphingolipid classes, with some analogs preferentially routed into sphingomyelin and others into both sphingomyelin and complex glycosphingolipids.

We further couple metabolic selectivity to chemical functionalization. Bifunctional lipid analogs bearing a routing element and a bioorthogonal handle enable delivery of chemical functionality to defined lipid pools in living cells, allowing selective installation of probes, such as fluorophores or other functional groups, into newly synthesized cellular lipids. Together, our findings establish substrate-directed metabolic control as a broad strategy for selectively modulating the mammalian lipidome. Precision control of the lipidome expands our ability to interrogate lipid function and may enable selective lipidome reprogramming in disease.

## Synthetic Chemistry

### General Practice

Unless otherwise noted, all non-aqueous reactions were carried out under Nitrogen atmosphere, in oven dried glassware. Reagents were purchased from commercial suppliers (ABCR, ACROS, Sigma Aldrich, Ambeed, TCI, Strem, Alfa, Combi-Blocks or Fluorochem) and used without further purification. Anhydrous solvents over molecular sieves were purchased from Acros and used as received. Analytical thin layer chromatography (TLC) was performed on Merck silica gel 60 F254 TLC glass plates and visualized with 254 nm light and potassium permanganate (1.50 g KMnO_4_, 10.0 g K_2_CO_3_, 1.25 mL 10% NaOH, 200 mL water), or ceric ammonium molybdate (10.0 g Cerium(IV)sulfate, 25.0 g phosphomolybdic acid, 940 mL water 60.0 mL conc. sulfuric acid) staining solutions followed by heating. Organic solutions were concentrated by rotary evaporation at 40 °C. Chromatographic purification of reaction products was carried out by flash chromatography using Millipore Silica gel 60 (0.063-0.200 mm), under 0.3–0.5 bar overpressure.

### NMR Spectroscopy

^1^H NMR spectra were recorded on a Bruker AVIII 400 MHz spectrometer with a prodigy cryoprobe and are reported in ppm with the solvent resonance as the reference (CDCl_3_ at 7.26 ppm, MeOH-d_4_ at 3.31 ppm, DMSO-d_6_ at 2.50). Peaks and their apparent multiplicities are reported as (s = singlet, d = doublet, t = triplet, q = quartet, m = multiplet, br = broad signal, coupling constant(s) in Hz, integration). ^13^C NMR spectra were recorded with 1H-decoupling on Bruker AVIII 101 MHz spectrometer prodigy cryoprobe and are reported in ppm with the solvent resonance as the reference unless noted otherwise (CDCl_3_ at 77.16 ppm, MeOH-d_4_ at 49.00 ppm DMSO-d_6_ at 39.52 ppm).

### HRMS

High resolution mass spectrometric data were obtained on an Agilent 6230 LC-TOF and are reported as (*m/z*).

### Cell culture

All cell lines, including HeLa, A549, Caco-2, and HepG2 cells, were cultured in T75 flasks (Fisher Scientific, FB012937) in Dulbecco’s Modified Eagle Medium (DMEM; high glucose, sodium pyruvate, and L-glutamine; GenClone, 25-500) supplemented with 10% (v/v) fetal bovine serum (FBS; GenClone, 25-550) and 1% (v/v) penicillin–streptomycin (P/S). HeLa SK1/2 KO were previously described.^40^

## Lipidomic Analysis

### Materials

Materials used for LC–MS were water, Optima™ LC/MS Grade (Fisher Scientific, W6-4), acetonitrile, Optima™ LC/MS Grade (Fisher Scientific, A955-4), Isopropanol, Optima™ LC/MS Grade (Fisher Scientific, A461-A), ammonium formate (Sigma-Aldrich, 70221-25G-F) and formic acid Optima™ LC/MS Grade (Fisher Scientific, A117-50). Solvents for lipid extraction were tert-butyl methyl ether (Sigma-Aldrich, 34875) and methanol (Fisher Scientific, A456). The lipid internal standard mixture was Deuterated Lipidomics MaxSpec ® Mixture (Cayman Chemical, 40974).

### Lipid extraction

Lipid extraction was performed using a tert-butyl methyl ether (MTBE)-based protocol.^41^ Cells were washed twice with ice-cold PBS, harvested from 6-well plates, and pelleted at 800 × g for 5 min at 4 °C. Pellets were resuspended in 70 µL water, followed by addition of 232 µL methanol and 20 µL (10× diluted) internal standard mixture. Subsequently, 840 µL MTBE was added and samples were incubated for 1 h at room temperature with shaking. Phase separation was induced by adding 150 µL water, followed by incubation for 5 min at room temperature and centrifugation at 1,000 × g for 10 min. The upper organic phase (750 µL) was collected and transferred to glass autosampler vials (Thermo, 6PSV9-1PG) and dried overnight under vacuum.

### Liquid Chromatography

Dried lipid extracts were reconstituted in 500 µL of Solvent B (isopropanol/acetonitrile/ water, 88:10:2, vol/vol) containing 10 mM ammonium formate and 0.1% (vol/vol) formic acid. Lipids were separated by reversed-phase LC on a Vanquish Core system (Thermo Fisher Scientific) equipped with an Accucore C30 column (150 × 2.1 mm, 2.6 mm, 150 Å, Thermo Fisher Scientific). Mobile phase A from LC system was acetonitrile/water, 60:40, vol/vol with 10 mM ammonium formate and 0.1% (vol/vol) formic acid and Mobile phase B from LC system was isopropanol/acetonitrile/water, 88:10:2, vol/vol both containing 10 mM ammonium formate and 0.1% (vol/vol) formic acid. Separation was performed at 45 °C at a flow rate of 0.26 mL/min using the following gradient: 0–2 min, 30–43% B (curve 5); 2.1–12 min, 43–55% B (curve 5); 12–18 min, 65–85% B (curve 5); 18–20 min, 85–100% B (curve 5); 20–25 min, 100% B isocratic; 25.1–28 min, 100%–30% B (curve 5) followed by 7 min re-equilibration at 30% B.

### Mass spectrometry

LC was coupled to an Orbitrap Exploris™ 240 mass spectrometer (Thermo Fisher Scientific) with a heated electrospray ionization (HESI) source. Mass spectra were acquired in positive and negative modes with the following ESI parameters: sheath gas, 40 (Arb); auxiliary gas, 10 (Arb); sweep gas, 0 (Arb); spray voltage, (+)3.25 kV (positive ion mode); (−)3 kV (negative ion mode); Ion transfer tube temperature, 300 °C; S-lens RF level, 70; Vaporizer temperature, 275 °C. Data acquisition for lipid identification was performed in data-dependent acquisition mode (DDA) full scan with/without MS/MS. The full scan has the resolution of 120,000, AGC target set to standard mode, maximum injection time set to auto mode in a scan range of m/z 100–1,200. Data-dependent MS/MS scans were acquired with a resolution of 15,000, AGC target set to standard mode, maximum injection time set to auto mode, isolation window of 1 m/z and stepped normalized collision energies of 20, 30 and 40. A data-dependent MS2 was triggered (Cycle time of 1.2 s) when an AGC target of 2.5e3 was reached followed by a dynamic exclusion for 10 s. All isotopes and charge states >1 were excluded. Full scan spectra were acquired in profile type and MS/MS scans were acquired in centroid type. Data for lipid quantification in individual samples were acquired in full scan MS mode with the following parameters: resolution of 120,000, AGC target set to standard mode, maximum injection time set to auto mode in a scan range of m/z 100-1200.

### Lipid identification and quantification

Raw data were processed using Compound Discoverer v3.4 (Thermo Scientific) with a modified MetID Mass List Search workflow using the following parameters: m/z tolerance for precursor mass selection set to 5 ppm; minimum peak intensity threshold 30,000; S/N threshold set to 1.5; Peak rating threshold set to 0.4; Gap fill enabled with a 1.5 S/N threshold, 3 minimum scans per peak. A mass list having exact masses of both endogenous lipids and chemically modified lipids was imported for analysis. For lipid identification, triacylgylcerols, diacylglycerols were identified as [M + NH_4_]^+^ adducts. Lysophosphatidylcholines, and acyl-, ether- and vinyl ether-PC, acylcarnitines, ceramides and sphingomyelins were analyzed as [M + H]^+^ adducts. Lyso-phosphatidylethanolamines, and acyl-, ether- and vinyl ether-PE, phosphatidylserines, phosphatidylinositols and phosphatidylglycerol were analyzed as [M−H]^-^ adducts. Acyl-, ether- and vinyl ethers were identified as [M−H]^-^ adducts.^42^ Ether lipids and plasmalogens were identified based on MS/MS fragmentation, where diagnostic sn2 acyl chain fragments were used to determined fatty acyl composition. The lipid classes BMP and PG are isomeric and could not be distinguished with high confidence under the conditions used. Quantification was performed by peak integration of the extracted ion chromatograms of most common ion adducts. Peak integration was first performed by CD then manually curated and adjusted. Identified lipids were normalized to peak areas of added internal standards to decrease analytical variation. Raw lipidomics data were analyzed using Compound Discoverer and exported as .csv files and plotted in GraphPad Prism 11.0.0 (GraphPad Software).

### Lipid Droplet Imaging and Quantification

HeLa cells were seeded in 18-well Cellvis 18 Chambered Coverglass System (C18-1.5H), coated with poly-L-lysine, at the density of 15k cells/well for 12 hours prior to treatment in DMEM (10% FBS, 1% P/S). Cells were incubated with DMEM (10% FBS, 1% P/S) containing 50 μM of π-FA1 and π-FA3 respectively for 6 hours. After incubation, the cells were washed with warm PBS and fixed with 4% (w/v) PFA for 8 minutes. Lipid droplets were stained with 1 μM BODIPY 493/503 at 37°C for 20 minutes and the nuclei were stained with 1 μg/mL Hoechst 33342 at 37°C for 15 minutes. Prior to imaging, cells were washed with PBS. The fixed cells were imaged using Zeiss LSM800 with a 40X water objective. Acquired images were saved as .Tif files for analysis. All images were processed with Fiji^43^ (ImageJ v2.16.0/1.54p). For each channel in the representative fluorescence microscopy image, the images are first converted to 8-bit format and adjusted the brightness (min:32, max:255). The merged images were generated by merging the figure using processed images obtained from the DAPI and BODIPY channel. For nuclei quantification, the original image from the DAPI channel were converted into 8-bit format and set with adjusted threshold (min:46, max:255). The function, find edge, was applied prior to using analyze particle function (size: 500-infinity pixel) to quantify the number of nuclei. For LD quantification, the original image from BODFI channel were converted into 8-bit format and set with adjusted threshold (min: 70, max: 255). The image is converted to mask and quantified using the analyze particle function (size: 0-Infinity, circularity: 0.20-1.00 pixel). The LDs/cell was calculated by dividing the number of LDs in each image by the number of nuclei in each image. The data was plotted using GraphPad Prism. Unpaired t-test with Welch’s correction was used to determine if the difference between treatments were significant.

### SPAAC Imaging

HeLa cells were seeded in 8-well chamber with glass imaging slide (Cellvis #C8-1.5H-N) 24h before imaging experiment in DMEM (10% FBS, 1% P/S), such that cells grow to approximately 70% confluency at the time of imaging. The fatty acid (clSA, cl-2MeSA, cl-10MeSA) as a DMSO stock was diluted to a final concentration of 10 µM in warm DMEM (10% FBS, 1% P/S) with vehicle (DMSO) or oleic acid (40 µM) and a final concentration of DMSO of 0.5%, 200 µL of this solution was added to cells and cells incubated for 4 h at 37 °C and 5% CO_2_ (v/v). At this time, media is removed and replaced with 200 µL of 5 µM CO-1 in full media.^36^ The cells were incubated for 1h at 37°C and 5% CO2 (v/v). Media is then removed and replaced with 200 µL of 1 mg/mL Hoechst-33342 in phenol red-free DMEM media (10% FBS, 1% P/S) for 20 min or, for co-staining experiments, the organelle tracking dyes were used at the following concentration diluted in phenol red-free DMEM media (without FBS or P/S) and incubated for 10 min, BioTracker 405 Blue Mitochondria Dye (Millipore Sigma #SCT135, 100 nM) and CellMask™ Plasma Membrane Deep Red (Thermo Fisher C10046, 2 μg/mL) or ER-Tracker red (MCE #HY-D1431, 2 μM) and LipidSpotTM 610 (Biotium #70069-T, 1/1000 dilution). Cells were washed once with phenol red free DMEM media (10% FBS, 1% P/S) and replaced with fresh phenol red free DMEM media (10% FBS, + 1% P/S) for imaging. Cells were imaged with a Zeiss LSM 800, AxioObserver Z1 Inverted with Definite focus confocal microscope equipped with an incubation chamber, 405 nm, 488 nm, 561 nm and 633 nm excitation laser lines, GaAsP detectors, and a 63x oil immersion objective. In the image processing, FIJI was used to calculate fluorescence intensity for images of each probe. Cell regions to be quantified were selected by setting a threshold based after a Gaussian blur filter (sigma=1) was applied. Then “Create Mask” and “Create Selection” functions were used to select the area to be quantified based on the determined threshold. The mean fluorescence was then quantified for an image, which represented n=1, an ordinary one-way ANOVA was the statistical method used to compare the means of the treatment groups. For colocalization, the coloc2 plugin with FIJI was used to calculate Pearson’s correlation coefficient, where correlation is calculated for individual cells by drawing masks around individual cells, n=12 cells quantified per treatment condition, an ordinary one-way ANOVA was the statistical method used to compare the means of the colocalization coefficients.

## Associated Content

Supporting information include synthetic procedures and NMR characterization data.

## Competing Interests

A provisional patent application related to this work has been filed by the California Institute of Technology, on which B.W., L.L., and J.M. are listed as inventors.

## Author Contributions

B.W., L.L. and J.M. conceived the project and designed the study. B.W., L.L., L.T., L.Q., conducted experiments. T.H. and K.E. contributed to lipidomic analysis. B.W. and J.M. wrote the manuscript. All authors edited the manuscript.

## Acknowledgments

J.M. acknowledges the National Cancer Institute (NCI) for supporting this work (R00CA277358). L.L. thanks the Swiss National Science Foundation (SNSF) for a Postdoc.Mobility fellowship (P500-2_239150). The Biological Imaging Facility at Caltech is acknowledged for providing equipment. We thank Howard Riezman and Peter Dervan for helpful discussion of this study.

---

**Extended Figure S1.**
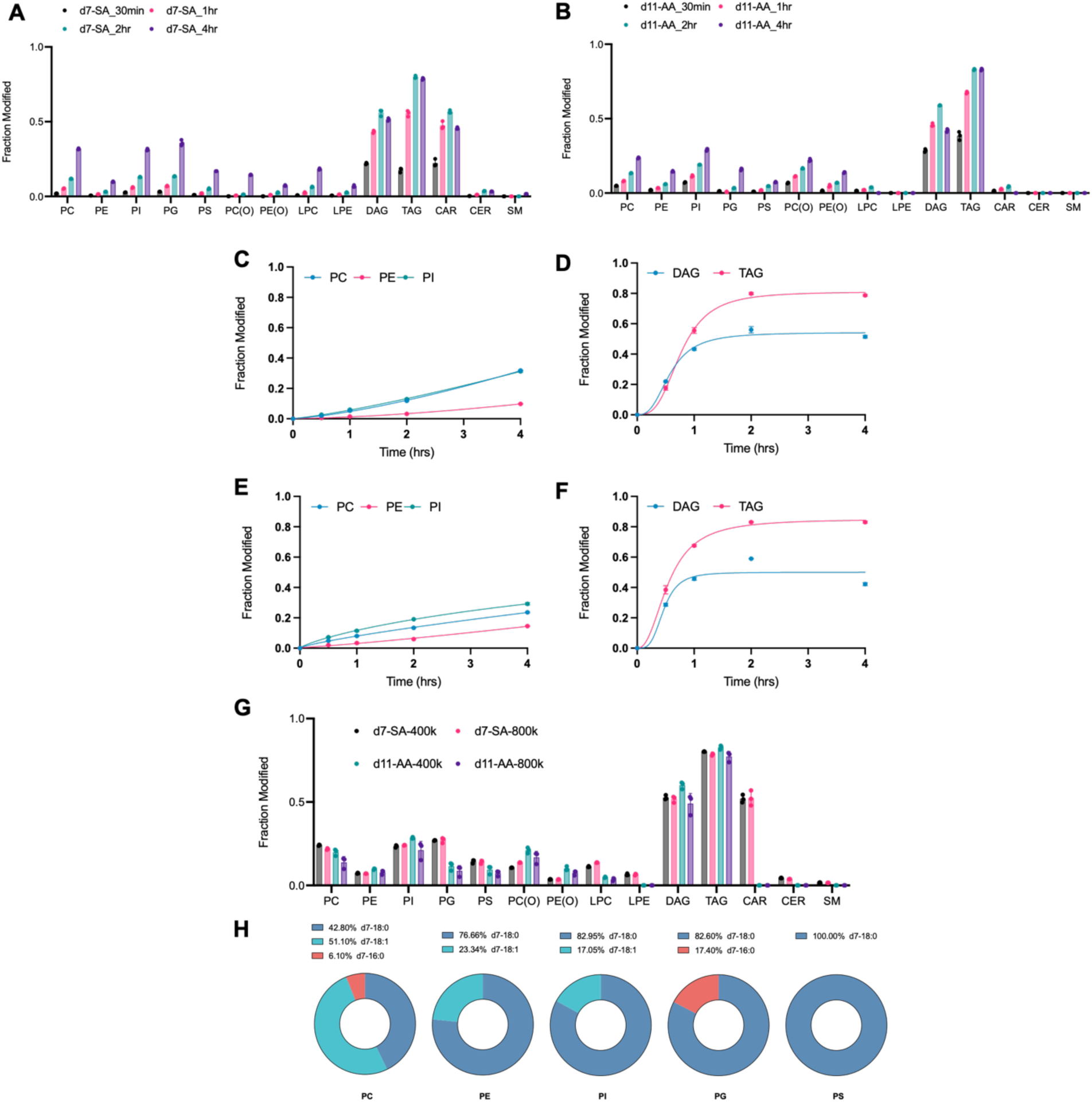
Expanded analysis of lipid tail-dependent metabolic routing. **(A)** Lipidomics analysis of incorporation of d7-stearic acid (d7-SA; 50 μM) into major lipid classes in HeLa cells at different treatment times, shown as the fraction of modified lipids relative to endogenous levels (n=3; mean ± s.d.).**(B)** Lipidomics analysis of incorporation of d11-arachidonic acid (d11-AA; 50 μM) into major lipid classes at different treatment times, shown as the fraction of modified lipids relative to endogenous levels (n = 3; mean ± s.d.). **(C, D)** Nonlinear regression analysis of d7-SA incorporation kinetics into phospholipids (C) and neutral lipids (D). **(E, F)** Nonlinear regression analysis of d11-AA incorporation kinetics into phospholipids (E) and neutral lipids (F). **(G)** Lipidomics analysis of d7-SA and d11-AA incorporation at different HeLa cell culture confluencies (50 μM, 4 h), shown as the fraction of modified lipids relative to endogenous levels (mean ± s.d., n = 3). **(H)** Lipidomics analysis of d7-SA-derived tail-modified species incorporated into major lipid classes in HeLa cells following 4 h treatment at 50 μM (n = 3, mean ± s.d.). Analysis was based on MS/MS acyl chain composition.

**Extended Figure S2.**
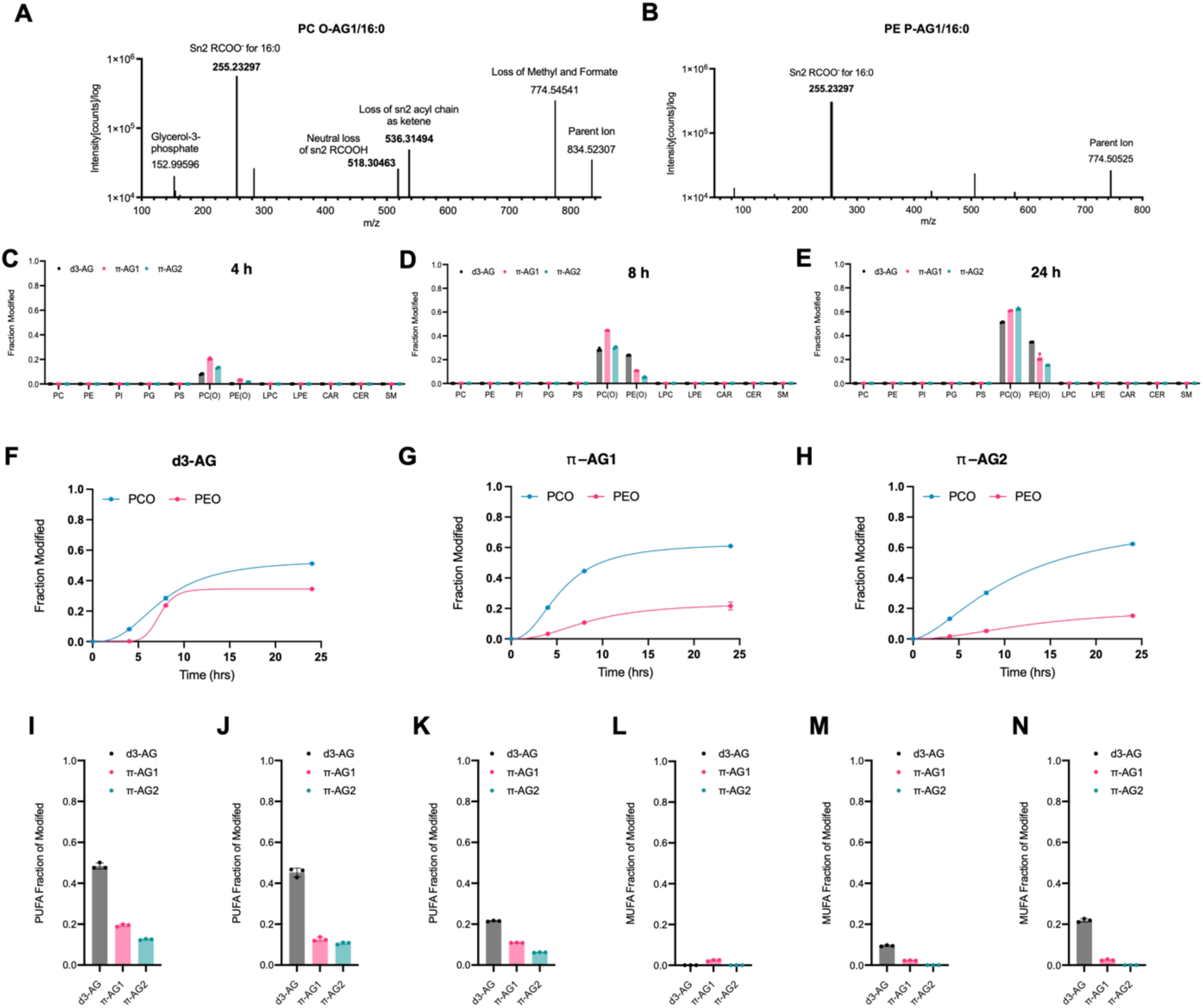
Expanded analysis of chemical ether lipid editing. **(A, B)** Representative MS/MS spectra of modified alkylglycerol-incorporated ether phosphatidylcholine (PC) (**A**) and ether phosphatidylethanolamine (PE) (**B**). **(C–E)** Lipidomics analysis of incorporation of chemically modified alkylglycerols (50 μM) into major lipid classes over time, shown as the fraction of modified lipids relative to endogenous levels (*n* = 3; mean ± s.d.). **(F–H)** Nonlinear regression analysis of alkylglycerol incorporation kinetics. **(I–K)** Fraction of polyunsaturated fatty acyl chains (PUFAs) in modified ether lipids after 4h **(I)**, 8h **(J)**, and 24h **(K)**. **(L–N)** Fraction of monounsaturated fatty acyl chains (MUFAs) in modified ether lipids after 4h **(L)**, 8h **(M)**, and 24h **(N)**.

**Extended Figure S3.**
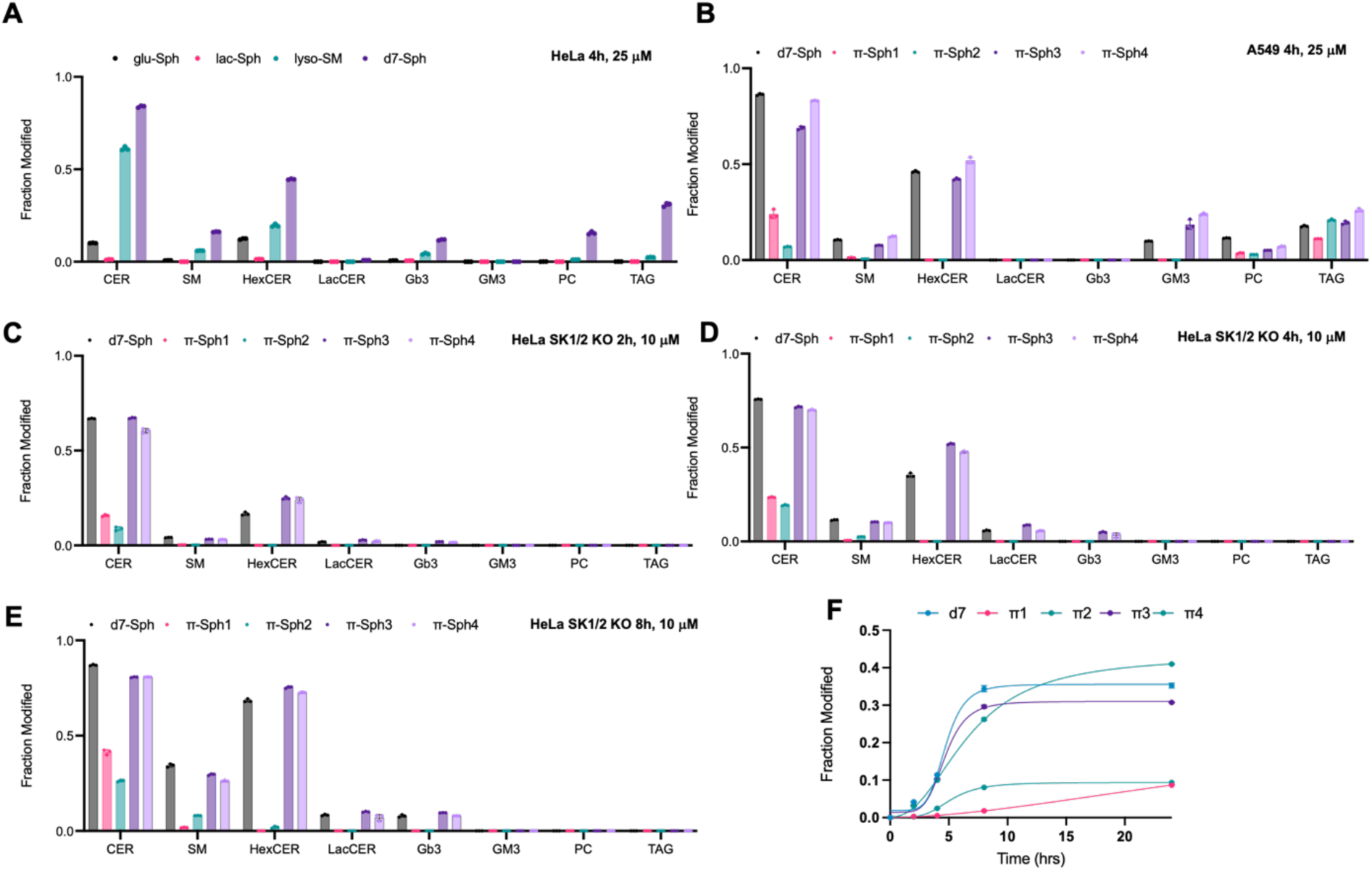
Expanded analysis of chemical sphingolipid editing. **(A).** Lipidomics analysis of incorporation of lyso-sphingolipids (25 μM, 4 h in HeLa cells) into major lipid classes, shown as the fraction of modified lipids relative to endogenous levels (*n* = 3; mean ± s.d.) **(B).** Lipidomics analysis of incorporation of aromatic sphingosines (25 μM, 4 h in A549 cells) into major lipid classes, shown as the fraction of modified lipids relative to endogenous levels (*n* = 3; mean ± s.d.). **(C-E).** Lipidomics analysis of incorporation of aromatic sphingosines (10 μM in HeLa cells) into major lipid classes across different treatment time, shown as the fraction of modified lipids relative to endogenous levels (*n* = 3; mean ± s.d.) **(F)** Nonlinear regression analysis of sphingomyelin (SM) incorporation kinetics.

**Extended Figure S4.**
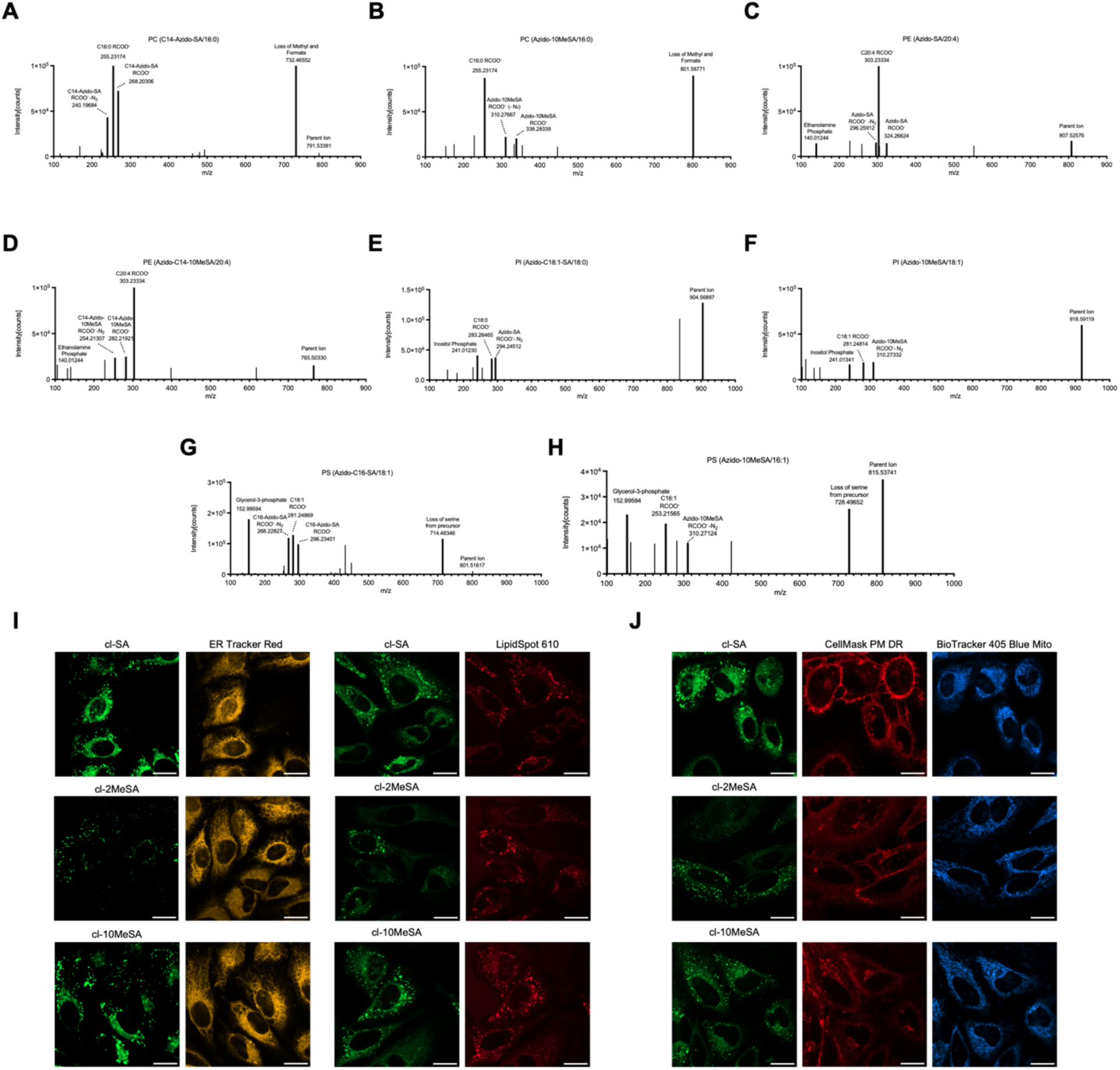
Expanded analysis of in situ lipid functionalization. **(A, B)** Confocal imaging of bifunctional fatty acid analogues (**cl-SA**, **cl-2MeSA**, and **cl-10MeSA**) in HeLa cells after 4 hour incubation and subsequent SPAAC labeling with BODIPY-BCN (CO-1^38^). Images were taken with co-treatment of oleic acid to induce lipid droplet formation. **(A)** Co-staining with LipidSpot 610 (1/1000 dilution) and ER-Tracker Red (1 μM). **(B)** Co-staining with CellMask Plasma Membrane Deep Red (2 μg/mL) and BioTracker 405 Blue Mitochondria (100 nM). **(C–J)** Representative MS/MS spectra of modified glycerophospholipids incorporating azido-functionalized fatty acids and branched analogues. **(C, D)** MS/MS spectra of modified PC species. **(E, F)** MS/MS spectra of modified PE species. **(G, H)** MS/MS spectra of modified PI species. **(I, J)** MS/MS spectra of PS species. Diagnostic fragment ions are highlighted to support lipid class identification and incorporation of modified acyl chains.

## Separation Workflow

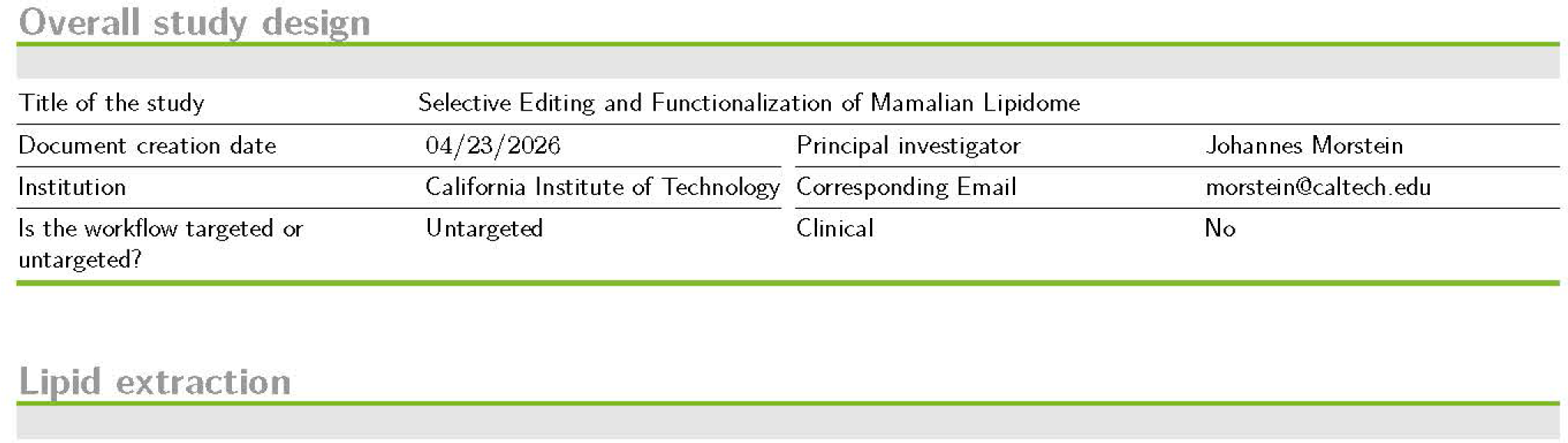
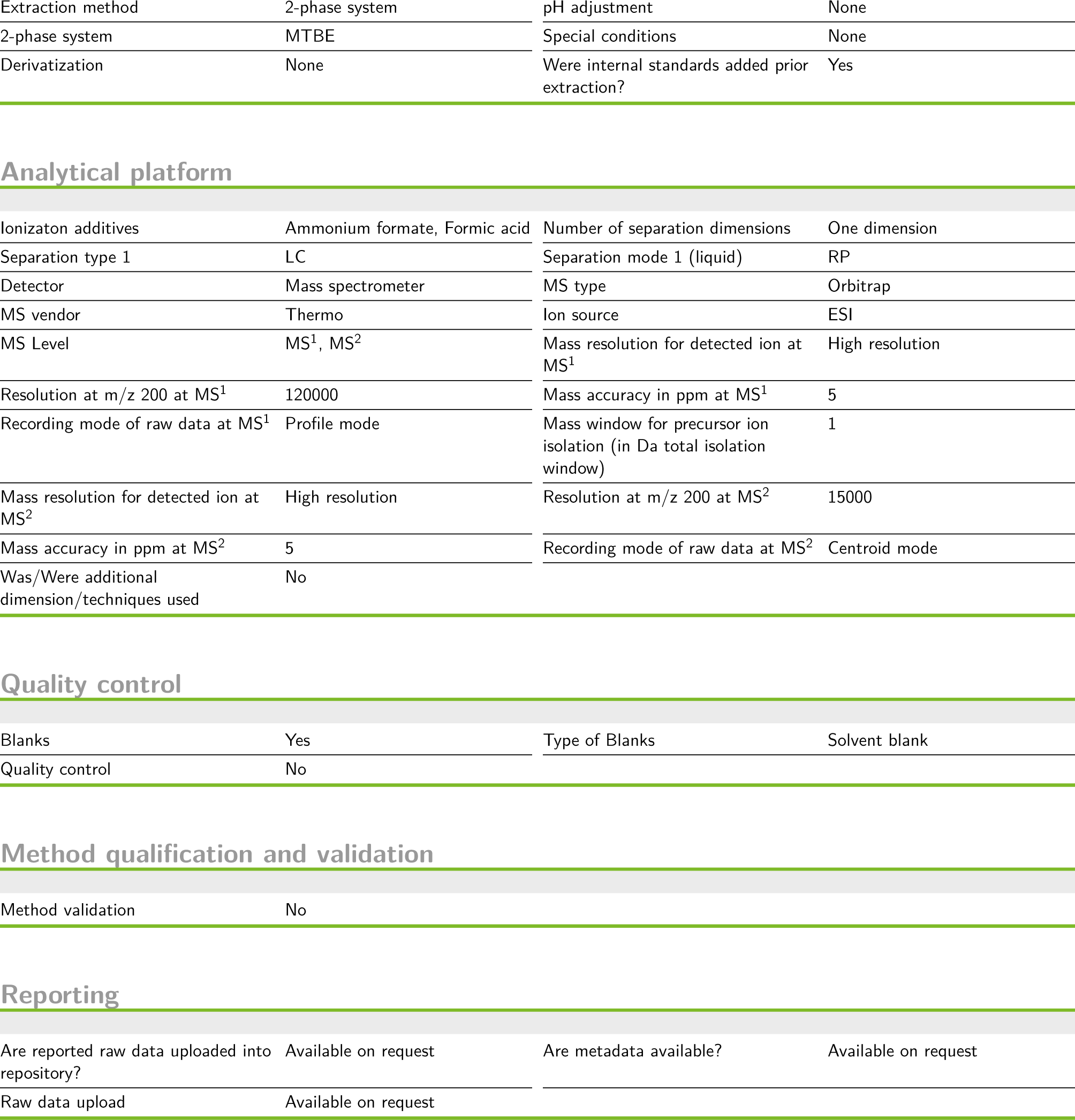

Overall study design
|  |  |  |  |
| --- | --- | --- | --- |
| Title of the study | Selective Editing and Functionalization of Mamalian Lipidome |  |  |
| Document creation date | 04/23/2026 | Principal investigator | Johannes Morstein |
| Institution | California Institute of Technology | Corresponding Email |  |
| Is the workflow targeted or untargeted? | Untargeted | Clinical | No |
| Lipid extraction |  |  |  |
| Extraction method | 2-phase system | pH adjustment | None |
| 2-phase system | MTBE | Special conditions | None |
| Derivatization | None | Were internal standards added prior extraction? | Yes |
| Analytical platform |  |  |  |
| Ionization additives | Ammonium formate, Formic acid | Number of separation dimensions | One dimension |
| Separation type 1 | LC | Separation mode 1 (liquid) | RP |
| Detector | Mass spectrometer | MS type | Orbitrap |
| MS vendor | Thermo | Ion source | ESI |
| MS Level | MS<sup>1</sup>, MS<sup>2</sup> | Mass resolution for detected ion at MS<sup>1</sup> | High resolution |
| Resolution at m/z 200 at MS<sup>1</sup> | 120000 | Mass accuracy in ppm at MS<sup>1</sup> | 5 |
| Recording mode of raw data at MS<sup>1</sup> | Profile mode | Mass window for precursor ion isolation (in Da total isolation window) | 1 |
| Mass resolution for detected ion at MS<sup>2</sup> | High resolution | Resolution at m/z 200 at MS<sup>2</sup> | 15000 |
| Mass accuracy in ppm at MS<sup>2</sup> | 5 | Recording mode of raw data at MS<sup>2</sup> | Centroid mode |
| Was/Were additional dimension/techniques used | No |  |  |
| Quality control |  |  |  |
| Blanks | Yes | Type of Blanks | Solvent blank |
| Quality control | No |  |  |

Method qualification and validation
|  |  |
| --- | --- |
| Method validation | No |
| Reporting |  |  |  |
| Are reported raw data uploaded into repository? | Available on request | Are metadata available? | Available on request |
| Raw data upload | Available on request |  |  |

## Sample Descriptions

epithelial cells / Human / Cells
|  |  |  |  |
| --- | --- | --- | --- |
| Storage and collection conditions | Available | Provided preanalytical information | Time to separate plasma/serum (min), Storage time (month), Freeze-thaw cycles, Preservation method |
| Temperature handling original sample | 4-8 °C | Instant sample preparation | Yes |
| Storage temperature | -80 °C | Storage time (month) | 0 |
| Freeze-thaw cycles | 0 | Additives | None |
| Were samples stored under inert gas? | No | Additional preservation methods | No |
| Biobank samples | No |  |  |

## Lipid Class Descriptions

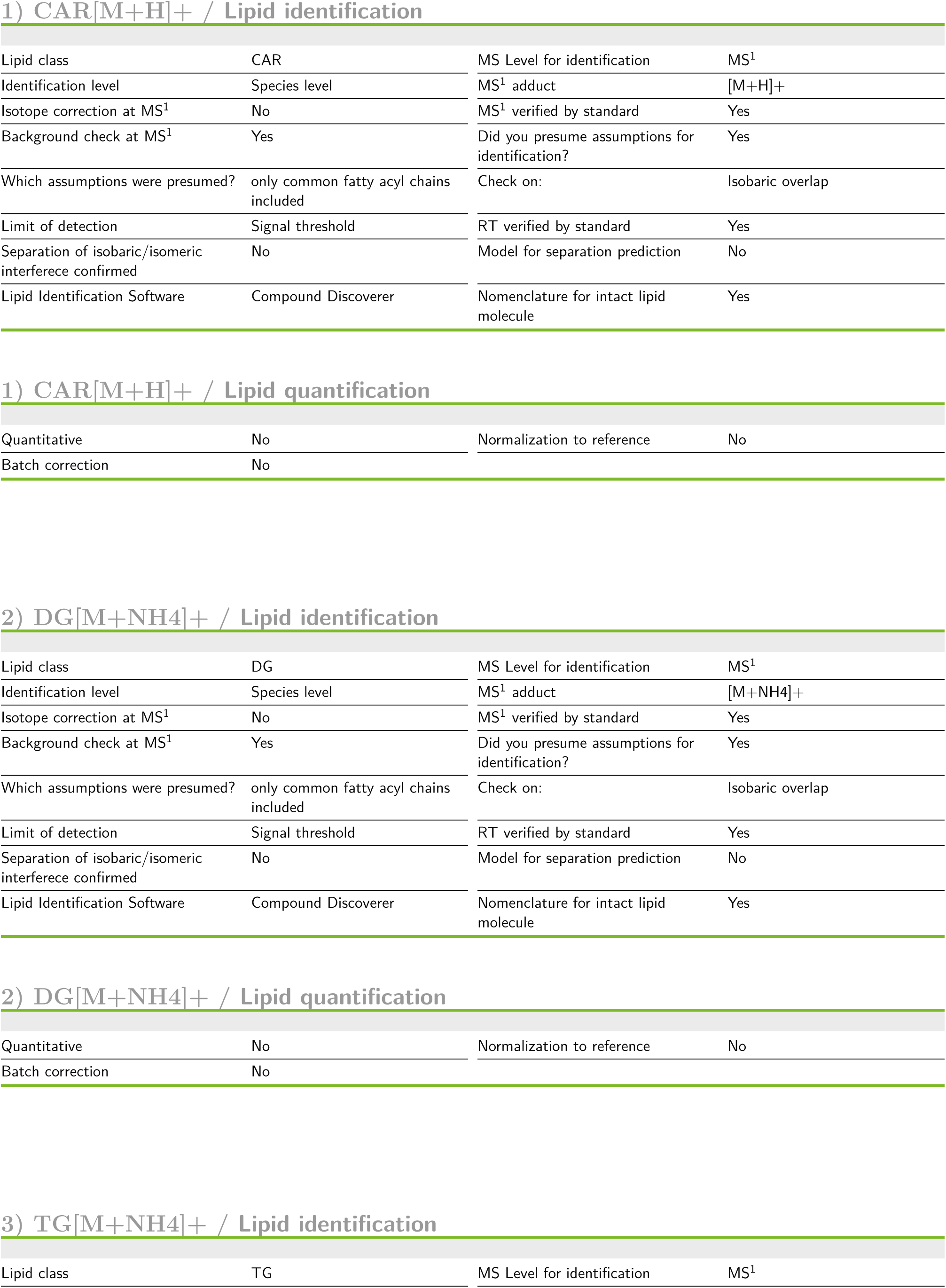
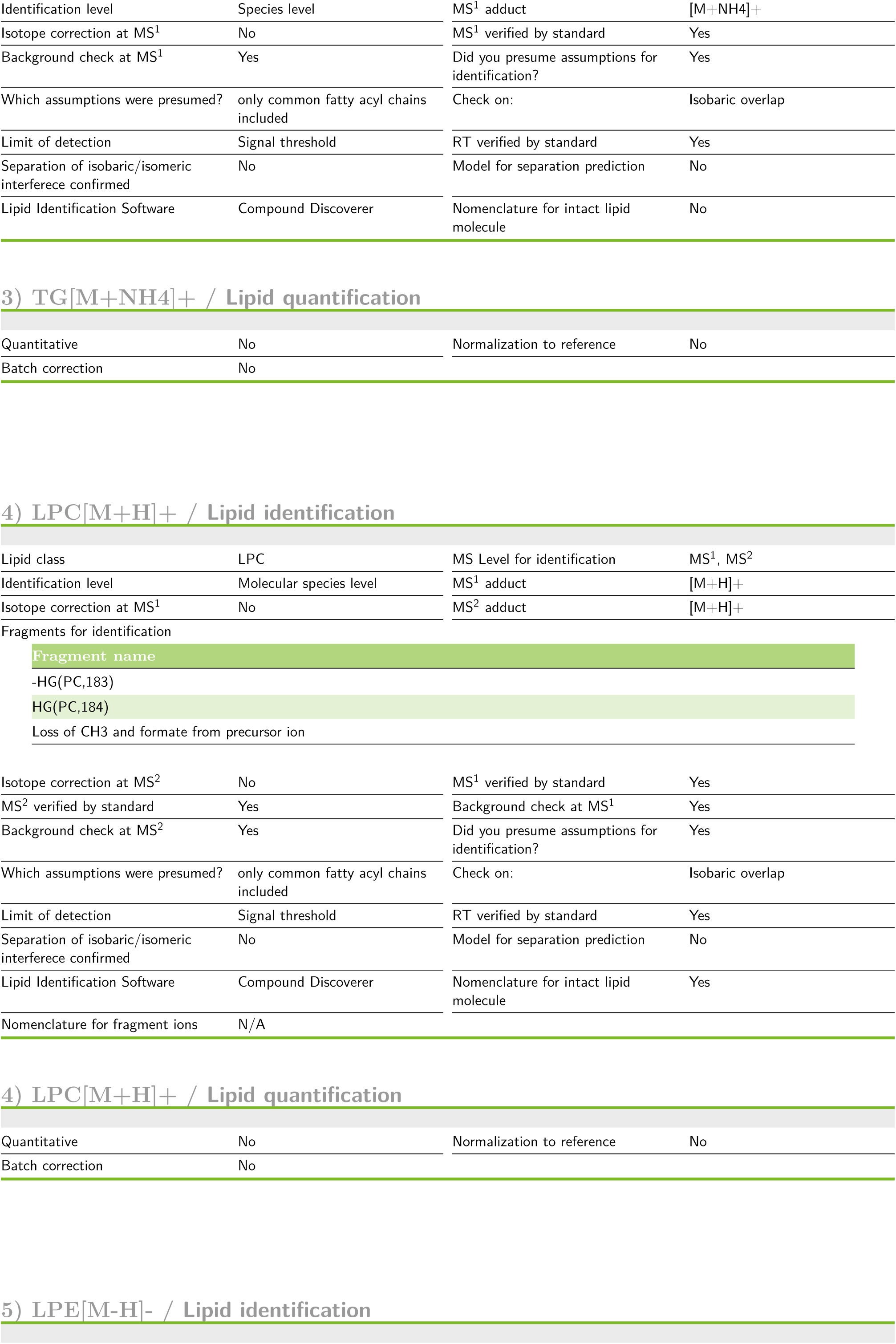
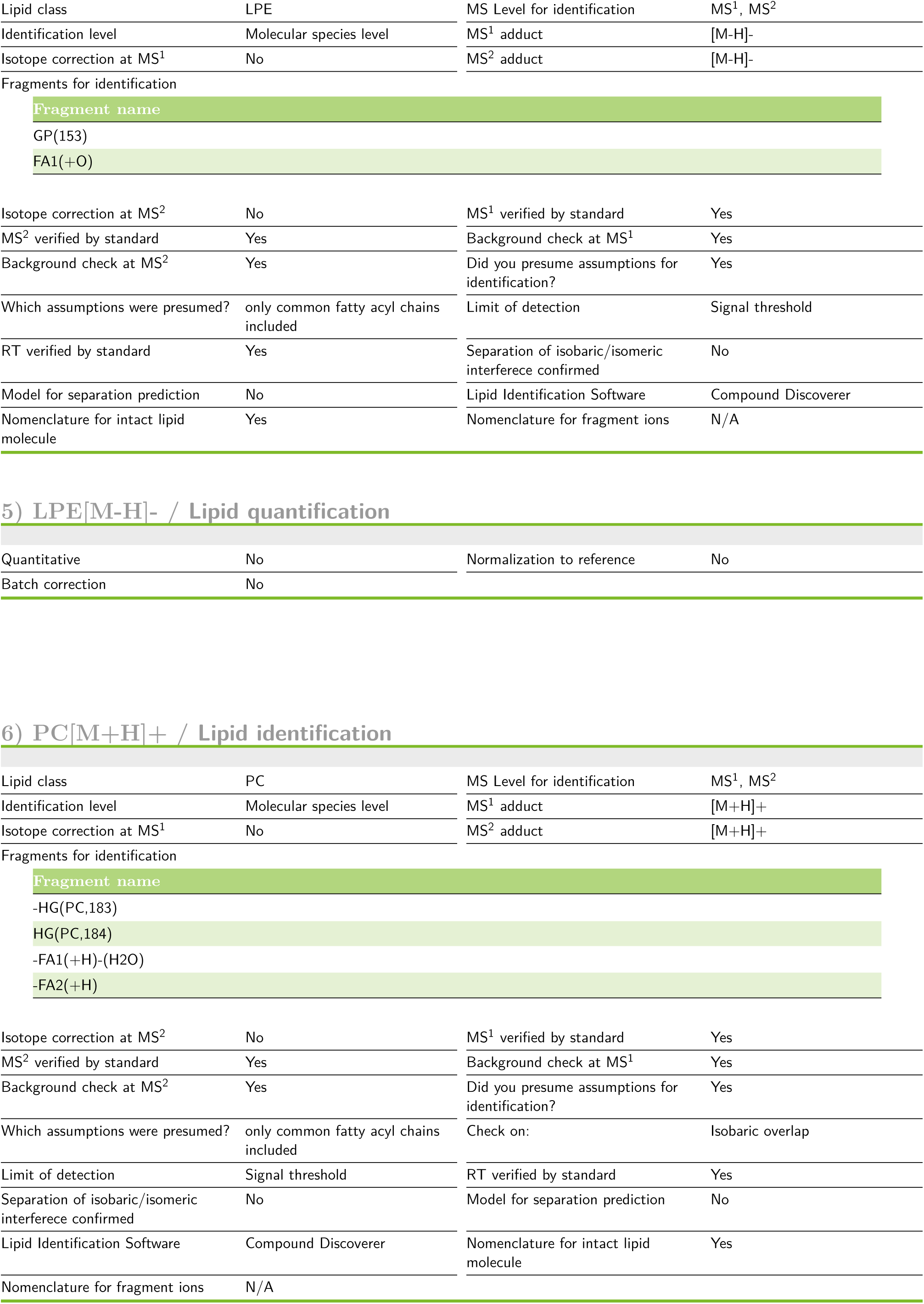
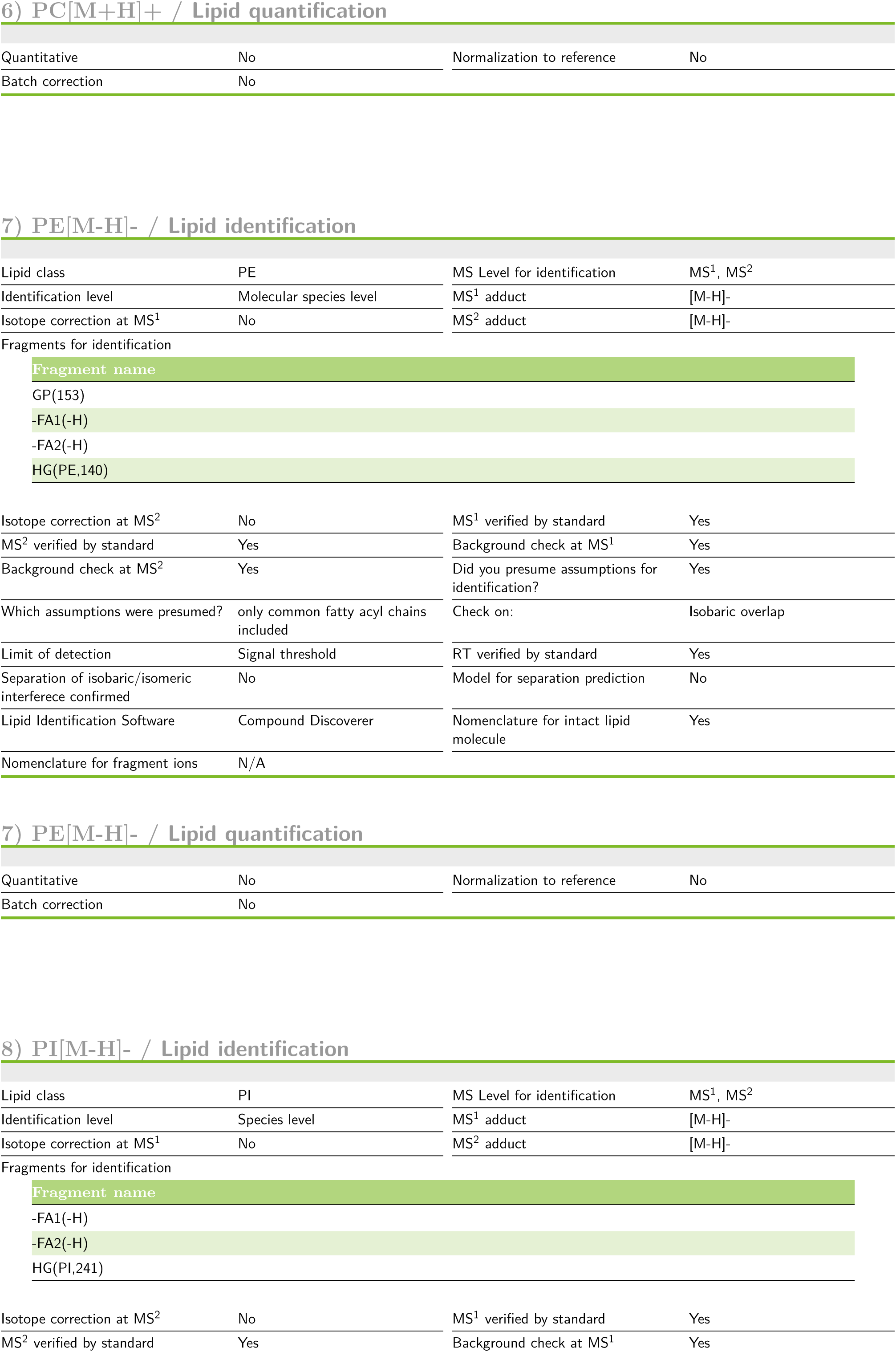
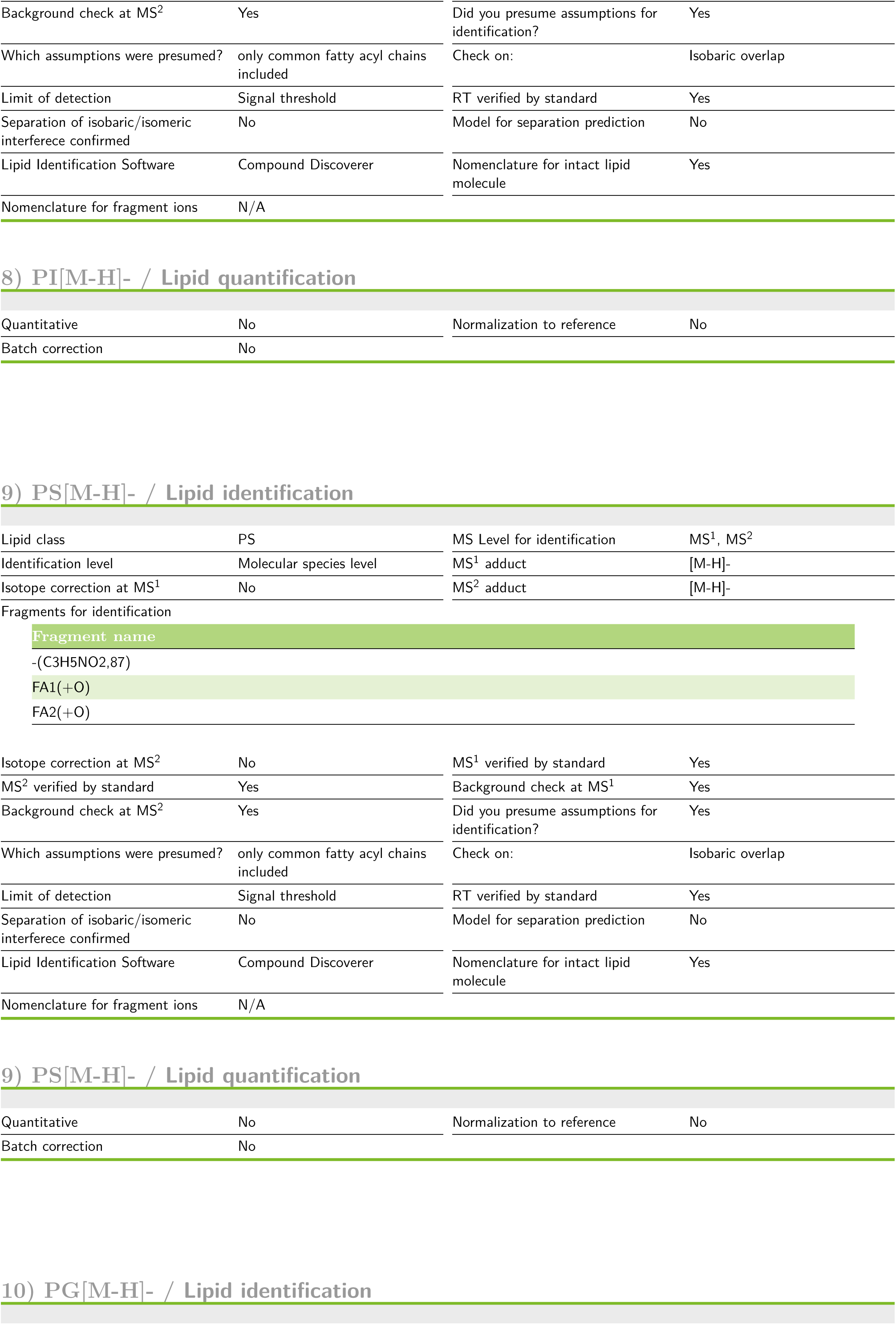
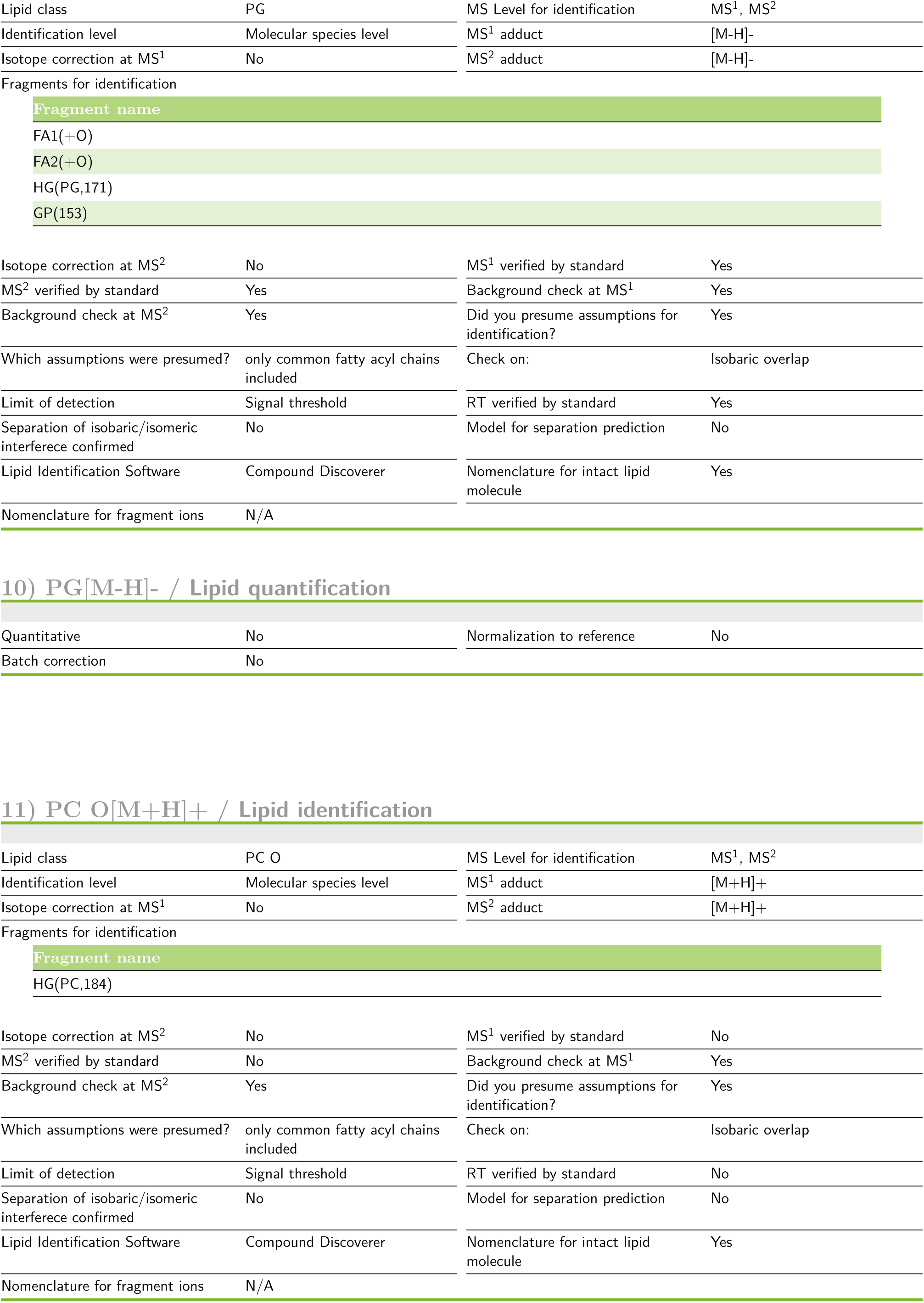
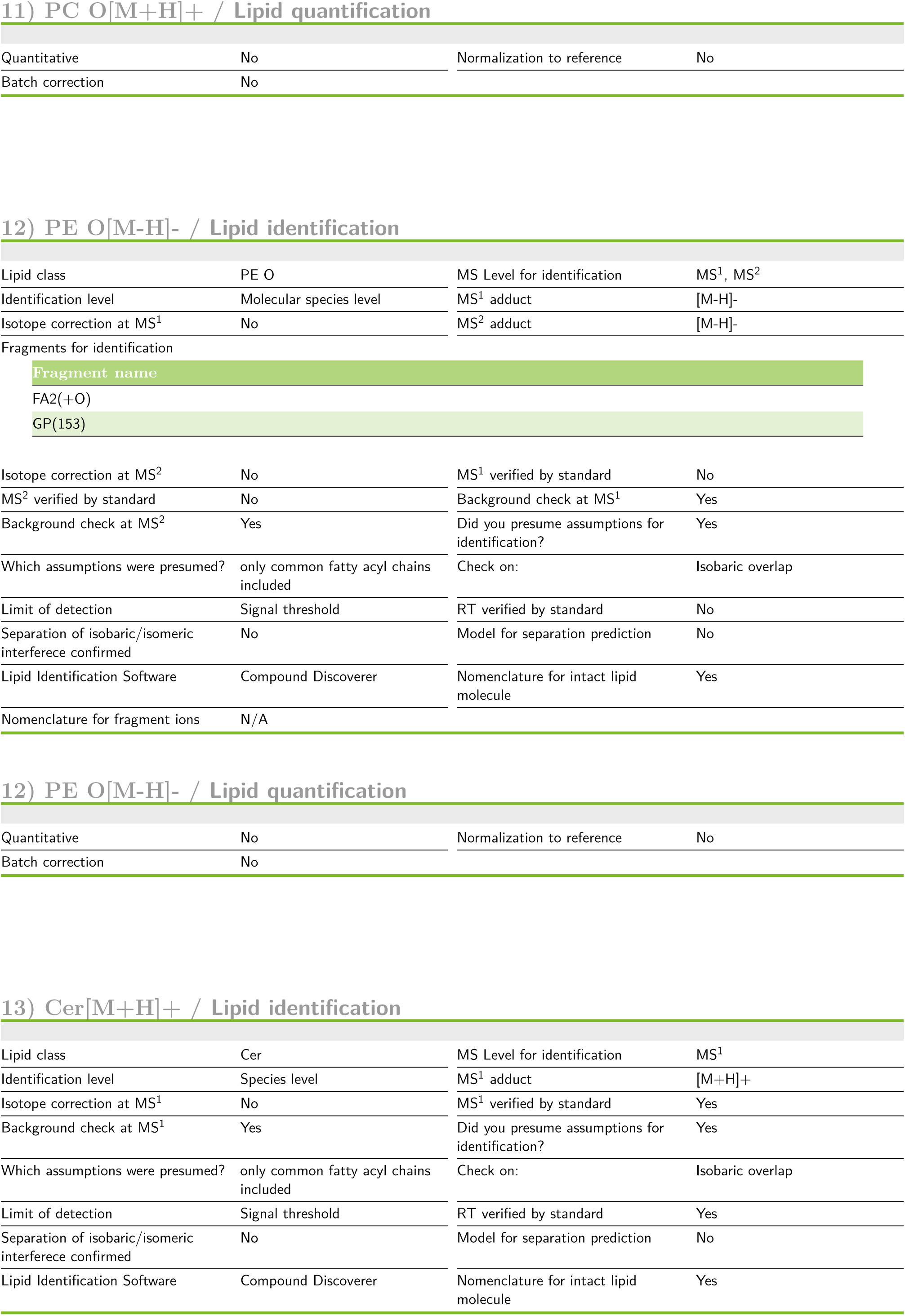
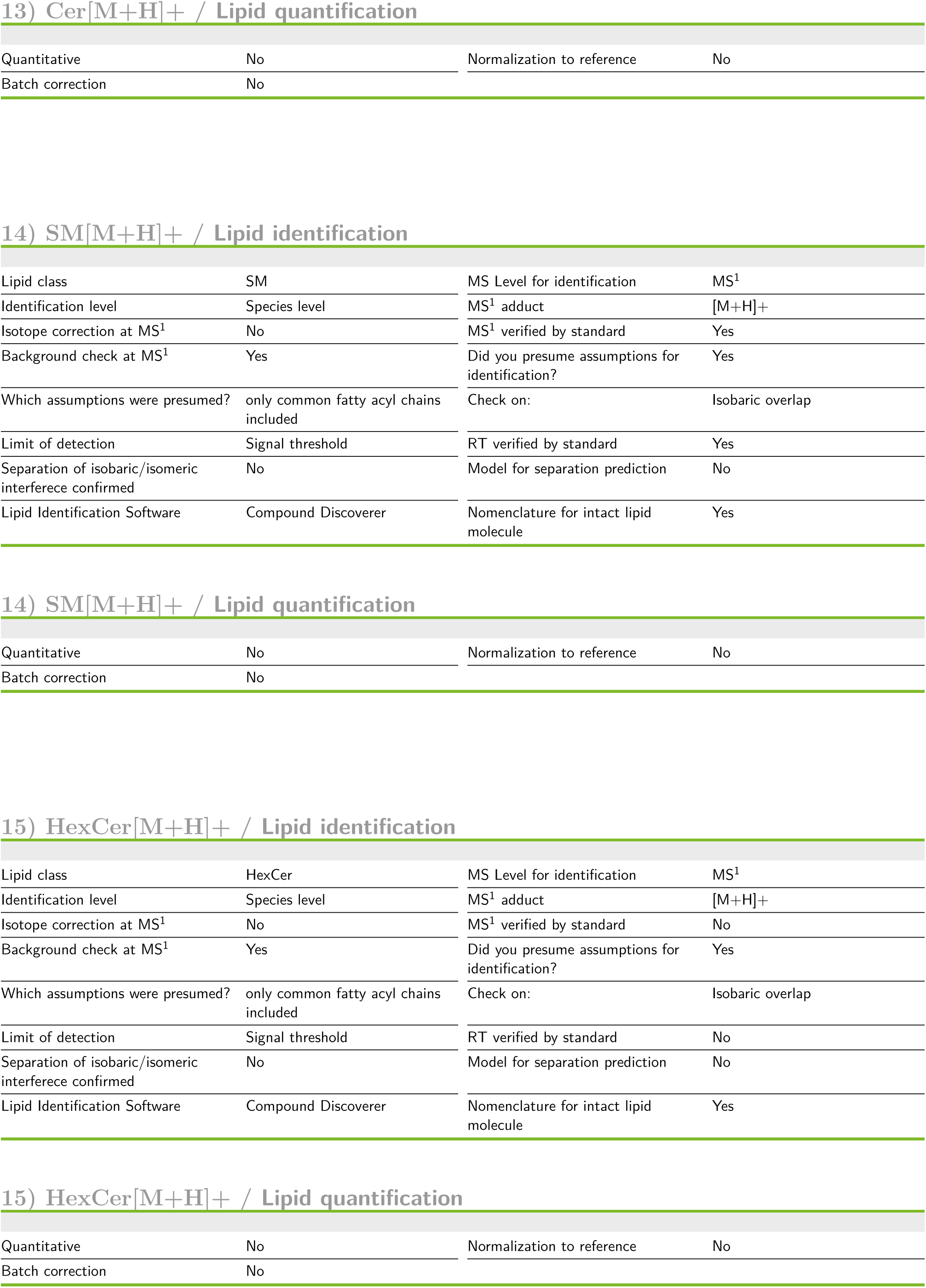
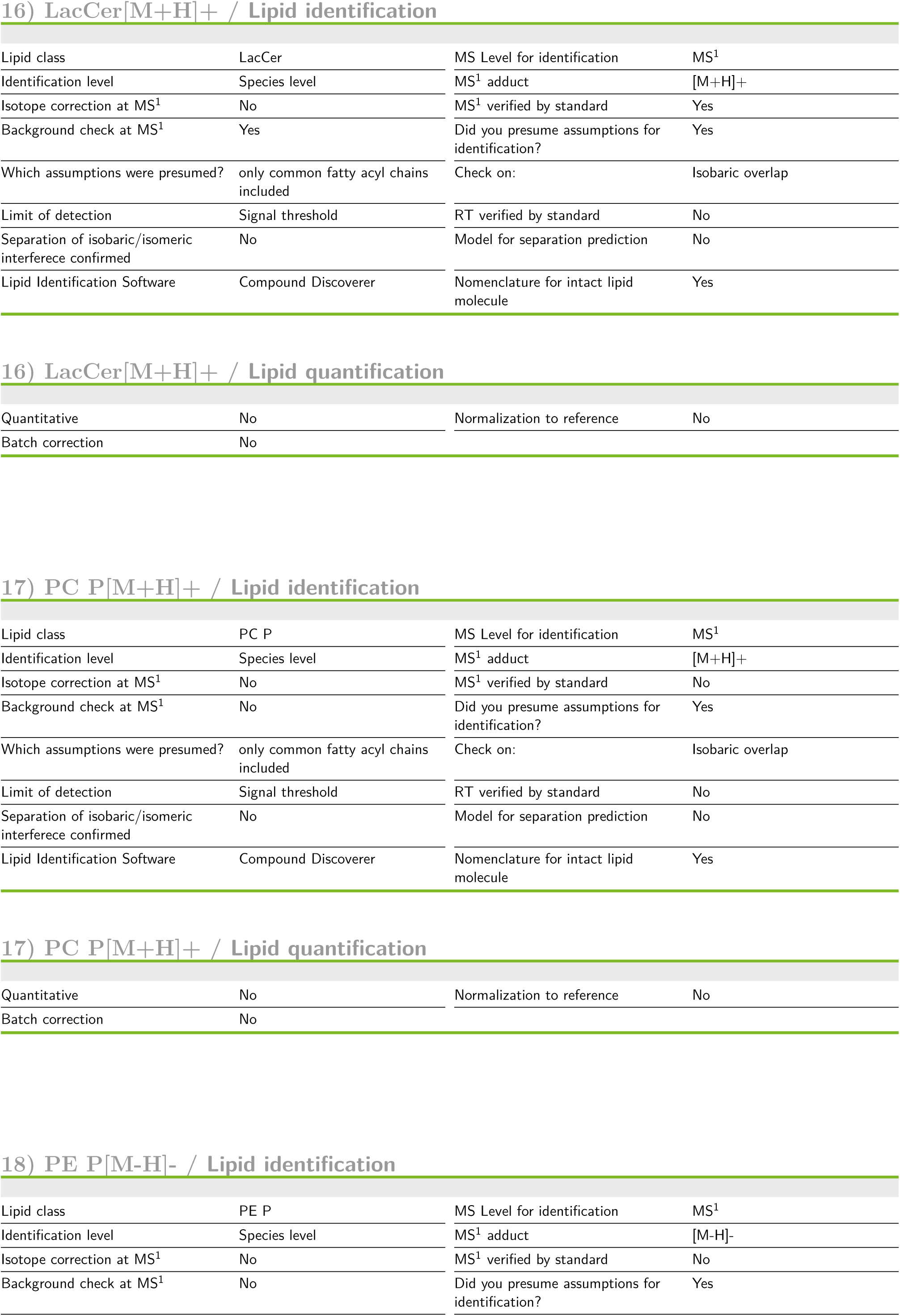
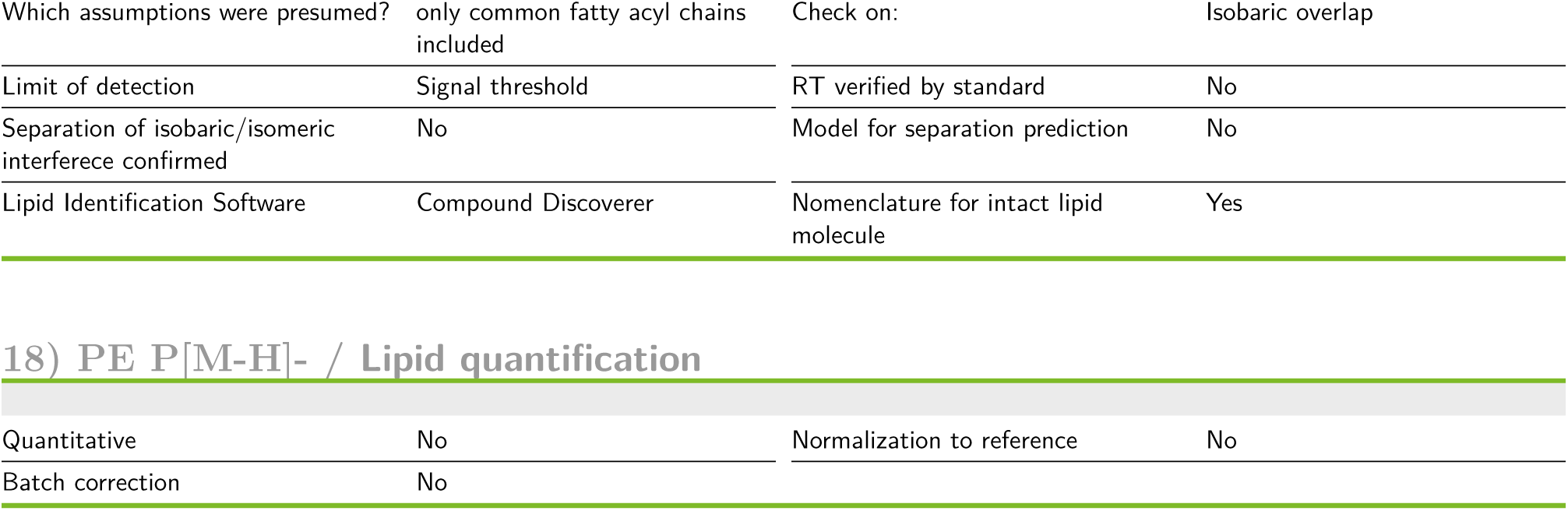

Lipid Class Descriptions 1) CAR[M+H]<sup>+</sup> / Lipid identification
| Lipid class | CAR | MS Level for identification | MS<sup>1</sup> |
| --- | --- | --- | --- |
| Identification level | Species level | MS<sup>1</sup> adduct | [M+H]<sup>+</sup> |
| Isotope correction at MS<sup>1</sup> | No | MS<sup>1</sup> verified by standard | Yes |
| Background check at MS<sup>1</sup> | Yes | Did you presume assumptions for identification? | Yes |
| Which assumptions were presumed? | only common fatty acyl chains included | Check on: | Isobaric overlap |
| Limit of detection | Signal threshold | RT verified by standard | Yes |
| Separation of isobaric/isomeric interference confirmed | No | Model for separation prediction | No |
| Lipid Identification Software | Compound Discoverer | Nomenclature for intact lipid molecule | Yes |
| 1) CAR[M+H]<sup>+</sup> / Lipid quantification |  |  |  |
| Quantitative | No | Normalization to reference | No |
| Batch correction | No |  |  |

2) DG[M+NH<sub>4</sub>]<sup>+</sup> / Lipid identification
| Lipid class | DG | MS Level for identification | MS<sup>1</sup> |
| --- | --- | --- | --- |
| Identification level | Species level | MS<sup>1</sup> adduct | [M+NH<sub>4</sub>]<sup>+</sup> |
| Isotope correction at MS<sup>1</sup> | No | MS<sup>1</sup> verified by standard | Yes |
| Background check at MS<sup>1</sup> | Yes | Did you presume assumptions for identification? | Yes |
| Which assumptions were presumed? | only common fatty acyl chains included | Check on: | Isobaric overlap |
| Limit of detection | Signal threshold | RT verified by standard | Yes |
| Separation of isobaric/isomeric interference confirmed | No | Model for separation prediction | No |
| Lipid Identification Software | Compound Discoverer | Nomenclature for intact lipid molecule | Yes |
| 2) DG[M+NH<sub>4</sub>]<sup>+</sup> / Lipid quantification |  |  |  |
| Quantitative | No | Normalization to reference | No |
| Batch correction | No |  |  |

3) TG[M+NH<sub>4</sub>]<sup>+</sup> / Lipid identification
| Lipid class | TG | MS Level for identification | MS<sup>1</sup> |
| --- | --- | --- | --- |
| Identification level | Species level | MS<sup>1</sup> adduct | [M+NH<sub>4</sub>]<sup>+</sup> |
| Isotope correction at MS<sup>1</sup> | No | MS<sup>1</sup> verified by standard | Yes |
| Background check at MS<sup>1</sup> | Yes | Did you presume assumptions for identification? | Yes |
| Which assumptions were presumed? | only common fatty acyl chains included | Check on: | Isobaric overlap |
| Limit of detection | Signal threshold | RT verified by standard | Yes |
| Separation of isobaric/isomeric interferece confirmed | No | Model for separation prediction | No |
| Lipid Identification Software | Compound Discoverer | Nomenclature for intact lipid molecule | No |
| 3) TG[M+NH<sub>4</sub>]<sup>+</sup> / Lipid quantification |  |  |  |
| Quantitative | No | Normalization to reference | No |
| Batch correction | No |  |  |
| 4) LPC[M+H]<sup>+</sup> / Lipid identification |  |  |  |
| Lipid class | LPC | MS Level for identification | MS<sup>1</sup>, MS<sup>2</sup> |
| Identification level | Molecular species level | MS<sup>1</sup> adduct | [M+H]<sup>+</sup> |
| Isotope correction at MS<sup>1</sup> | No | MS<sup>2</sup> adduct | [M+H]<sup>+</sup> |
| Isotope correction at MS<sup>2</sup> | No | MS<sup>1</sup> verified by standard | Yes |
| MS<sup>2</sup> verified by standard | Yes | Background check at MS<sup>1</sup> | Yes |
| Background check at MS<sup>2</sup> | Yes | Did you presume assumptions for identification? | Yes |
| Which assumptions were presumed? | only common fatty acyl chains included | Check on: | Isobaric overlap |
| Limit of detection | Signal threshold | RT verified by standard | Yes |
| Separation of isobaric/isomeric interferece confirmed | No | Model for separation prediction | No |
| Lipid Identification Software | Compound Discoverer | Nomenclature for intact lipid molecule | Yes |
| Nomenclature for fragment ions | N/A |  |  |
| 4) LPC[M+H]<sup>+</sup> / Lipid quantification |  |  |  |
| Quantitative | No | Normalization to reference | No |
| Batch correction | No |  |  |
| 5) LPE[M-H]<sup>-</sup> / Lipid identification |  |  |  |
| Lipid class | LPE | MS Level for identification | MS<sup>1</sup>, MS<sup>2</sup> |
| Identification level | Molecular species level | MS<sup>1</sup> adduct | [M-H]- |
| Isotope correction at MS<sup>1</sup> | No | MS<sup>2</sup> adduct | [M-H]- |
| Fragments for identification |  |  |  |
| Fragment name |  |  |  |
| GP(153) |  |  |  |
| FA1(+O) |  |  |  |
| Isotope correction at MS<sup>2</sup> | No | MS<sup>1</sup> verified by standard | Yes |
| MS<sup>2</sup> verified by standard | Yes | Background check at MS<sup>1</sup> | Yes |
| Background check at MS<sup>2</sup> | Yes | Did you presume assumptions for identification? | Yes |
| Which assumptions were presumed? | only common fatty acyl chains included | Limit of detection | Signal threshold |
| RT verified by standard | Yes | Separation of isobaric/isomeric interferece confirmed | No |
| Model for separation prediction | No | Lipid Identification Software | Compound Discoverer |
| Nomenclature for intact lipid molecule | Yes | Nomenclature for fragment ions | N/A |
| 5) LPE[M-H]- / Lipid quantification |  |  |  |
| Quantitative | No | Normalization to reference | No |
| Batch correction | No |  |  |
| 6) PC[M+H]<sup>+</sup> / Lipid identification |  |  |  |
| Lipid class | PC | MS Level for identification | MS<sup>1</sup>, MS<sup>2</sup> |
| Identification level | Molecular species level | MS<sup>1</sup> adduct | [M+H]<sup>+</sup> |
| Isotope correction at MS<sup>1</sup> | No | MS<sup>2</sup> adduct | [M+H]<sup>+</sup> |
| Fragments for identification |  |  |  |
| Fragment name |  |  |  |
| -HG(PC,183) |  |  |  |
| HG(PC,184) |  |  |  |
| -FA1(+H)-(H2O) |  |  |  |
| -FA2(+H) |  |  |  |
| Isotope correction at MS<sup>2</sup> | No | MS<sup>1</sup> verified by standard | Yes |
| MS<sup>2</sup> verified by standard | Yes | Background check at MS<sup>1</sup> | Yes |
| Background check at MS<sup>2</sup> | Yes | Did you presume assumptions for identification? | Yes |
| Which assumptions were presumed? | only common fatty acyl chains included | Check on: | Isobaric overlap |
| Limit of detection | Signal threshold | RT verified by standard | Yes |
| Separation of isobaric/isomeric interferece confirmed | No | Model for separation prediction | No |
| Lipid Identification Software | Compound Discoverer | Nomenclature for intact lipid molecule | Yes |
| Nomenclature for fragment ions | N/A |  |  |
| 6) PC[M+H]<sup>+</sup> / Lipid quantification |  |  |  |
| Quantitative | No | Normalization to reference | No |
| Batch correction | No |  |  |
| 7) PE[M-H]<sup>-</sup> / Lipid identification |  |  |  |
| Lipid class | PE | MS Level for identification | MS<sup>1</sup>, MS<sup>2</sup> |
| Identification level | Molecular species level | MS<sup>1</sup> adduct | [M-H]<sup>-</sup> |
| Isotope correction at MS<sup>1</sup> | No | MS<sup>2</sup> adduct | [M-H]<sup>-</sup> |
| Fragments for identification |  |  |  |
| Fragment name |  |  |  |
| GP(153) |  |  |  |
| -FA1(-H) |  |  |  |
| -FA2(-H) |  |  |  |
| HG(PE,140) |  |  |  |
| Isotope correction at MS<sup>2</sup> | No | MS<sup>1</sup> verified by standard | Yes |
| MS<sup>2</sup> verified by standard | Yes | Background check at MS<sup>1</sup> | Yes |
| Background check at MS<sup>2</sup> | Yes | Did you presume assumptions for identification? | Yes |
| Which assumptions were presumed? | only common fatty acyl chains included | Check on: | Isobaric overlap |
| Limit of detection | Signal threshold | RT verified by standard | Yes |
| Separation of isobaric/isomeric interference confirmed | No | Model for separation prediction | No |
| Lipid Identification Software | Compound Discoverer | Nomenclature for intact lipid molecule | Yes |
| Nomenclature for fragment ions | N/A |  |  |
| 7) PE[M-H]<sup>-</sup> / Lipid quantification |  |  |  |
| Quantitative | No | Normalization to reference | No |
| Batch correction | No |  |  |
| 8) PI[M-H]<sup>-</sup> / Lipid identification |  |  |  |
| Lipid class | PI | MS Level for identification | MS<sup>1</sup>, MS<sup>2</sup> |
| Identification level | Species level | MS<sup>1</sup> adduct | [M-H]<sup>-</sup> |
| Isotope correction at MS<sup>1</sup> | No | MS<sup>2</sup> adduct | [M-H]<sup>-</sup> |
| Fragments for identification |  |  |  |
| Fragment name |  |  |  |
| -FA1(-H) |  |  |  |
| -FA2(-H) |  |  |  |
| HG(PI,241) |  |  |  |
| Isotope correction at MS<sup>2</sup> | No | MS<sup>1</sup> verified by standard | Yes |
| MS<sup>2</sup> verified by standard | Yes | Background check at MS<sup>1</sup> | Yes |
| Background check at MS<sup>2</sup> | Yes | Did you presume assumptions for identification? | Yes |
| Which assumptions were presumed? | only common fatty acyl chains included | Check on: | Isobaric overlap |
| Limit of detection | Signal threshold | RT verified by standard | Yes |
| Separation of isobaric/isomeric interference confirmed | No | Model for separation prediction | No |
| Lipid Identification Software | Compound Discoverer | Nomenclature for intact lipid molecule | Yes |
| Nomenclature for fragment ions | N/A |  |  |
| 8) PI[M-H]- / Lipid quantification |  |  |  |
| Quantitative | No | Normalization to reference | No |
| Batch correction | No |  |  |
| 9) PS[M-H]- / Lipid identification |  |  |  |
| Lipid class | PS | MS Level for identification | MS<sup>1</sup>, MS<sup>2</sup> |
| Identification level | Molecular species level | MS<sup>1</sup> adduct | [M-H]- |
| Isotope correction at MS<sup>1</sup> | No | MS<sup>2</sup> adduct | [M-H]- |
| Isotope correction at MS<sup>2</sup> | No | MS<sup>1</sup> verified by standard | Yes |
| MS<sup>2</sup> verified by standard | Yes | Background check at MS<sup>1</sup> | Yes |
| Background check at MS<sup>2</sup> | Yes | Did you presume assumptions for identification? | Yes |
| Which assumptions were presumed? | only common fatty acyl chains included | Check on: | Isobaric overlap |
| Limit of detection | Signal threshold | RT verified by standard | Yes |
| Separation of isobaric/isomeric interference confirmed | No | Model for separation prediction | No |
| Lipid Identification Software | Compound Discoverer | Nomenclature for intact lipid molecule | Yes |
| Nomenclature for fragment ions | N/A |  |  |
| 9) PS[M-H]- / Lipid quantification |  |  |  |
| Quantitative | No | Normalization to reference | No |
| Batch correction | No |  |  |
| 10) PG[M-H]- / Lipid identification |  |  |  |
| Lipid class | PG | MS Level for identification | MS<sup>1</sup>, MS<sup>2</sup> |
| Identification level | Molecular species level | MS<sup>1</sup> adduct | [M-H]- |
| Isotope correction at MS<sup>1</sup> | No | MS<sup>2</sup> adduct | [M-H]- |
| Fragments for identification |  |  |  |
| Fragment name |  |  |  |
| FA1(+O) |  |  |  |
| FA2(+O) |  |  |  |
| HG(PG,171) |  |  |  |
| GP(153) |  |  |  |
| Isotope correction at MS<sup>2</sup> | No | MS<sup>1</sup> verified by standard | Yes |
| MS<sup>2</sup> verified by standard | Yes | Background check at MS<sup>1</sup> | Yes |
| Background check at MS<sup>2</sup> | Yes | Did you presume assumptions for identification? | Yes |
| Which assumptions were presumed? | only common fatty acyl chains included | Check on: | Isobaric overlap |
| Limit of detection | Signal threshold | RT verified by standard | Yes |
| Separation of isobaric/isomeric interferece confirmed | No | Model for separation prediction | No |
| Lipid Identification Software | Compound Discoverer | Nomenclature for intact lipid molecule | Yes |
| Nomenclature for fragment ions | N/A |  |  |
| 10) PG[M-H]- / Lipid quantification |  |  |  |
| Quantitative | No | Normalization to reference | No |
| Batch correction | No |  |  |
| 11) PC O[M+H]<sup>+</sup> / Lipid identification |  |  |  |
| Lipid class | PC O | MS Level for identification | MS<sup>1</sup>, MS<sup>2</sup> |
| Identification level | Molecular species level | MS<sup>1</sup> adduct | [M+H]<sup>+</sup> |
| Isotope correction at MS<sup>1</sup> | No | MS<sup>2</sup> adduct | [M+H]<sup>+</sup> |
| Fragments for identification |  |  |  |
| Fragment name |  |  |  |
| HG(PC,184) |  |  |  |
| Isotope correction at MS<sup>2</sup> | No | MS<sup>1</sup> verified by standard | No |
| MS<sup>2</sup> verified by standard | No | Background check at MS<sup>1</sup> | Yes |
| Background check at MS<sup>2</sup> | Yes | Did you presume assumptions for identification? | Yes |
| Which assumptions were presumed? | only common fatty acyl chains included | Check on: | Isobaric overlap |
| Limit of detection | Signal threshold | RT verified by standard | No |
| Separation of isobaric/isomeric interferece confirmed | No | Model for separation prediction | No |
| Lipid Identification Software | Compound Discoverer | Nomenclature for intact lipid molecule | Yes |
| Nomenclature for fragment ions | N/A |  |  |
| 11) PC O[M+H]<sup>+</sup> / Lipid quantification |  |  |  |
| Quantitative | No | Normalization to reference | No |
| Batch correction | No |  |  |
| 12) PE O[M-H]<sup>-</sup> / Lipid identification |  |  |  |
| Lipid class | PE O | MS Level for identification | MS<sup>1</sup>, MS<sup>2</sup> |
| Identification level | Molecular species level | MS<sup>1</sup> adduct | [M-H]<sup>-</sup> |
| Isotope correction at MS<sup>1</sup> | No | MS<sup>2</sup> adduct | [M-H]<sup>-</sup> |
| Fragments for identification |  |  |  |
| Fragment name |  |  |  |
| FA2(+O) |  |  |  |
| GP(153) |  |  |  |
| Isotope correction at MS<sup>2</sup> | No | MS<sup>1</sup> verified by standard | No |
| MS<sup>2</sup> verified by standard | No | Background check at MS<sup>1</sup> | Yes |
| Background check at MS<sup>2</sup> | Yes | Did you presume assumptions for identification? | Yes |
| Which assumptions were presumed? | only common fatty acyl chains included | Check on: | Isobaric overlap |
| Limit of detection | Signal threshold | RT verified by standard | No |
| Separation of isobaric/isomeric interference confirmed | No | Model for separation prediction | No |
| Lipid Identification Software | Compound Discoverer | Nomenclature for intact lipid molecule | Yes |
| Nomenclature for fragment ions | N/A |  |  |
| 12) PE O[M-H]<sup>-</sup> / Lipid quantification |  |  |  |
| Quantitative | No | Normalization to reference | No |
| Batch correction | No |  |  |
| 13) Cer[M+H]<sup>+</sup> / Lipid identification |  |  |  |
| Lipid class | Cer | MS Level for identification | MS<sup>1</sup> |
| Identification level | Species level | MS<sup>1</sup> adduct | [M+H]<sup>+</sup> |
| Isotope correction at MS<sup>1</sup> | No | MS<sup>1</sup> verified by standard | Yes |
| Background check at MS<sup>1</sup> | Yes | Did you presume assumptions for identification? | Yes |
| Which assumptions were presumed? | only common fatty acyl chains included | Check on: | Isobaric overlap |
| Limit of detection | Signal threshold | RT verified by standard | Yes |
| Separation of isobaric/isomeric interference confirmed | No | Model for separation prediction | No |
| Lipid Identification Software | Compound Discoverer | Nomenclature for intact lipid molecule | Yes |
| 13) Cer[M+H]<sup>+</sup> / Lipid quantification |  |  |  |
| Quantitative | No | Normalization to reference | No |
| Batch correction | No |  |  |
| 14) SM[M+H]<sup>+</sup> / Lipid identification |  |  |  |
| Lipid class | SM | MS Level for identification | MS<sup>1</sup> |
| Identification level | Species level | MS<sup>1</sup> adduct | [M+H]<sup>+</sup> |
| Isotope correction at MS<sup>1</sup> | No | MS<sup>1</sup> verified by standard | Yes |
| Background check at MS<sup>1</sup> | Yes | Did you presume assumptions for identification? | Yes |
| Which assumptions were presumed? | only common fatty acyl chains included | Check on: | Isobaric overlap |
| Limit of detection | Signal threshold | RT verified by standard | Yes |
| Separation of isobaric/isomeric interference confirmed | No | Model for separation prediction | No |
| Lipid Identification Software | Compound Discoverer | Nomenclature for intact lipid molecule | Yes |
| 14) SM[M+H]<sup>+</sup> / Lipid quantification |  |  |  |
| Quantitative | No | Normalization to reference | No |
| Batch correction | No |  |  |
| 15) HexCer[M+H]<sup>+</sup> / Lipid identification |  |  |  |
| Lipid class | HexCer | MS Level for identification | MS<sup>1</sup> |
| Identification level | Species level | MS<sup>1</sup> adduct | [M+H]<sup>+</sup> |
| Isotope correction at MS<sup>1</sup> | No | MS<sup>1</sup> verified by standard | No |
| Background check at MS<sup>1</sup> | Yes | Did you presume assumptions for identification? | Yes |
| Which assumptions were presumed? | only common fatty acyl chains included | Check on: | Isobaric overlap |
| Limit of detection | Signal threshold | RT verified by standard | No |
| Separation of isobaric/isomeric interference confirmed | No | Model for separation prediction | No |
| Lipid Identification Software | Compound Discoverer | Nomenclature for intact lipid molecule | Yes |
| 15) HexCer[M+H]<sup>+</sup> / Lipid quantification |  |  |  |
| Quantitative | No | Normalization to reference | No |
| Batch correction | No |  |  |
| 16) LacCer[M+H]<sup>+</sup> / Lipid identification |  |  |  |
| Lipid class | LacCer | MS Level for identification | MS<sup>1</sup> |
| Identification level | Species level | MS<sup>1</sup> adduct | [M+H]<sup>+</sup> |
| Isotope correction at MS<sup>1</sup> | No | MS<sup>1</sup> verified by standard | Yes |
| Background check at MS<sup>1</sup> | Yes | Did you presume assumptions for identification? | Yes |
| Which assumptions were presumed? | only common fatty acyl chains included | Check on: | Isobaric overlap |
| Limit of detection | Signal threshold | RT verified by standard | No |
| Separation of isobaric/isomeric interference confirmed | No | Model for separation prediction | No |
| Lipid Identification Software | Compound Discoverer | Nomenclature for intact lipid molecule | Yes |
| 16) LacCer[M+H]<sup>+</sup> / Lipid quantification |  |  |  |
| Quantitative | No | Normalization to reference | No |
| Batch correction | No |  |  |
| 17) PC P[M+H]<sup>+</sup> / Lipid identification |  |  |  |
| Lipid class | PC P | MS Level for identification | MS<sup>1</sup> |
| Identification level | Species level | MS<sup>1</sup> adduct | [M+H]<sup>+</sup> |
| Isotope correction at MS<sup>1</sup> | No | MS<sup>1</sup> verified by standard | No |
| Background check at MS<sup>1</sup> | No | Did you presume assumptions for identification? | Yes |
| Which assumptions were presumed? | only common fatty acyl chains included | Check on: | Isobaric overlap |
| Limit of detection | Signal threshold | RT verified by standard | No |
| Separation of isobaric/isomeric interference confirmed | No | Model for separation prediction | No |
| Lipid Identification Software | Compound Discoverer | Nomenclature for intact lipid molecule | Yes |
| 17) PC P[M+H]<sup>+</sup> / Lipid quantification |  |  |  |
| Quantitative | No | Normalization to reference | No |
| Batch correction | No |  |  |
| 18) PE P[M-H]<sup>-</sup> / Lipid identification |  |  |  |
| Lipid class | PE P | MS Level for identification | MS<sup>1</sup> |
| Identification level | Species level | MS<sup>1</sup> adduct | [M-H]<sup>-</sup> |
| Isotope correction at MS<sup>1</sup> | No | MS<sup>1</sup> verified by standard | No |
| Background check at MS<sup>1</sup> | No | Did you presume assumptions for identification? | Yes |
| Which assumptions were presumed? | only common fatty acyl chains included | Check on: | Isobaric overlap |
| Limit of detection | Signal threshold | RT verified by standard | No |
| Separation of isobaric/isomeric interferece confirmed | No | Model for separation prediction | No |
| Lipid Identification Software | Compound Discoverer | Nomenclature for intact lipid molecule | Yes |
| 18) PE P[M-H]<sup>-</sup> / Lipid quantification |  |  |  |
| Quantitative | No | Normalization to reference | No |
| Batch correction | No |  |  |

## Chemical Synthesis Materials and Methods

### π-Fatty Acids

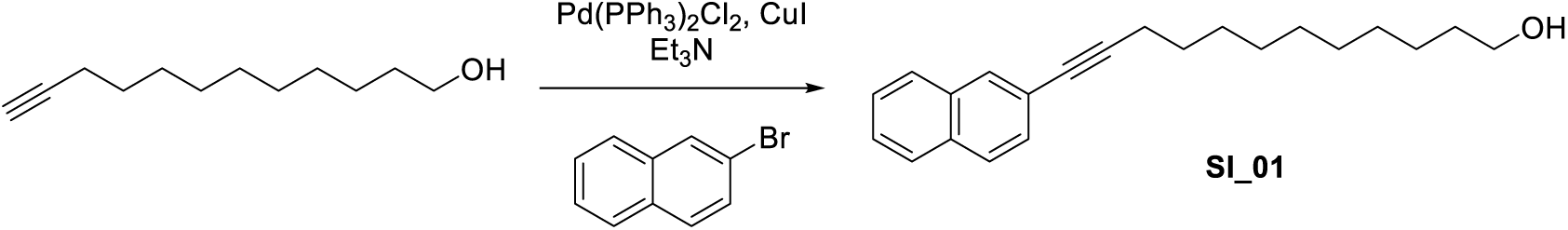

**12-(naphthalen-2-yl)dodec-11-yn-1-ol (SI_01):** 2-Bromonaphthalene (5.01 g, 24.2 mmol, 1.00 equiv) was dissolved in 30.0 mL triethylamine, followed by the addition of dodec-11-yn-1-ol (6.60 g, 36.2 mmol, 1.50 equiv), Pd(PPh_3_)_2_Cl_2_ (850 mg, 1.21 mmol, 0.05 equiv) and CuI (460 mg, 2.42 mmol, 0.10 equiv) under nitrogen atmosphere. The reaction mixture was heated to 80 °C and stirred for 2 h, then allowed to cool to ambient temperature. The reaction mixture was diluted with water and extracted three times with ethyl acetate. The combined organic layers were washed with brine, dried over Na_2_SO_4_, filtered and concentrated under reduced pressure. The residue was purified by silica gel chromatography (hexanes/ethyl acetate, 0-5%) to afford the title compound as a clear oil (7.38 g, 24.2 mmol, 99% yield).

**LCMS** *m/z* calculated for C_22_H_28_O^+^ ([M+H]^+^): 309.2 found: 309.2.

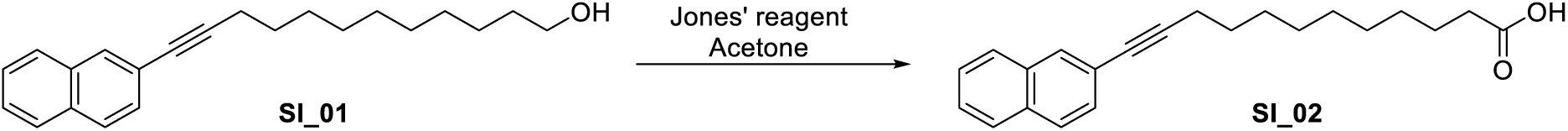

**12-(naphthalen-2-yl)dodec-11-ynoic acid (SI_02):** 12-(Naphthalen-2-yl)dodec-11-yn-1-ol (**SI_01**, 1.00 g, 3.24 mmol, 1.00 equiv) was dissolved in 10.0 mL acetone and cooled to 0 °C. Jones’ reagent (3.24 mL, 6.48 mmol, 2.00 equiv) was added, and the cooling bath was removed and the mixture allowed to warm to ambient temperature and stirred for 2 h. The reaction mixture was diluted with water and extracted with three times with ethyl acetate. The combined organic layers were washed with brine, dried over Na_2_SO_4_, filtered and concentrated under reduced pressure. The residue was purified by silica gel chromatography (hexanes/EtOAc, 5-20%) to afford the title compound **SI_02** (400 mg, 1.24 mmol, 38% yield).

**LCMS** *m/z* calculated for C_22_H_32_O^+^ ([M+H]^+^): 323.2 found: 323.2.

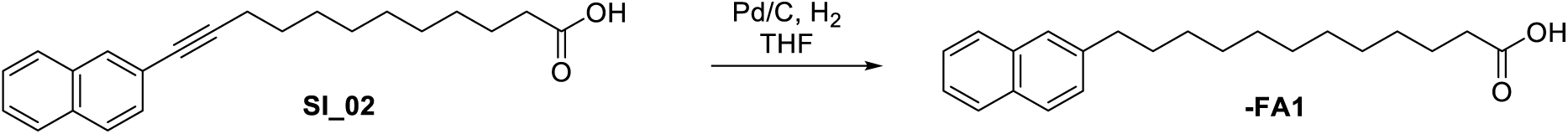

**π-FA1:** A mixture of 12-(naphthalen-2-yl)dodec-11-ynoic acid (**SI_02**, 330 mg, 1.02 mmol, 1.00 equiv) and 10% palladium on carbon (218 mg, 0.24 mmol, 0.24 equiv) in 5.0 mL tetrahydrofuran was stirred under hydrogen atmosphere (50 psi) overnight. The reaction mixture was filtered through a celite pad and the filtrate was evaporated to dryness. The residue was purified by C18 column chromatography (MeCN/water, 0-100%, with 0.1% formic acid) to afford **π-FA1** as a white solid (220 mg, 0.67 mmol, 66% yield).

**^1^H NMR** (400 MHz, CDCl_3_) δ 10.33 (bs, 1H) 7.81 – 7.75 (m, 3H), 7.60 (s, 1H), 7.48 – 7.37

(m, 2H), 7.33 (dd, *J* = 8.4, 1.6 Hz, 1H), 2.82 – 2.71 (t, *J* = 7.6 Hz, 2H), 2.34 (t, *J* = 7.6 Hz,

2H), 1.73-1.58 (m, 4H), 1.40 – 1.26 (m, 14H).

**^13^C NMR** (101 MHz, CDCl_3_) δ 179.6, 140.6, 133.8, 132.1, 127.8, 127.7, 127.6, 127.5, 126.4,

125.9, 125.1, 36.3, 34.1, 31.5, 29.70, 29.68, 29.65, 29.55, 29.47, 29.4, 29.2, 24.8.

**LCMS** *m/z* calculated for C_22_H29O_2_^-^ ([M-H]^-^): 327.2 found: 327.2.

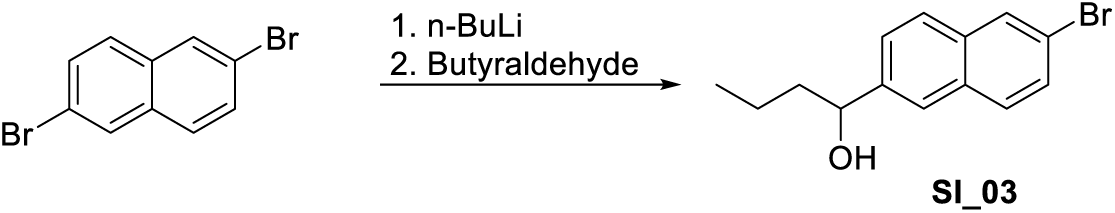

**1-(6-bromonaphthalen-2-yl)butan-1-ol (SI_03):** To a solution of 2,6-dibromonaphthalene (5.00 g, 17.5 mmol, 1.00 equiv) in 20.0 mL THF was added dropwise n-butyllithium (7.6 mL, 19.2 mmol, 1.10 equiv) at –78 °C. The reaction mixture was stirred at –78 °C for 30 min, then butyraldehyde (1.26 g, 17.5 mmol, 1.00 equiv) was added and the solution was allowed to warm up to room temperature and stirred for another 16 h. The reaction was quenched with saturated NH_4_Cl and extracted three times with EtOAc. The combined organic layers were washed with brine, dried over Na_2_SO_4_, filtered and concentrated under reduced pressure. The residue was purified by silica gel chromatography (hexanes/EtOAc, 0 - 20%) to afford the title compound **SI_03** as a yellow solid (2.4 g, 8.6 mmol, 49%).

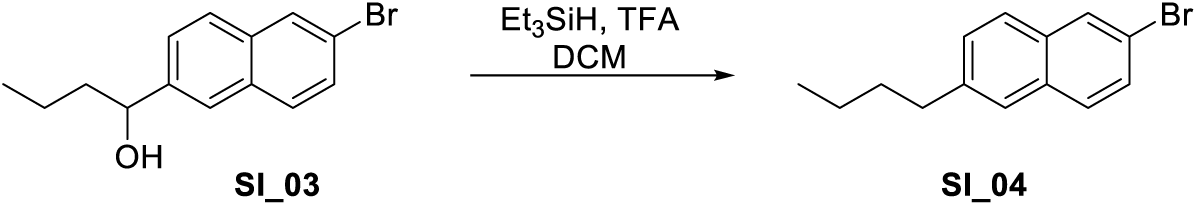

**2-Bromo-6-butylnaphthalene (SI_04):** To a solution of 1-(6-bromonaphthalen-2-yl)butan-1-ol (**SI_03**, 4.93 g, 17.7 mmol) in 20.0 mL dichloromethane, was added triethylsilane (10.3 g, 88.3 mmol, 5.00 equiv) and trifluoroacetic acid (10.1 g, 88.3 mmol, 5.00 equiv). The mixture was stirred at room temperature for 4 h. The reaction quenched by the addition of water and the resulting mixture extracted three times with ethyl acetate. The combined organic layers were washed with brine, dried over Na_2_SO_4_, filtered and concentrated under reduced pressure. The residue was purified by silica gel chromatography (hexanes/EtOAc, 0-5%) to afford the title compound **SI_04** as a white solid (2.4 g, 9.1 mmol, 52% yield).

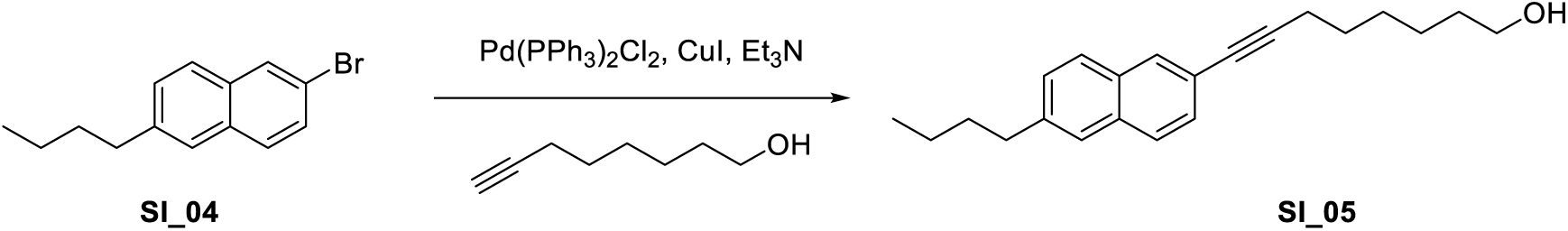

**8-(6-Butylnaphthalen-2-yl)oct-7-yn-1-ol (SI_05):** To a mixture of 2-bromo-6-butylnaphthalene (**SI_04**, 1.7 g, 6.46 mmol, 1.00 equiv) in 17.0 mL triethylamine were added oct-7-yn-1-ol (1.22 g, 9.69 mmol, 1.50 equiv), Pd(PPh_3_)_2_Cl_2_ (0.23 g, 0.32 mmol, 0.05 equiv) and CuI (0.12 g, 0.65 mmol, 0.10 equiv). The reaction mixture was stirred for 16 h at 80 ℃ under nitrogen atmosphere. The reaction quenched by the addition of water and the resulting mixture extracted three times with ethyl acetate. The combined organic layers were washed with brine, dried over Na_2_SO_4_, filtered and concentrated under reduced pressure. The residue was purified by silica gel chromatography (hexanes/EtOAc, 0-15%) to afford the title compound **SI_05** as a yellow solid (1.75 g, 5.67 mmol, 88% yield).

**LCMS** *m/z* calculated for C_22_H_29_O^+^ ([M+H]^+^): 309.21 found: 309.2.

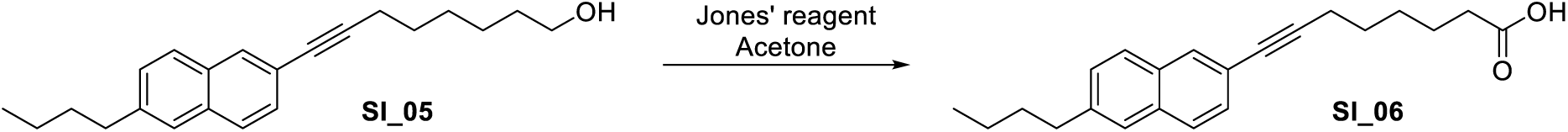

**8-(6-Butylnaphthalen-2-yl)oct-7-ynoic acid (SI_06):** To a mixture of 8-(6-butylnaphthalen-2-yl)oct-7-yn-1-ol (**SI_05**, 1.30 g, 4.21 mmol, 1.00 equiv) in 3.0 mL acetone at 0 °C was added Jones’ reagent (1.00 g, 5.05 mmol, 1.20 equiv) at 0℃. The mixture was stirred at 25 ℃ for 2 h. The reaction quenched by the addition of water and the ensuing mixture extracted twice with ethyl acetate. The combined organic layers were washed with brine, dried over Na_2_SO_4_, filtered and concentrated under reduced pressure. The residue was purified by silica gel chromatography (hexanes/EtOAc, 0-35%) to afford the title compound **SI_06** as a yellow solid (1.20, 3.72 mmol, 88%).

**LCMS** *m/z* calculated for C_22_H_25_O_2_ ^-^([M-H]^-^): 321.19 found: 321.0.

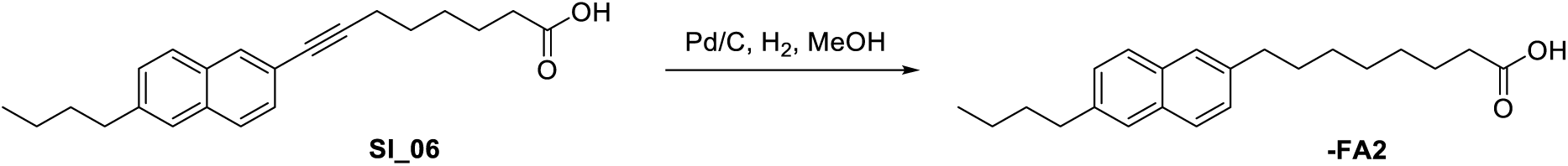

**π-FA2:** To a solution of 8-(6-butylnaphthalen-2-yl)oct-7-ynoic acid (**SI_06**, 400 mg, 1.24 mmol) in 10.0 mL methanol was added 10% palladium on carbon (264 mg, 0.248 mmol) and the reaction stirred under H_2_ atmosphere (50 psi) overnight. The reaction mixture was then filtered through a celite pad and the filtrate was evaporated to dryness. The residue was purified by C18 column chromatography (MeCN/water, 0-75%, with 0.1% formic acid) to afford **π-FA2** as a white solid (230 mg, 0.703 mmol, 57% yield).

**^1^H NMR** (400 MHz, DMSO-*d*_6_) δ 11.95 (s, 1H), 7.74 (d, *J* = 8.8 Hz, 2H), 7.61 (s, 2H), 7.33

(d, *J* = 8.4 Hz, 2H), 2.72-2.67 (m, 4H), 2.18 (t, *J* = 7.2 Hz, 2H), 1.69 – 1.58 (m, 4H), 1.53 –

1.43 (m, 2H), 1.40 – 1.21 (m, 8H), 0.91 (t, *J* = 7.2 Hz, 3H).

**^13^C NMR** (101 MHz, DMSO-*d*_6_) δ 174.5, 139.02, 139.00, 131.7, 127.3, 127.1, 125.7, 35.2,

34.9, 33.7, 33.0, 30.8, 28.58, 28.56, 28.51, 24.48, 21.8, 13.8

*Note: Four resonances corresponding to the naphthalene fragment are not visible in the*

*^13^C NMR due to overlap*.

**LCMS** *m/z* calculated for C_22_H_29_O_2_^-^ ([M-H]^-^): 325.22 found: 325.1.

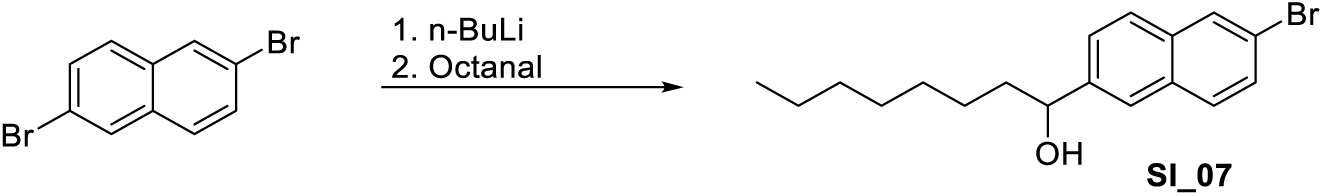

**1-(6-Bromonaphthalen-2-yl)octan-1-ol (SI_07):** To a solution of 2,6-dibromonaphthalene (10.0 g, 35.0 mmol, 1.00 equiv) in 100.0 mL THF was added dropwise n-butyllithium (15.3 mL, 38.4 mmol, 1.10 equiv, 2.50 M) at –78 °C. The reaction mixture was stirred at –78 °C for 50 min, then octanal (4.48 g, 35.0 mmol, 1.00 equiv) was added and the solution was allowed to warm up to room temperature and stirred for another 16 h. The reaction was quenched with saturated NH_4_Cl and extracted three times with EtOAc. The combined organic layers were washed with brine, dried over Na_2_SO_4_, filtered and concentrated under reduced pressure. The residue was purified by silica gel chromatography (hexanes/EtOAc, 0 - 20%) to afford the title compound **SI_07** as a yellow solid (5.29 g, 15.8 mmol, 49%).

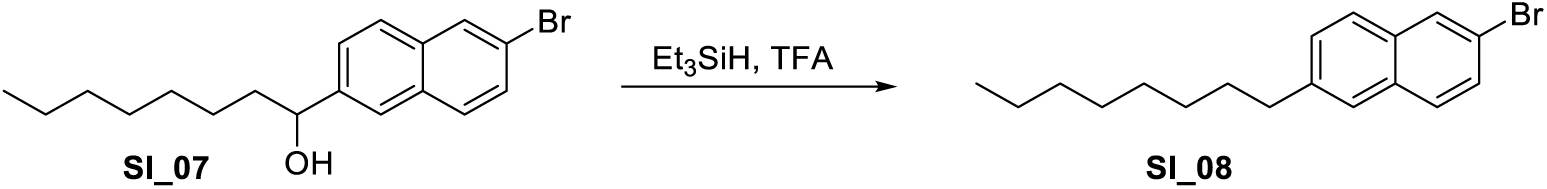

**2-Bromo-6-octylnaphthalene (SI_08):** To a solution of 1-(6-bromonaphthalen-2-yl)octan-1-ol (**SI_07**, 5.29 g, 15.8 mmol, 1.00 equiv) in 53.0 mL dichloromethane was added triethylsilane (9.20 g, 79.1 mmol, 5.00 equiv) and trifluoroacetic acid (9.01 g, 79.1 mmol, 5.00 equiv). The mixture was stirred at room temperature for 16 h. The reaction quenched by the addition of water and the resulting mixture extracted three times with ethyl acetate. The combined organic layers were washed with brine, dried over Na_2_SO_4_, filtered and concentrated under reduced pressure. The residue was purified by silica gel chromatography (hexanes/EtOAc, 0-5%) to afford the title compound **SI_08** as a white solid (4.58 g, 14.4 mmol, 91% yield).

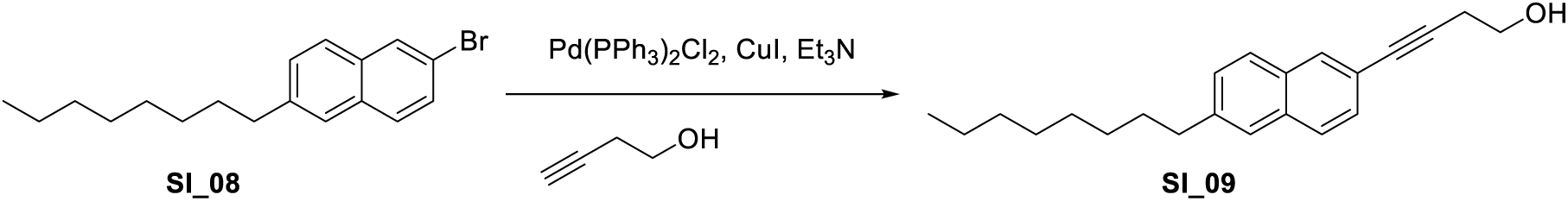

**4-(6-Octylnaphthalen-2-yl)but-3-yn-1-ol (SI_09):** To a mixture of 2-bromo-6-octylnaphthalene (**SI_08**, 1.9 g, 5.95 mmol, 1.00 equiv) in 10.0 mL triethylamine were added but-3-yn-1-ol (0.63 g, 8.93 mmol, 1.50 equiv), Pd(PPh_3_)_2_Cl_2_ (0.21 g, 0.30 mmol, 0.05 equiv) and CuI (0.11 g, 0.65 mmol, 0.10 equiv). The reaction mixture was stirred for 2 h at 80 ℃ under nitrogen atmosphere. The reaction quenched by the addition of water and the resulting mixture extracted three times with ethyl acetate. The combined organic layers were washed with brine, dried over Na_2_SO_4_, filtered and concentrated under reduced pressure. The residue was purified by silica gel chromatography (hexanes/EtOAc, 0-15%) to afford the title compound **SI_09** as a clear oil (1.1 g, 3.56 mmol, 60% yield).

**LCMS** *m/z* calculated. for C_22_H_28_O^+^ ([M+H]^+^): 309.2 found: 309.2.

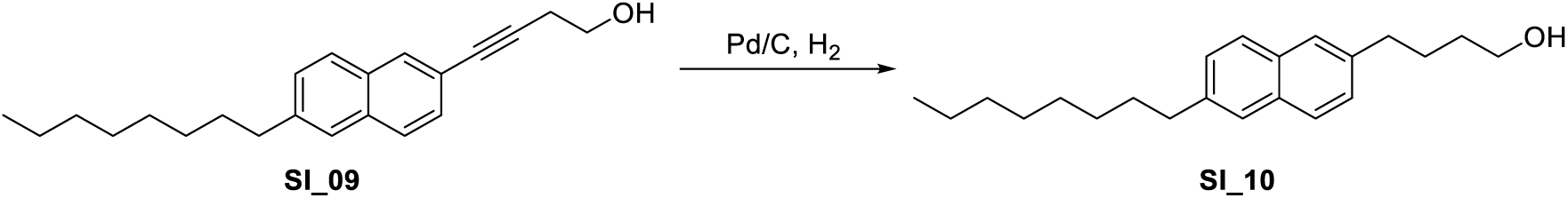

**4-(6-Octylnaphthalen-2-yl)butan-1-ol SI_10:** To a solution of 4-(6-octylnaphthalen-2-yl)but-3-yn-1-ol (**SI_09**, 1.00 g, 3.24 mmol, 1.00 equiv) in 20.0 mL THF was added 10% palladium on carbon (345 mg, 3.24 mmol, 1.00 equiv) and the reaction stirred under H_2_ atmosphere (50 psi) overnight. The reaction mixture was then filtered through a celite pad and the filtrate was evaporated to dryness. The residue was used in the next step without further purification.

**LCMS** *m/z* calculated. for C_22_H_32_O^+^ ([M+H]^+^): 313.2 found: 313.3.

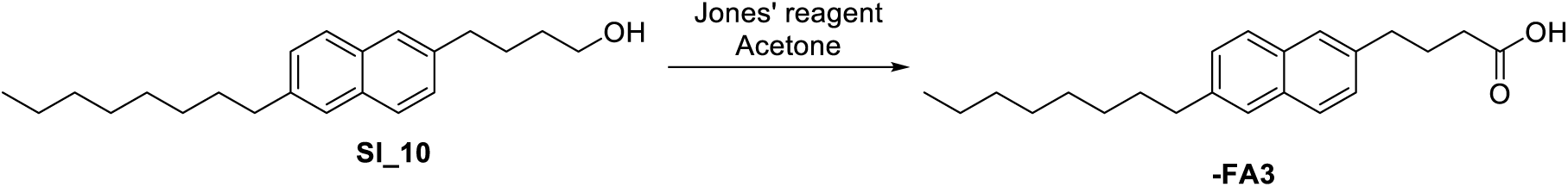

**π-FA3:** To a mixture of 4-(6-octylnaphthalen-2-yl)butan-1-ol (**SI_10**, 600 mg, 1.92 mmol, 1.00 equiv) in 10.0 mL acetone at 0 °C was added Jones’ reagent (1.2 mL, 2.50 mmol, 1.30 equiv) at 0℃. The mixture was stirred at 25 ℃ for 2 h. The reaction quenched by the addition of water and the ensuing mixture extracted twice with ethyl acetate. The combined organic layers were washed with brine, dried over Na_2_SO_4_, filtered and concentrated under reduced pressure. The residue was purified by silica gel chromatography (hexanes/EtOAc, 0-55%) to afford the title compound **π-FA3** as a yellow solid (1.20, 3.72 mmol, 88%).

**^1^H NMR** (400 MHz, CDCl_3_) δ 7.70 (m, 2H), 7.57 (s, 2H), 7.32–7.28 (m, 2H), 2.82 (m, 2H),

2.74(t, *J* = 7.6 Hz, 2H), 2.40 (t, *J* = 7.4 Hz, 2H), 2.12 – 1.99 (m, 2H), 1.72– 1.65 (m, 2H), 1.35 – 1.27 (m, 10H), 0.87 (t, *J* = 6.4 Hz, 3H).

**^13^C NMR** (101 MHz, CDCl_3_) δ 179.7, 140.0, 137.9, 132.4, 132.2, 127.8, 127.7, 127.4, 127.3,

126.5, 126.2, 36.2, 35.2, 33.4, 32.0, 31.6, 29.7, 29.5, 29.4, 26.3, 22.8, 14.3.

**LCMS** *m/z* calculated for C_22_H_29_O_2_^-^ ([M-H]^-^): 325.2 found: 325.2.

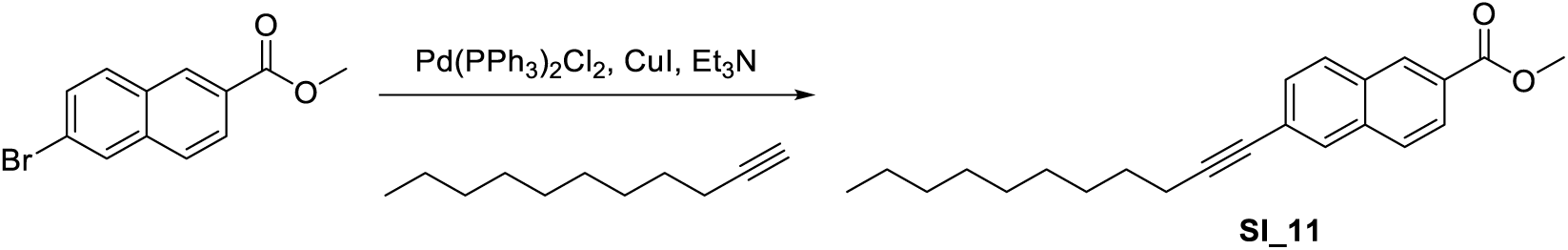

**Methyl 6-(undec-1-yn-1-yl)-2-naphthoate (SI_11):** To a mixture of methyl 6-bromo-2-naphthoate (2.0 g, 7.54 mmol, 1.00 equiv) in 15.0 mL triethylamine were added undec-1-yne (1.72 g, 11.3 mmol, 1.50 equiv), Pd(PPh_3_)_2_Cl_2_ (0.26 g, 0.38 mmol, 0.05 equiv) and CuI (0.14 g, 0.75 mmol, 0.10 equiv) under N_2_. The mixture was stirred for 12 h at 80 ℃ under nitrogen atmosphere. The reaction quenched by water and extracted three times with ethyl acetate. The combined organic layers were washed with brine, dried over anhydrous sodium sulfate and concentrated under reduced pressure. The residue was purified by silica gel chromatography (hexanes/EtOAc, 0 - 5%) to afford the title compound **SI_11** as a yellow solid (1.60 g, 4.75 mmol, 63% yield).

**LCMS** *m/z* calculated for C_23_H_29_O_2_^+^ ([M+H]^+^): 337.2 found: 337.1.

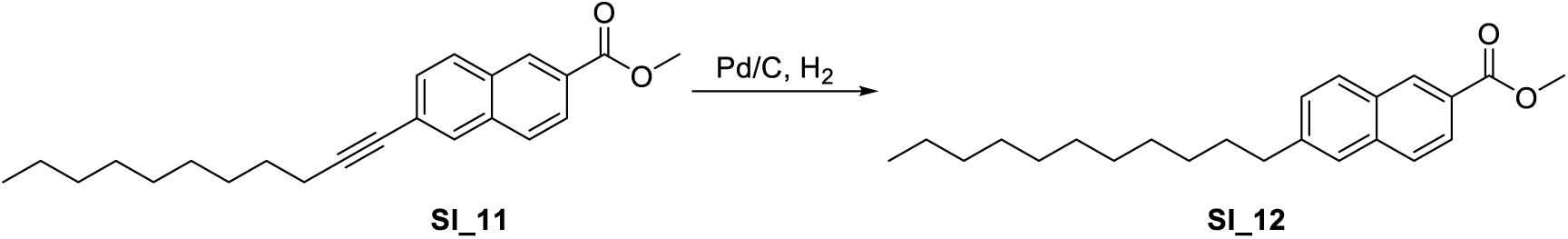

**Methyl 6-undecyl-2-naphthoate (SI_12):** To a mixture of methyl 6-(undec-1-yn-1-yl)-2-naphthoate (**SI_11**, 1.6 g, 4.76 mmol, 1.00 equiv) in 8.0 mL methanol 10% palladium on carbon (0.54 g, 0.51 mmol, 0.11 equiv) was added. The reaction mixture was stirred overnight under hydrogen atmosphere (50 psi). The reaction mixture was filtered through a celite pad and the filtrate was evaporated to dryness. The residue was used in the next step directly without further purification.

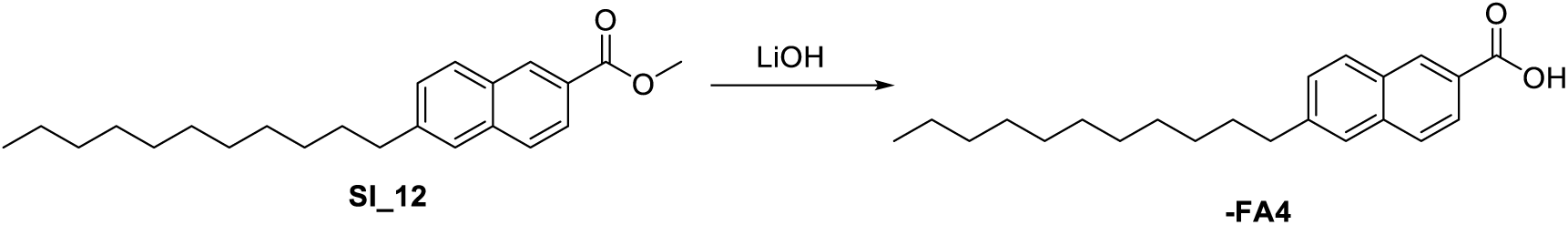

**π-FA4:** To a solution of methyl 6-undecyl-2-naphthoate (**SI_12**, 500 mg, 1.47 mmol, 1.00 equiv) in a mixture of methanol and water (1:1, 4.0 mL total) was added LiOH·H_2_O (123.3 mg, 2.94 mmol, 2.00 equiv) and the reaction stirred for 12 h at room temperature. The reaction mixture was diluted with water, the residue was adjusted to pH = 1, which was extracted three times with ethyl acetate. The combined organic layers were washed with brine, dried over Na_2_SO_4_, filtered and concentrated under reduced pressure. The residue was purified by silica gel chromatography (hexanes/EtOAc, 10 - 55%) to afford the title product **π-FA4** as a white solid (223 mg, 0.66 mmol, 45% yield).

**^1^H NMR** (400 MHz, DMSO-d_6_) δ 12.96 (s, 1H), 8.54 (s, 1H), 8.01 (d, J = 8.4 Hz, 1H), 7.97 –

7.87 (m, 2H), 7.75 (s, 1H), 7.46 (d, J = 8.4 Hz, 1H), 2.75 (t, J = 7.5 Hz, 3H), 1.71 – 1.60 (m,

3H), 1.21 (s, 14H), 0.83 (t, J = 6.8 Hz, 3H).

**^13^C NMR** (101 MHz, DMSO-d_6_) δ 167.5, 142.8, 135.2, 130.6, 130.3, 129.2, 128.2, 127.6,

127.3, 125.9, 125.2, 35.4, 31.3, 30.6, 29.01, 28.97, 28.96, 28.8, 28.70, 28.66, 22.1, 13.9.

**LCMS** *m/z* calculated for C_22_H_31_O_2_^+^ ([M+H]^+^): 327.2 found:327.1.

### Alkyl Glycerol Probes

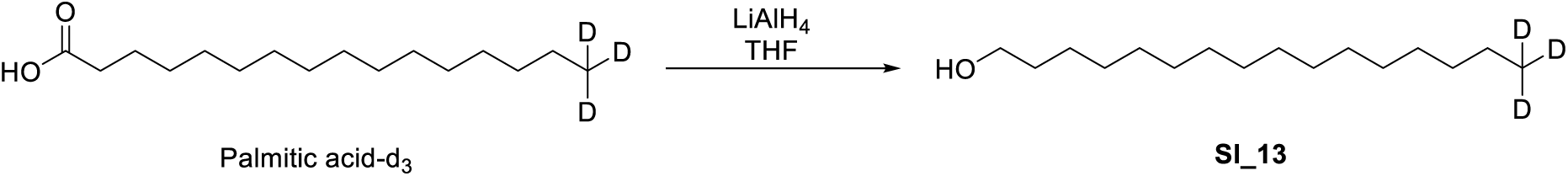

**Palmytol-d_3_ (SI-13)**: Palmitic acid-d_3_ (50.0 mg, 0.192 mmol, 1.00 equiv) was dissolved in

0.40 mL THF and added to a cooled (0 °C) solution of LiAlH_4_ (11.0 mg, 0.290 mmol, 1.50 equiv) in 0.6 mL THF. The resulting solution slowly warmed to ambient temperature over two hours. The reaction was quenched by the addition of sat. NH_4_Cl, and the mixture extracted with MBTE (3x). The combined organic solvents were dried over Na_2_SO_4_, filtered and evaporated to a crude which was immediately used in the next reaction

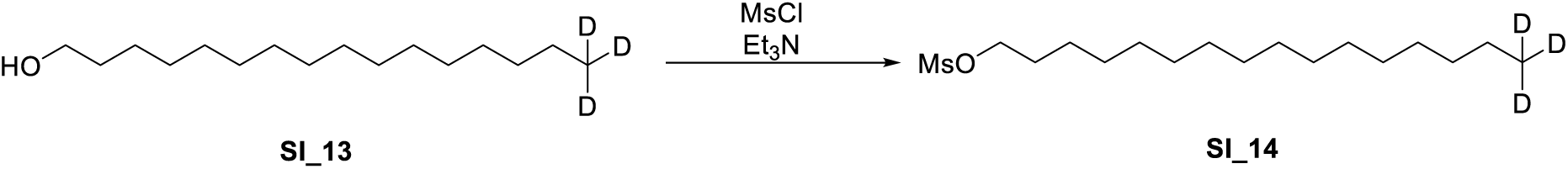

**Palmytol-d_3_ mesylate (SI_14):** Unpurified alcohol (assumed quant., 0.192 mmol, 1.00 equiv) was dissolved in 3.0 mL CH_2_Cl_2_ and cooled to 0°C using an ice bath. Sequentially a solution of Et_3_N (53.5 µL in 540 µL CH_2_Cl_2_, 0.384 mmol, 2.00 equiv) and MsCl (22.3 µL in 450 µL CH_2_Cl_2_, 0.288 mmol, 1.50 equiv) were added. Upon stirring for 30 min, the reaction was quenched by the addition of sat. NaHCO_3_ and the resulting mixture extracted 3x with CH_2_Cl_2_. The combined organic phases were dried over Na_2_SO_4_, filtered and evaporated to give the crude mesylate **SI-14**, which was immediately employed in the subsequent reaction.

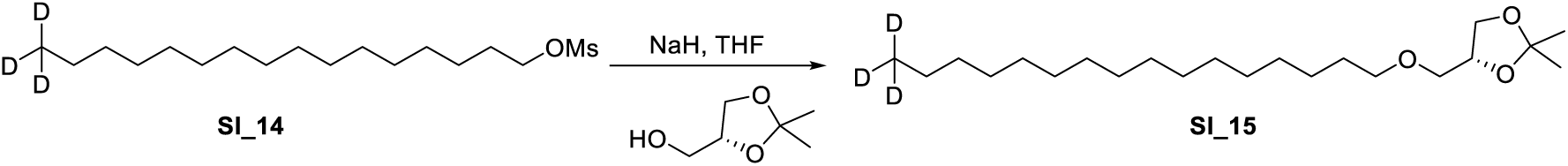

**D_3_-Pa-Alkylated (SI_15):** (R)-(2,2-dimethyl-1,3-dioxolan-4-yl)methanol (75 mg, 0.576 mmol, 3.00 equiv) was dissolved in 1.0 mL THF and stirred. NaH (24 mg, 0576 mmol, 3.00 equiv, 60% in oil) was added at once and the resulting mixture stirred for 10 minutes. Then, a solution of Mesylate **SI_14** (unpurified, assumed quant., 0.192 mmol, 1.00 equiv) in 0.4 mL THF (flask rinsed 4x with 0.05 mL THF) was added and the resulting mixture heated to 50 °C overnight. The reaction was then cooled to ambient temperature and quenched by the addition of sat. NH_4_Cl. The mixture was extracted 3x with MTBE, the combined organic phases were dried over Na_2_SO_4_, filtered and evaporated to a crude. Purification by column chromatography gave desired product **SI_15** as a clear oil (13 mg, 36.2 µmol, 19% yield over three steps).

**^1^H NMR** (400 MHz, CDCl_3_) δ 4.26 (p, *J* = 6.0 Hz, 1H), 4.06 (dd, *J* = 8.2, 6.4 Hz, 1H), 3.73

(dd, *J* = 8.2, 6.4 Hz, 1H), 3.56 – 3.37 (m, 4H), 1.65 – 1.51 (m, 2H), 1.42 (s, 3H), 1.36 (s, 3H), 1.25 (s, 26H).

**^13^C NMR** (101 MHz, CDCl_3_) δ 109.5, 74.9, 72.1, 72.0, 67.1, 32.0, 29.85, 29.83, 29.81, 29.76,

29.75, 29.70, 29.6, 29.5, 26.9, 26.2, 25.6, 22.6.

*Note: The resonance corresponding to the CD_3_ is not visible in the ^13^C NMR, also 3 resonances corresponding to carbon atoms within the fatty acid chain are not visible due to overlap*.

**HRMS** *m/z* calculated for C_22_H_42_D_3_O_3_ [M+H]^+^: 360.3552, found: 360.3552.

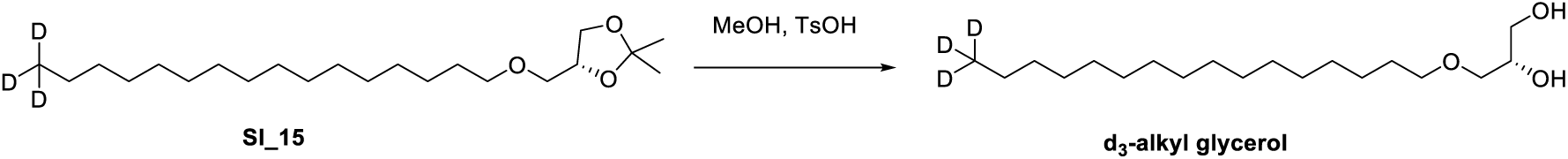

**d_3_-Alkyl glycerol:** Acetonide **SI_15** (13 mg, 36.2 µmol, 1.00 equiv) was dissolved in 0.9 mL MeOH and stirred. TsOH (7.0 mg, 36.2 µmol, 1.00 equiv) was added at once and the reaction mixture warmed to 35°C. After two hours the reaction was cooled to ambient temperature and quenched by the addition of sat. NaHCO3. The mixture was extracted 3x with EtOAc, the combined organic phases filtered and evaporated to a crude. Purification by column chromatography gave desired product **d_3_-alkyl glycerol** as a white solid (10.2 mg, 36.2 µmol, 88%).

**^1^H NMR (**400 MHz, CDCl_3_) δ 3.86 (tt, *J* = 5.6, 4.0 Hz, 1H), 3.72 (dd, *J* = 11.4, 3.9 Hz, 1H),

3.64 (dd, *J* = 11.4, 5.2 Hz, 1H), 3.57 – 3.39 (m, 4H), 1.57 (p, *J* = 6.7 Hz, 2H), 1.25 (s, 26H).

**^13^C NMR** (101 MHz, CDCl_3_) δ 72.7, 72.0, 70.6, 64.5, 32.0, 29.84, 29.82, 29.81, 29.76, 29.73,

29.6, 29.5, 26.2, 22.6.

*Note: The resonance corresponding to the CD_3_ is not visible in the ^13^C NMR, also 4 resonances corresponding to carbon atoms within the fatty acid chain are not visible due to overlap*.

**HRMS** *m/z* calculated for C_19_H_38_D_3_O_3_ [M+H]^+^: 320.3239, found: 320.3239.

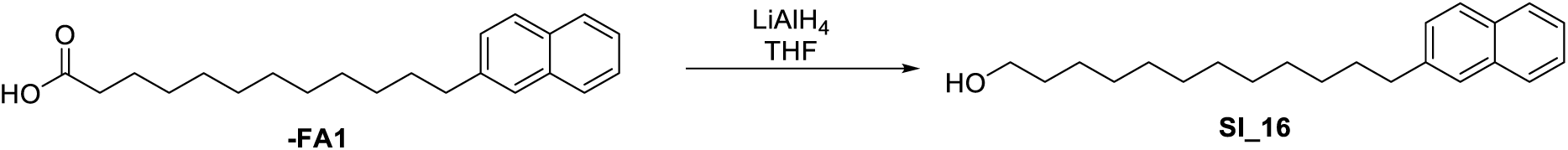

**π-FA1-alcohol (SI_16): π-FA1** (10.0 mg, 0.031 mmol, 1.00 equiv) was dissolved in 0.60 mL THF and cooled to 0 °C using an ice-water bath. LiAlH_4_ (3.5 mg, 0.092 mmol, 3.00 equiv) was added and the resulting solution slowly warmed to ambient temperature over 2 hours. The reaction was quenched by the addition of sat. NH_4_Cl, and the mixture extracted with MBTE (3x). The combined organic solvents were dried over Na_2_SO_4_, filtered and evaporated to a crude which was immediately used in the next reaction:

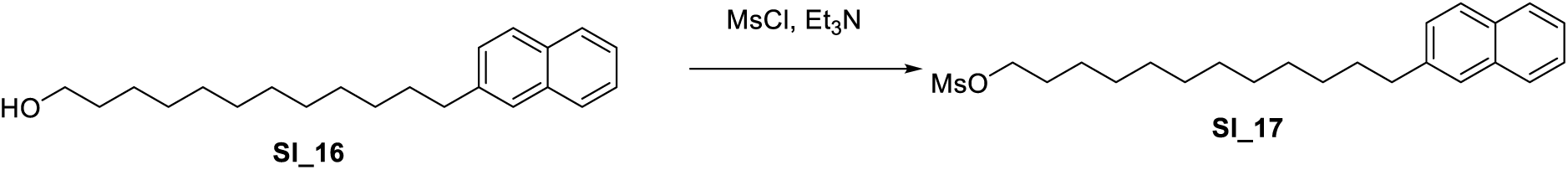

**π-FA1-mesylate (SI_17):** Unpurified alcohol **SI_16** (assumed quant., 0.031 mmol, 1.00 equiv) was dissolved in 0.6 mL CH_2_Cl_2_ and cooled to 0°C using an ice bath. Sequentially a solution of Et_3_N (8.5 µL in 85 µL CH_2_Cl_2_, 0.061 mmol, 2.00 equiv) and MsCl (3.6 µL in 72 µL CH_2_Cl_2_, 0.046 mmol, 1.50 equiv) were added. Upon stirring for 30 min, the reaction was quenched by the addition of sat. NaHCO_3_ and the resulting mixture extracted 3x with CH_2_Cl_2_. The combined organic phases were dried over Na_2_SO_4_ and purified by preparative TLC (6:2:2 Hexanes/CH_2_Cl_2_/MTBE) to give the desired compound **SI_17** as a colorless solid (7.4 mg, 0.019 mmol, 61%).

**^1^H NMR** (400 MHz, CDCl_3_) δ 7.71 (ddd, *J* = 13.9, 8.0, 3.0 Hz, 3H), 7.53 (s, 1H), 7.41 – 7.29 (m, 2H), 7.26 (dd, *J* = 8.4, 1.7 Hz, 1H), 4.14 (t, *J* = 6.6 Hz, 2H), 2.92 (d, *J* = 0.8 Hz, 3H), 2.73 – 2.65 (m, 2H), 1.72 – 1.57 (m, 4H), 1.40 – 1.05 (m, 16H).

**^13^C NMR** (101 MHz, CDCl_3_) δ 140.5, 133.6, 131.9, 127.7, 127.6, 127.5, 127.4, 126.3, 125.8,

125.0, 70.2, 37.4, 36.1, 31.4, 29.59, 29.56, 29.54, 29.51, 29.42, 29.35, 29.1, 29.0, 25.4.

*Note: HMRS data for the parent compound could not be detected, only an ion consistent with loss of MsO^-^:*

**HRMS** *m/z* calculated for C_22_H_31_ [M-MsO^-^]^+^: 295.2420, found: 295.2418.

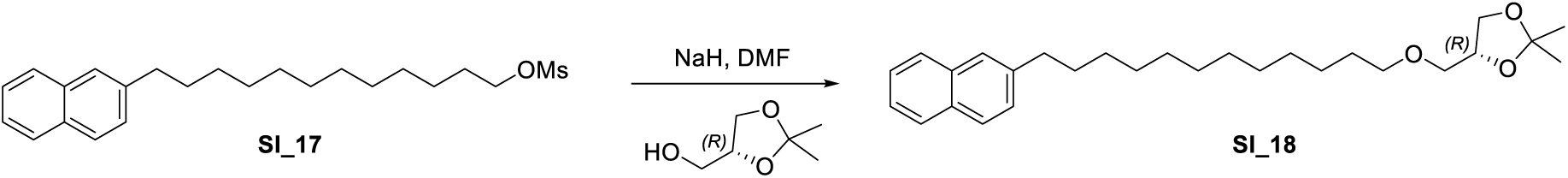

**π-FA1-acetonide (SI_18):** (R)-(2,2-dimethyl-1,3-dioxolan-4-yl)methanol (12.5 mg, 0.095 mmol, 5.00 equiv) was dissolved in 0.30 mL THF and cooled to 0 °C using an ice-water bath. NaH (3.9 mg mg, 0.095 mmol, 5.00 equiv, 60%) was added and the resulting mixture stirred until a clear solution was obtained (10 min). Then, Mesylate **SI_17** (7.4 mg, 0.019 mmol, 1.00 equiv) was added as a solution in THF (0.2 mL for dissolution, rinsed 4x with 0.05 mL each). The resulting solution was then concentrated to ca. A third of its volume under a stream of N_2_, sealed and heated to 45 °C overnight. TLC analysis showed incomplete conversion, so the reaction was further heated to 55 °C for 2 hours. The mixture was then quenched by the addition of sat. NaHCO_3_ and the resulting mixture extracted 3x with MTBE. The combined organic phases were dried over Na_2_SO_4_, filtered and evaporated to a crude. Purification by pTLC gave desired product **SI_18** as a colorless solid (4.6 mg, 10.8 µmol, 57%).

**^1^H NMR** (400 MHz, CDCl_3_) δ 7.78 (ddd, *J* = 14.2, 7.9, 3.2 Hz, 3H), 7.61 (d, *J* = 1.7 Hz, 1H), 7.48 – 7.38 (m, 2H), 7.33 (dd, *J* = 8.4, 1.8 Hz, 1H), 4.26 (p, *J* = 6.0 Hz, 1H), 4.06 (dd, *J* = 8.3, 6.4 Hz, 1H), 3.73 (dd, *J* = 8.2, 6.4 Hz, 1H), 3.56 – 3.36 (m, 4H), 2.81 – 2.72 (m, 2H), 1.70 (p, *J* = 7.5 Hz, 2H), 1.56 (m, 2H), 1.42 (s, 3H), 1.36 (s, 3H), 1.32 – 1.20 (m, 16H).

**^13^C NMR** (101 MHz, CDCl_3_) δ 140.6, 133.8, 132.0, 127.8, 127.7, 127.6, 127.5, 126.4, 125.9, 125.1, 109.5, 74.9, 72.04, 71.96, 67.1, 36.3, 31.5, 29.8, 29.73, 29.73, 29.70, 29.69, 29.6, 29.5, 26.9, 26.2, 25.6.

**HRMS** *m/z* calculated for C_28_H_43_O_3_ [M+H]^+^: 427.3207, found: 427.3210.

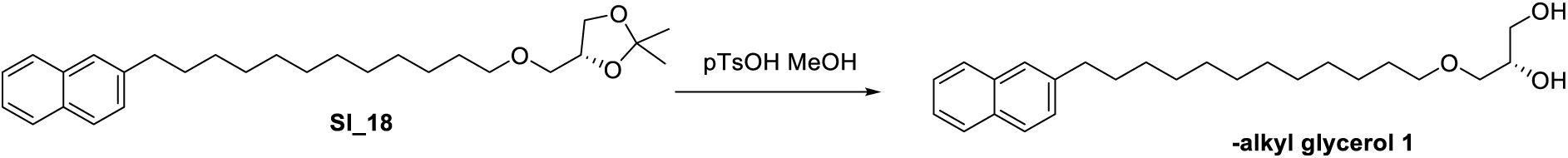

**π-Alkyl glycerol 1:** Acetonide **SI_18** (4.6 mg, 10.8 umol, 1.00 equiv) was dissolved in 0.3 mL MeOH and stirred. TsOH (2.1 mg, 10.8 umol, 1.00 equiv) was added at once, and the reaction mixture warmed to 35 °C. Upon stirring for 3 hours, the reaction was quenched by the addition of sat. NaHCO_3_ and the resulting mixture extracted 3x with EtOAc. The combined organic phases were dried over Na_2_SO_4_, filtered and evaporated to a crude. Purification by pTLC gave desired product **π-alkyl glycerol 1** as a white solid (2.5 mg, 6.5 µmol, 60%).

**^1^H NMR** (400 MHz, CDCl_3_) δ 7.78 (ddd, *J* = 14.1, 7.9, 3.1 Hz, 3H), 7.61 (d, *J* = 1.7 Hz, 1H), 7.50 – 7.37 (m, 2H), 7.33 (dd, *J* = 8.3, 1.7 Hz, 1H), 3.85 (d, *J* = 7.2 Hz, 1H), 3.72 (d, *J* = 11.6 Hz, 1H), 3.65 (dd, *J* = 11.3, 5.0 Hz, 1H), 3.58 – 3.39 (m, 4H), 2.81 – 2.72 (m, 2H), 2.57 (s, 1H), 2.14 (s, 1H), 1.70 (p, *J* = 7.5 Hz, 2H), 1.57 (t, *J* = 6.7 Hz, 4H), 1.39 – 1.17 (m, 14H).

**^13^C NMR** (101 MHz, CDCl_3_) δ 140.6, 133.8, 132.0, 127.8, 127.7, 127.6, 127.5, 126.4, 125.9, 72.7, 72.0, 70.5, 64.5, 36.3, 31.5, 29.9, 29.8, 29.72 (2C, overlap), 29.71, 29.68, 29.6, 29.5, 26.2.

**HRMS** *m/z* calculated for C_25_H_39_O_3_ [M+H]^+^: 387.2894, found: 387.2871.

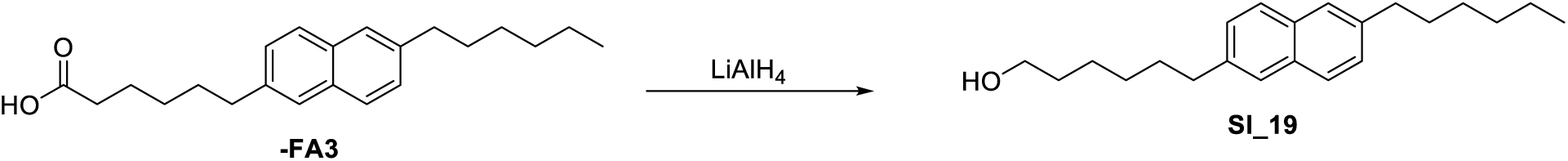

**π-FA3-alcohol (SI_19)**: **π-FA3** (20.0 mg, 0.062 mmol, 1.00 equiv) was dissolved in 1.20 mL THF and cooled to 0 °C using an ice-water bath. LiAlH_4_ (7.00 mg, 0.186 mmol, 3.00 equiv) was added and the resulting solution slowly warmed to ambient temperature over 3 hours. The reaction was quenched by the addition of sat. NH_4_Cl, and the mixture extracted with MBTE (3x). The combined organic solvents were dried over Na_2_SO_4_, filtered and evaporated to give crude **SI_19** which was immediately used in the next reaction:

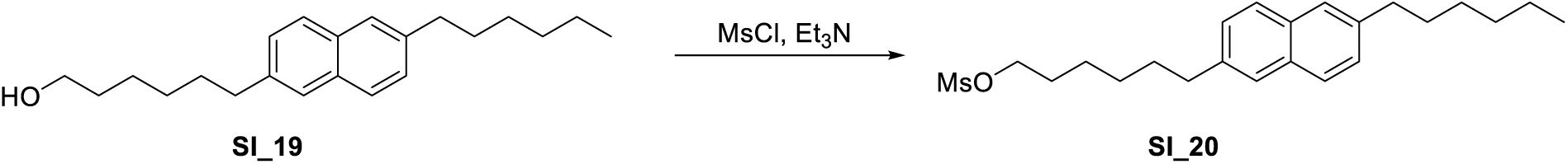

**π-FA1-mesylate (SI_20):** Unpurified alcohol **SI_19** (assumed quant., 0.062 mmol, 1.00 equiv) was dissolved in 1.20 mL CH_2_Cl_2_ and cooled to 0°C using an ice bath. Sequentially, Et_3_N (19 µL 0.124 mmol, 2.00 equiv) and MsCl (7.2 µL 0.093 mmol, 1.50 equiv) were added. Upon stirring for 30 min, the reaction was quenched by the addition of sat. NaHCO_3_ and the resulting mixture extracted 3x with CH_2_Cl_2_. The combined organic phases were dried over Na_2_SO_4_ and purified by preparative TLC (6:2:2 Hexanes/CH_2_Cl_2_/MTBE) to give the desired compound **SI_20** as a colorless solid (20.6 mg, 52.7 µmol, 85% over two steps).

**^1^H NMR** (400 MHz, CDCl_3_) δ 7.70 (dd, *J* = 8.3, 5.6 Hz, 2H), 7.57 (d, *J* = 2.1 Hz, 2H), 7.30 (ddd, *J* = 12.7, 8.4, 1.7 Hz, 2H), 4.28 – 4.21 (m, 2H), 2.97 (s, 3H), 2.86 – 2.79 (m, 2H), 2.79 – 2.70 (m, 2H), 1.82 (ttd, *J* = 7.7, 4.6, 1.9 Hz, 4H), 1.71 – 1.65 (m, 2H), 1.41 – 1.22 (m, 10H), 0.96 – 0.84 (m, 3H).

**^13^C NMR** (101 MHz, CDCl_3_) δ 140.1, 138.2, 132.4, 132.1, 127.8, 127.7, 127.4, 127.2, 126.4, 126.2, 70.0, 37.5, 36.2, 35.4, 32.0, 31.6, 29.7, 29.5, 29.4, 28.7, 27.2, 22.8, 14.3.

*Note: HMRS data for the parent compound could not be detected, only an ion consistent with loss of MsO^-^:*

**HRMS** *m/z* calculated for C_22_H_31_ [M-MsO^-^]^+^: 295.2420, found: 295.2421.

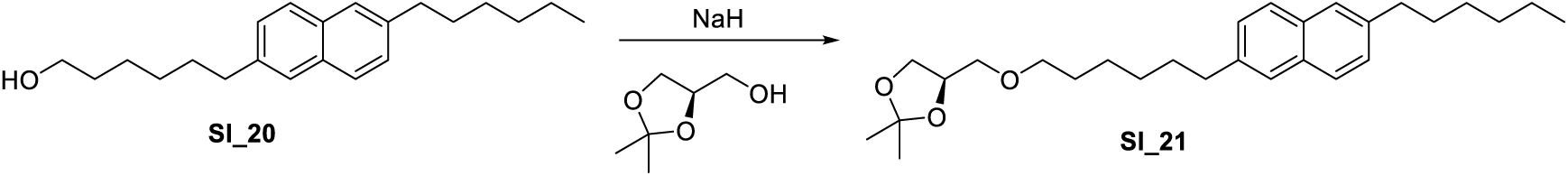

**π-FA3-acetonide (SI_21):** (R)-(2,2-dimethyl-1,3-dioxolan-4-yl)methanol (34.9 mg, 0.264 mmol, 5.00 equiv) was dissolved in 0.80 mL THF and cooled to 0 °C using an ice-water bath. NaH (6.3 mg mg, 0.264 mmol, 5.00 equiv, 60%) was added and the resulting mixture stirred until a clear solution was obtained (10 min). Then, mesylate **SI_20** (20.6 mg, 0.053 mmol, 1.00 equiv) was added as a solution in THF (0.4 mL for dissolution, rinsed 4x with 0.05 mL each). The resulting solution was then sealed and heated to 50 °C overnight. The mixture was then quenched by the addition of sat. NaHCO_3_ and the resulting mixture extracted 3x with MTBE. The combined organic phases were dried over Na_2_SO_4_, filtered and evaporated to a crude. Purification by pTLC gave desired product **SI_21** as a colorless solid (12.6 mg, 29.5 µmol, 56%).

**^1^H NMR** (400 MHz, CDCl_3_) δ 7.69 (dd, *J* = 8.5, 2.2 Hz, 2H), 7.56 (Bs, 2H), 7.29 (ddd, *J* = 8.4, 4.3, 1.6 Hz, 2H), 4.31 – 4.20 (m, 1H), 4.05 (dd, *J* = 8.3, 6.4 Hz, 1H), 3.72 (dd, *J* = 8.3, 6.4 Hz, 1H), 3.56 – 3.44 (m, 3H), 3.41 (dd, *J* = 9.9, 5.5 Hz, 1H), 2.76 (dt, *J* = 11.4, 7.6 Hz, 6H), 1.82 – 1.59 (m, 6H), 1.42 (s, 3H), 1.36 (s, 3H), 1.35 – 1.22 (m, 8H), 0.93 – 0.85 (m, 3H).

**^13^C NMR** (101 MHz, CDCl_3_) δ 139.8, 139.1, 132.3, 132.2, 127.6, 127.5, 127.43, 127.36, 126.3, 126.2, 109.5, 74.9, 72.0, 71.8, 67.1, 36.2, 35.9, 32.0, 31.6, 29.7, 29.5, 29.4, 29.3, 28.0, 26.9, 25.6, 22.8, 14.3.

**HRMS** *m/z* calculated for C_28_H_43_O_3_ [M+H]^+^: 427.3207, found: 427.3210.

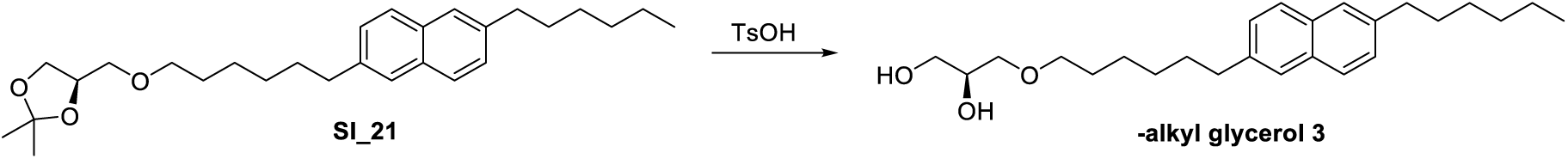

**π-Alkyl glycerol 3:** Acetonide **SI_21** (12.6 mg, 29.5 umol, 1.00 equiv) was dissolved in 1.20 mL MeOH and stirred. TsOH (5.6 mg, 29.5 umol, 1.00 equiv) was added at once, and the reaction mixture warmed to 35 °C. Upon stirring for 3 hours, the reaction was quenched by the addition of sat. NaHCO_3_ and the resulting mixture extracted 3x with EtOAc. The combined organic phases were dried over Na_2_SO_4_, filtered and evaporated to a crude. Purification by pTLC gave desired product **π-alkyl glycerol 3** as a white solid (8.7 mg, 23 µmol, 76%).

**^1^H NMR** (400 MHz, CDCl_3_) δ 7.39 (dd, *J* = 8.3, 3.1 Hz, 2H), 7.26 (s, 2H), 6.99 (ddd, *J* = 8.4, 6.5, 1.7 Hz, 2H), 3.54 (ddd, *J* = 7.8, 5.7, 3.9 Hz, 1H), 3.40 (dd, *J* = 11.4, 3.9 Hz, 1H), 3.33 (dd, *J* = 11.4, 5.2 Hz, 1H), 3.26 – 3.13 (m, 4H), 2.53 – 2.40 (m, 4H), 2.27 (s, 1H), 1.84 (s, 1H), 1.49 – 1.31 (m, 6H), 1.13 – 0.92 (m, 10H), 0.69 – 0.53 (m, 3H).

**^13^C NMR** (101 MHz, CDCl_3_) δ 139.9, 138.9, 132.3, 132.2, 127.7, 127.5, 127.4, 126.3, 126.2, 72.6, 71.7, 70.6, 64.4, 36.2, 35.9, 32.0, 31.6, 29.9, 29.7, 29.5, 29.4, 29.3, 27.9, 22.8, 14.3.

**HRMS** *m/z* calculated for C_25_H_39_O_3_ [M+H]^+^: 387.2894, found: 387.2891.

### π-Sphingosines

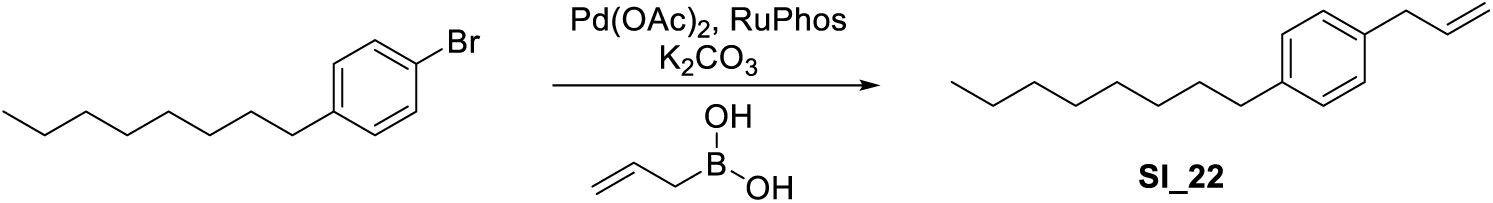

**1-Allyl-4-octylbenzene (SI_22):** 1-Bromo-4-octylbenzene (2.00 g, 7.43 mmol, 1.00 equiv) was dissolved in a mixture of PhMe and H_2_O (22 mL, 10:1). Allylboronic acid (0.77 g, 8.91 mmol, 1.20 equiv), K_2_CO_3_ (3.08 g, 22.3 mmol, 3.00 equiv), Pd(OAc)_2_ (197 mg, 0.88 mmol, 0.12 equiv), and Ruphos (819 mg, 1.76 mmol, 0.24 equiv) were added and the reaction mixture was stirred for 5 h at 80°C. EtOAc and water were added, the phases separated and the aqueous phase twice extracted with EtOAc. The combined organic phases were dried over Na_2_SO_4_, filtered, and concentrated. The mixture was purified by flash column chromatography (hexanes) to yield the title compound **SI_22** as colorless oil (1.20 g, 5.20 mmol, 70%).

**^1^H NMR** (400 MHz, CDCl_3_) δ 7.11 (s, 4H), 5.97 (ddt, *J* = 16.9, 10.0, 6.7 Hz, 1H), 5.13 – 5.01 (m, 2H), 3.36 (d, *J* = 6.7 Hz, 2H), 2.63 – 2.50 (m, 2H), 1.65 – 1.56 (m, 2H), 1.32 – 1.23 (m, 10H), 0.87 (d, *J* = 7.0 Hz, 3H).

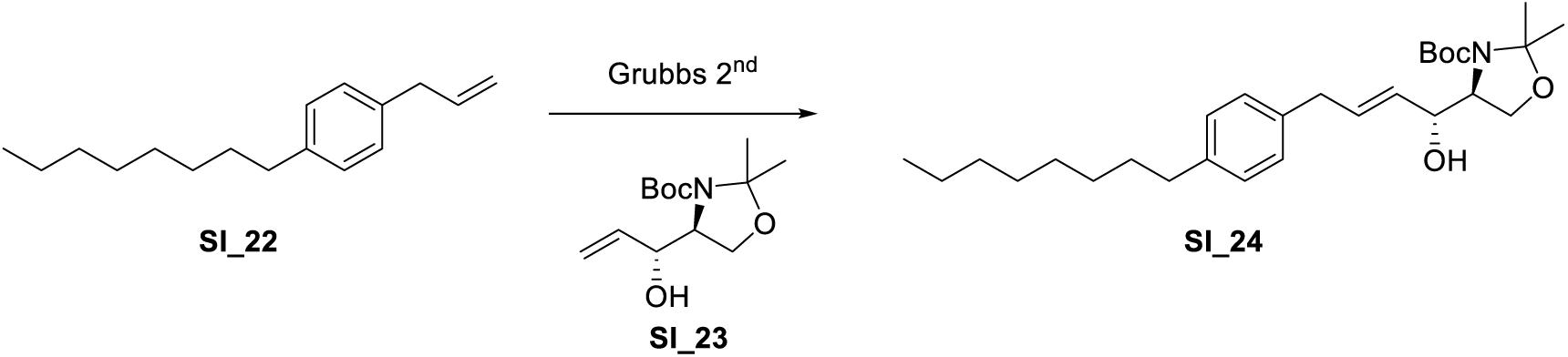

**tert-butyl (S)-4-((R,E)-1-hydroxy-4-(4-octylphenyl)but-2-en-1-yl)-2,2-dimethyloxazol-idine-3-carboxylate (SI_24):** 1-Allyl-4-octylbenzene (100 mg, 0.43 mmol, 1.00 equiv) was dissolved in CH_2_Cl_2_ (5 mL). The vinyl adduct of Garner’s aldehyde (**SI_23**, 223 mg, 0.866 mmol, 2.00 equiv) and HG-II (5 mg, 5.89 µmol, 1.37 %) were added and the reaction mixture was stirred for 16h at 40°C. The reaction mixture was concentrated and purified by flash column chromatography (PE/EA: 0% ∼ 10%) to yield **SI_24** as colorless oil (80 mg, 0.174 mmol 40%).

**LCMS**: *m/z* calculated for C_27_H_46_NO_4_Na^+^ ([M+Na]^+^): 482.3 found: 482.4.

*Note: **SI_23** was prepared according to the procedure outlined in J. Org. Chem. **1998**, 22, 7999*.

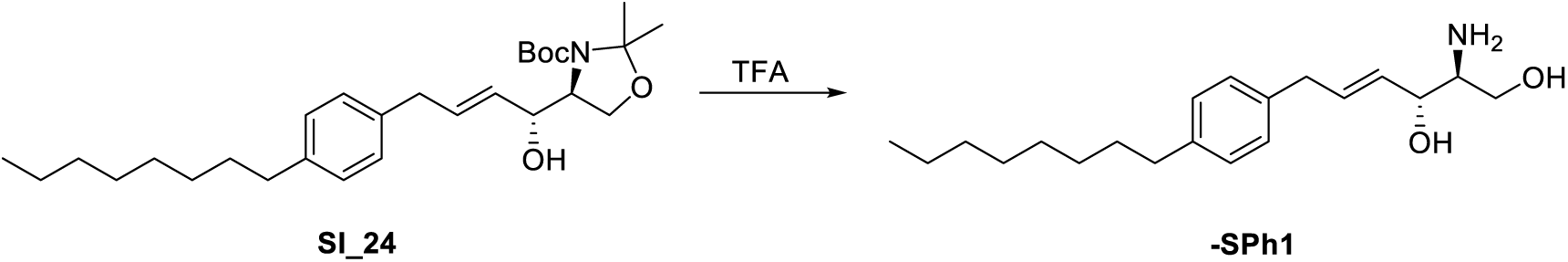

**π-Sph1: SI_24** (80.0 mg, 0.17 mmol, 1.00 equiv) was dissolved in 15.0 mL DCM. 3.00 mL TFA were added and the reaction mixture stirred for 1 hour at ambient temperature, then it was concentrated and the residue directly purified by HPLC (C18, MeCN/H_2_O 40-50% over 16 min, R_t_ = 8.3-9.5 min) to give the title compound **π-Sph1** as a colorless solid (26.5 mg, 0.082 mmol, 48%).

**^1^H NMR** (400 MHz, MeOD) δ 7.10 (s, 4H), 5.99 (dt, *J* = 14.1, 6.8 Hz, 1H), 5.54 (dd, *J* = 15.3, 6.7 Hz, 1H), 4.36 – 4.28 (m, 1H), 3.79 (dd, *J* = 11.6, 4.0 Hz, 1H), 3.66 (dd, *J* = 11.6, 8.4 Hz, 1H), 3.39 (d, *J* = 6.8 Hz, 2H), 3.22 (dt, *J* = 8.5, 4.3 Hz, 1H), 2.61 – 2.52 (m, 2H), 1.59 (d, *J* = 7.1 Hz, 2H), 1.30 (m, 10H), 0.89 (t, *J* = 6.8 Hz, 3H).

**LCMS** *m/z* calculated for C_20_H_34_NO_2_^+^ ([M+H]^+^): 320.3 found: 320.2.

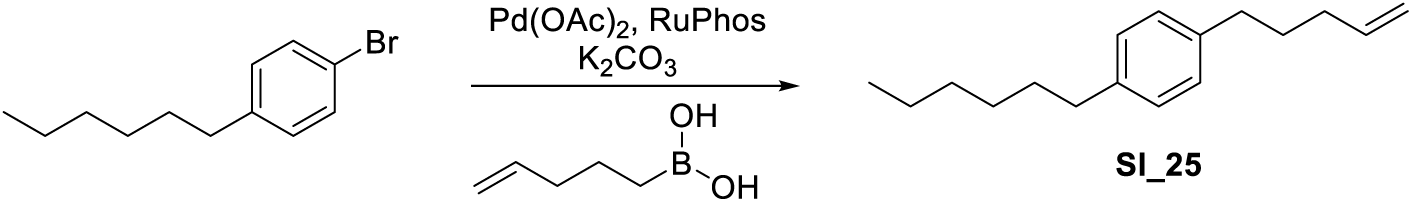

**1-Hexyl-4-(pent-4-en-1-yl)benzene (SI_25):** 1-Bromo-4-hexylbenzene (3.10 g, 21.1 mmol, 1.00 equiv) was dissolved in a mixture of PhMe and H_2_O (22 mL, 10:1). 4-Pentenylboronic acid (2.00 g, 17.6 mmol, 0.83 equiv), K_2_CO_3_ (7.30 g, 52.7 mmol, 2.50 equiv), Pd(OAc)_2_ (197 mg, 0.88 mmol, 0.042 equiv), and Ruphos (819 mg, 1.76 mmol, 0.083 equiv) were added and the reaction mixture was stirred for 5 h at 80°C. EtOAc and water were added, the phases separated and the aqueous phase twice extracted with EtOAc. The combined organic phases were dried over Na_2_SO_4_, filtered, and concentrated. The mixture was purified by flash column chromatography (hexanes) to yield the title compound as colorless oil (1.50 g, 7.81 mmol, 37%).

**^1^H NMR** (400 MHz, CDCl_3_) δ 7.02 (s, 4H), 5.77 (ddt, *J* = 17.0, 10.1, 6.6 Hz, 1H), 5.00 – 4.86

(m, 2H), 2.51 (q, *J* = 8.4 Hz, 4H), 2.02 (q, *J* = 7.1 Hz, 2H), 1.62 (q, *J* = 7.5 Hz, 2H), 1.55 –

1.49 (m, 2H), 1.25 (d, *J* = 9.4 Hz, 6H), 0.81 (t, *J* = 6.5 Hz, 3H).

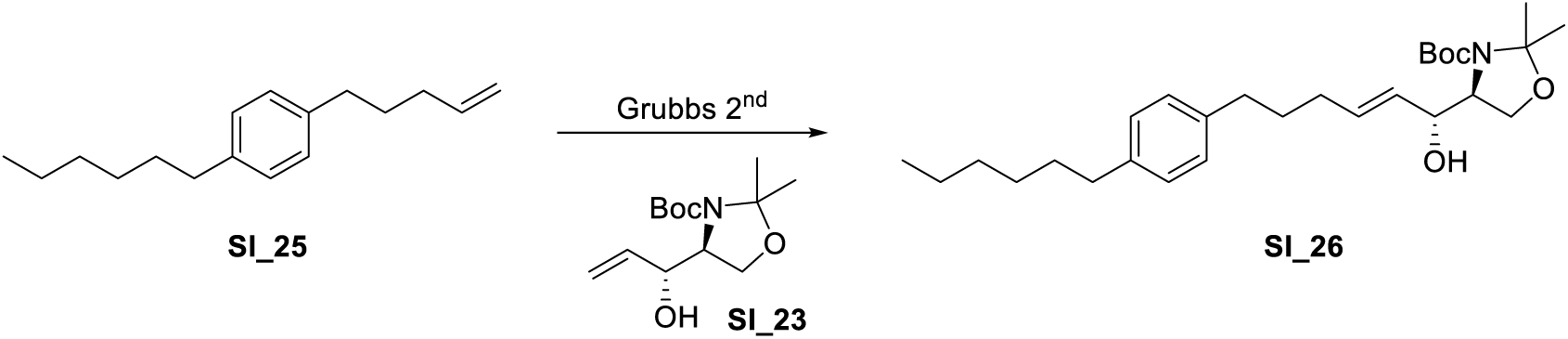

**tert-butyl (S)-4-((R,E)-6-(4-hexylphenyl)-1-hydroxyhex-2-en-1-yl)-2,2-dimethyloxazoli-dine-3-carboxylate (SI_26): SI_25** (1.50 g, 6.51 mmol, 1.00 equiv) was dissolved in CH_2_Cl_2_ (15 mL). The vinyl adduct of Garner’s aldehyde **SI_23** (2.50 g, 9.77 mmol, 1.50 equiv) and HG-II (50 mg, 0.0589 mmol, 0.90 mol%) were added and the reaction mixture was stirred for

16 h at 40°C. The reaction mixture was concentrated and purified by flash column chromatography (PE/EA: 0% ∼ 10%) to yield **SI_26** as colorless oil (1.90 g, 2.60 mmol, 40%).

**LCMS** *m/z* calculated for C_27_H_46_NO_4_Na^+^ ([M+Na]^+^): 482.3 found: 482.4.

*Note: **SI_23** was prepared according to the procedure outlined in J. Org. Chem. **1998**, 22, 7999*.

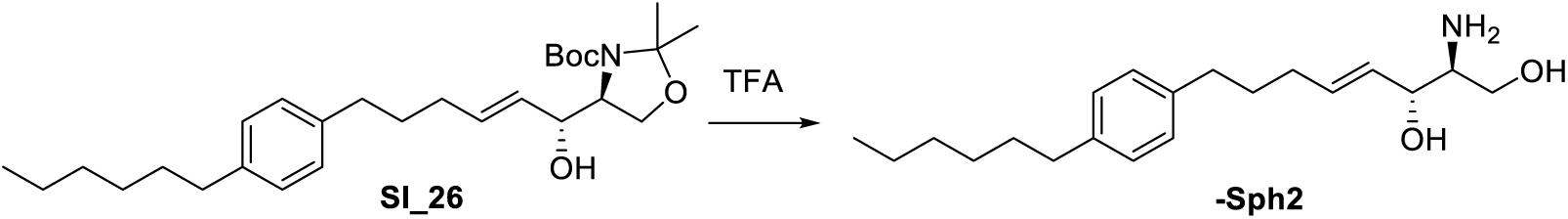

**π-Sph2: SI_26** (200 mg, 0.43 mmol, 1.00 equiv) was dissolved in 15.0 mL DCM. 3.0 mL TFA was added and the resulting solution stirred for 1 hour at ambient temperature, then it was concentrated and directly purified by HPLC (C18, MeCN/H_2_O 42-52% over 16 min, R_t_ = 7.5-8.0 min) to give the title compound **π-Sph2** as a colorless solid (34.5 mg, 0.11 mmol, 25%).

**^1^H NMR** (400 MHz, DMSO-*d*_6_) δ 7.75 (s, 2H), 7.08 (s, 4H), 5.73 (dt, *J* = 13.9, 6.5 Hz, 1H), 5.45 (dd, *J* = 17.6, 5.5 Hz, 2H), 5.14 (t, *J* = 4.7 Hz, 1H), 4.18 (d, *J* = 5.5 Hz, 1H), 3.60 (dt, *J* = 8.1, 4.1 Hz, 1H), 3.52 – 3.41 (m, 1H), 3.03 (s, 1H), 2.55 (d, *J* = 7.6 Hz, 2H), 2.03 (q, *J* = 6.7 Hz, 2H), 1.63 (p, *J* = 7.5 Hz, 2H), 1.52 (d, *J* = 7.3 Hz, 2H), 1.27 (s, 6H), 0.89 – 0.81 (m, 3H).

**LCMS** *m/z* calculated for C_20_H_34_NO_2_^+^ [M+H]^+^: 320.3 found: 320.2.

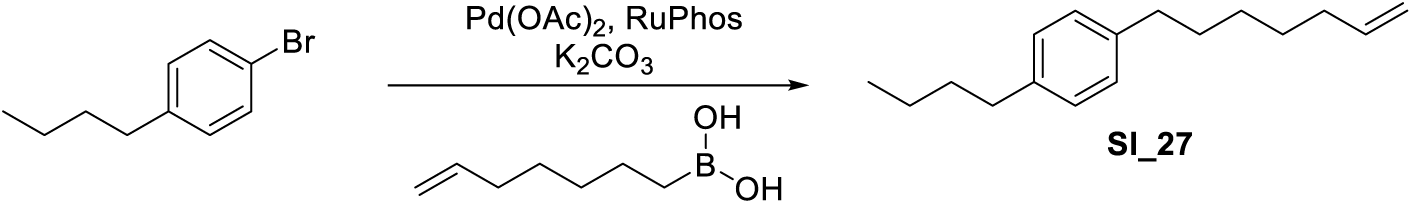

**1-Butyl-4-(hept-6-en-1-yl)benzene (SI_27):** 1-Bromo-4-butylbenzene (2.40 g, 16.9 mmol, 1.00 equiv) was dissolved in a mixture of PhMe and H2O (22 mL, 10:1). 6-heptenylboronic acid (3.00 g, 14.1 mmol, 0.83 equiv), K_2_CO_3_ (5.80 g, 42.2 mmol, 2.50 equiv), Pd(OAc)_2_ (159 mg, 0.70 mmol, 0.042 equiv), and Ruphos (657 mg, 1.76 mmol, 0.083 equiv) were added and the reaction mixture was stirred for 5 h at 80°C. EtOAc and water were added, the phases separated and the aqueous phase twice extracted with EtOAc. The combined organic phases were dried over Na_2_SO_4_, filtered, and concentrated. The mixture was purified by flash column chromatography (hexanes) to yield the title compound **SI_27** as colorless oil (1.20 g, 6.25 mmol, 37%).

**^1^H NMR** (400 MHz, CDCl_3_) δ 7.09 (s, 4H), 5.81 (ddt, *J* = 17.0, 10.1, 6.7 Hz, 1H), 5.04 – 4.90 (m, 1H), 2.58 (td, *J* = 8.0, 2.4 Hz, 4H), 2.05 (q, *J* = 6.9 Hz, 2H), 1.65 – 1.57 (m, 4H), 1.38 (ddq, *J* = 22.2, 14.6, 7.5 Hz, 6H), 0.93 (t, *J* = 7.3 Hz, 3H).

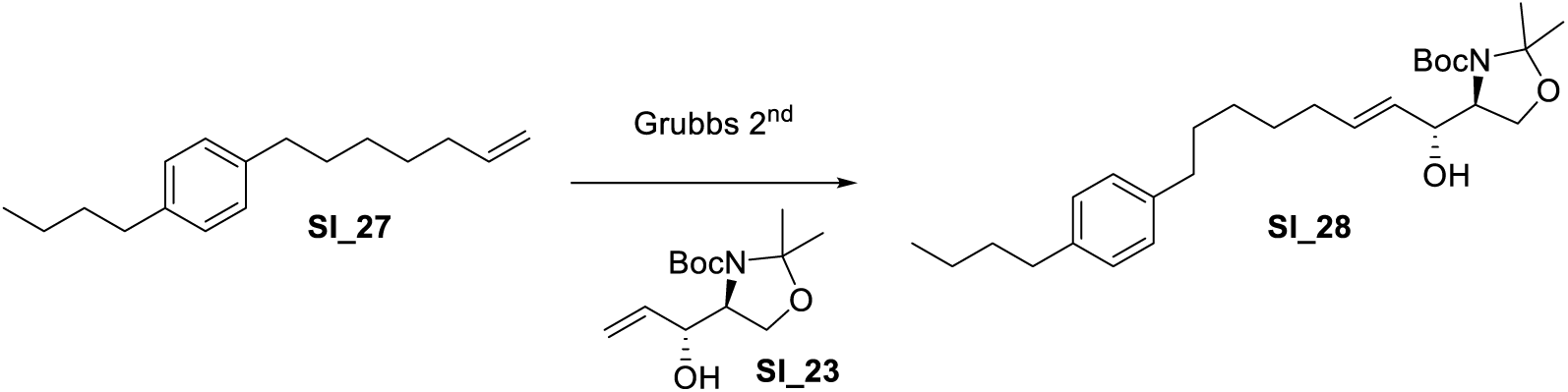

**tert-tert-butyl (S)-4-((R,E)-8-(4-butylphenyl)-1-hydroxyoct-2-en-1-yl)-2,2-dimethyloxa-zolidine-3-carboxylate (SI_28): SI_27** (1.00 g, 4.30 mmol, 1.00 equiv) was dissolved in 10.0 mL CH_2_Cl_2_. The vinyl adduct of Garner’s aldehyde (1.60 g, 6.51 mmol, 1.51 equiv) and HG-II (60 mg, 0.071 mmol, 1.64 mol%) were added and the reaction mixture was stirred for 16 h at 40°C. The reaction mixture was concentrated and purified by flash column chromatography (PE/EA: 0% ∼ 10%) to yield **SI_28** as colorless oil (1.10 g, 2.37 mmol, 40%).

**LCMS** *m/z* calculated for C_27_H_46_NO_4_Na^+^ ([M+Na]^+^): 482.3 found: 482.4.

*Note: **SI_23** was prepared according to the procedure outlined in J. Org. Chem. **1998**, 22, 7999*.

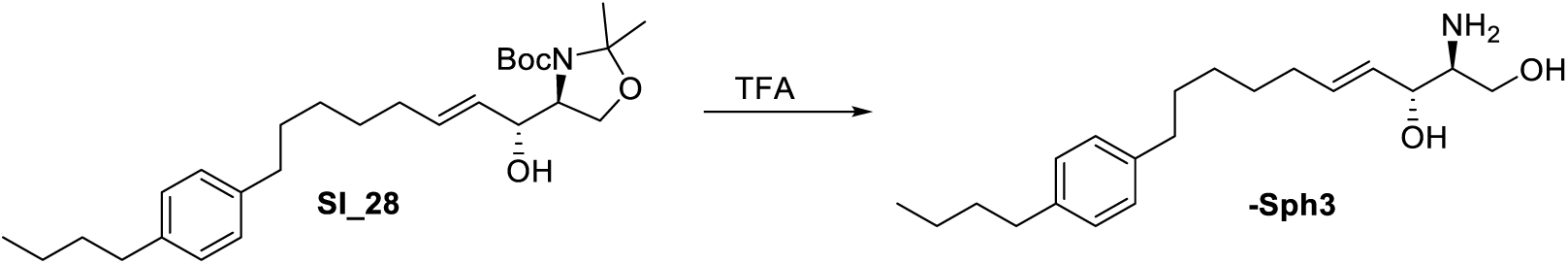

**π-Sph3: SI_28** (200 mg, 0.43 mmol, 1.00 equiv) was dissolved in 15.0 mL DCM. 3.0 mL TFA was added and the resulting solution stirred for 1 hour at ambient temperature, then it was concentrated and directly purified by HPLC (C18, MeCN/H_2_O 46-56% over 16 min, R_t_ = 7.0-8.7 min) to give the title compound **π-Sph3** as a colorless solid (37.2 mg, 0.12 mmol, 27%).

**^1^H NMR** (400 MHz, DMSO-d_6_) δ 7.72 (s, 2H), 7.08 (s, 4H), 5.69 (dt, *J* = 14.0, 6.8 Hz, 1H), 5.43 (dd, *J* = 14.5, 5.5 Hz, 2H), 5.16 – 5.09 (m, 1H), 4.17 (d, *J* = 5.4 Hz, 1H), 3.64 – 3.56 (m, 1H), 3.49 – 3.42 (m, 1H), 3.05 – 2.99 (m, 1H), 2.54 (s, 2H), 2.01 (q, *J* = 6.7 Hz, 2H), 1.52 (dt, *J* = 15.3, 7.4 Hz, 4H), 1.31 (ddt, *J* = 21.8, 14.5, 7.1 Hz, 6H), 0.89 (t, *J* = 7.3 Hz, 3H).

**LCMS** *m/z* calculated for C_20_H_33_NO_2_^+^ ([M+H]^+^): 319.5 found: 320.2.

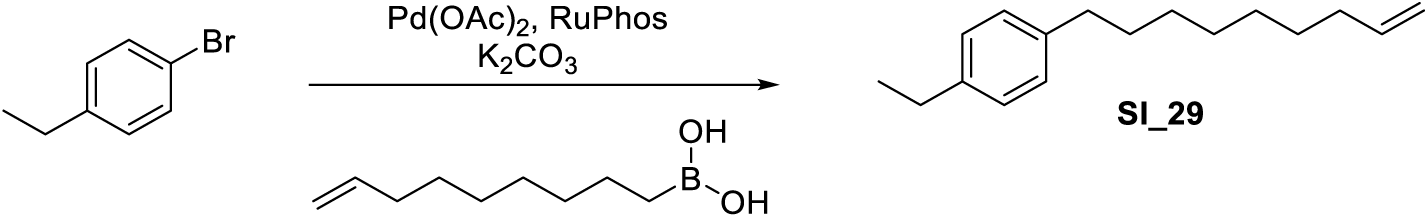

**1-Ethyl-4-(non-8-en-1-yl)benzene (SI_29):** 1-Bromo-4-ethylbenzene (15.7 g, 84.7 mmol, 1.00 equiv) was dissolved in a mixture of PhMe and H_2_O (143 mL, 10:1). 8-nonenylboronic acid (12.0 g, 70.6 mmol, 0.83 equiv), K_2_CO_3_ (29.2 g, 211 mmol, 2.50 equiv), Pd(OAc)_2_ (792 mg, 3.52 mmol, 0.50 equiv), and Ruphos (3.28 g, 8.82 mmol, 0.10 equiv) were added and the reaction mixture was stirred for 5 h at 80°C. EtOAc and water were added, the phases separated and the aqueous phase twice extracted with EtOAc. The combined organic phases were dried over Na_2_SO_4_, filtered, and concentrated. The mixture was purified by flash column chromatography (hexanes) to yield the title compound **SI_29** as colorless oil (13.0 g, 73.7 mmol, 37%).

**^1^H NMR** (400 MHz, CDCl_3_) δ 7.15 – 7.04 (m, 4H), 5.81 (ddt, J = 16.9, 10.2, 6.7 Hz, 1H), 5.04 – 4.89 (m, 2H), 2.67 – 2.53 (m, 4H), 2.04 (q, J = 6.9 Hz, 2H), 1.59 (q, J = 7.3 Hz, 2H), 1.35 (d, J = 24.2 Hz, 8H), 1.23 (t, J = 7.6 Hz, 3H).

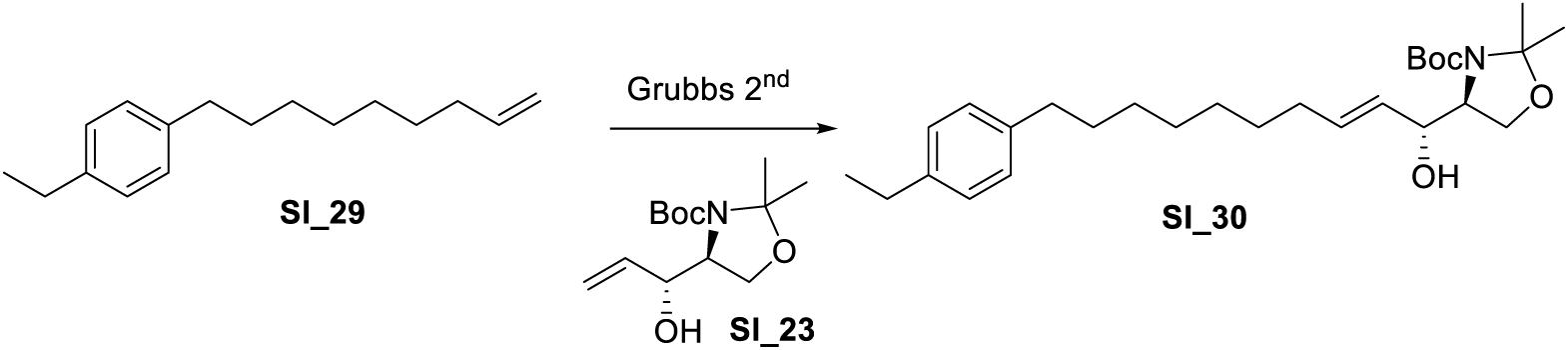

**Tert-butyl (S)-4-((R,E)-10-(4-ethylphenyl)-1-hydroxydec-2-en-1-yl)-2,2-dimethyloxazo-lidine-3-carboxylate (SI_30): SI_29** (2.00 g, 8.70 mmol, 1.00 equiv) was dissolved in 20.0 mL DCM. The vinyl adduct of Garner’s aldehyde **SI_23** (2.00 g, 7.90 mmol, 0.88 equiv) and HG-II (120 mg, catalytic) were added and the reaction mixture stirred at 40 °C for 16 hours. The mixture was concentrated and purified by column chromatography (hexanes/EtOAc 0-10%) to give the title compound **SI_29** as a colorless oil (1.45 g, 3.13 mmol, 36%).

**LCMS** *m/z* calculated for C_27_H_46_NO_4_Na^+^ ([M+Na]^+^): 482.3 found: 482.4.

*Note: **SI_23** was prepared according to the procedure outlined in J. Org. Chem. **1998**, 22, 7999*.

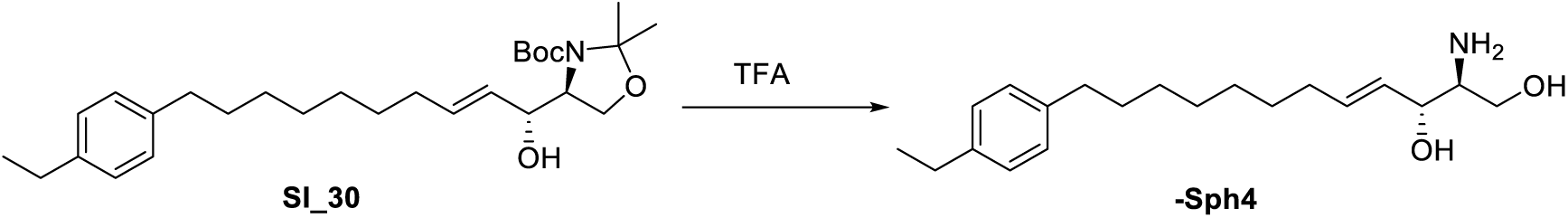

**π-Sph4: SI_30** (300 mg, 0.65 mmol, 1.00 equiv) was dissolved in 10.0 mL DCM. 2.0 mL TFA was added and the resulting solution stirred for 1 hour at ambient temperature, then it was concentrated and directly purified by HPLC (C18, MeCN/H_2_O 43-53% over 16 min, R_t_ = 7.8-10.0 min) to give the title compound **π-Sph4** as a colorless solid (34.3 mg, 0.11 mmol, 17%).

**^1^H NMR** (400 MHz, MeOD) δ 7.11 – 7.02 (m, 4H), 5.90 – 5.78 (m, 1H), 5.47 (dd, J =15.4, 6.9 Hz, 1H), 4.30 – 4.23 (m, 1H), 3.79 (dd, J = 11.6, 4.1 Hz, 1H), 3.65 (dd, J =11.6, 8.3 Hz, 1H), 3.18 (dt, J = 8.6, 4.2 Hz, 1H), 2.64 – 2.51 (m, 4H), 2.10 (q, J = 6.9Hz, 2H), 1.59 (t, J = 6.8 Hz, 2H), 1.45 – 1.38 (m, 2H), 1.34 (d, J = 4.5 Hz, 6H), 1.20 (t,J = 7.6 Hz, 3H).

**LCMS** *m/z* calculated for C_20_H_33_NO_2_^+^ ([M+H]^+^): 319.5 found: 320.2.

### Azido-Stearic Acids

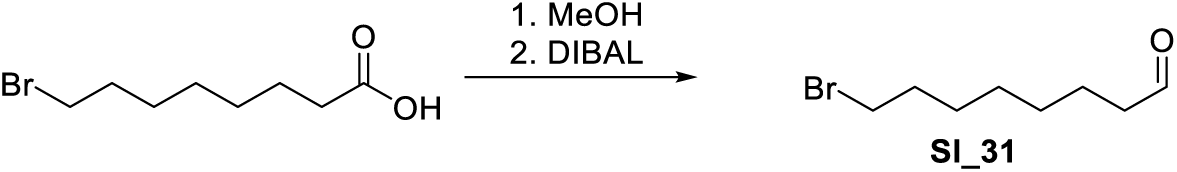

**8-Bromooctanal (SI_31):** 8-Bromooctanoic acid (1.88 g, 8.43 mmol, 1.00 equiv) was dissolved in 25.6 mL MeOH. Dropwise, sulfuric acid (0.23 mL, 4.22 mmol, 0.50 equiv) was added and the resulting solution stirred at ambient temperature for 1 hour. The mixture was then poured into sat. NaHCO_3_ and the resulting mixture extracted 3x with EtOAc. The combined organic phases were dried over Na_2_SO_4_, filtered and evaporated to a crude, which was directly subjected to the next reaction.

Methyl-8-bromooctanoate (assumed quant. 8.43 mmol, 1.00 equiv) was dissolved in 40.0 mL DCM and cooled to – 78 °C. Dropwise, DIBAL (9.0 mL, 9.02 mmol, 1.07 equiv, 1 M solution in PhMe) was added and the resulting mixture stirred for 1 hour at −78 C. The reaction was quenched by the addition of 1.7 mL Methanol to the reaction, then warmed to ambient temperature and poured into 1 M HCl. The reaction mixture was extracted 3x with DCM, the combined organic phases were dried over Na_2_SO_4_, filtered and evaporated to a crude. Purification by column chromatography gave the desired product **SI_31** as a colorless oil (910 mg, 4.39 mmol, 52% over two steps.)

**^1^H NMR** (400 MHz, CDCl_3_) δ 9.76 (t, *J* = 1.8 Hz, 1H), 3.40 (t, *J* = 6.8 Hz, 2H), 2.43 (td, *J* = 7.3, 1.8 Hz, 2H), 1.85 (p, *J* = 6.9 Hz, 2H), 1.69 – 1.55 (m, 2H), 1.50 – 1.40 (m, 2H), 1.34 (p, *J* = 3.6 Hz, 4H).

**^13^C NMR** (101 MHz, CDCl_3_) δ 202.9, 44.0, 34.0, 32.8, 29.1, 28.6, 28.1, 22.1.

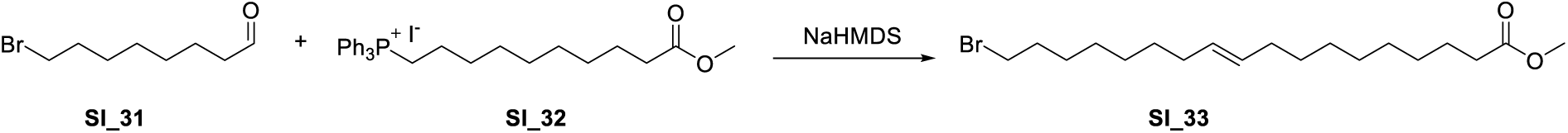

**Methyl (E)-18-bromooctadec-10-enoate (SI_33):** Wittig reagent **SI_32** (505 mg, 0.878 mmol, 1.30 equiv) was azeotroped 3x with PhMe, then dissolved in 5.00 mL THF and cooled to 0 °C. Dropwise, NaHMDS (0.88 mL, 0.878 mmol, 1.30 equiv) was added and the resulting bright orange solution stirred for 10 minutes. The solution containing the ylide was then cooled to −78 °C and a solution of 8-Bromooctanal (**SI_31**, 140.0 mg, 0.676 mmol, 1.00 equiv) in THF (0.7 mL for dissolution, 3x 0.1 mL rinse) was added to the mixture. The reaction was then slowly warmed to ambient temperature over 3 hours, then quenched by pouring the reaction mixture into sat. NH_4_Cl. The resulting mixture was extracted 3x with EtOAc, the combined organic phases were dried over Na_2_SO_4_, filtered and evaporated to a crude. Purification by column chromatography gave desired product as a colorless oil (170 mg, 0.453 mmol, 67%, mixture of E/Z isomers).

**^1^H NMR** (400 MHz, CDCl_3_) δ 5.39 – 5.25 (m, 2H), 3.66 (s, 3H), 3.40 (t, *J* = 6.9 Hz, 2H), 2.30 (t, *J* = 7.5 Hz, 2H), 2.07 – 1.95 (m, *J* = 3.0 Hz, 4H), 1.85 (dt, *J* = 14.3, 6.9 Hz, 2H), 1.60 (dt, *J* = 9.9, 6.9 Hz, 2H), 1.49 – 1.39 (m, 2H), 1.37 – 1.20 (m, 16H).

**^13^C NMR** (101 MHz, CDCl_3_) δ 174.5, 130.1, 129.9, 51.6, 34.24, 34.15, 33.0, 29.9, 29.7, 29.5, 29.4 (2C), 29.3, 29.2, 28.8, 28.3, 27.33, 27.26, 25.1.

*Note: Wittig reagent **SI_32** was prepared according to Helvetica Chimica Acta, **1974**, 57, 434*.

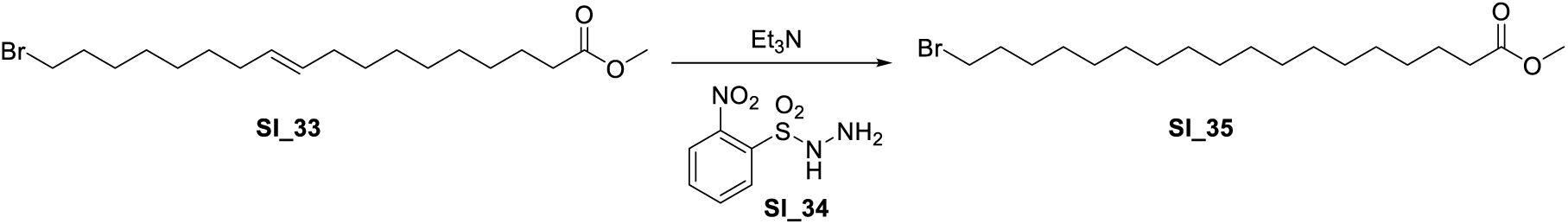

**Methyl 18-bromooctadecanoate (SI_35): SI_33** (100.0 mg, 0.266 mmol, 1.00 equiv) was dissolved in 2.50 mL DCM. Sequentially, Et_3_N (0.37 mL, 2.66 mmol, 10.0 equiv) and Hydrazide **SI_34** (289 mg, 1.33 mmol, 5.00 equiv) were added and the resulting suspension stirred overnight. The mixture was then poured into sat. NaHCO_3_ and the phases separated. The aqueous phase was further extracted 3x with DCM, the combined organic phases were dried over Na_2_SO_4_, filtered and evaporated to a crude. Purification by column chromatography (hexanes/EtOAc) gave desired product **SI_35** as a colorless oil (69.7 mg, 0.19 mmol, 69%).

**^1^H NMR** (400 MHz, CDCl_3_) δ 3.66 (s, 3H), 3.41 (t, *J* = 6.9 Hz, 2H), 2.30 (t, *J* = 7.5 Hz, 2H), 1.85 (p, *J* = 7.0 Hz, 2H), 1.62 (p, *J* = 7.2 Hz, 2H), 1.41 (q, *J* = 7.1 Hz, 2H), 1.26 (d, *J* = 10.3 Hz, 24H).

**^13^C NMR** (101 MHz, CDCl_3_) δ 174.5, 51.6, 34.3, 34.2, 33.0, 29.81, 29.81, 29.80, 29.79, 29.76, 29.74, 29.69, 29.60, 29.59, 29.4, 29.3, 28.9, 28.3, 25.1.

**HRMS** *m/z* calculated for C_19_H_38_BrO_2_ [M+H]^+^: 377.2050, found: 377.2050.

*Note: Hydrazide **SI_34** was prepared according to J. Am. Chem. Soc. **2017**, 139, 15636*.

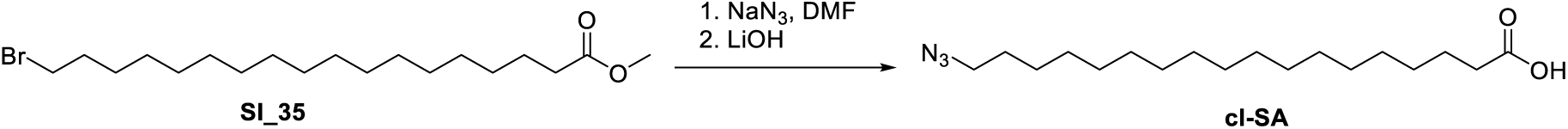

**18-Azido-stearic acid (cl-SA)**: **SI_35** (68 mg, 0.18 mmol, 1.00 equiv) was dissolved in

0.75 mL DMF, followed by the addition of NaN_3_ (59 mg, 0.90 mmol, 5.00 equiv). The resulting mixture was stirred for 24 hours, then poured into sat. NaHCO_3_. The mixture was extracted 3x with EtOAc, the combined organic phases were dried over Na_2_SO_4_, filtered and evaporated to a crude. The crude was azeotroped 3x with heptanes, then directly subjected to the next reaction.

Unpurified 18-azido-stearic acid methyl ester (assumed quant. 0.18 mmol, 1.00 equiv) was dissolved in a mixture of 1.00 mL THF, 0.40 mL MeOH and 0.20 mL water and cooled to 0°C. LiOH·H_2_O (23.0 mg, 0.54 mmol, 3.00 equiv) was added and the mixture slowly allowed to warm to ambient temperature. After stirring for four hours, the reaction mixture was poured into 1 M HCl and the ensuing mixture extracted with DCM 3x. The combined organic phases were dried over Na_2_SO_4_, filtered and evaporated to a solid, which was recrystallized from hexanes to give the desired product **cl-SA** as a white solid (29 mg, 0.089 mmol, 49% over two steps).

**^1^H NMR** (400 MHz, CDCl_3_) δ 10.30 (bs, 1H), 3.25 (t, *J* = 7.0 Hz, 2H), 2.35 (t, *J* = 7.5 Hz, 2H), 1.62 (dp, *J* = 14.4, 7.3 Hz, 4H), 1.27 (d, *J* = 8.2 Hz, 26H).

**^13^C NMR** (101 MHz, CDCl_3_) δ 178.7, 51.7, 33.9, 29.80, 29.78, 29.77, 29.73, 29.69, 29.63, 29.58, 29.4, 29.3, 29.2, 29.0, 26.9, 24.8.

*Note: 2 Resonances corresponding to the alkyl chain are not visible in the ^13^C NMR due to overlap*.

**HRMS** *m/z* calculated for C_19_H_34_N_3_O_2_ [M-H]^-^: 324.2657, found: 324.2657.

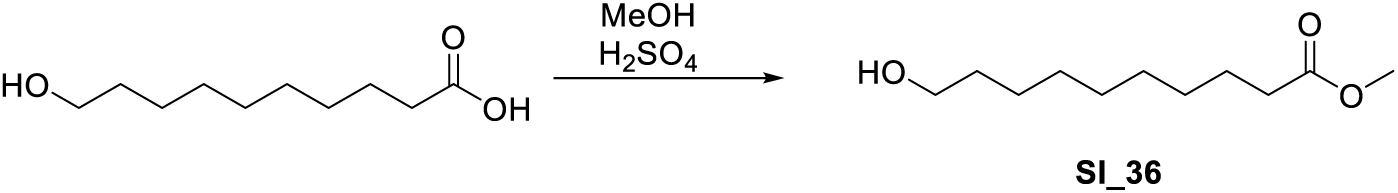

**10-Hydroxydecanoic acid methyl ester (SI_36):** 10-hydroxydecanoic acid (6.00 g, 31.9 mmol, 1.00 equiv) was suspended in 130 mL MeOH and stirred vigorously. Dropwise, H_2_SO_4_ (0.85 mL, 15.9 mmol, 0.50 equiv) was added. The resulting mixture was then heated to reflux for 30 min, cooled to ambient temperature and poured into sat. NaHCO_3_. The mixture was extracted 3x with EtOAc, the combined organic phases were washed with water and brine, dried over Na_2_SO_4_, filtered and evaporated to give **SI_36** (6.27 g, 31.0 mmol, 97%) as a pale yellow oil, which was immediately used in subsequent reactions.

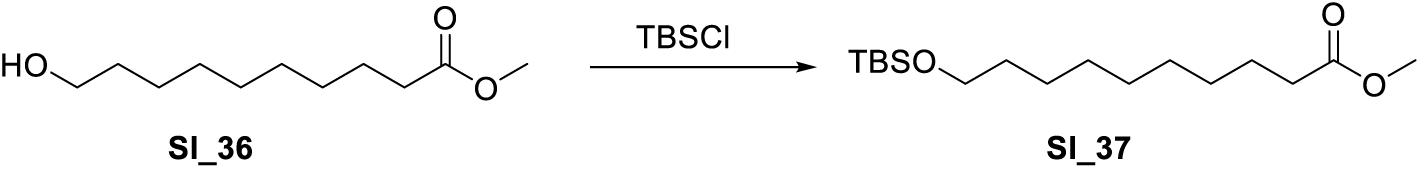

**Methyl 10-((tert-butyldimethylsilyl)oxy)decanoate (SI_37): SI_36** (5.27 g, 26.1 mmol, 1.00 equiv) was dissolved in 60.0 mL DCM and cooled to 0 °C. Sequentially, Imidazole (4.61 g, 67.7 mmol, 2.60 equiv) and TBSCl (5.10 g, 33.9 mmol, 1.30 equiv) was added and the resulting solution stirred for 2 hours, slowly warming to ambient temperature. The reaction was then poured onto sat. NH_4_Cl, the resulting mixture extracted 3x with DCM and the combined organic phases dried over Na_2_SO_4_, filtered and evaporated to a crude. Purification by column chromatography gave desired product **SI_37** as colorless oil (6.80 g, 21.5 mmol, 83%).

**^1^H NMR** (400 MHz, CDCl_3_) δ 3.66 (d, *J* = 0.7 Hz, 3H), 3.59 (t, *J* = 6.6 Hz, 2H), 2.30 (t, *J* = 7.6 Hz, 2H), 1.67 – 1.56 (m, 3H), 1.50 (p, *J* = 7.0 Hz, 2H), 1.29 (d, *J* = 5.5 Hz, 9H), 0.89 (d, *J* = 0.7 Hz, 9H), 0.04 (s, 6H).

**^13^C NMR** (101 MHz, CDCl_3_) δ 174.5, 63.5, 51.6, 34.3, 33.0, 29.57, 29.51, 29.34, 29.28, 26.1, 25.9, 25.1, 18.5, −5.1.

Identical characterization as reported in: *Chemistry and Physics of Lipids*, **1999**, *97*, 87-91.

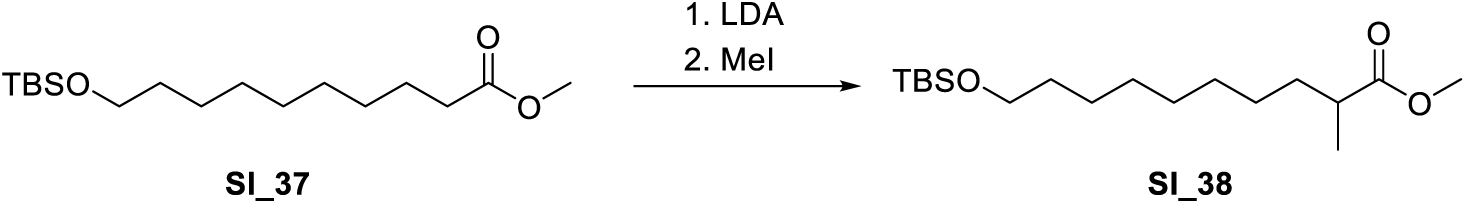

**Methyl 10-((tert-butyldimethylsilyl)oxy)-2-methyldecanoate (SI_38):** A LDA solution was prepared by dissolving *i*-Pr_2_NH (0.98 mL, 6.95 mmol, 1.10 equiv) in 13.0 mL THF and cooling to 0 °C, followed by the dropwise addition of n-BuLi (4.3 mL, 6.95 mmol, 1.10 equiv). After stirring for 20 min, a solution of **SI_37** (2.00 g, 6.32 mmol, 1.00 equiv) in 12 mL THF was added and the resulting mixture stirred at 0 °C for 30 min. Then, MeI (0.43 mL, 6.95 mmol, 1.00 equiv) was added and the reaction stirred for 2 hours, slowly warming to ambient temperature. The mixture was then poured onto sat. NH_4_Cl and the mixture extracted three times with EtOAc. The combined organic phases were dried over Na_2_SO_4_, filtered and evaporated to a crude. Purification by column chromatography (hexanes/EtOAc) gave desired product as a colorless oil (1.35 g, 4.08 mmol, 65%).

**^1^H NMR** (400 MHz, CDCl_3_) δ 3.67 (s, 3H), 3.59 (t, J = 6.6 Hz, 2H), 2.43 (h, J = 7.0 Hz, 1H), 1.71 – 1.60 (m, 2H), 1.56 – 1.44 (m, 2H), 1.28 (d, J = 4.7 Hz, 10H), 1.14 (d, J = 7.0 Hz, 3H), 0.89 (s, 9H), 0.04 (s, 6H).

**^13^C NMR** (101 MHz, CDCl_3_) δ 177.6, 63.5, 51.6, 39.6, 34.0, 33.0, 29.6, 29.5, 27.4, 26.1, 25.9, 18.5, 17.2, −5.1.

Identical characterization as reported in: *Chemistry and Physics of Lipids*, **1999**, *97*, 87-91.

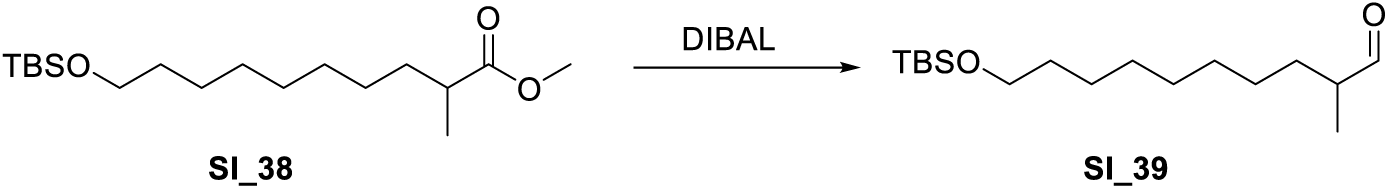

**10-((Tert-butyldimethylsilyl)oxy)-2-methyldecanal (SI_39): SI_38** (260 mg, 0.786 mmol, 1.00 equiv) was dissolved in 7.0 mL DCM and cooled to −78 °C. Dropwise, a solution of DIBAL-H (1.02 mL, 1.02 mmol, 1.30 equiv, 1 M in DCM) was added and the resulting solution stirred for 1 hour. The raction was then quenched by the rapid addition of MeOH at −78 °C. The mixture was then poured into sat. NH_4_Cl, and the phases separated. The aqueous phase was further extracted 3x with DCM, the combined organic phases were dried over Na_2_SO_4_, filtered and evaporated to a crude. Purification by column chromatography (hexanes/EtOAc) gave desired product **SI_39** as a colorless oil (150 mg, 0.499 mmol, 64%).

**^1^H NMR** (400 MHz, CDCl_3_) δ 9.61 (d, *J* = 2.0 Hz, 1H), 3.59 (t, *J* = 6.6 Hz, 2H), 2.33 (hd, *J* = 6.8, 2.0 Hz, 1H), 1.76 – 1.65 (m, 1H), 1.50 (p, *J* = 6.8 Hz, 2H), 1.38 – 1.22 (m, 11H), 1.08 (d, *J* = 7.0 Hz, 3H), 0.89 (s, 9H), 0.04 (s, 6H).

**^13^C NMR** (101 MHz, CDCl_3_) δ 205.6, 63.4, 46.5, 33.0, 30.7, 29.7, 29.6, 29.5, 27.1, 26.1, 25.9, 18.5, 13.5, −5.1.

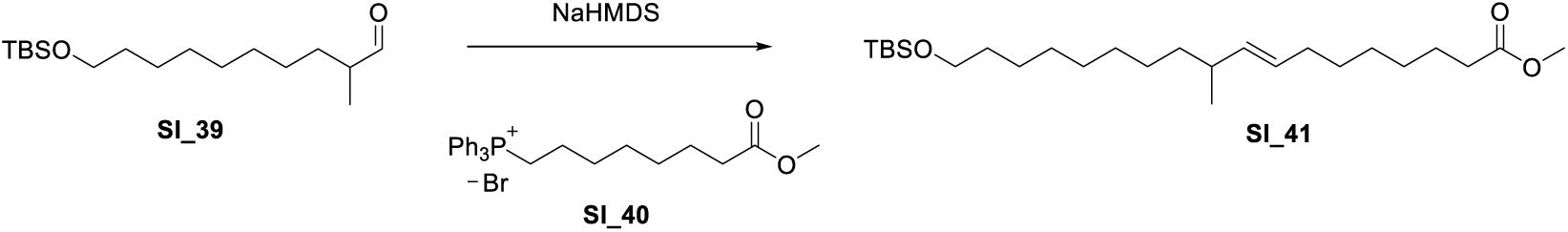

**Methyl (E)-18-((tert-butyldimethylsilyl)oxy)-12-methyloctadec-10-enoate (SI_41):** Wittig reagent **SI_40** (324 mg, 0.649 mmol, 1.30 equiv) was azeotroped 3x with PhMe, then dissolved in 3.00 mL THF and cooled to 0 °C. Dropwise, NaHMDS (0.65 mL, 0.649 mmol, 1.30 equiv) was added and the resulting bright orange solution stirred for 10 minutes. The solution containing the ylide was then cooled to −78 °C and a solution of **SI_39** (150.0 mg, 0.499 mmol, 1.00 equiv) in THF (0.7 mL for dissolution, 3x 0.1 mL rinse) was added to the mixture. The reaction was then slowly warmed to ambient temperature over 3 hours, then quenched by pouring the reaction mixture into sat. NH_4_Cl. The resulting mixture was extracted 3x with EtOAc, the combined organic phases were dried over Na_2_SO_4_, filtered and evaporated to a crude. Purification by column chromatography gave desired product **SI_41** as a colorless oil (165 mg, 0.374 mmol, 75%, mixture of E/Z isomers).

**^1^H NMR** (400 MHz, CDCl_3_) δ 5.32 – 5.21 (m, 1H), 5.10 (ddt, J = 11.0, 9.7, 1.5 Hz, 1H), 3.66 (s, 3H), 3.59 (t, J = 6.6 Hz, 2H), 2.38 (d, J = 9.3 Hz, 1H), 2.30 (t, J = 7.6 Hz, 2H), 2.06 – 1.96 (m, 2H), 1.72 – 1.57 (m, 2H), 1.49 (q, J = 6.8 Hz, 2H), 1.41 – 1.18 (m, 18H), 0.91 (d, J = 6.9 Hz, 3H), 0.89 (s, 9H), 0.04 (s, 6H).

**^13^C NMR** (101 MHz, CDCl_3_) δ 174.5, 136.7, 128.2, 63.5, 51.6, 37.7, 34.3, 33.0, 31.8, 29.94, 29.87, 29.8, 29.6, 29.2, 29.1, 27.7, 27.5, 26.1, 26.0, 25.1, 21.6, 18.5, −5.1.

**HRMS** *m/z* calculated for C_26_H_53_O_3_Si [M+H]^+^: 441.3759, found: 441.3758.

*Note: Wittig reagent **SI_40** was prepared according to: J. Org. Chem. **2001**, 66, 7765*.

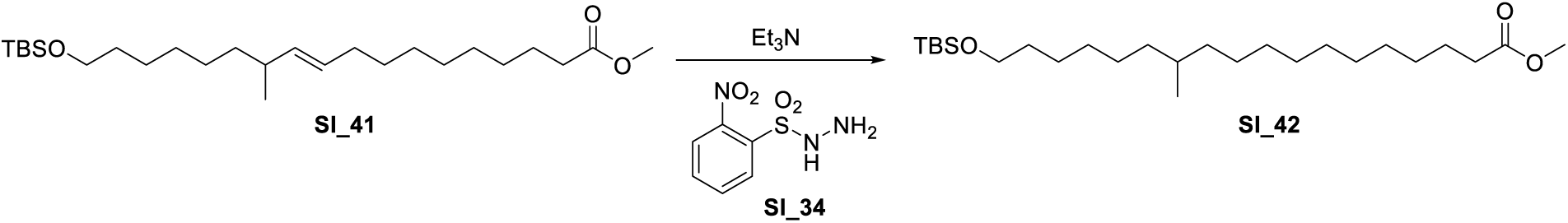

**Methyl 18-((tert-butyldimethylsilyl)oxy)-10-methyloctadecanoate (SI_42): SI_41** (90.0 mg, 0.204 mmol, 1.00 equiv) was dissolved in 1.60 mL DCM. Sequentially, Et_3_N (0.29 mL, 2.04 mmol, 10.0 equiv) and Hydrazide **SI_34** (222 mg, 1.02 mmol, 5.00 equiv) were added and the resulting suspension stirred overnight. The mixture was then poured into sat. NaHCO_3_ and the phases separated. The aqueous phase was further extracted 3x with DCM, the combined organic phases were dried over Na_2_SO_4_, filtered and evaporated to a crude. Purification by column chromatography (hexanes/EtOAc) gave desired product **SI_42** as a colorless oil (75 mg, 0.17 mmol, 83%).

**^1^H NMR** (400 MHz, CDCl_3_) δ 3.66 (s, 3H), 3.59 (t, *J* = 6.6 Hz, 2H), 2.30 (t, *J* = 7.6 Hz, 2H), 1.61 (q, *J* = 7.1 Hz, 2H), 1.51 (h, *J* = 6.5 Hz, 2H), 1.38 – 1.18 (m, 23H), 1.16 – 0.97 (m, 2H), 0.89 (s, 9H), 0.83 (d, *J* = 6.5 Hz, 3H), 0.04 (s, 6H).

**^13^C NMR** (101 MHz, CDCl_3_) δ 174.4, 63.4, 51.5, 37.1, 34.1, 32.9, 32.8, 29.98, 29.96, 29.7, 29.51, 29.48, 29.3, 29.2, 27.09, 27.07, 26.0, 25.8, 25.0, 19.7, 18.4, −5.2.

**HRMS** *m/z* calculated for C_26_H_55_O_3_Si [M+H]^+^: 443.3915, found: 443.3922.

*Note: Hydrazide **SI_34** was prepared according to J. Am. Chem. Soc. **2017**, 139, 15636*.

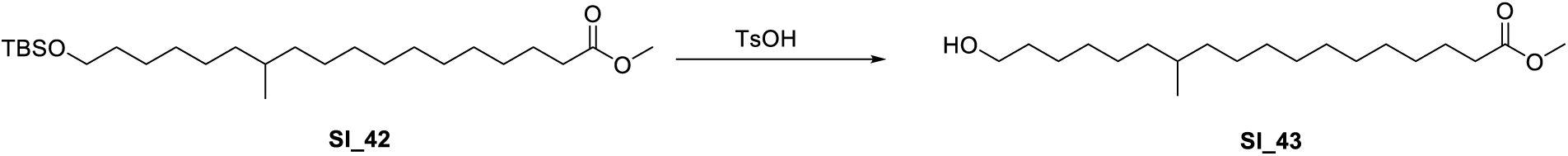

**Methyl 18-hydroxy-10-methyloctadecanoate (SI_43): SI_42** (75 mg, 0.17 mmol,1.00 equiv) was dissolved in 2.00 mL MeOH. Subsequently, pTsOH (32 mg, 0.17 mmol, 1.00 equiv) was added and the resulting mixture stirred at ambient temperature for 2 hours. The reaction was then poured into sat. NaHCO_3_, and the mixture extracted 3x with EtOAc. The combined organic phases were dried over Na_2_SO_4_, filtered and evaporated to a crude. Purification by column chromatography (hexanes/EtOAc) gave the desired product **SI_43** as a colorless oil (51 mg, 0.16 mmol, 91%).

**^1^H NMR** (400 MHz, CDCl_3_) δ 3.64 (s, 3H), 3.61 (t, J = 6.7 Hz, 2H), 2.28 (t, J = 7.5 Hz, 2H), 1.65 – 1.50 (m, 4H), 1.25 (m, 24H), 1.05 (dt, J = 11.3, 8.1 Hz, 2H), 0.81 (d, J = 6.5 Hz, 3H).

**^13^C NMR** (101 MHz, CDCl_3_) δ 174.5, 63.1, 51.6, 37.2, 34.2, 32.9, 32.8, 30.0, 29.8, 29.59, 29.56, 29.4, 29.3, 27.2, 27.1, 25.9, 25.1, 19.8.

**HRMS** *m/z* calculated for C_20_H_41_O_3_ [M+H]^+^: 329.3050, found: 329.3050.

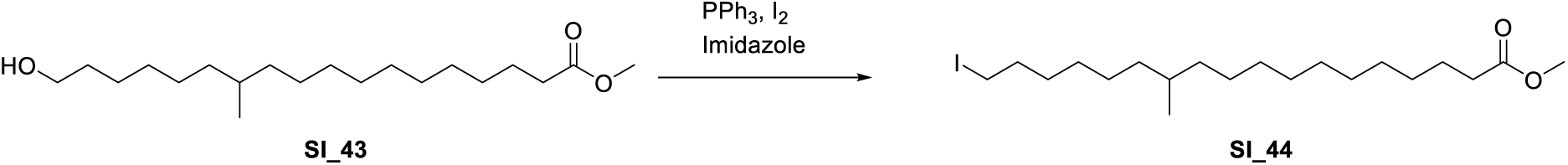

**Methyl 18-iodo-10-methyloctadecanoate (SI_44): SI_43** (51 mg, 0.16 mmol, 1.00 equiv) was dissolved in 0.40 mL DCM and cooled to 0 °C. Sequentially, imidazole (32 mg, 0.47 mmol, 3.00 equiv), PPh_3_ (61 mg, 0.23 mmol, 1.50 equiv) and I_2_ (59 mg, 0.23 mmol, 1.50 equiv) were added. After stirring at 0 °C for two hours, the reaction was quenched by the simultaneous addition of sat. NaHCO_3_ and sat. Na_2_S_3_O_3_. The ensuing mixture was extracted 3x with DCM, the combined organic phases were dried over Na_2_SO_4_, filtered and evaporated to a crude. Purification by column chromatography gave desired product **SI_44** as a colorless oil (65 mg, 0.16 mmol, 96%).

**^1^H NMR** (400 MHz, CDCl_3_) δ 3.65 (s, 3H), 3.18 (t, *J* = 7.1 Hz, 2H), 2.29 (t, *J* = 7.6 Hz, 2H), 1.81 (p, *J* = 7.1 Hz, 2H), 1.61 (p, *J* = 7.3 Hz, 2H), 1.43 – 1.16 (m, 24H), 1.06 (dt, *J* = 11.3, Hz, 2H), 0.82 (d, *J* = 6.5 Hz, 3H).

**^13^C NMR** (101 MHz, CDCl_3_) δ 174.4, 51.6, 37.18, 37.17, 34.2, 33.7, 32.9, 30.6, 30.1, 30.0, 29.62, 29.58, 29.4, 29.3, 28.7, 27.2, 27.1, 25.1, 19.8, 7.4.

**HRMS** *m/z* calculated for C_20_H_41_IO_2_ [M+H]^+^: 439.2068, found: 439.2057.

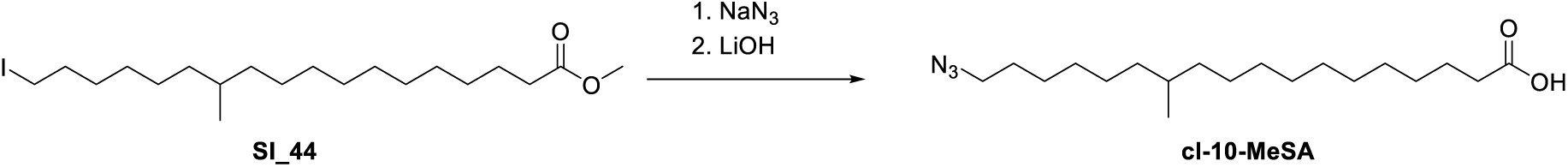

**18-azido-10-methyloctadecanoic acid (cl-10-MeSA): SI_44** (65 mg, 0.15 mmol, 1.00 equiv) was dissolved in 0.80 mL DMF. NaN_3_ (48 mg, 0.74 mmol, 5.00 equiv) was added and the resulting solution stirred for 16 hours. The reaction was then poured into sat. NaHCO_3_, and the ensuing mixture extracted 3x with EtOAc. The combined organic phases were dried over Na_2_SO_4_, filtered and evaporated to a crude, which immediately subjected to the next reaction:

Crude Methyl 18-azido-10-methyloctadecanoate (0.15 mmol, 1.00 equiv) was dissolved in a mixture of 1.0 mL THF, 0.4 mL MeOH and 0.2 mL water and cooled to 0 °C. LiOH·H_2_O (19 mg, 0.44 mmol, 3.00 equiv) was then added and the reaction mixture warmed to ambient temperature. After stirring for 16 hours the reaction was poured into 1 M HCl and the mixture extracted 5x with DCM. The combined organic phases were dried over Na_2_SO_4_, filtered and evaporated to a crude. Purification by column chromatography followed by preparative TLC (both hexanes/EtOAc with 1% AcOH) gave desired product **cl-10-MeSA** as a colorless oil (25 mg, 0.074 mmol, 50% over two steps).

**^1^H NMR** (400 MHz, CDCl_3_) δ 9.90 (s, 1H), 3.25 (t, *J* = 7.0 Hz, 2H), 2.35 (t, *J* = 7.5 Hz, 2H), 1.77 – 1.49 (m, 4H), 1.39 – 1.20 (m, 23H), 1.07 (dt, *J* = 11.5, 8.1 Hz, 2H), 0.83 (d, *J* = 6.5 Hz, 3H).

**^13^C NMR** (101 MHz, CDCl_3_) δ 180.1, 51.6, 37.2, 32.9, 30.1, 30.0, 29.7, 29.6, 29.4, 29.3, 29.2, 29.0, 27.19, 27.17, 26.9, 24.8, 19.9.

**HRMS** *m/z* calculated for C_19_H_36_N_3_O_2_ [M-H]^-^: 338.2813, found: 338.2821.

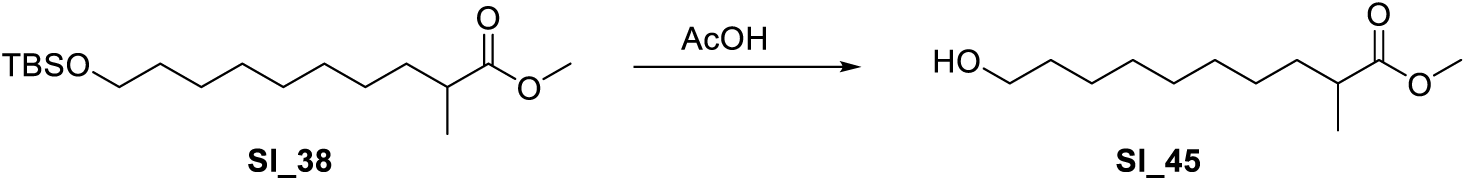

**Methyl 10-hydroxy-2-methyldecanoate (SI_45): SI_38** (1.04 g, 3.15 mmol, 1.00 equiv) was dissolved in a mixture of 10.0 mL AcOH, 3.30 mL THF and 3.30 mL Water and stirred at ambient temperature for 4 hours. Then, the reaction was poured into sat. NaHCO_3_, and the ensuing mixture extracted 3x with DCM. The combined organic phases were dried over Na_2_SO_4_, filtered and evaporated to a crude. Purification by column chromatography (hexanes/EtOAc) gave desired product **SI_45** as a clear oil (0.107 g, 0.495 mmol, 16%).

**^1^H NMR** (400 MHz, CDCl_3_) δ 3.66 (s, 3H), 3.63 (t, *J* = 6.6 Hz, 2H), 2.49 – 2.36 (m, 1H), 1.67 – 1.50 (m, 4H), 1.41 – 1.22 (m, 10H), 1.13 (d, *J* = 7.0 Hz, 3H).

**^13^C NMR** (101 MHz, CDCl_3_) δ 177.6, 63.2, 51.6, 39.6, 33.9, 32.9, 29.5, 29.50, 29.45, 27.3, 25.8, 17.2.

**HRMS** *m/z* calculated for C_12_H_24_NaO_3_ [M+Na]^+^: 239.1618, found: 239.1614.

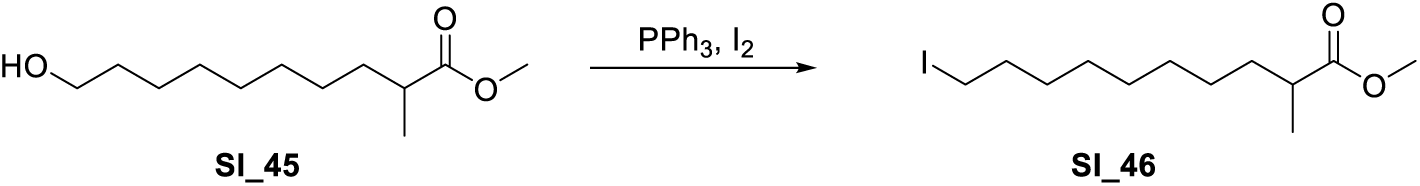

**Methyl 10-iodo-2-methyldecanoate SI_46:** Methyl 10-hydroxy-2-methyldecanoate (0.107 g, 0.495 mmol, 1.00 equiv) was dissolved in 5.0 mL DCM and cooled to 0 °C. Sequentially, imidazole (101 mg, 1.48 mmol, 3.00 equiv), PPh_3_ (143 mg, 0.544 mmol, 1.10 equiv) and iodine (138 mg, 0.544 mmol, 1.10 equiv) were added and the resulting solution stirred for 2 hours at 0 °C. The reaction was then quenched by the simultaneous addition of sat. Na_2_S_2_O_3_ and NaHCO_3_. The phases were separated and the aqueous phase further extracted 3x with DCM. The combined organic phases were dried over Na_2_SO_4_, filtered and evaporated to a crude. Purification by column chromatography (hexanes/EtOAc) gave desired product **SI_46** as a colorless oil (160 mg, 0.490 mmol, 99%).

**^1^H NMR** (400 MHz, CDCl_3_) δ 3.62 (s, 3H), 3.14 (t, *J* = 7.0 Hz, 2H), 2.38 (h, *J* = 7.0 Hz, 1H), 1.77 (p, *J* = 7.1 Hz, 2H), 1.65 – 1.52 (m, 1H), 1.34 (td, *J* = 8.3, 4.5 Hz, 3H), 1.27 (s, 8H), 1.09 (d, *J* = 7.0 Hz, 3H).

**^13^C NMR** (101 MHz, CDCl_3_) δ 177.3, 51.5, 39.5, 33.8, 33.5, 30.5, 29.4, 29.3, 28.5, 27.2, 17.1, 7.3.

**HRMS** *m/z* calculated for C_12_H_24_IO_2_ [M+H]^+^: 327.0816, found: 327.0811.

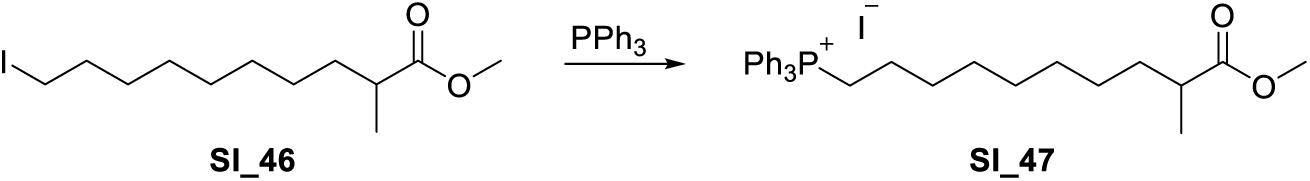

**Methyl 10-(iodotriphenyl-l5-phosphaneyl)-2-methyldecanoate SI_47: SI_46** (160 mg, 0.49 mmol, 1.00 equiv) was dissolved in 2.00 mL PhMe. PPh_3_ (180 mg, 0.687 mmol, 1.40 equiv) was added and the solution heated to reflux for 24 hours. The mixture was then cooled to ambient temperature and volatiles were removed under vacuo. The residue was dissolved in MeCN, and extracted 10x with hexanes, discarding the hexane layer. The MeCN solution was then evaporated, giving the wittig reagent **SI_47** as a colorless syrup (207 mg, 0.352 mmol, 72%) which was immediately employed in the subsequent reaction:

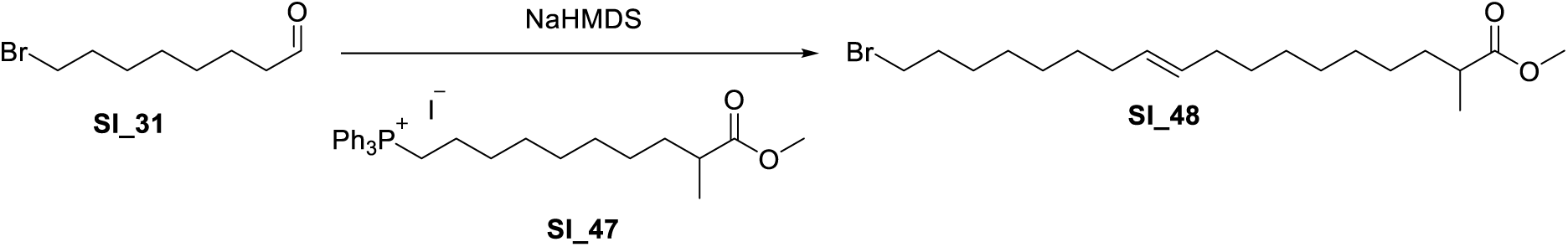

**Methyl (E)-18-bromo-2-methyloctadec-10-enoate (SI_48): SI_47** (207 mg, 0.352 mmol, 1.30 equiv) was azeotroped with PhMe (3x), then dissolved in 2.00 mL THF and cooled to 0 °C. Dropwise, NaHMDS (0.35 mL, 0.35 mmol, 1.30 equiv, 1M in THF) was added and the resulting bright orange solution stirred for 10 minutes, then cooled to −78 °C. Dropwise, a solution of **SI_31** (56 mg, 0.27 mmol, 1.00 equiv) in 1.0 mL THF (0.7 mL for dissolution, 3x 0.1 mL rinse) was added and the resulting solution stirred for 1 hour at −78 °C, then the cooling bath was removed and the solution slowly warmed to ambient temperature and further stirred for 1 hour. The reaction was then quenched by pouring into sat. NH_4_Cl and the ensuing mixture was extracted 3x with EtOAc. The combined organic phases were dried over Na_2_SO_4_, filtered and evaporated to a crude. Purification by column chromatography (hexanes/EtOAc) gave desired product **SI_48** as a colorless oil (66 mg, 0.17 mmol, 63%).

**^1^H NMR** (400 MHz, CDCl_3_) δ 5.45 – 5.27 (m, 2H), 3.66 (s, 3H), 3.40 (t, *J* = 6.9 Hz, 2H), 2.43 (h, *J* = 7.0 Hz, 1H), 2.01 (th, *J* = 5.6, 2.9 Hz, 4H), 1.91 – 1.81 (m, 2H), 1.71 – 1.55 (m, 1H), 1.42 (qd, *J* = 6.2, 3.1 Hz, 2H), 1.38 – 1.23 (m, 17H), 1.14 (d, *J* = 7.0 Hz, 3H).

**^13^C NMR** (101 MHz, CDCl_3_) δ 177.6, 130.1, 129.9, 51.6, 39.6, 34.2, 34.0, 33.0, 29.9, 29.8, 29.6, 29.5, 29.4, 29.2, 28.8, 28.3, 27.38, 27.35, 27.27, 17.2.

**HRMS** *m/z* calculated for C_20_H_38_BrO_2_ [M+H]^+^: 389.2950, found: 389.2050.

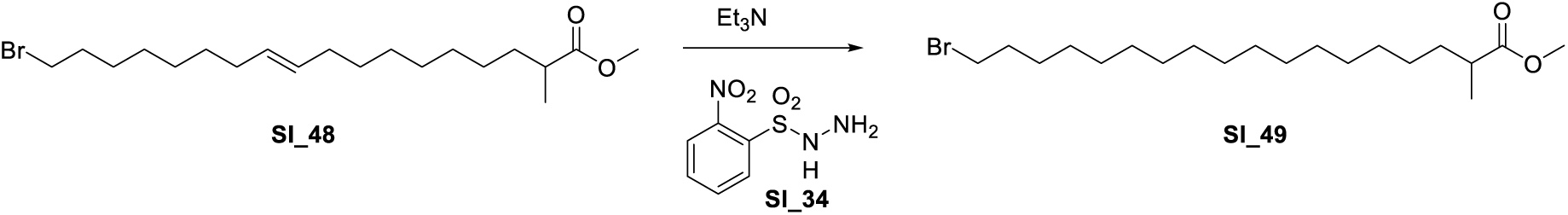

**Methyl 18-bromo-2-methyloctadecanoate (SI_49): SI_48** (66 mg, 0.17 mmol, 1.00 equiv) was dissolved in 1.50 mL DCM. Et_3_N (0.24 mL, 1.70 mmol, 10.0 equiv) and Hydrazide **SI_34** (184 mg, 0.85 mmol, 5.00 equiv) were added and the reaction left stirring overnight. The reaction was poured into sat NaHCO_3_ and the ensuing mixture extracted 3x with DCM. The combined organic phases were dried over Na_2_SO_4_, filtered and evaporated to a crude. Purification by column chromatography gave the desired product **SI_49** as a colorless oil (52 mg, 0.13 mmol, 78%).

**^1^H NMR** (400 MHz, CDCl_3_) δ 3.65 (s, 3H), 3.39 (t, *J* = 6.9 Hz, 2H), 2.42 (h, *J* = 7.0 Hz, 1H), 1.84 (dt, *J* = 14.6, 7.0 Hz, 2H), 1.72 – 1.56 (m, 1H), 1.39 (tq, *J* = 13.0, 6.6 Hz, 3H), 1.25 (d, *J* = 3.5 Hz, 24H), 1.13 (d, *J* = 6.9 Hz, 3H).

**^13^C NMR** (101 MHz, CDCl_3_) δ 177.5, 51.6, 39.6, 34.2, 34.0, 33.0, 29.79, 29.77, 29.75, 29.72, 29.68, 29.65, 29.62, 29.57, 28.9, 28.3, 27.4, 17.2.

**HRMS** *m/z* calculated for C_20_H_40_BrO_2_ [M+H]^+^: 391.2206, found: 391.2204.

*Note: Hydrazide **SI_34** was prepared according to J. Am. Chem. Soc. **2017**, 139, 15636*.

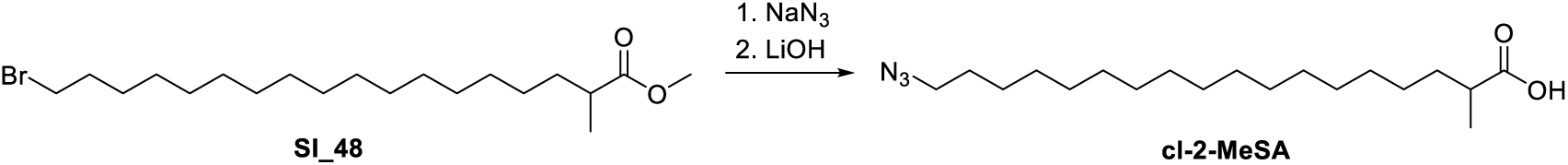

**18-Azido-2-methyloctadecanoic acid (cl-2-MeSA): SI_48** (52 mg, 0.133 mmol, 1.00 equiv) was dissolved in 0.65 mL DMF, followed by the addition of NaN_3_ (43.2 mg, 0.66 mmol, 5.00 equiv). The reaction was stirred for 24 hours, then poured into sat. NaHCO_3_ and the mixture extracted 3x with EtOAc. The combined organic phases were dried over Na_2_SO_4_, filtered and evaporated to a crude, which was directly dissolved in a mixture of 1.0 mL THF, 0.4 mL MeOH, 0.2 mL water and cooled to 0 °C. LiOH·H_2_O (16.7 mg, 0.40 mmol, 3.00 equiv) was added and the reaction allowed to warm to ambient temperature and stirred for 4 hours. The reaction was then poured into 1 M HCl and the mixture extracted 5x with DCM. The combined organic phases were dried over Na_2_SO_4_, filtered and evaporated to a crude, which was purified by crystallization from hexanes to give **cl-2-MeSA** as a white solid (19 mg, 0.056 mmol, 42% over two steps).

**^1^H NMR** (400 MHz, CDCl_3_) δ 3.25 (t, *J* = 7.0 Hz, 2H), 2.46 (h, *J* = 6.9 Hz, 1H), 1.73 – 1.53 (m, 3H), 1.45 – 1.22 (m, 28H), 1.17 (d, *J* = 6.9 Hz, 3H).

**^13^C NMR** (101 MHz, CDCl_3_) δ 182.6, 51.7, 33.7, 29.81, 29.77, 29.75, 29.70, 29.67, 29.63, 29.3, 29.0, 27.3, 26.9, 17.0.

**HRMS** *m/z* calculated for C_19_H_36_N_3_O_2_ [M-H]^-^: 338.2813, found: 338.2830.

## NMR Data

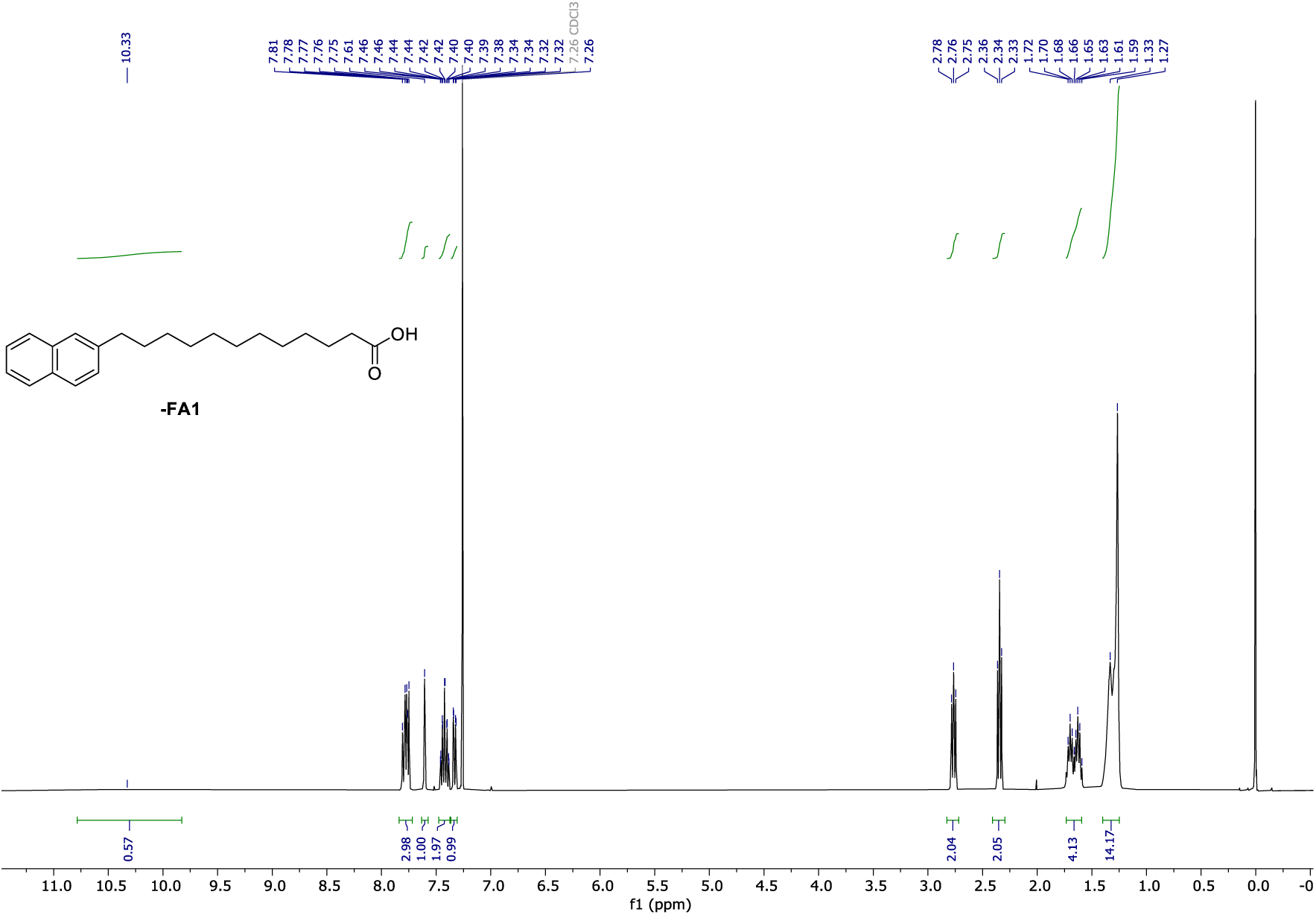

**^1^H NMR** (400 MHz, CDCl_3_) of **π-FA1**.

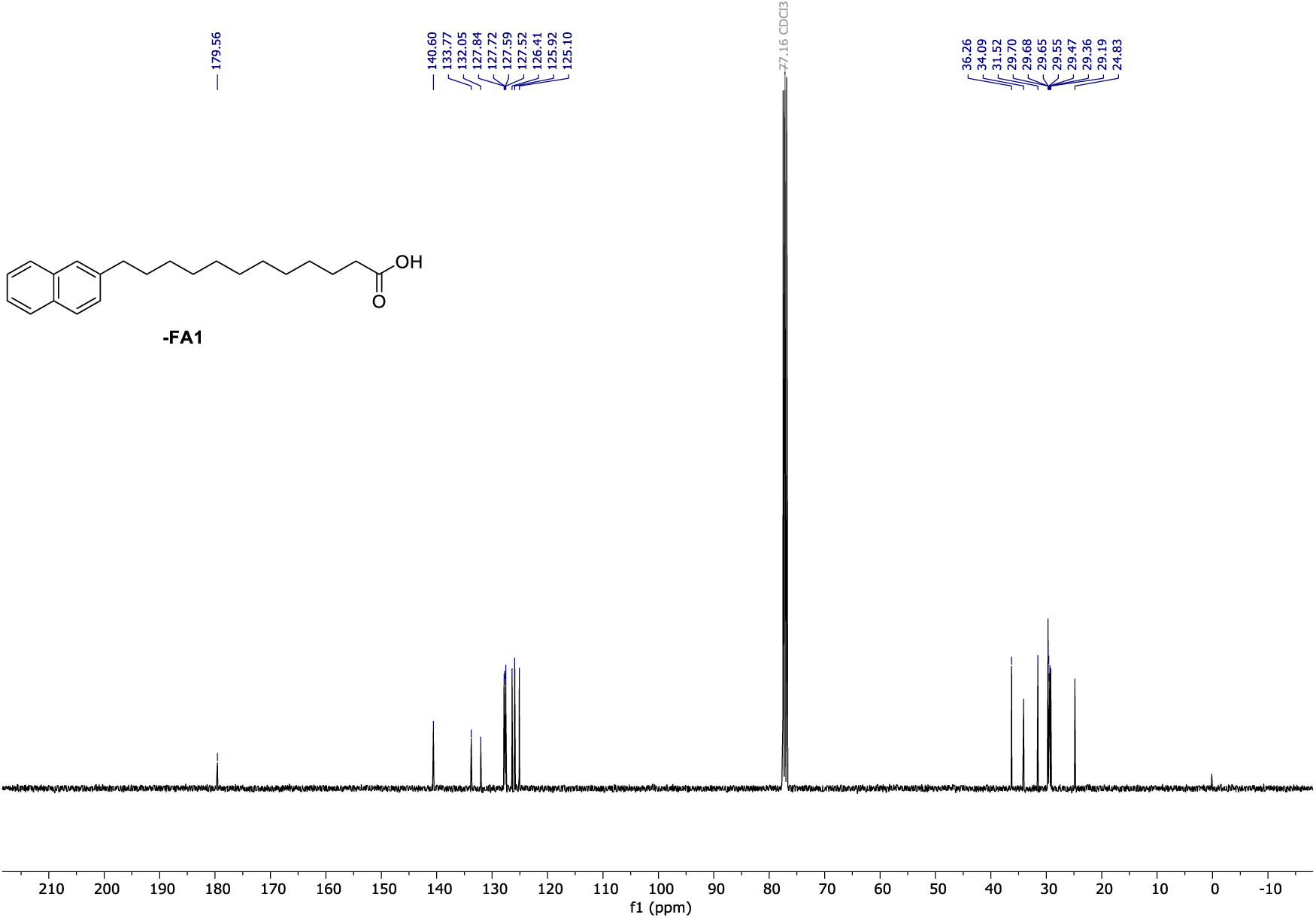

**^13^C NMR** (101 MHz, CDCl_3_) of **π-FA1**.

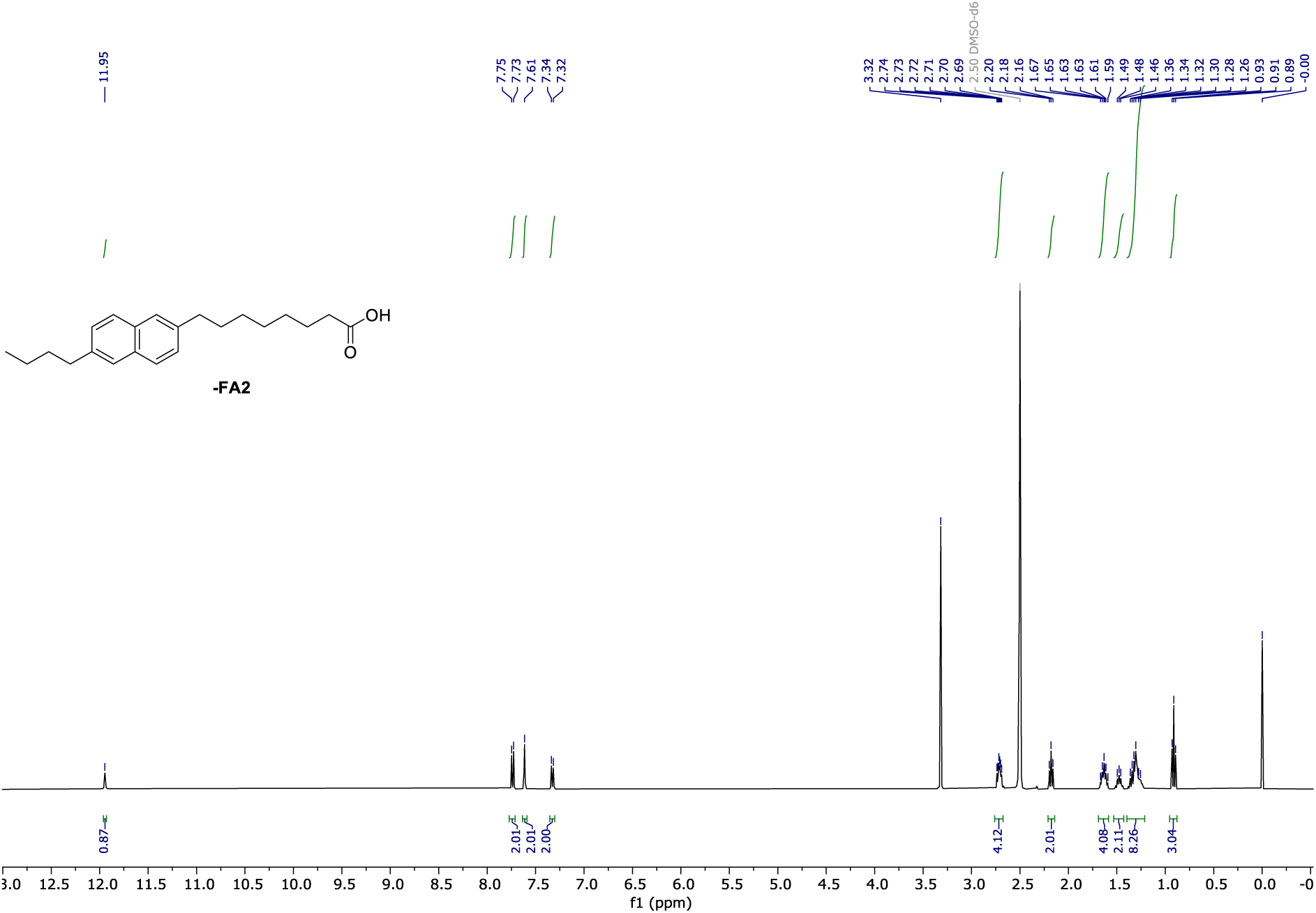

**^1^H NMR** (400 MHz, CDCl_3_) of **π-FA2**.

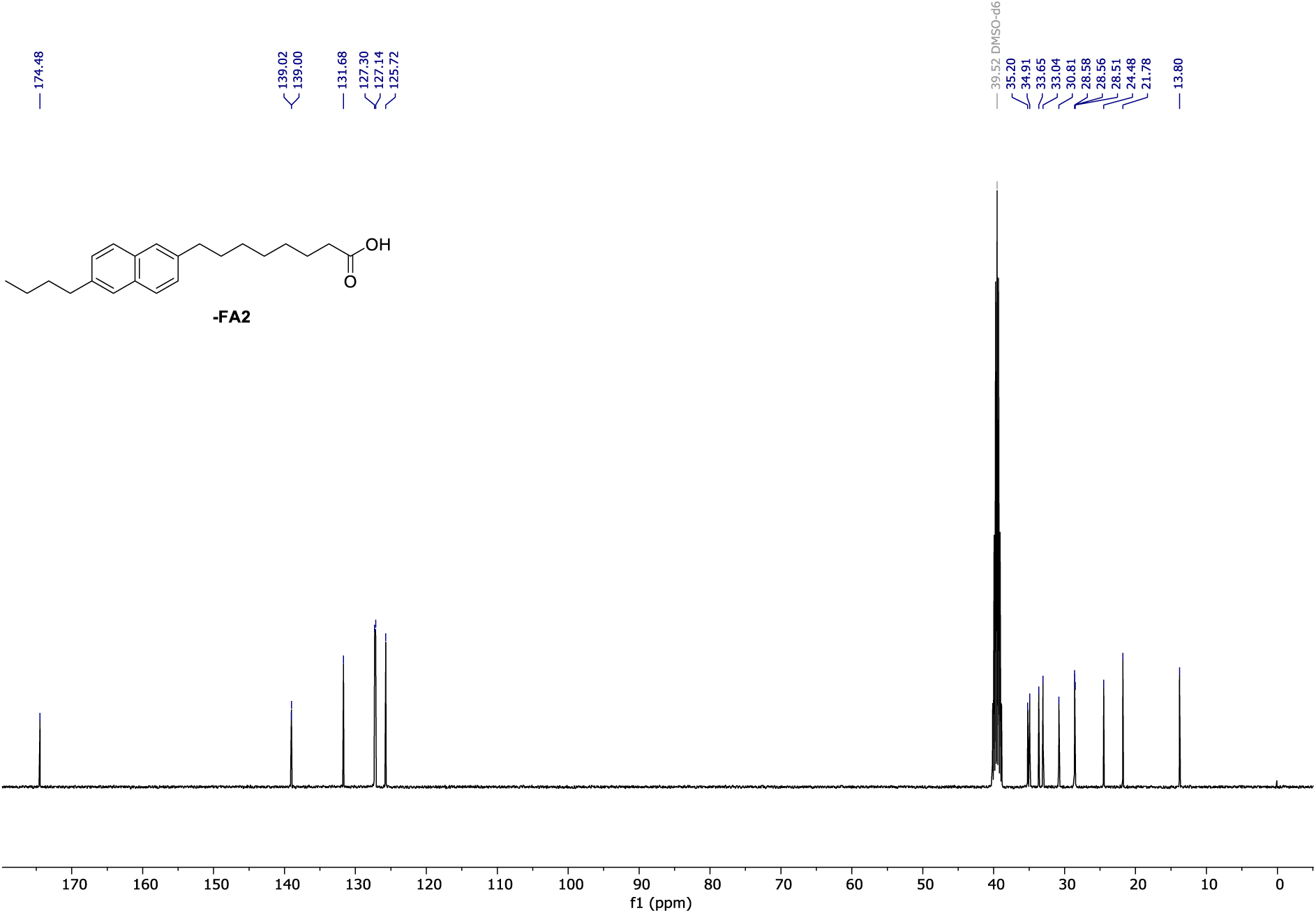

**^13^C NMR** (101 MHz, CDCl_3_) of **π-FA2**.

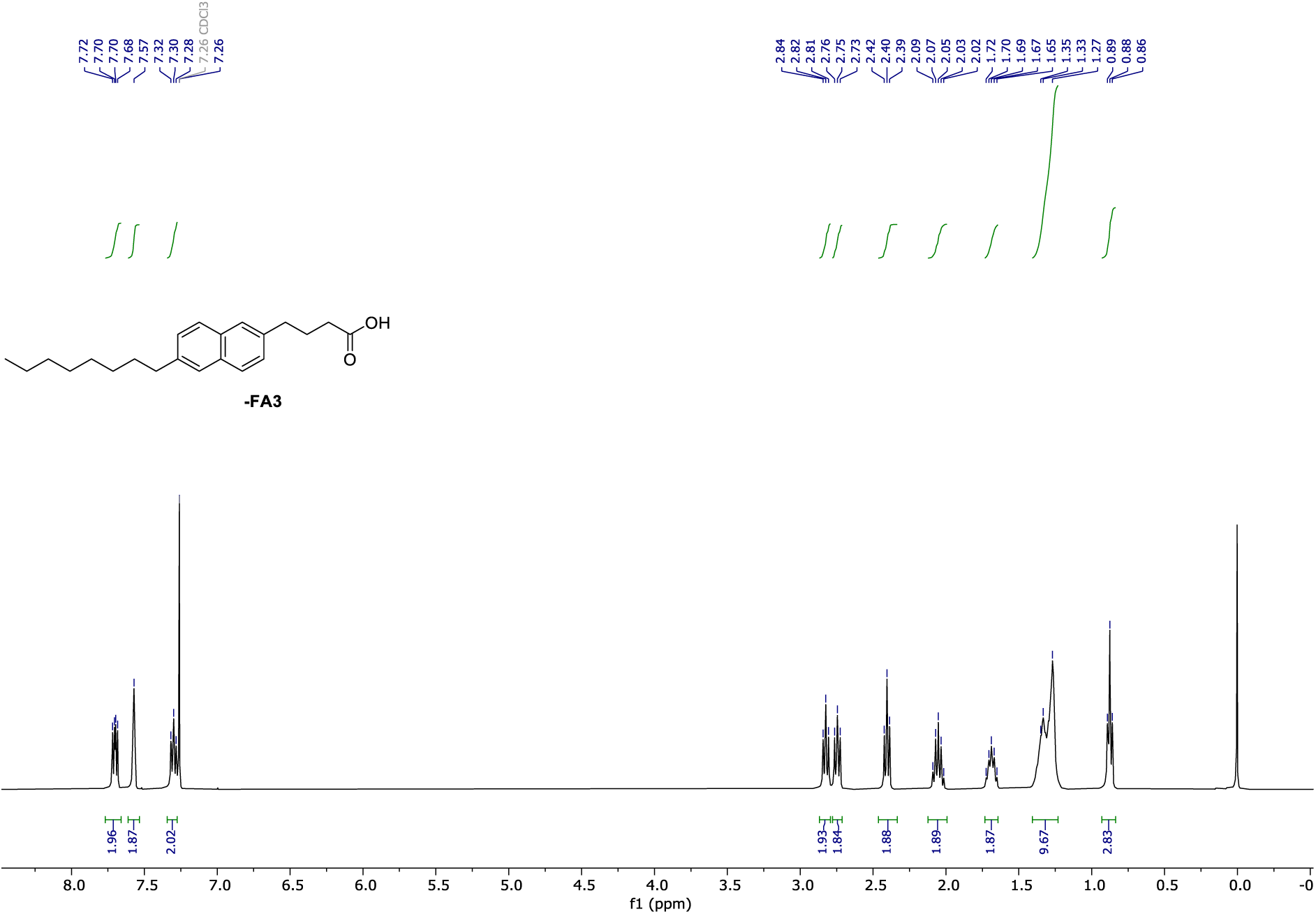

**^1^H NMR** (400 MHz, CDCl_3_) of **π-FA3**.

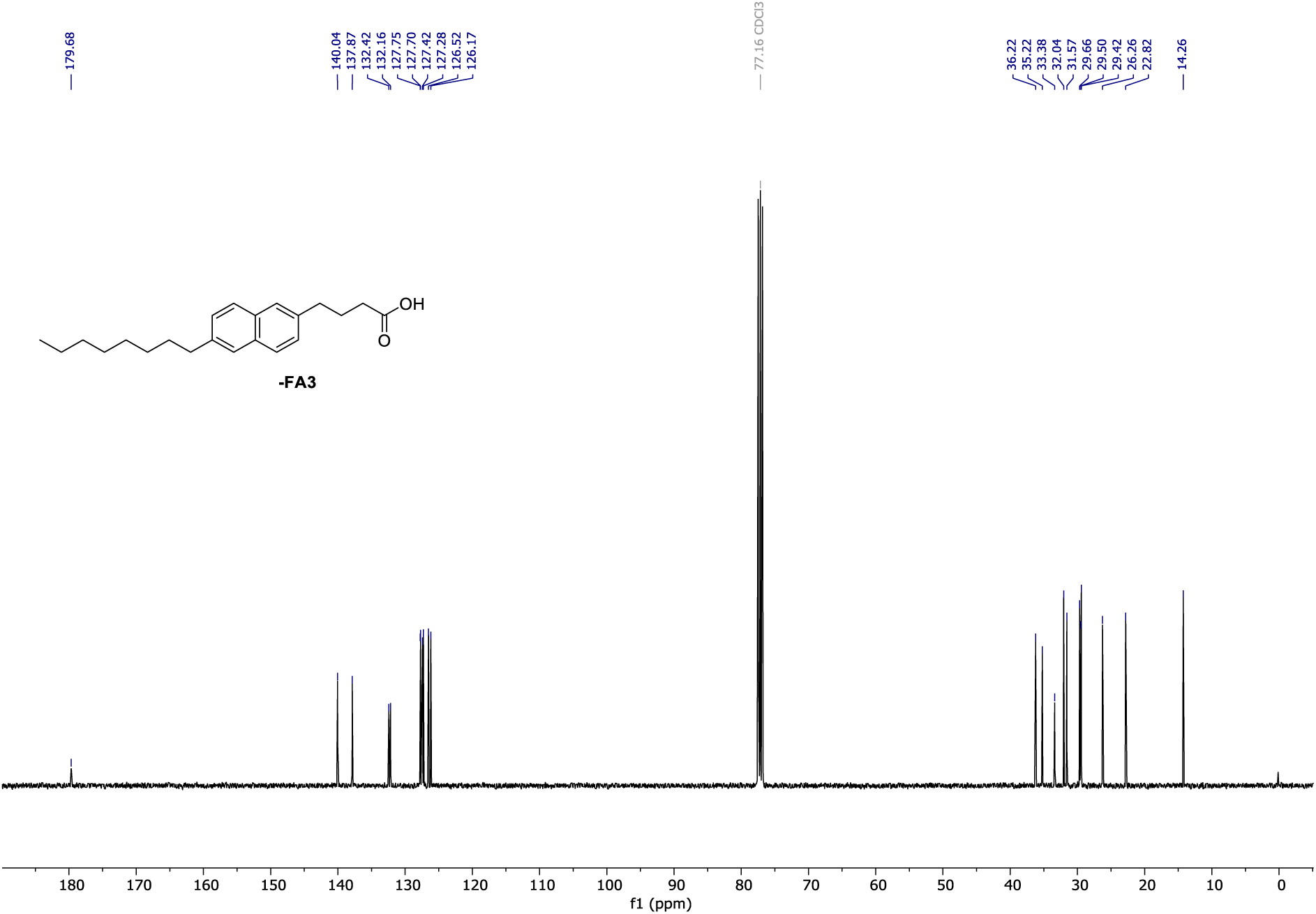

**^13^C NMR** (101 MHz, CDCl_3_) of **π-FA3**.

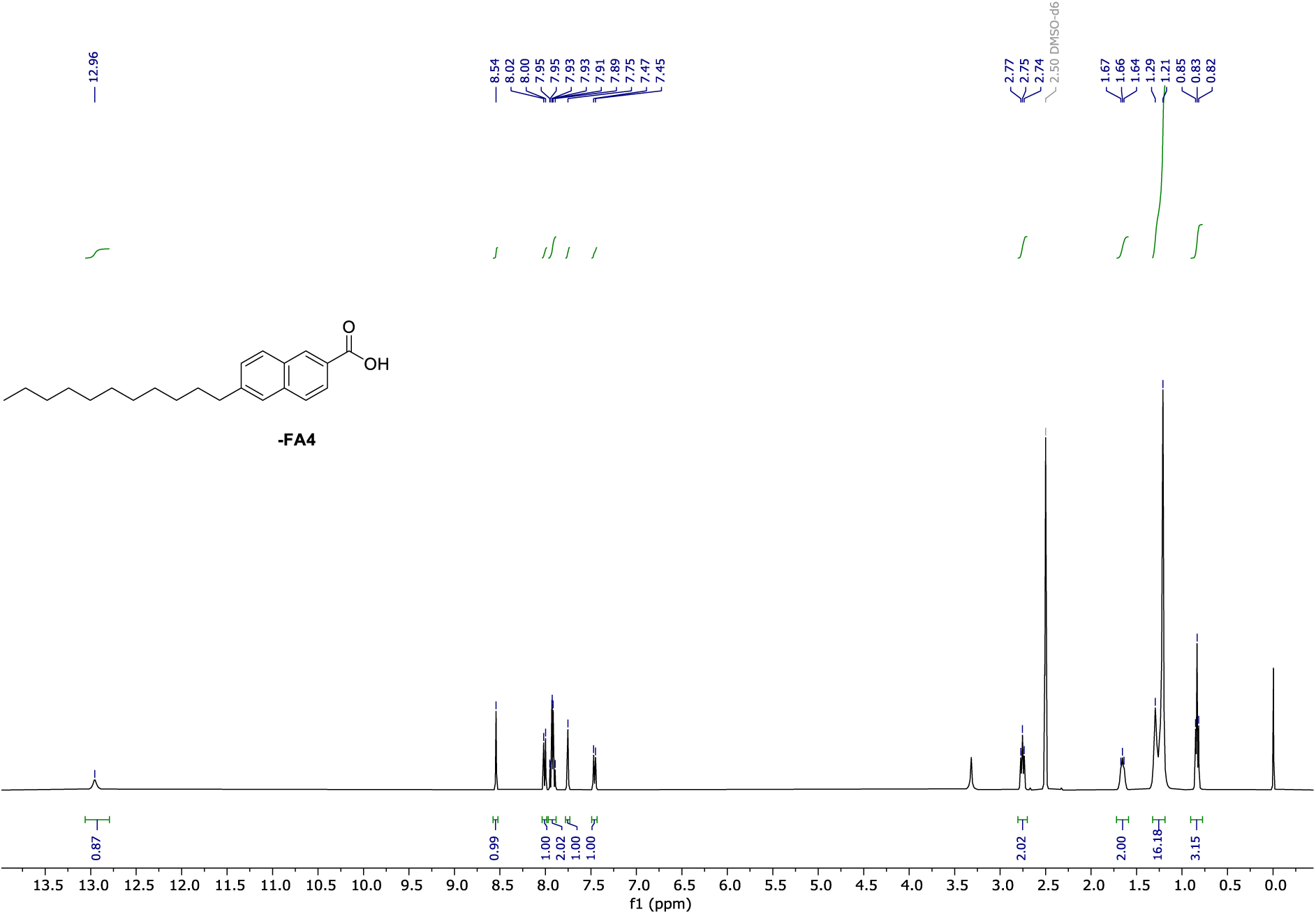

**^1^H NMR** (400 MHz, CDCl_3_) of **π-FA4**.

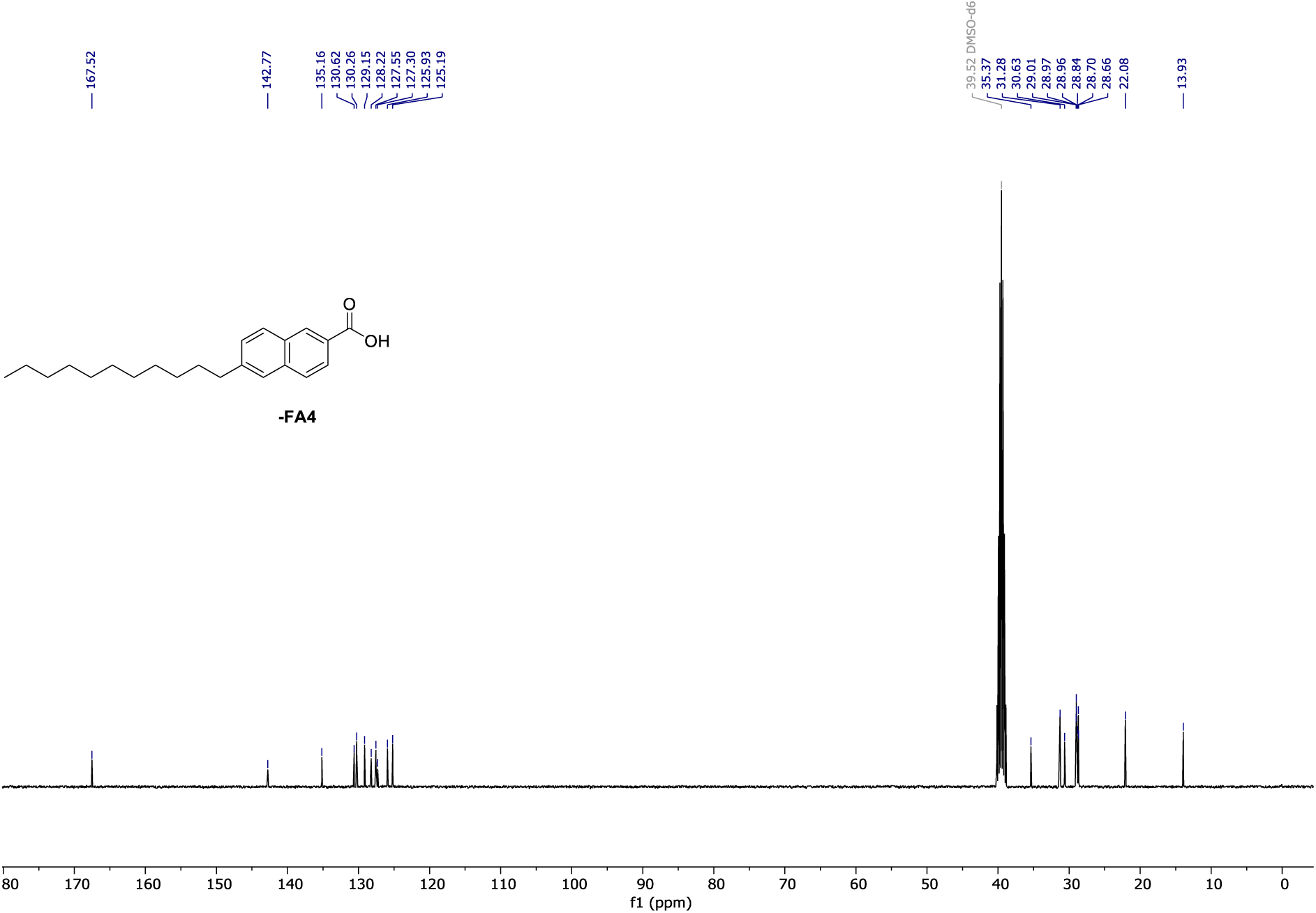

**^13^C NMR** (101 MHz, CDCl_3_) of **π-FA4**.

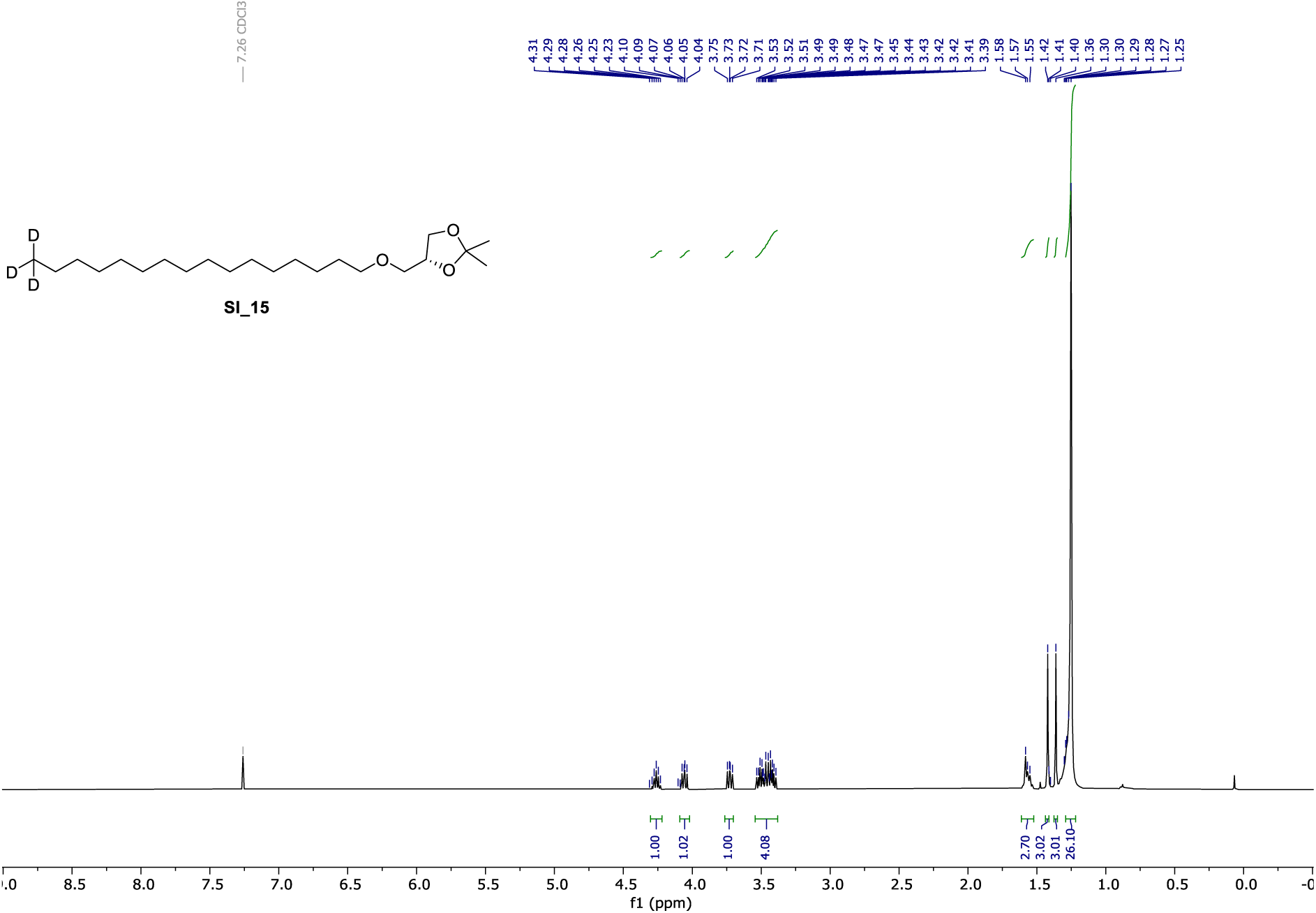

**^1^H NMR** (400 MHz, CDCl_3_) of **SI_15**.

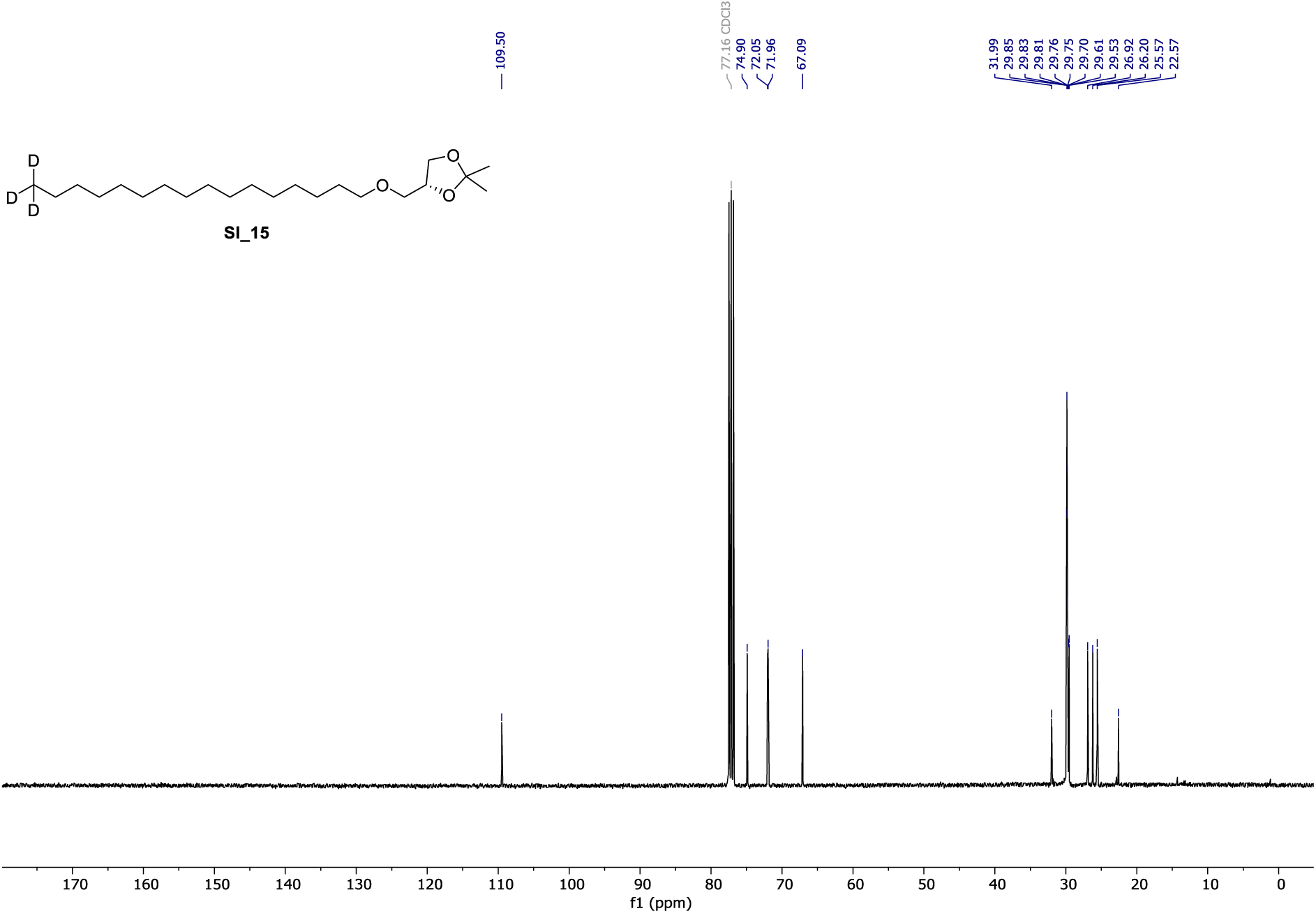

**^13^C NMR** (101 MHz, CDCl_3_) of **SI_15**.

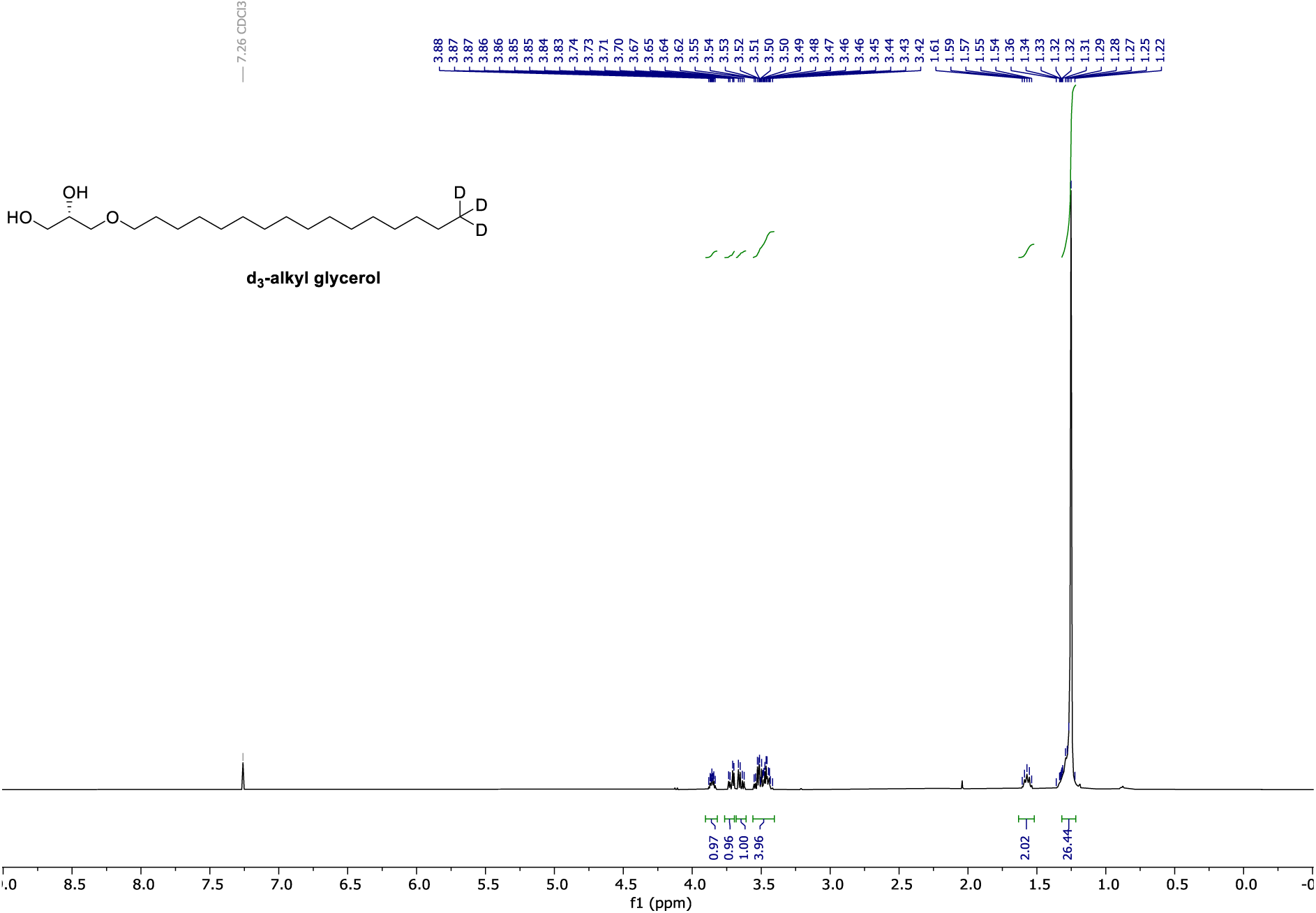

**^1^H NMR** (400 MHz, CDCl_3_) of **d3-alkyl glycerol**.

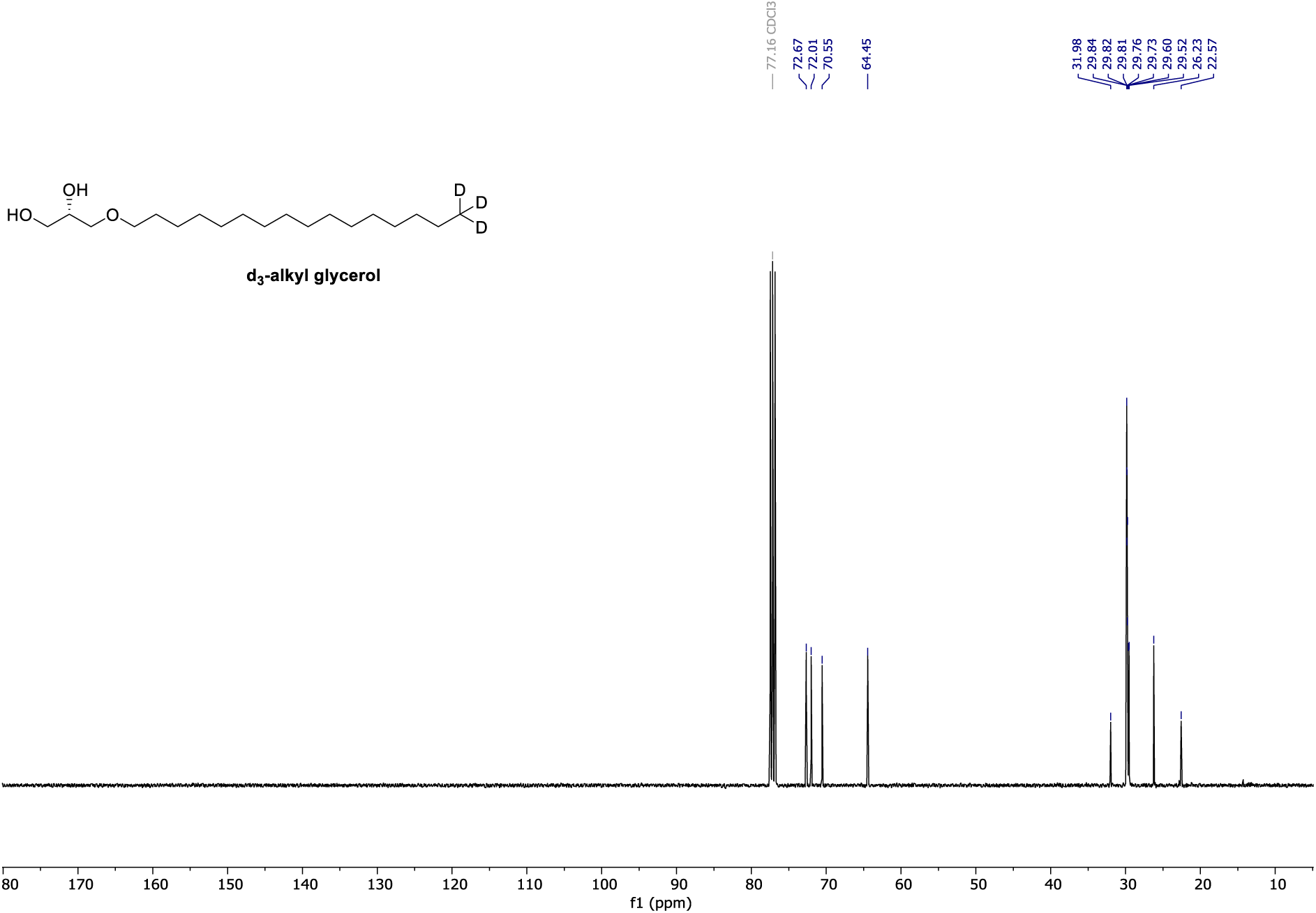

**^13^C NMR** (101 MHz, CDCl_3_) of **d3-alkyl glycerol**.

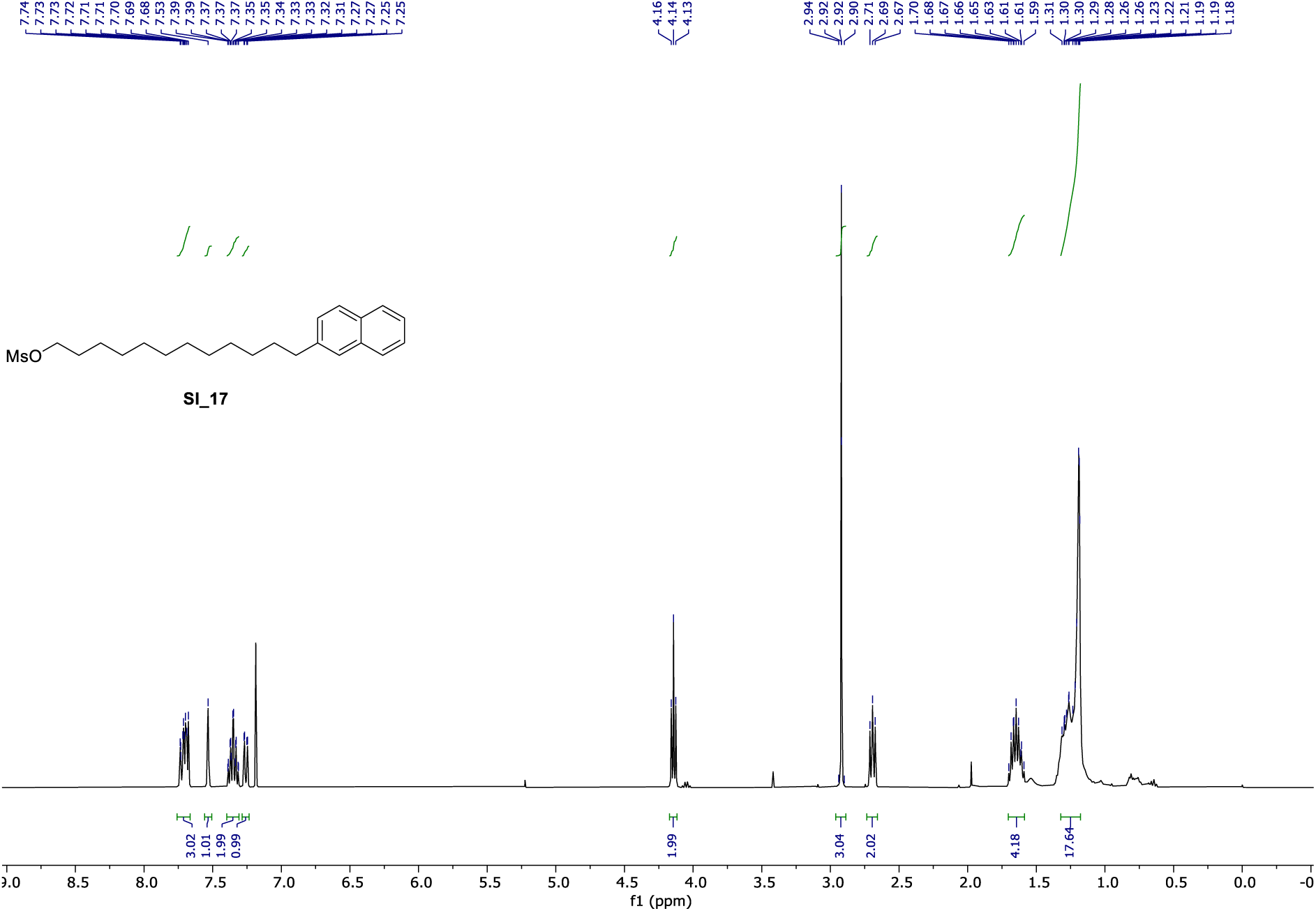

**^1^H NMR** (400 MHz, CDCl_3_) of **SI_17**.

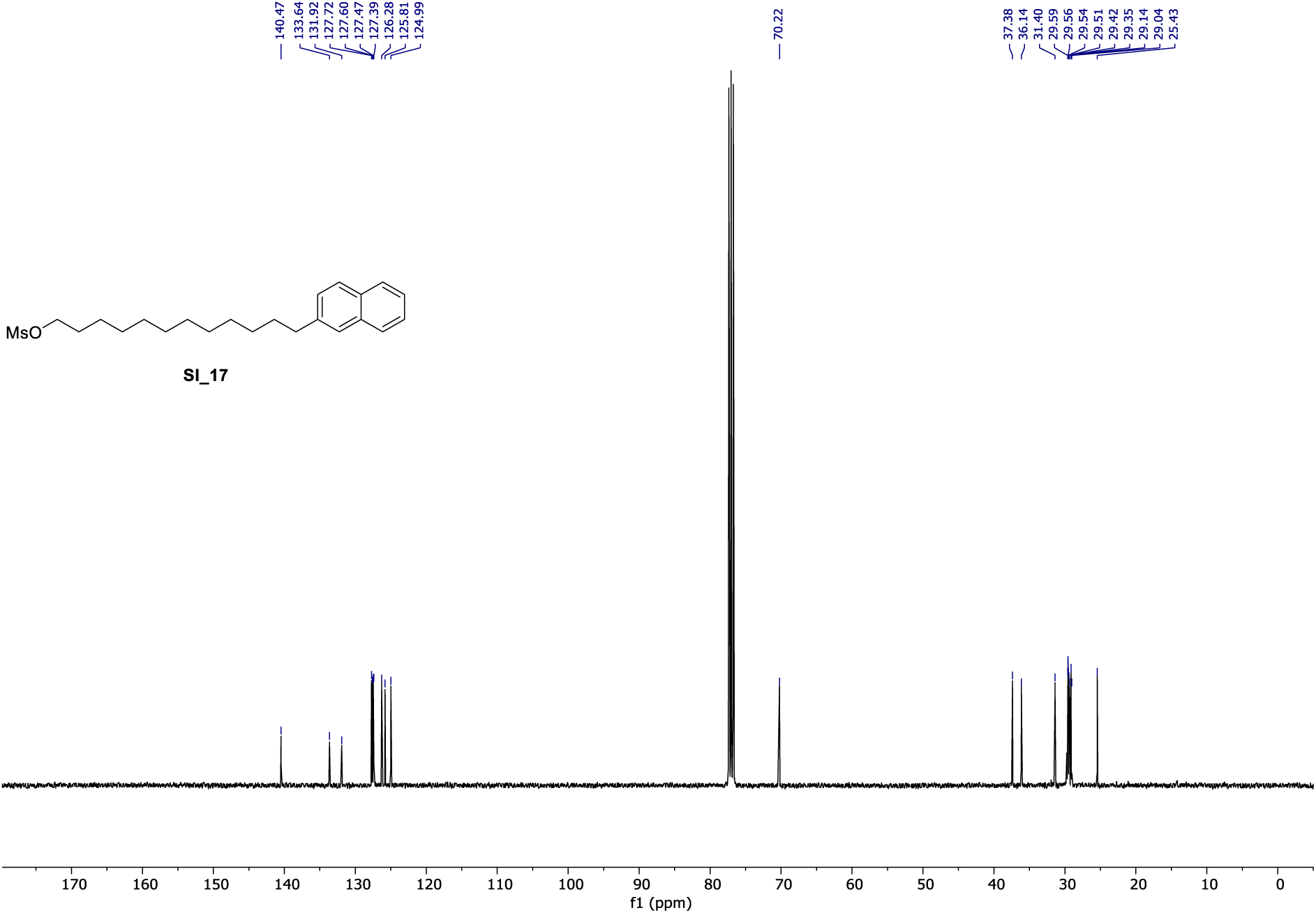

**^13^C NMR** (101 MHz, CDCl_3_) of **SI_17**.

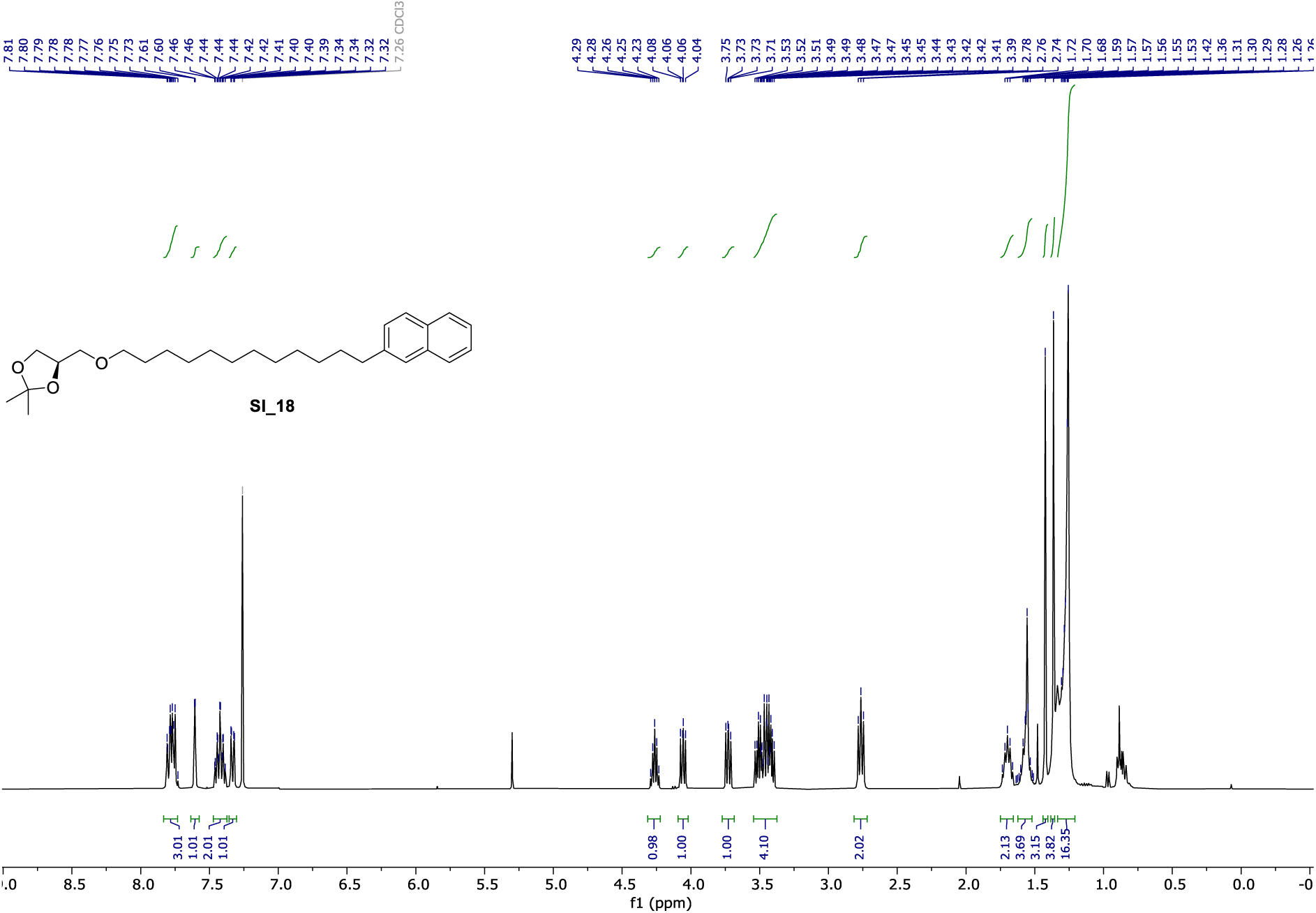

**^1^H NMR** (400 MHz, CDCl_3_) of **SI_18**.

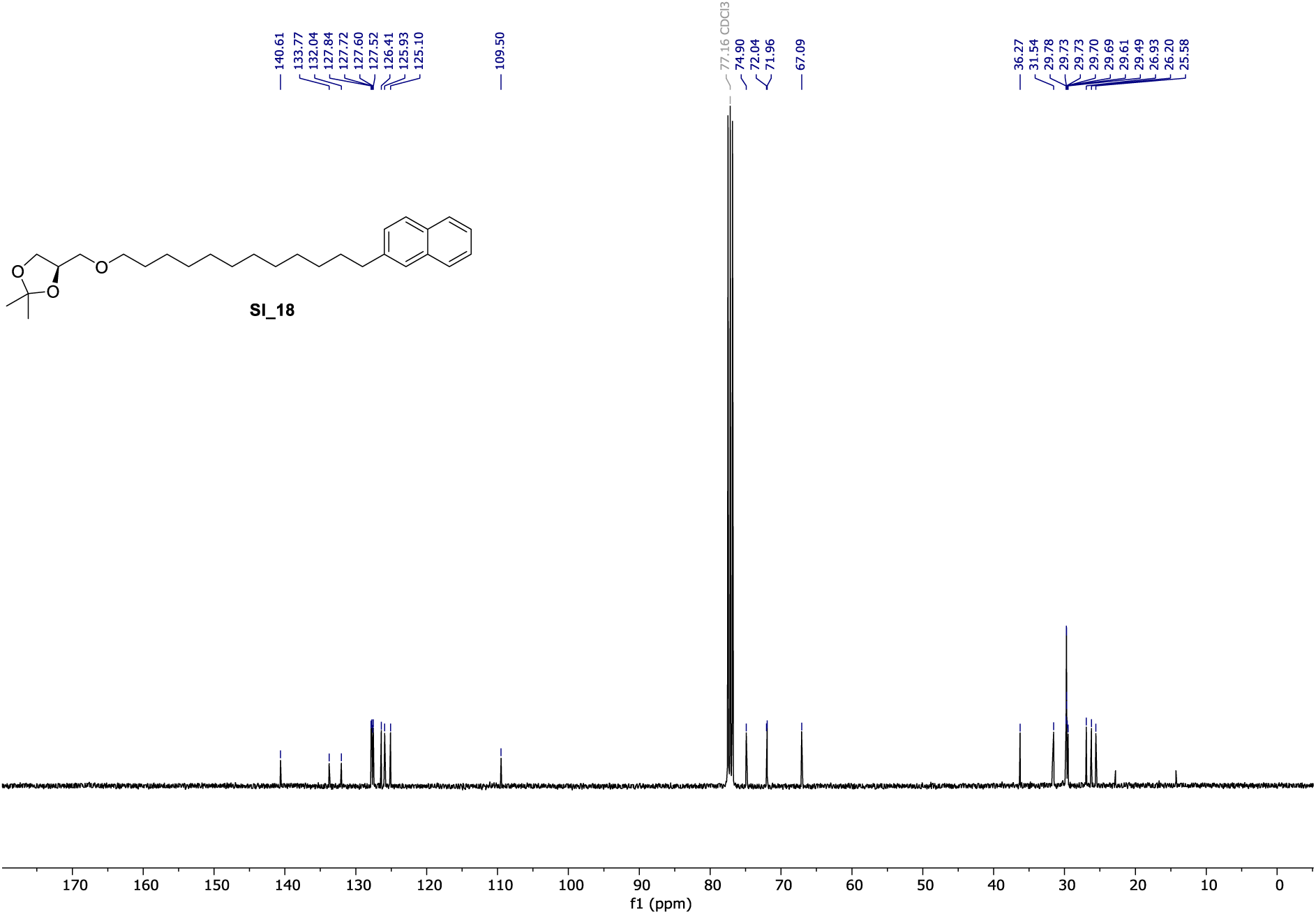

**^13^C NMR** (101 MHz, CDCl_3_) of **SI_18**.

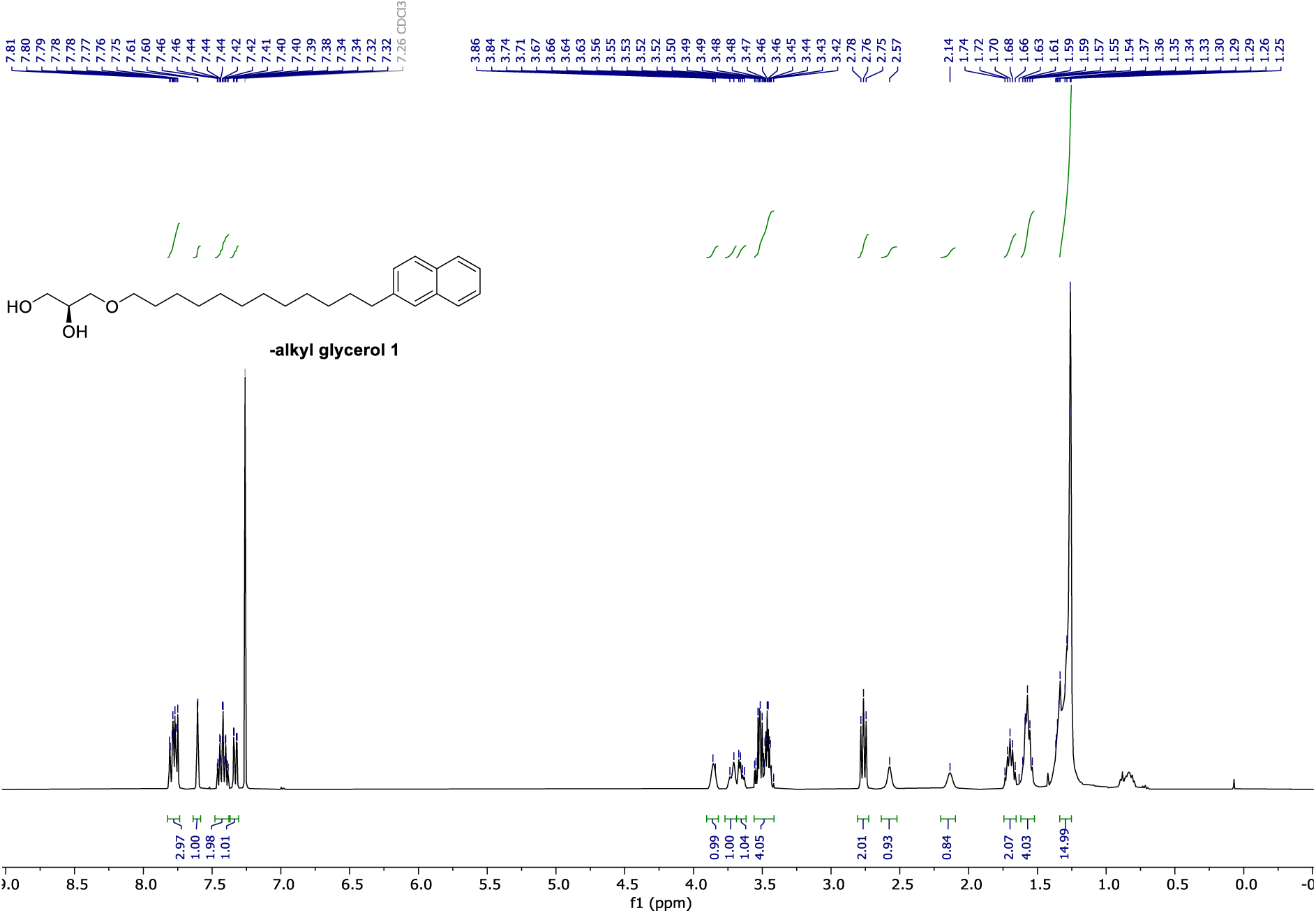

**^13^C NMR** (101 MHz, CDCl_3_) of **π-alkyl glycerol 1**.

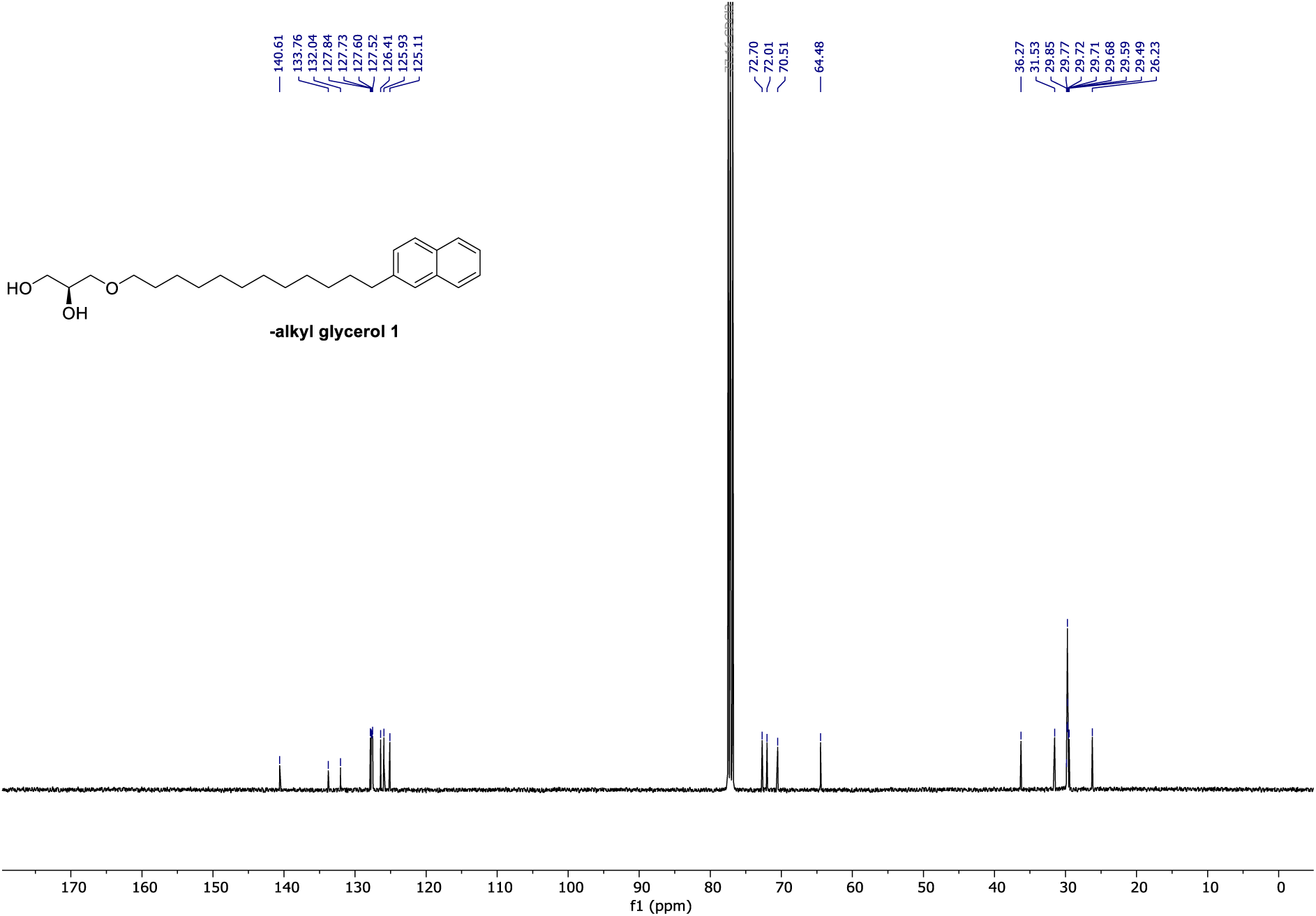

**^1^H NMR** (400 MHz, CDCl_3_) of **SI_20**.

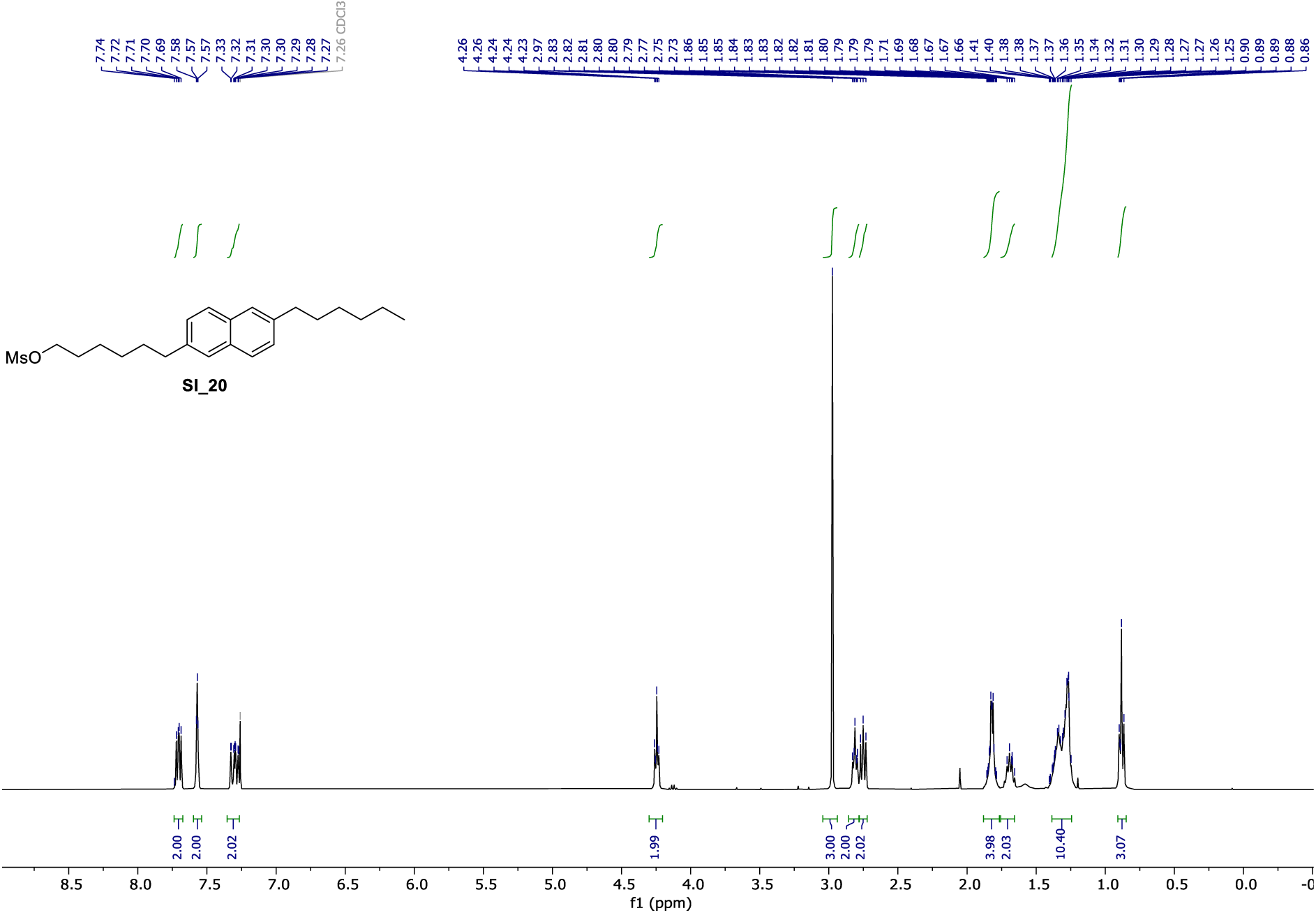

**^13^C NMR** (101 MHz, CDCl_3_) of **SI_20**.

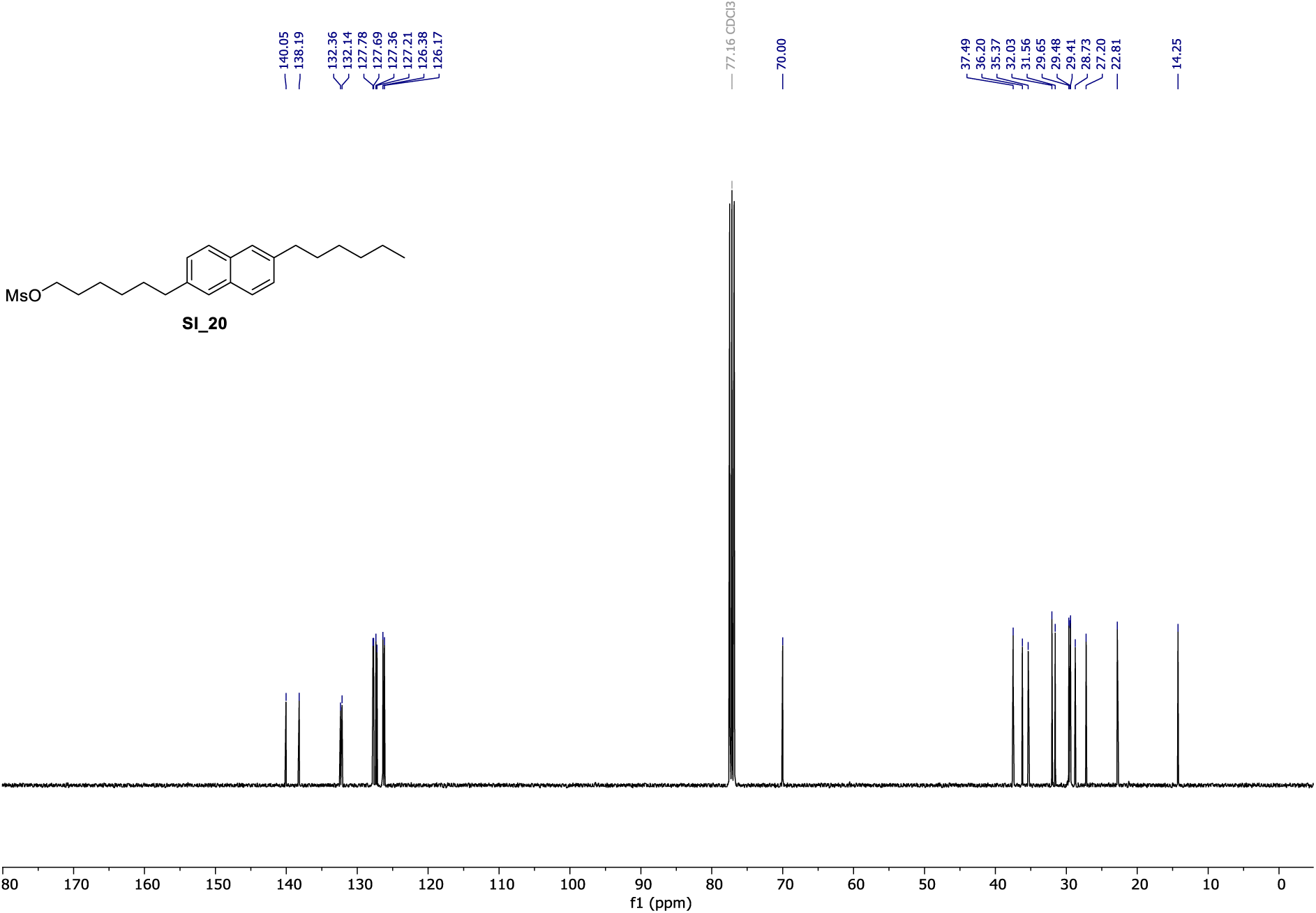

**^1^H NMR** (400 MHz, CDCl_3_) of **SI_21**.

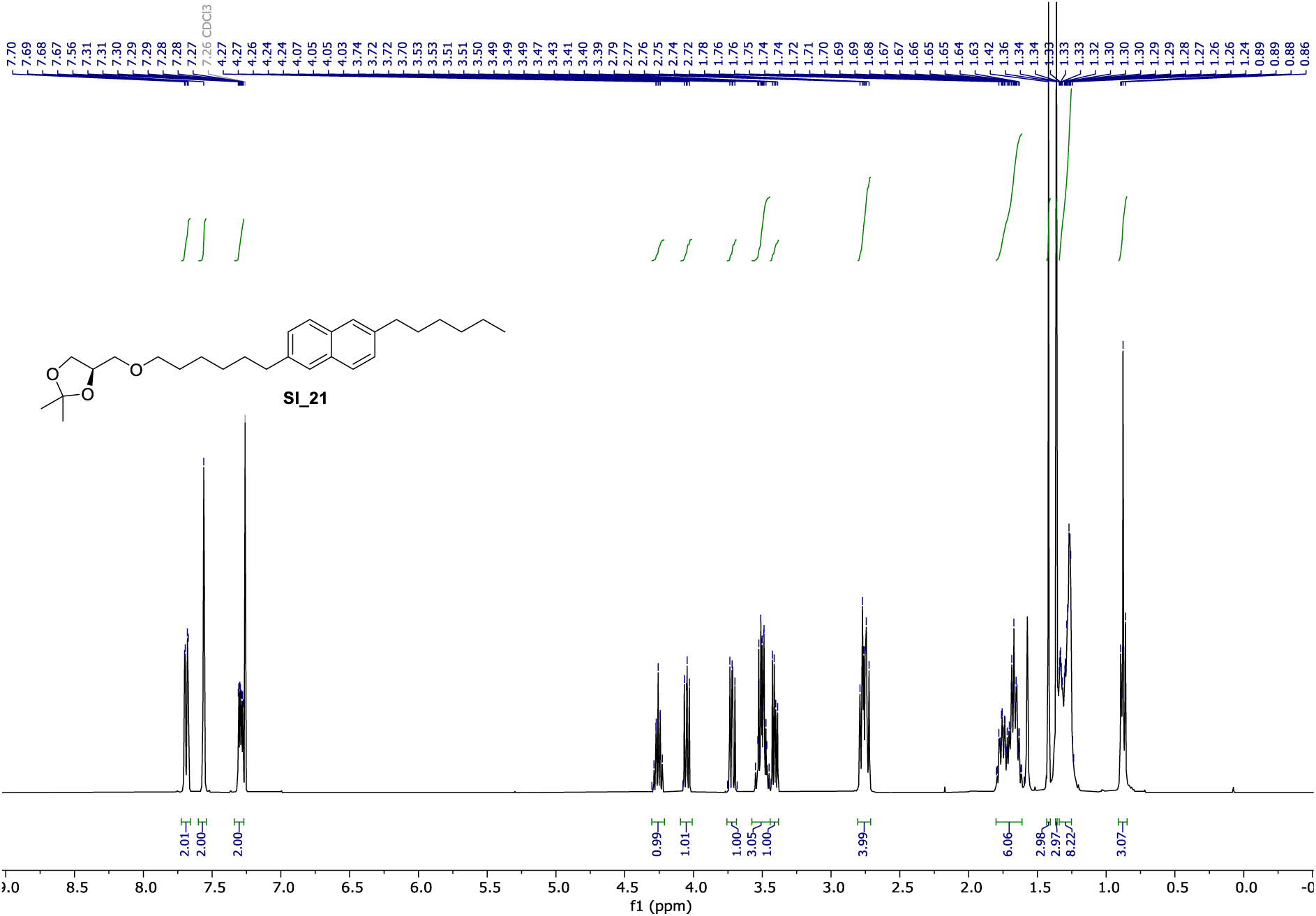

**^13^C NMR** (101 MHz, CDCl_3_) of **SI_21**.

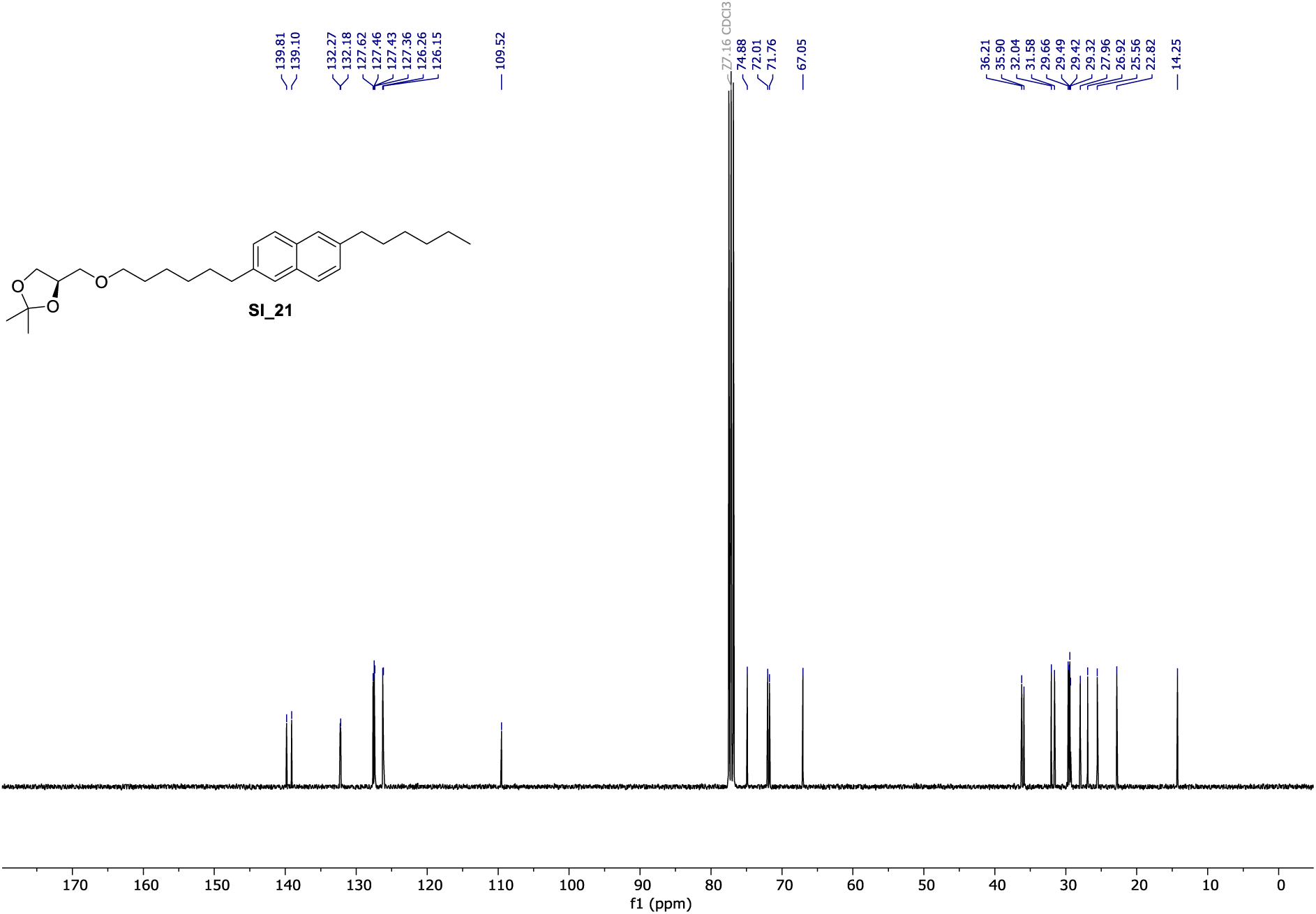

**^1^H NMR** (400 MHz, CDCl_3_) of **π-alkyl glycerol 3**.

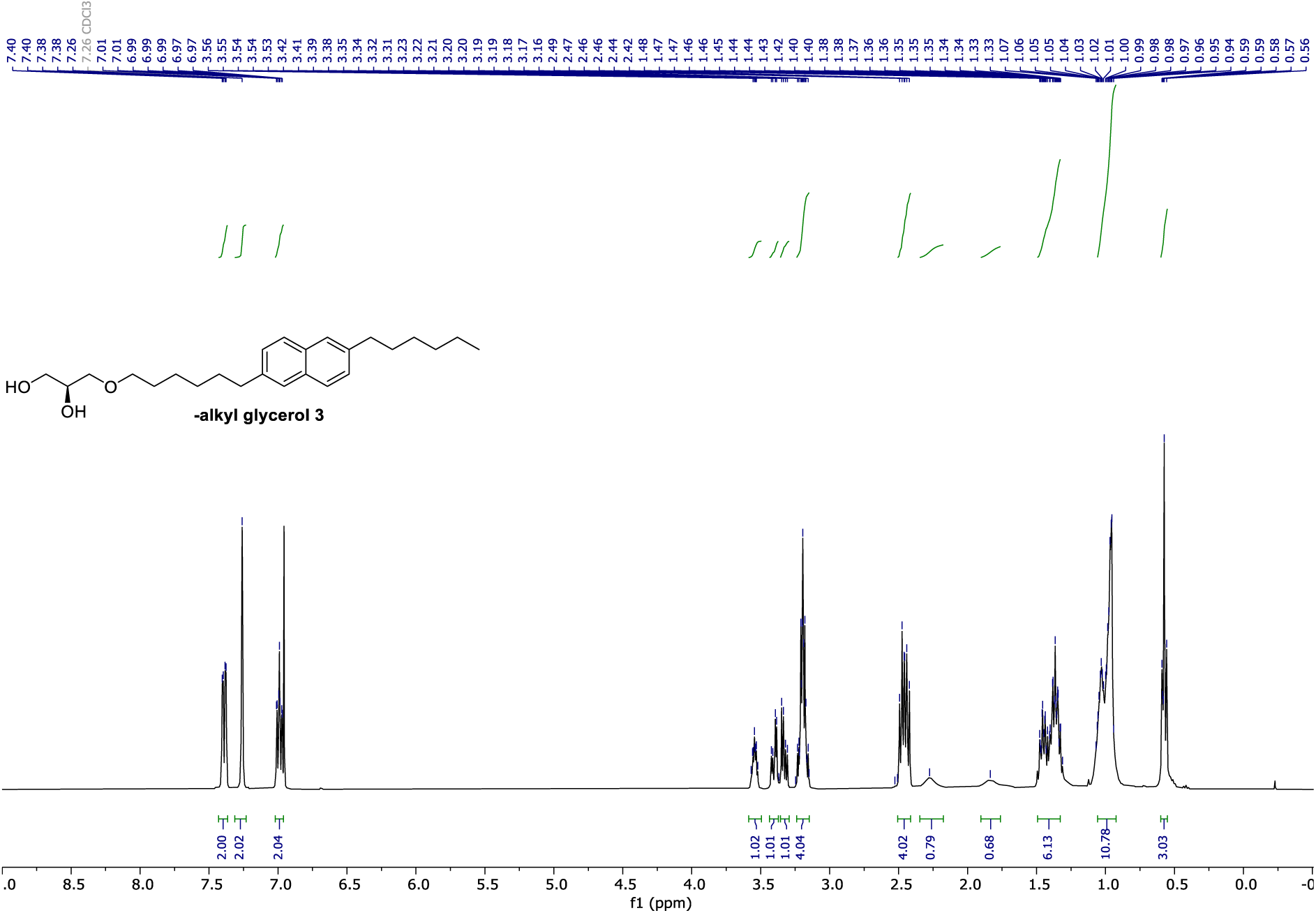

**^13^C NMR** (101 MHz, CDCl_3_) of **π-alkyl glycerol 3**.

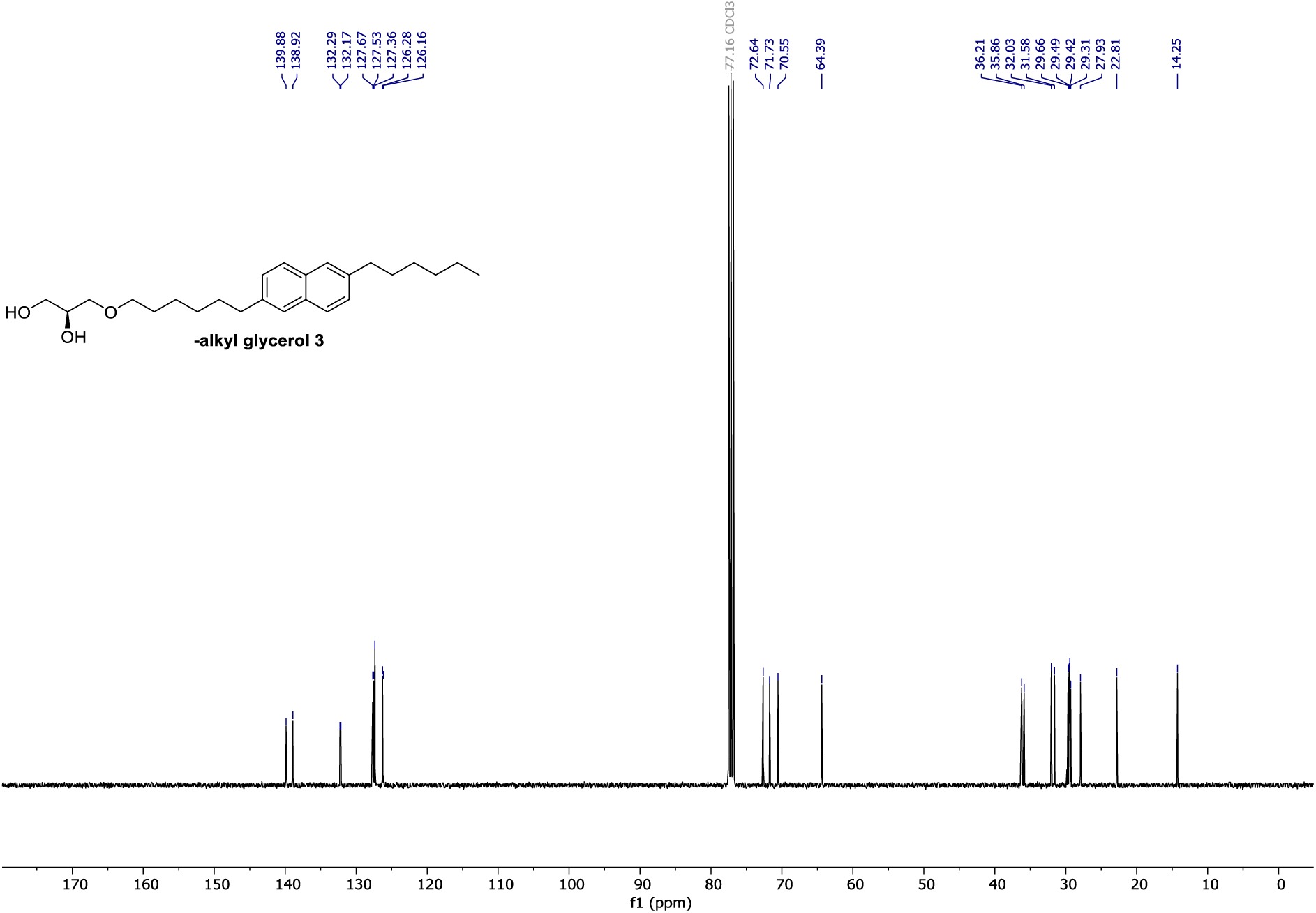

**^1^H NMR** (400 MHz, CDCl_3_) of **SI_22**.

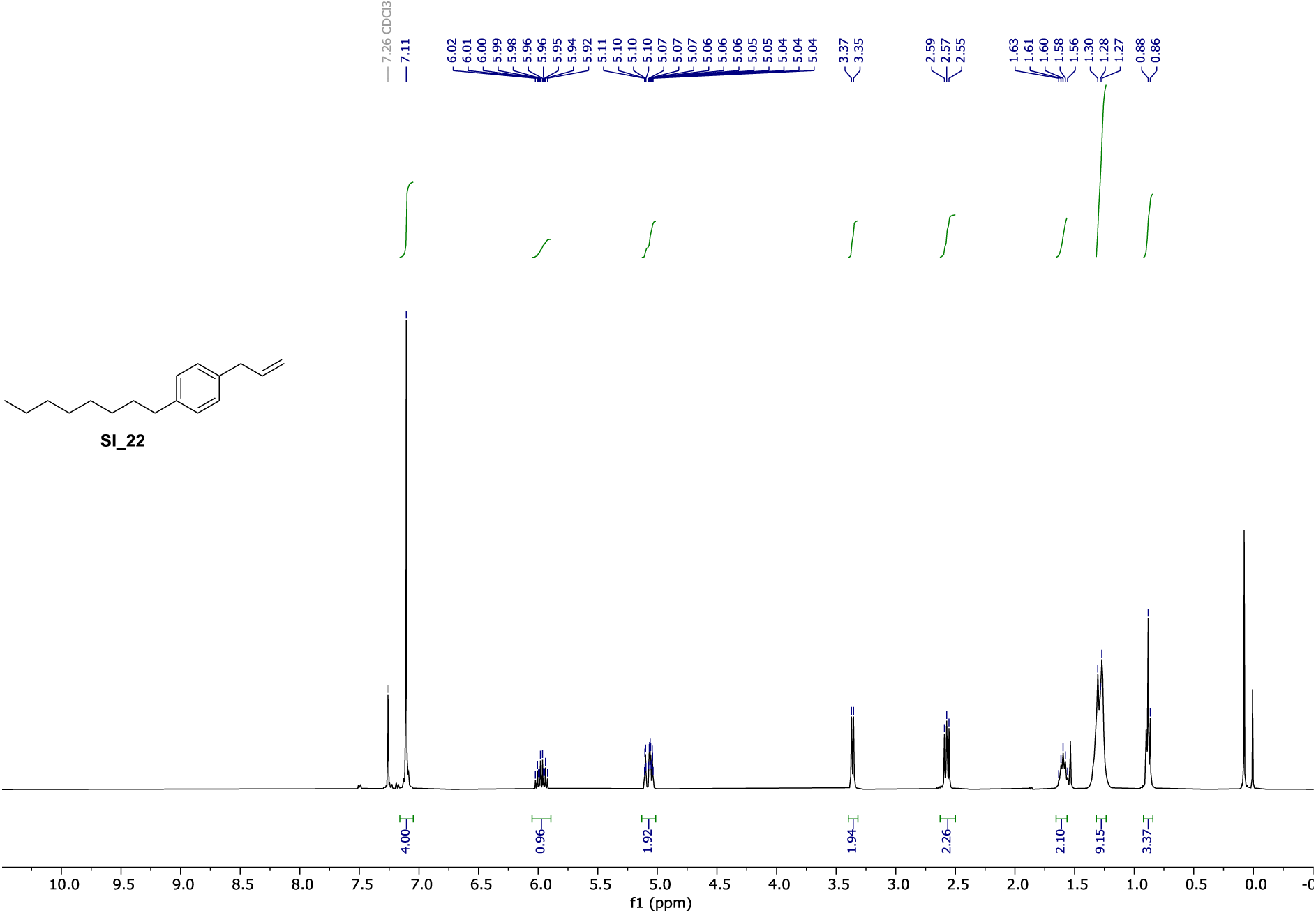

**^1^H NMR** (400 MHz, MeOH-d_4_) of **π-Sph-1**.

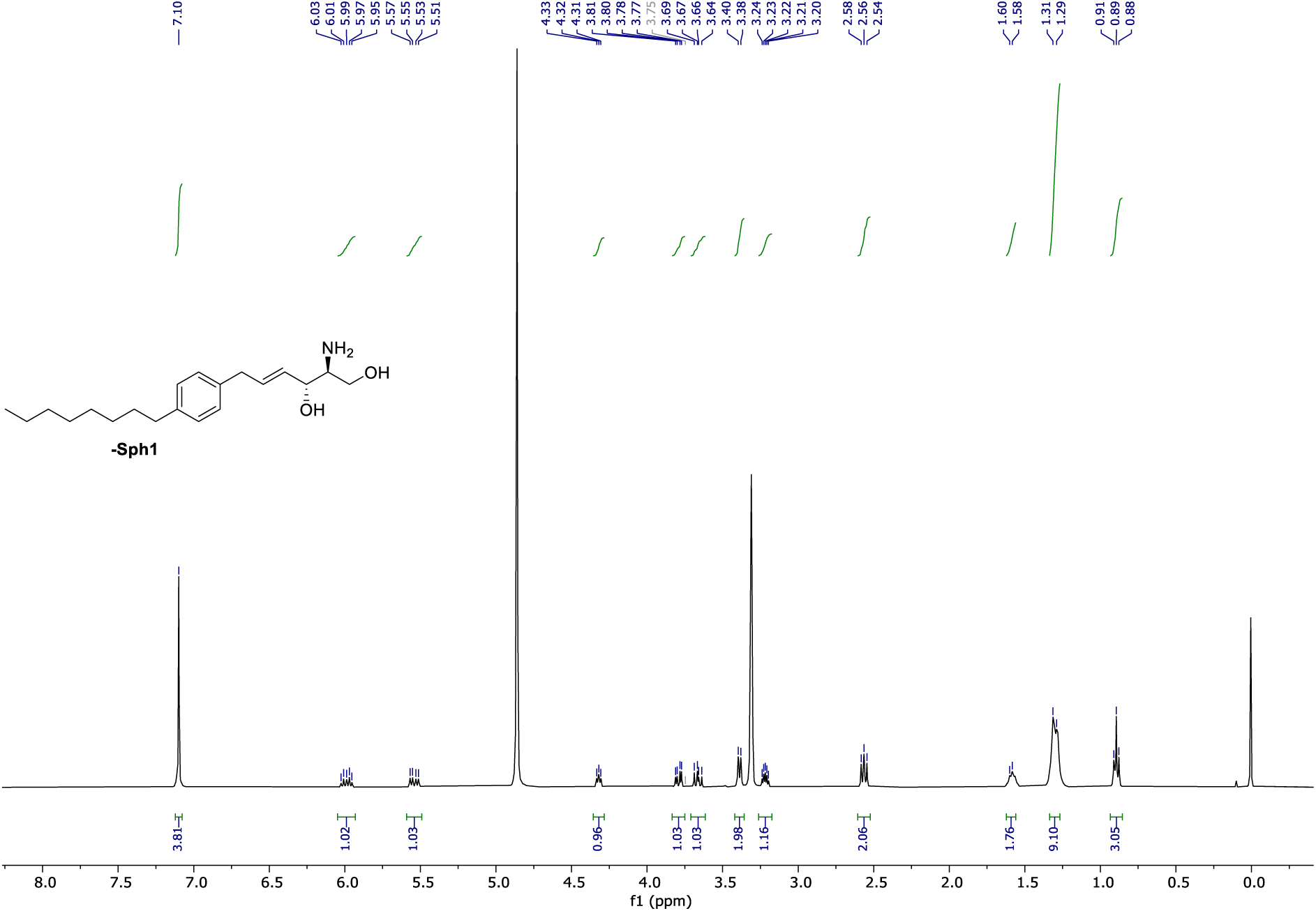

**^1^H NMR** (400 MHz, CDCl_3_) of **SI_25**.

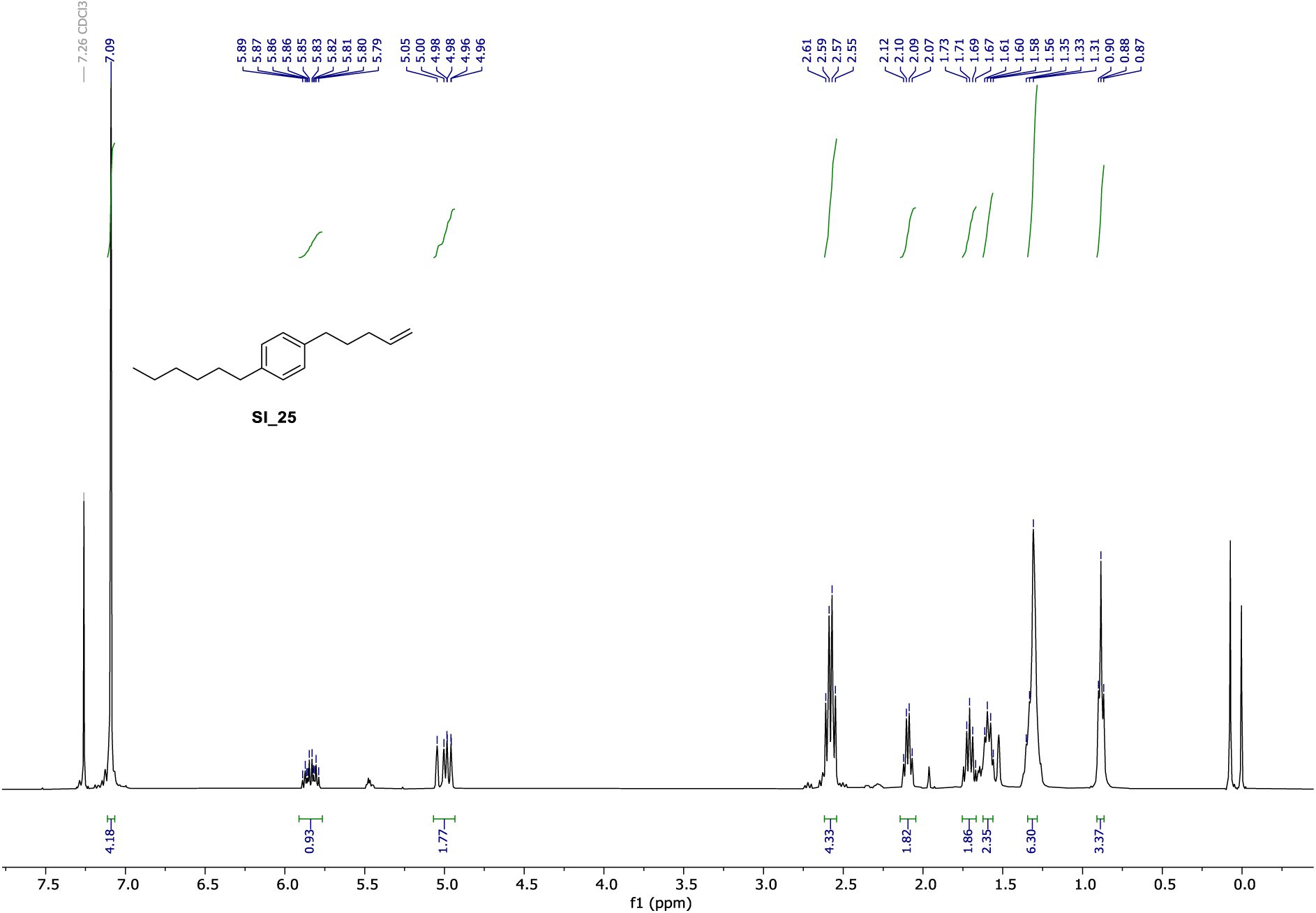

**^1^H NMR** (400 MHz, DMSO-d_6_) of **π-Sph2**.

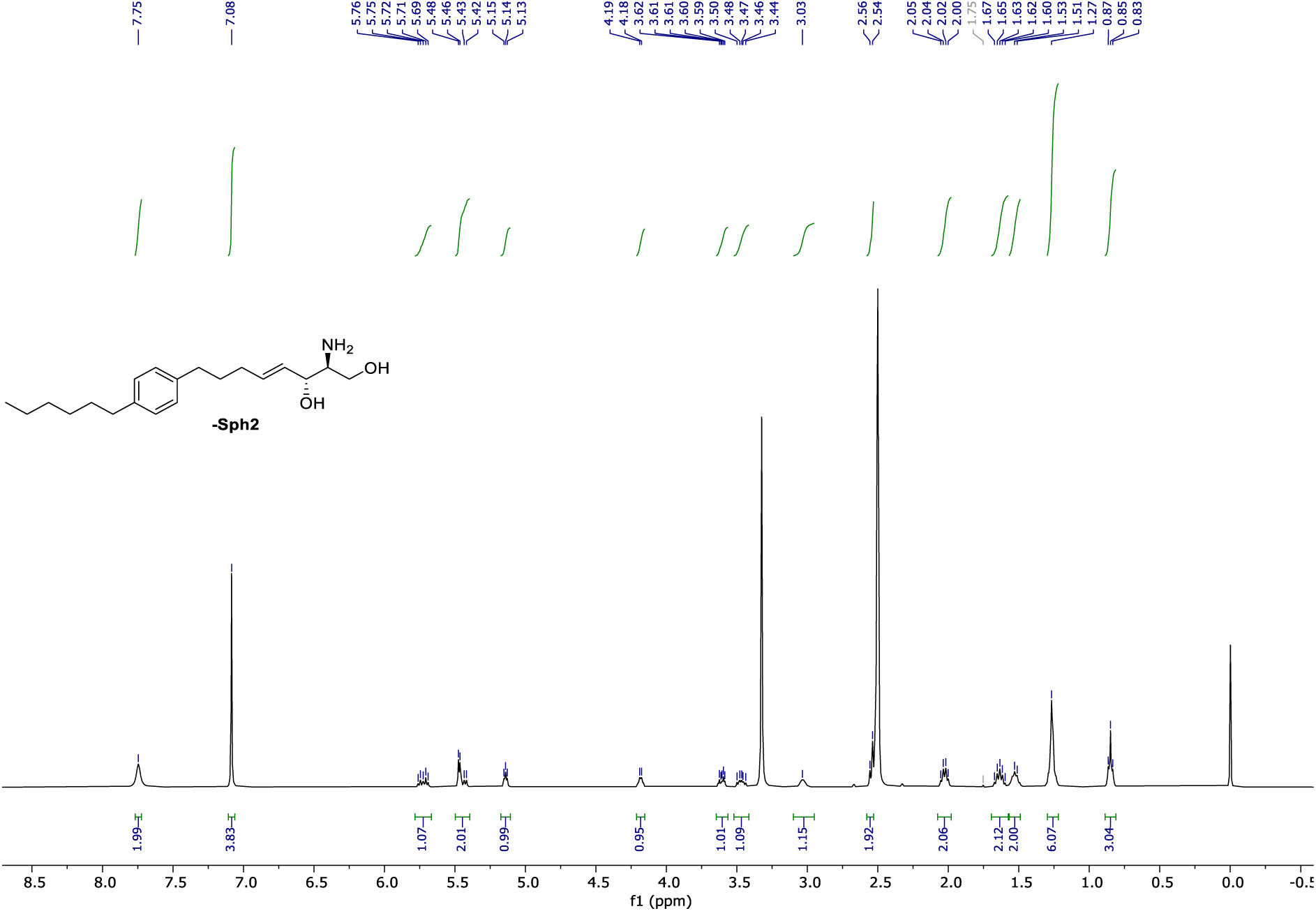

**^1^H NMR** (400 MHz, CDCl_3_) of **SI_27**.

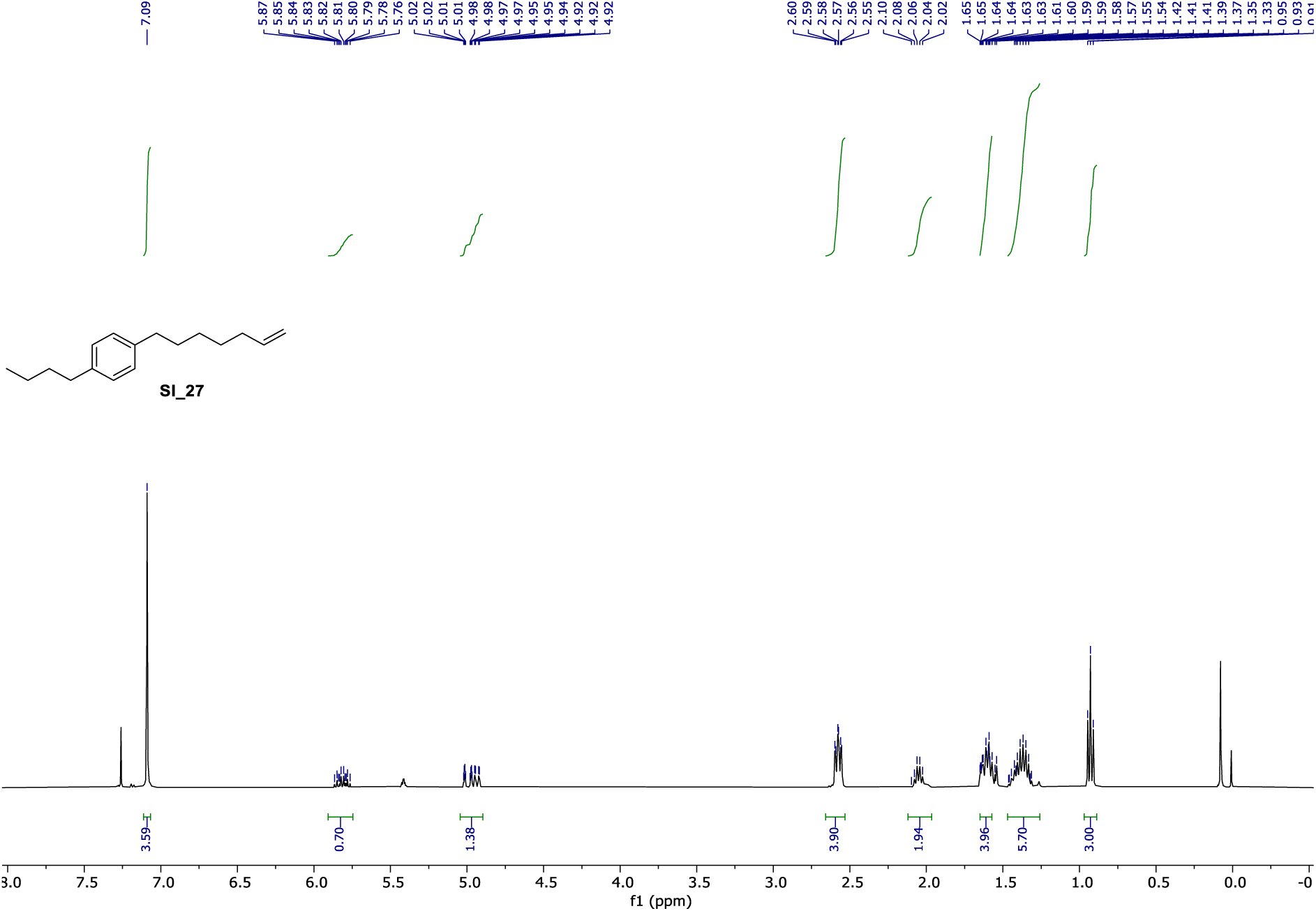

**^1^H NMR** (400 MHz, DMSO-d_6_) of **π-Sph3**.

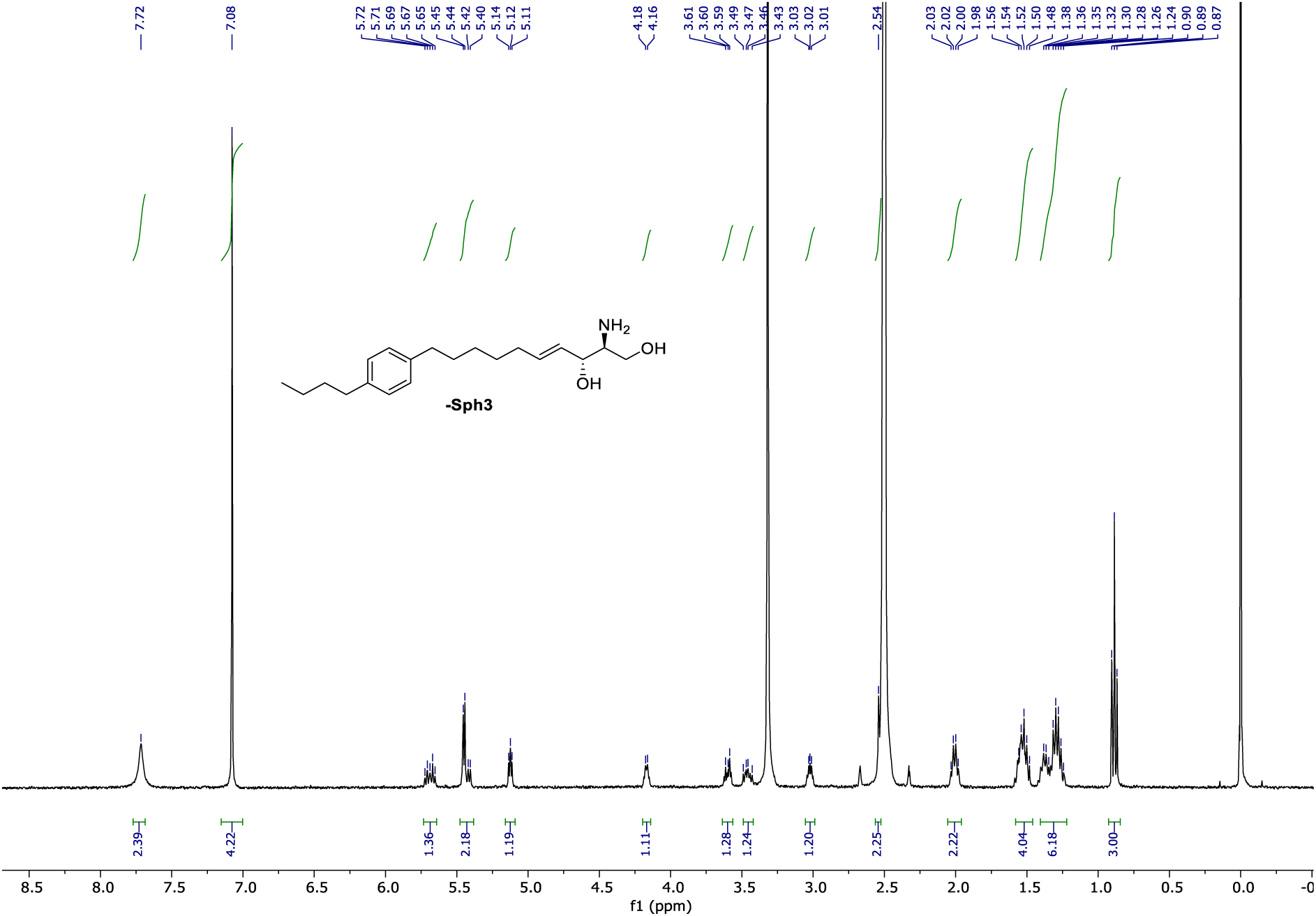

**^1^H NMR** (400 MHz, CDCl_3_) of **SI_29**.

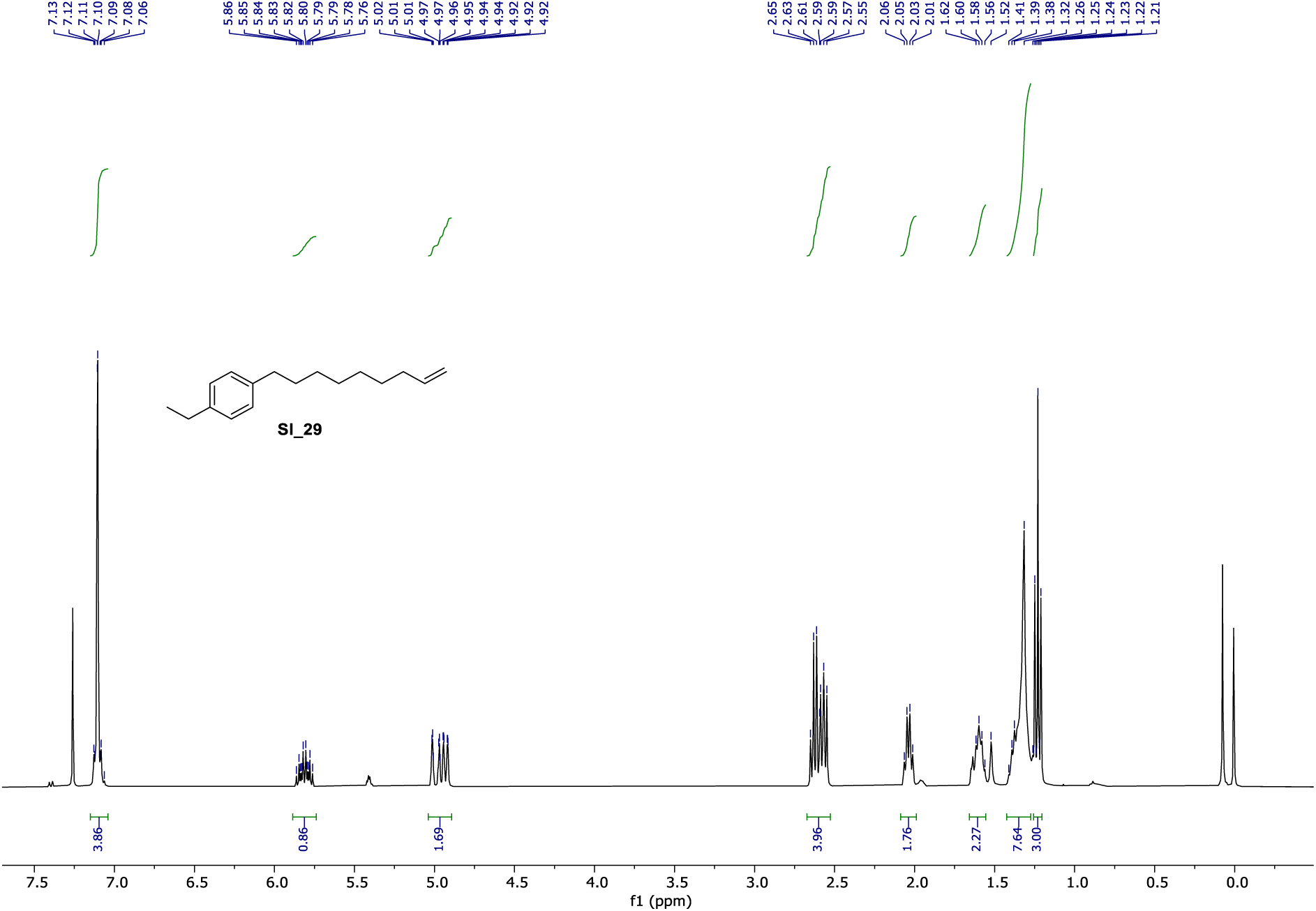

**^1^H NMR** (400 MHz, MeOH-d_4_) of **π-Sph4**.

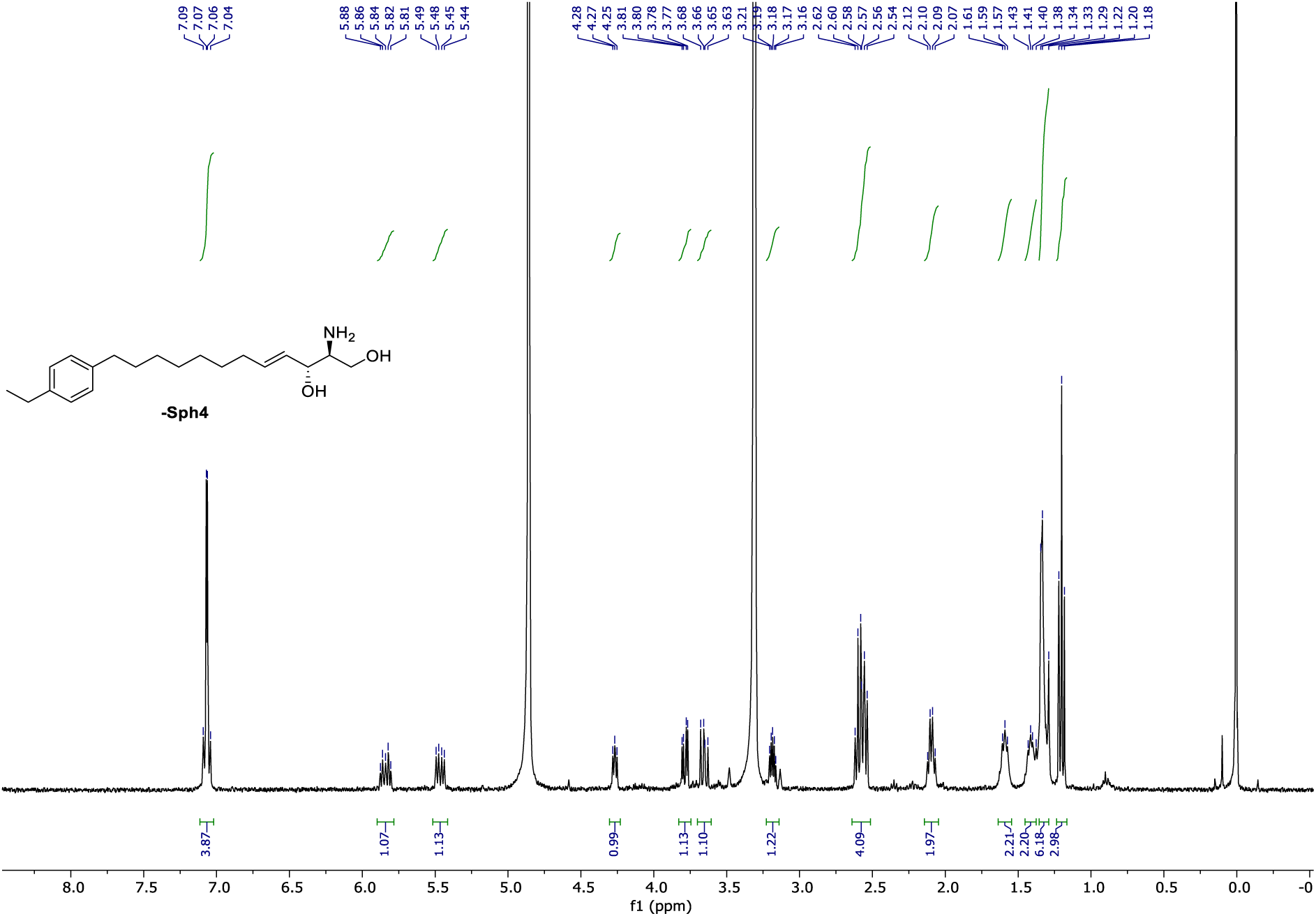

**^1^H NMR** (400 MHz, CDCl_3_) of **SI_31.**

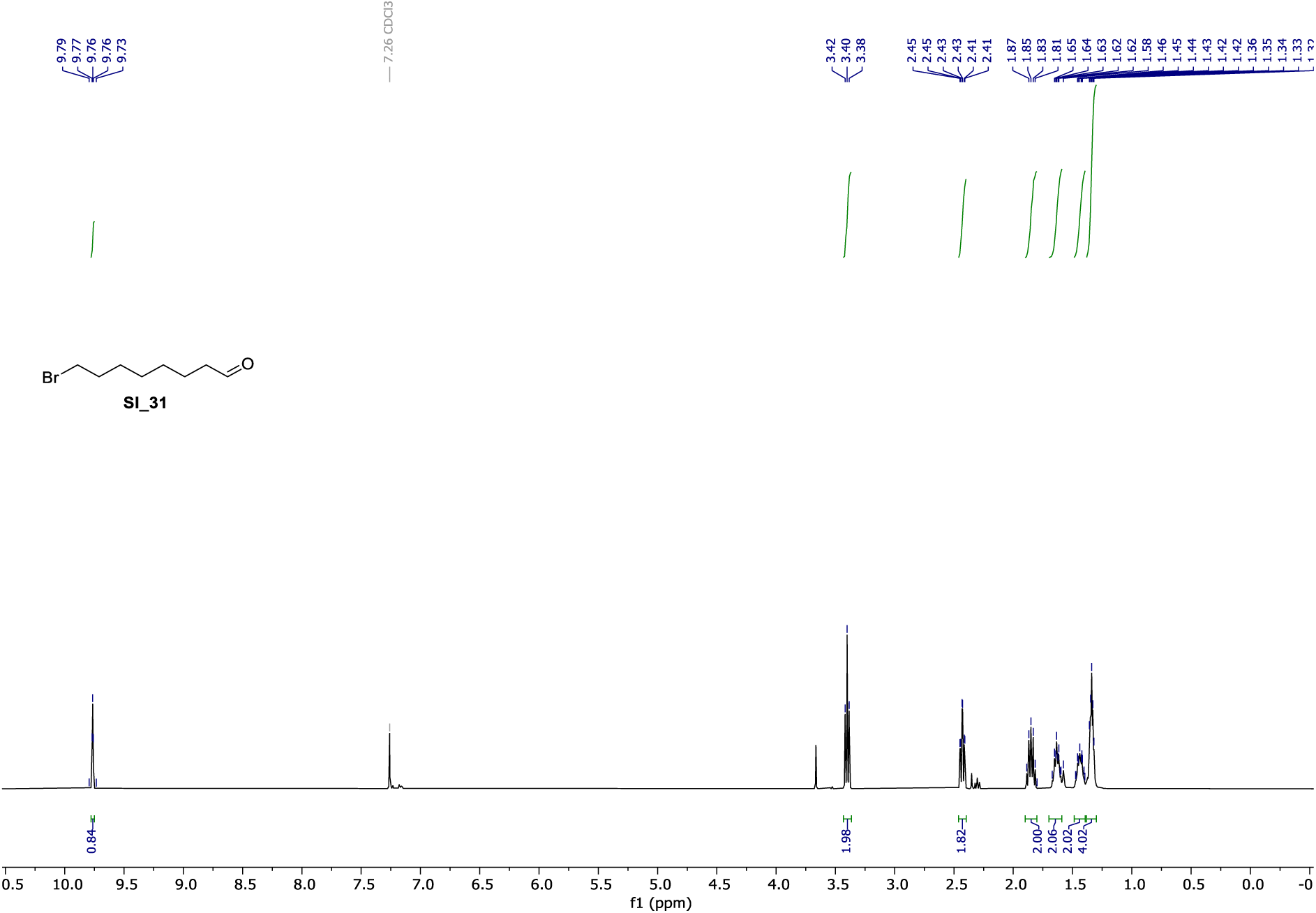

**^13^C NMR** (101 MHz, CDCl_3_) of **SI_31**.

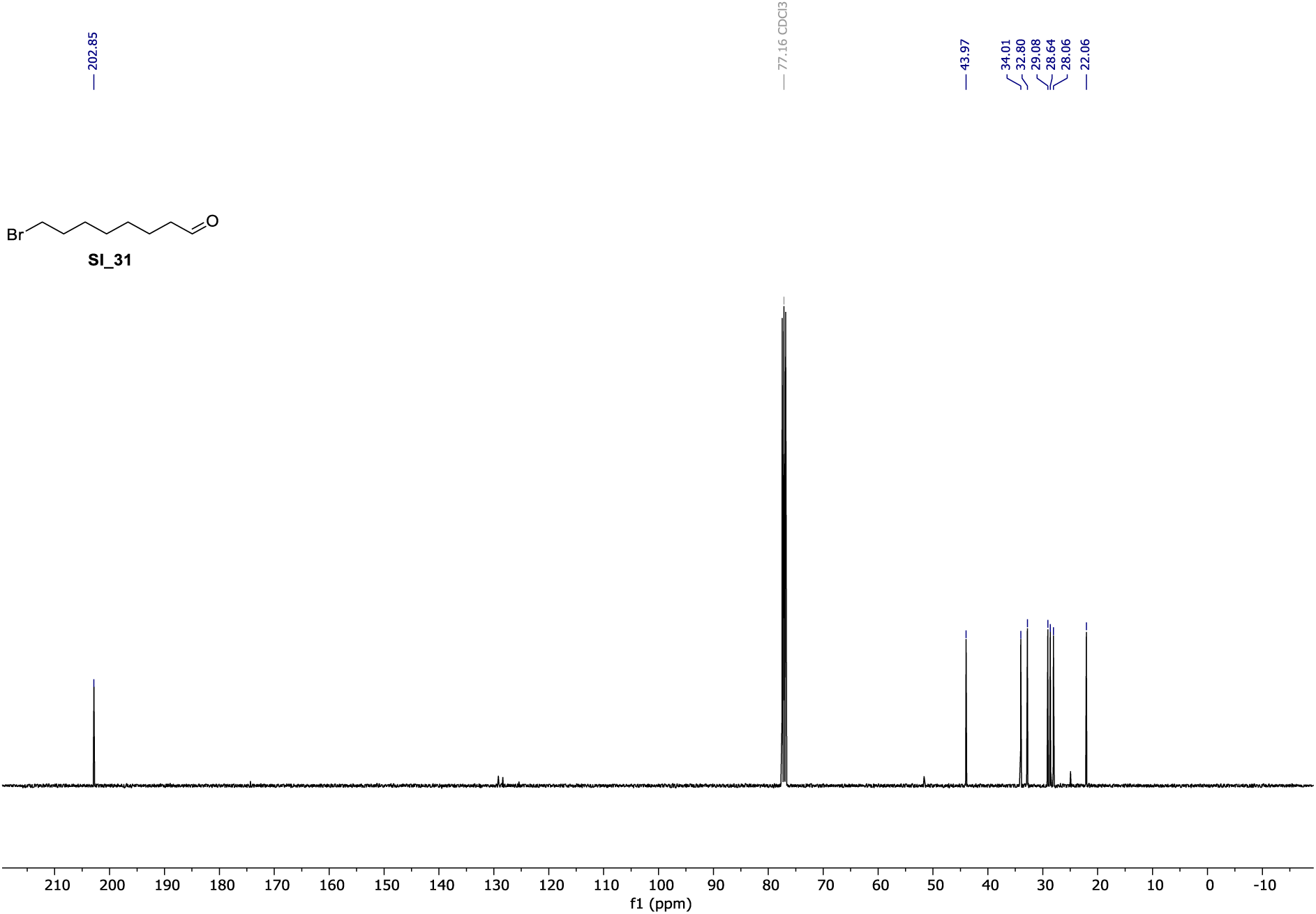

**^1^H NMR** (400 MHz, CDCl_3_) of **SI_33**.

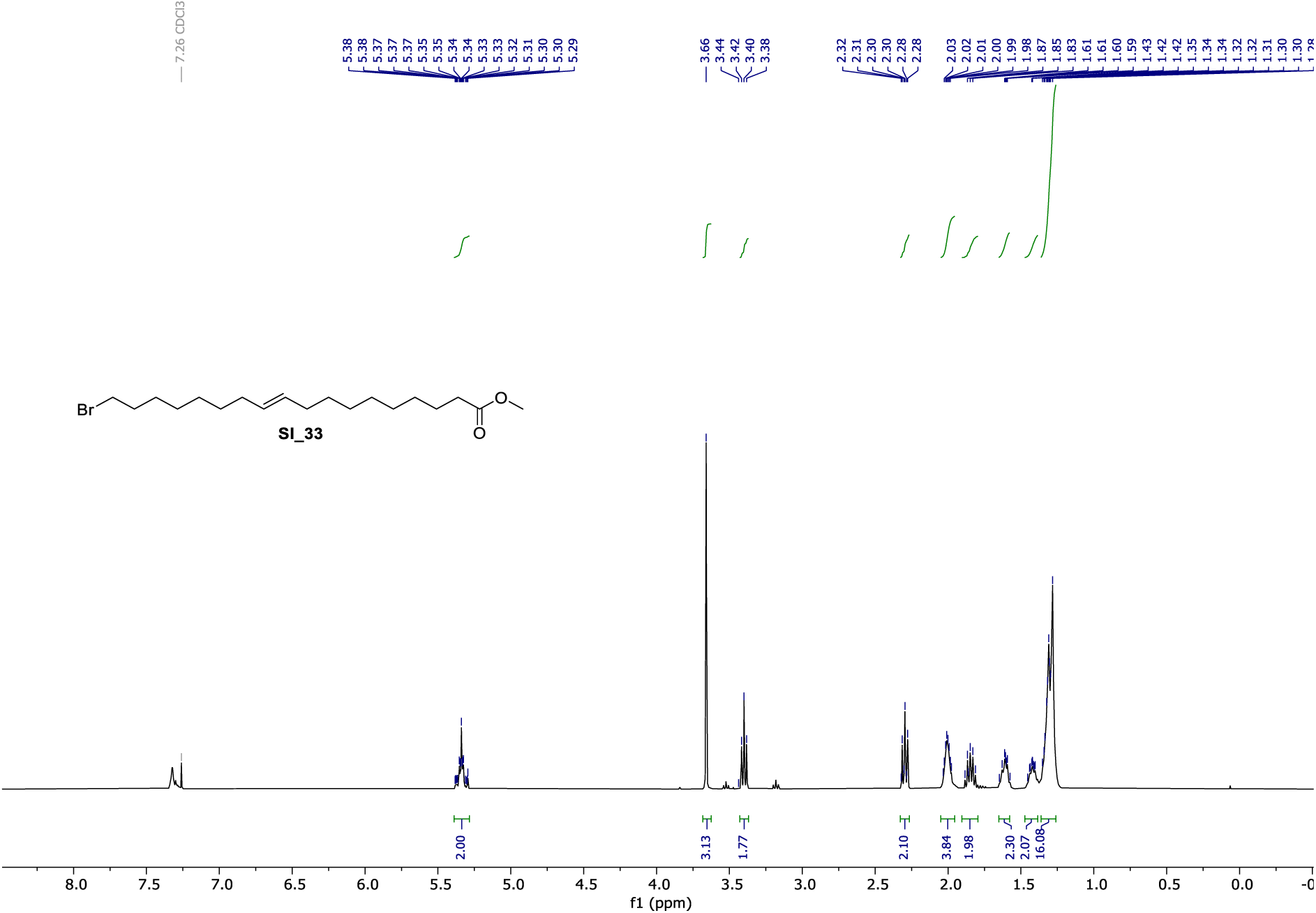

**^13^C NMR** (101 MHz, CDCl_3_) of **SI_33**.

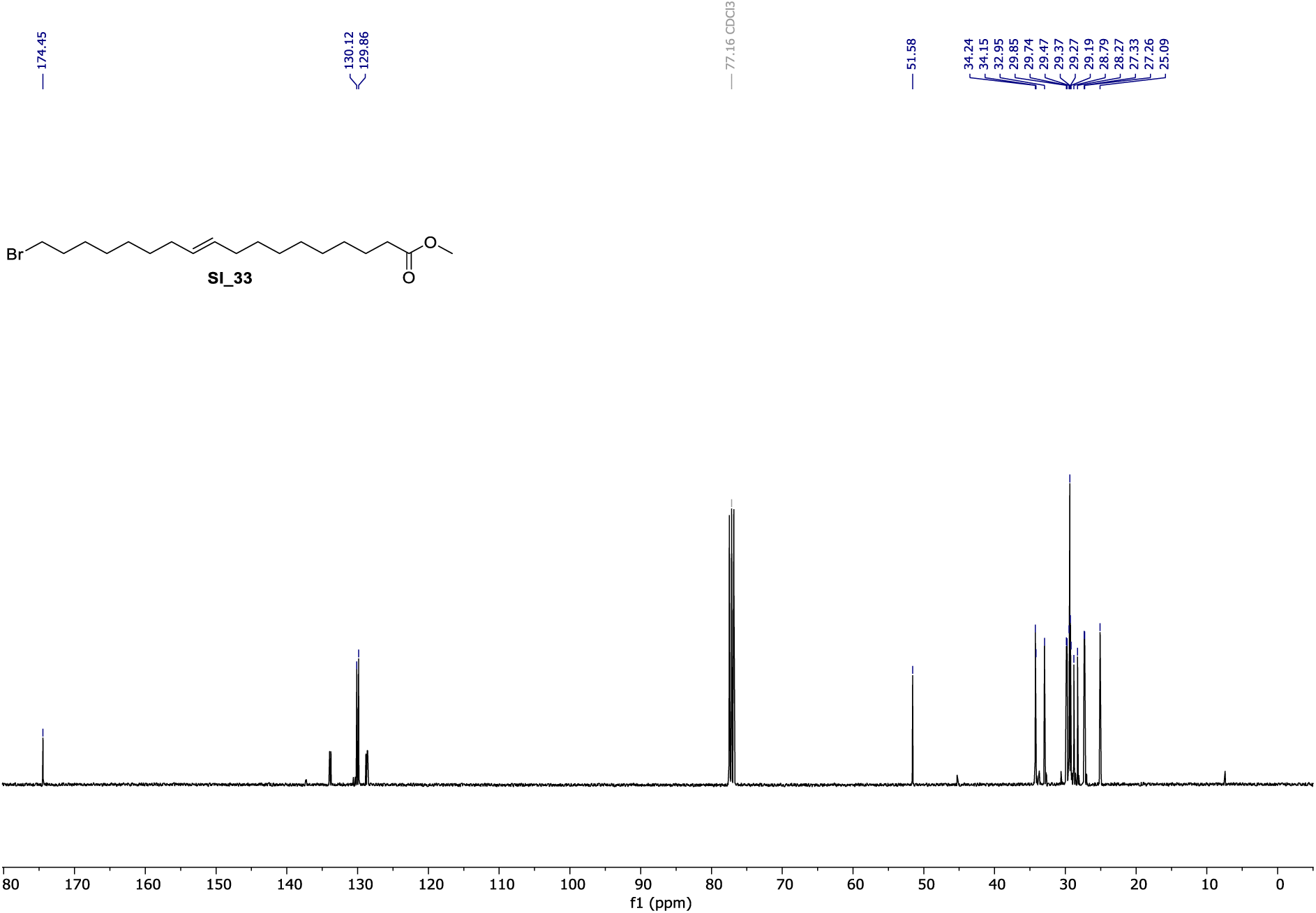

**^1^H NMR** (400 MHz, CDCl_3_) of **SI_35**.

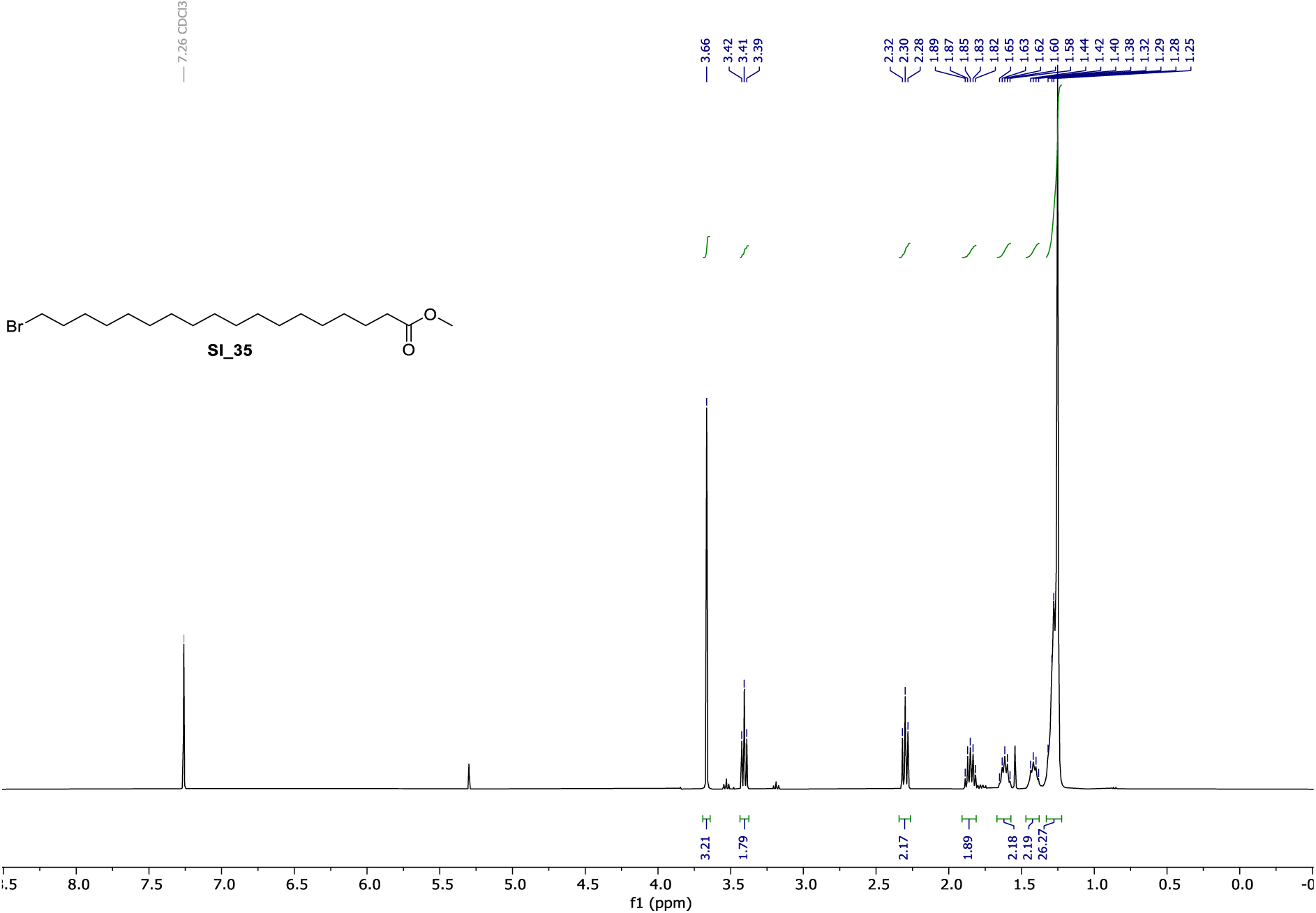

**^13^C NMR** (101 MHz, CDCl_3_) of **SI_35**.

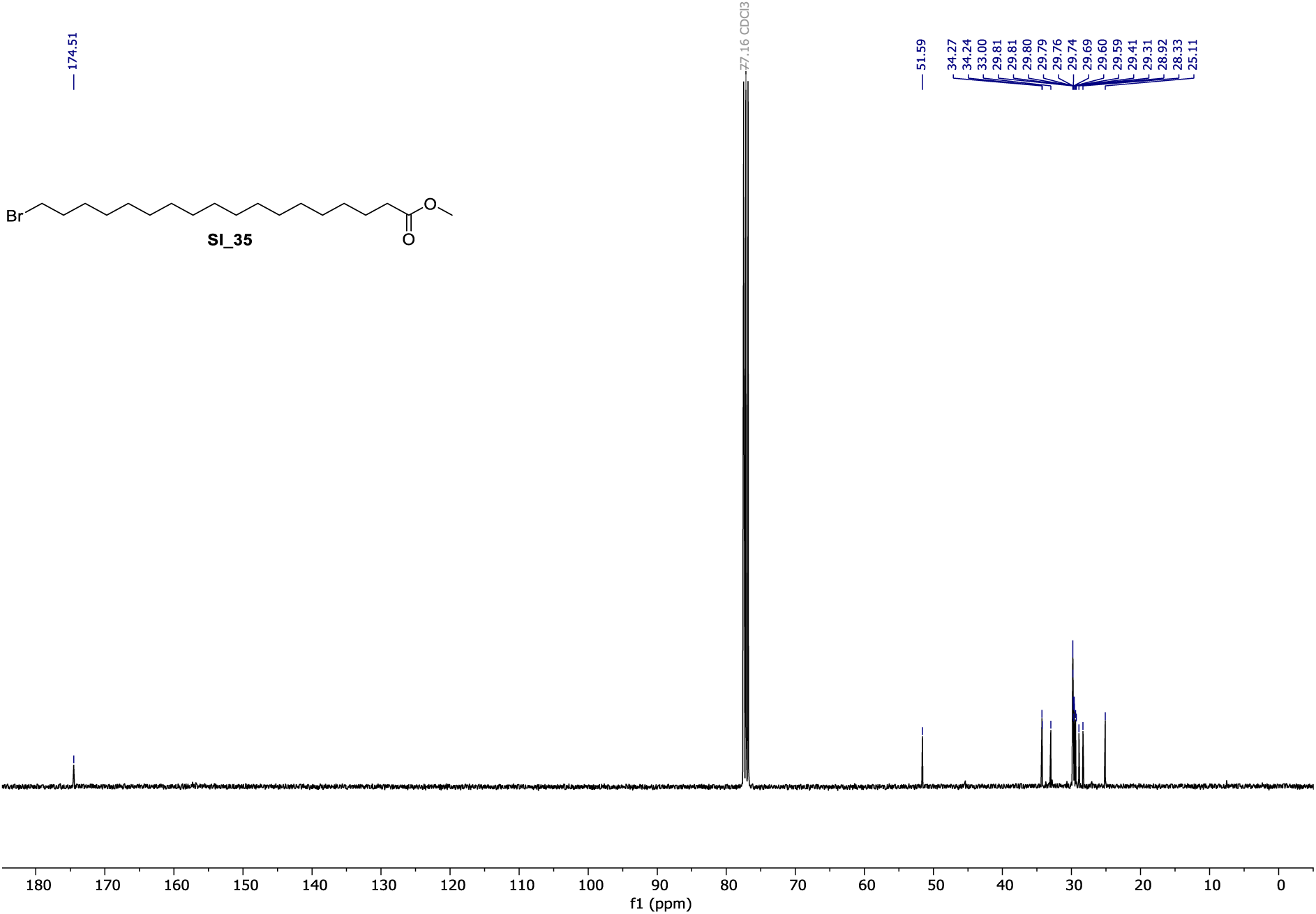

**^1^H NMR** (400 MHz, CDCl_3_) of **cl-SA**.

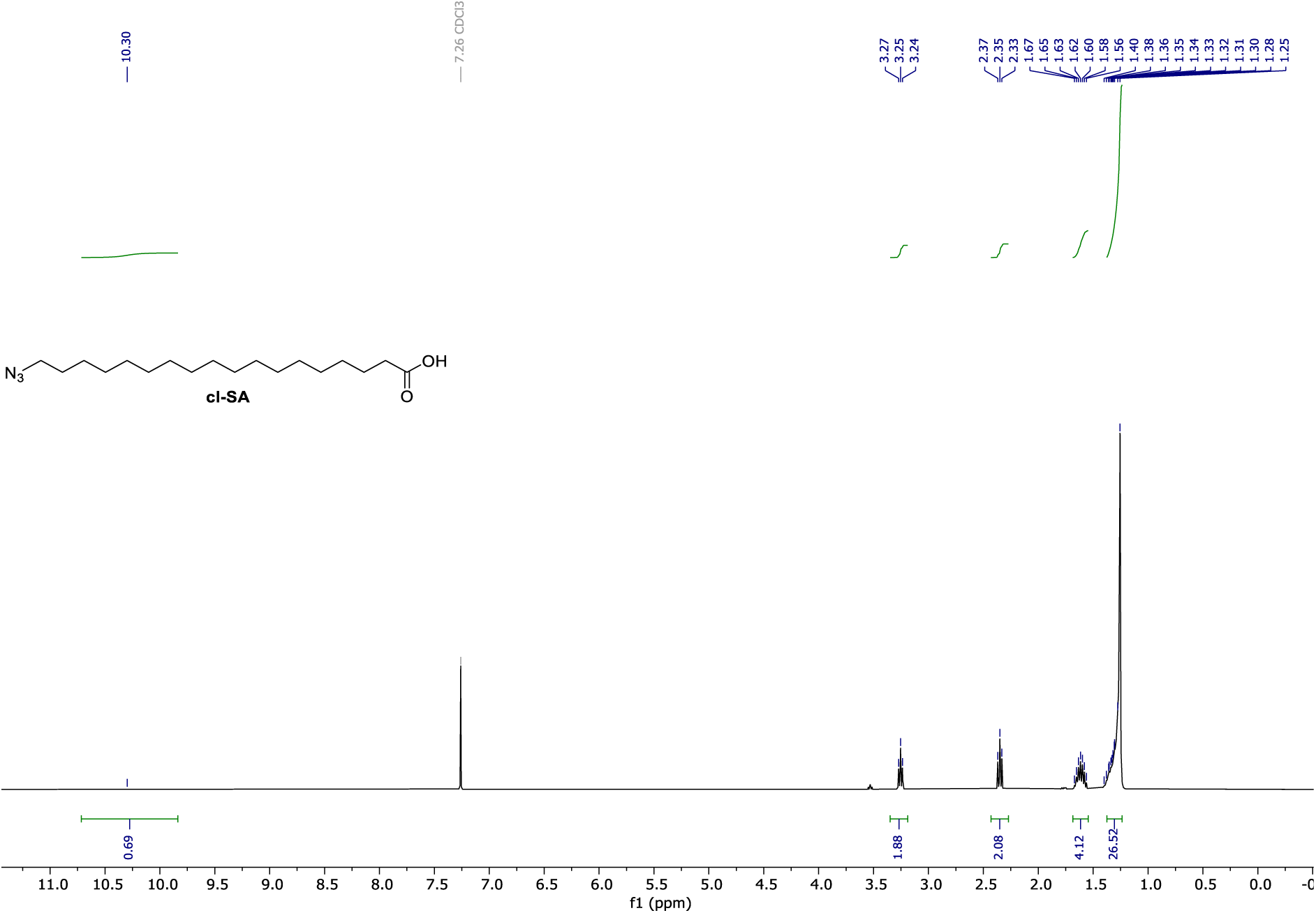

**^13^C NMR** (101 MHz, CDCl_3_) of **cl-SA**.

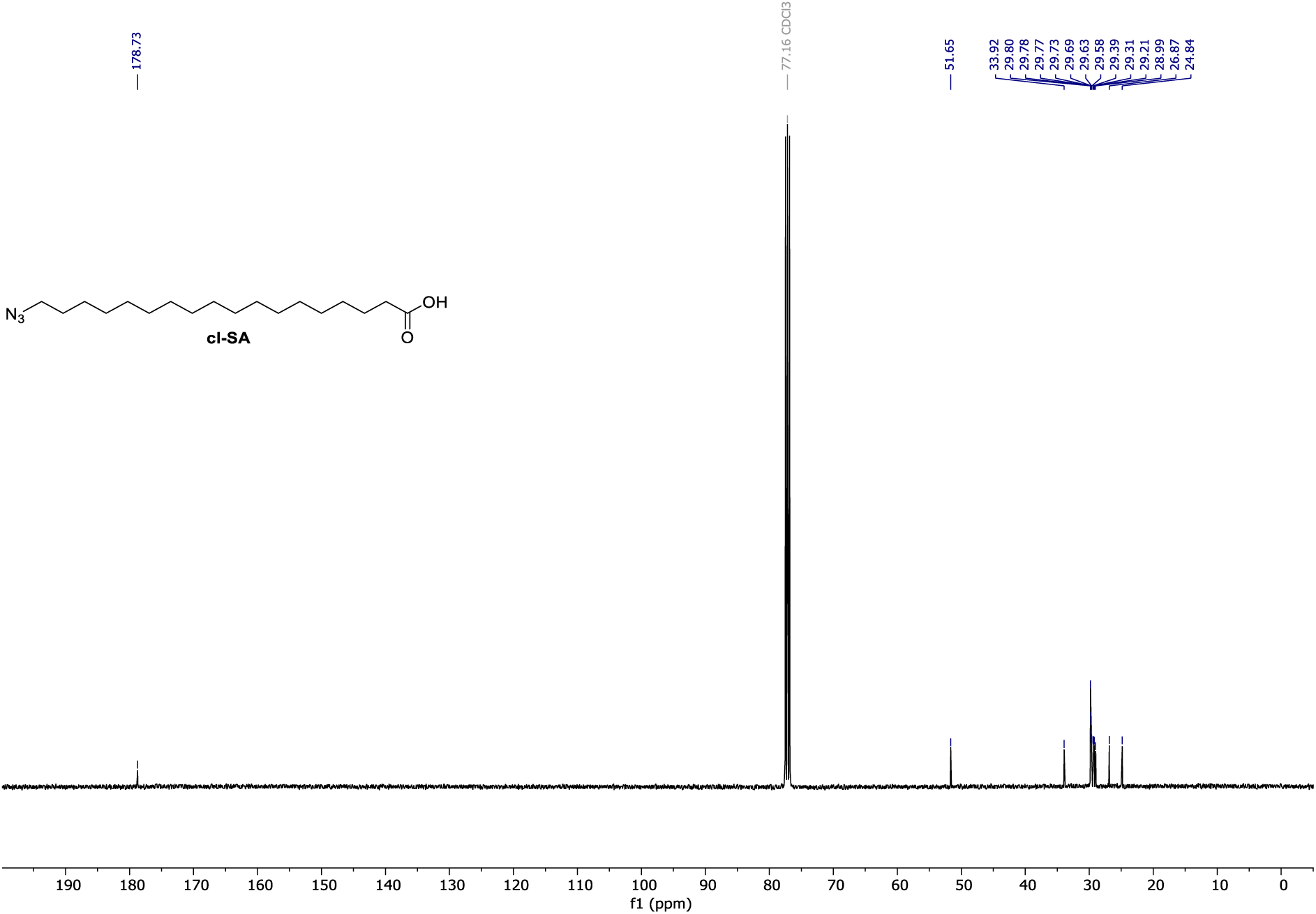

**^1^H NMR** (400 MHz, CDCl_3_) of **SI_37**.

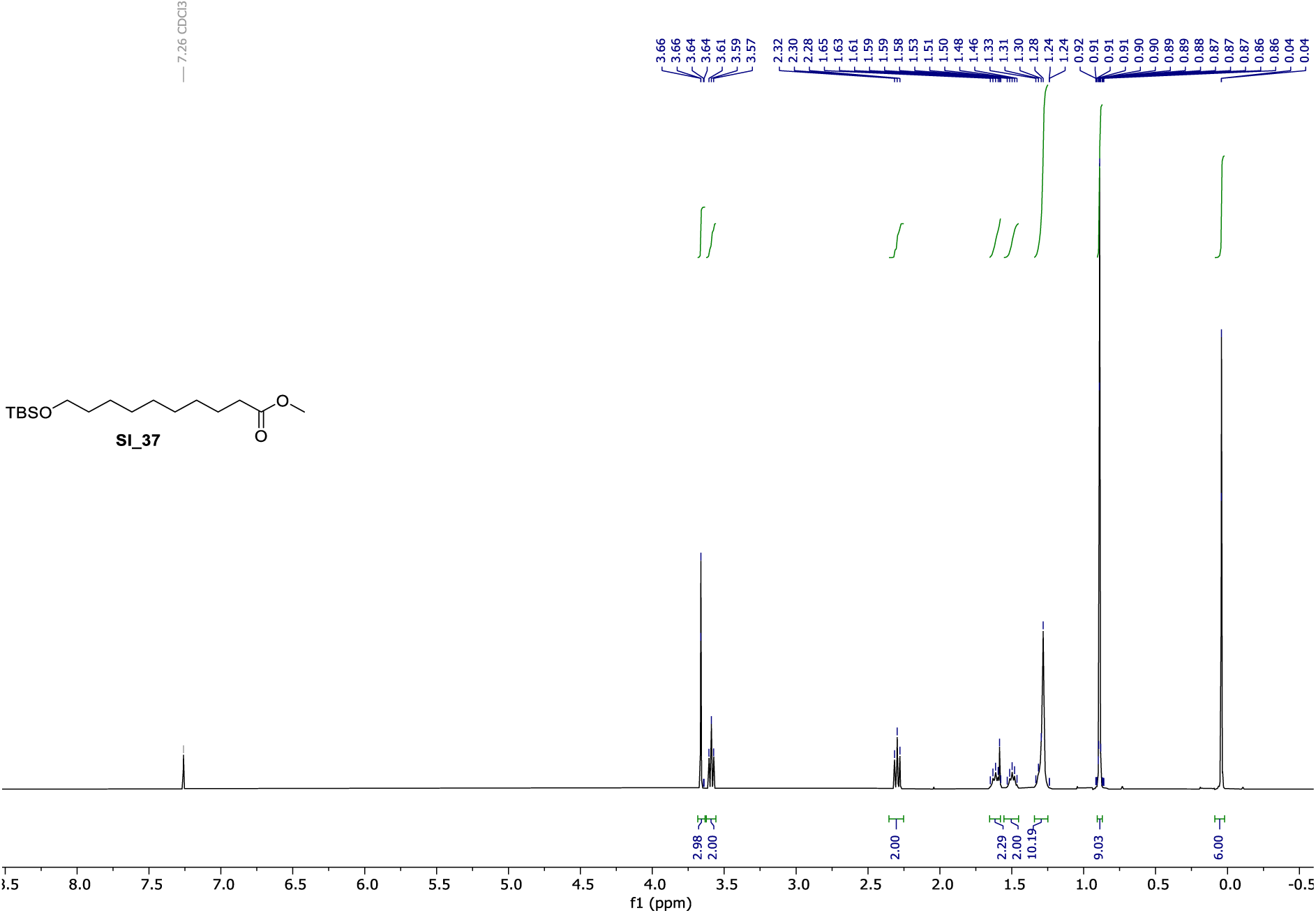

**^13^C NMR** (101 MHz, CDCl_3_) of **SI_37**.

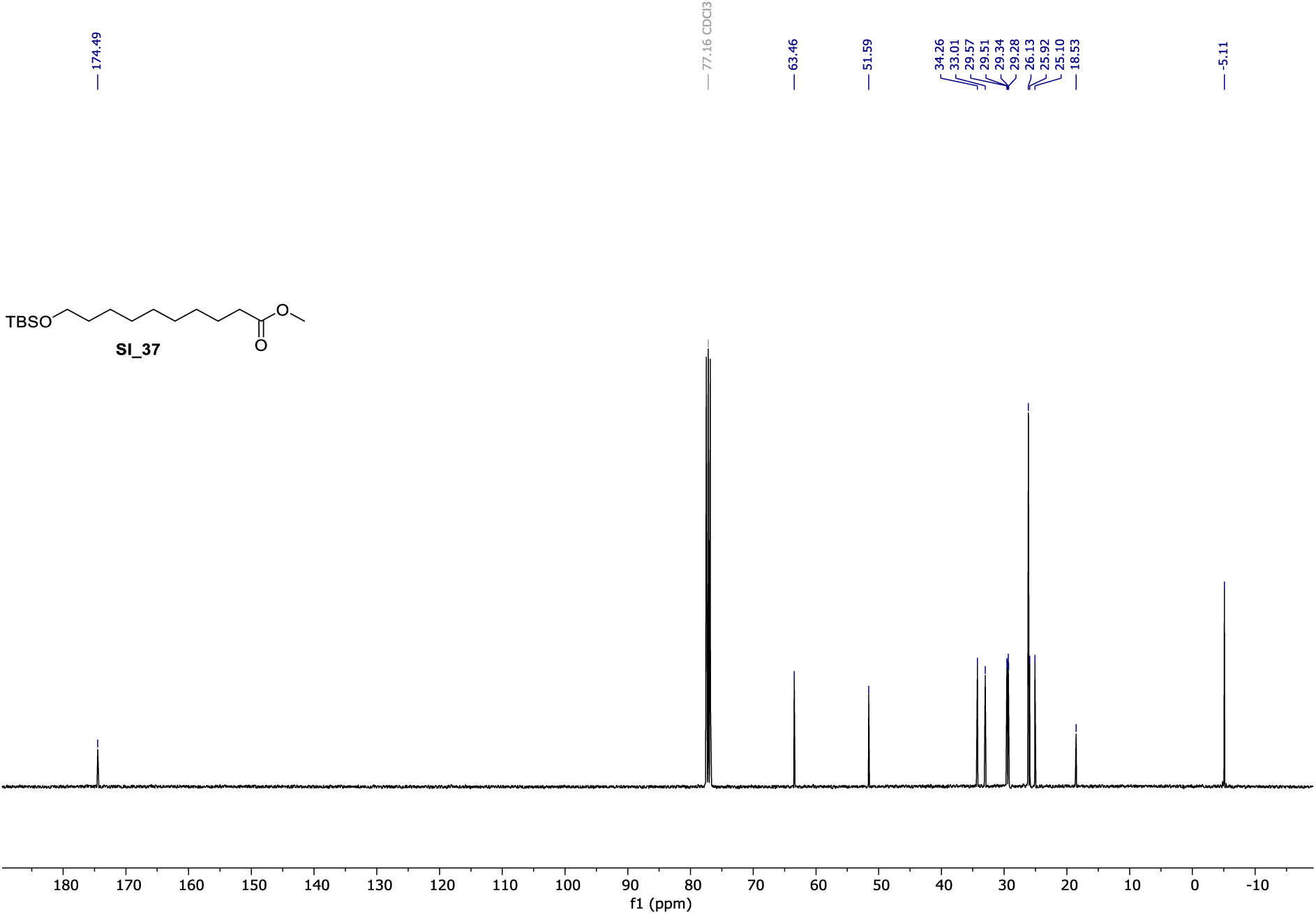

**^1^H NMR** (400 MHz, CDCl_3_) of **SI_38**.

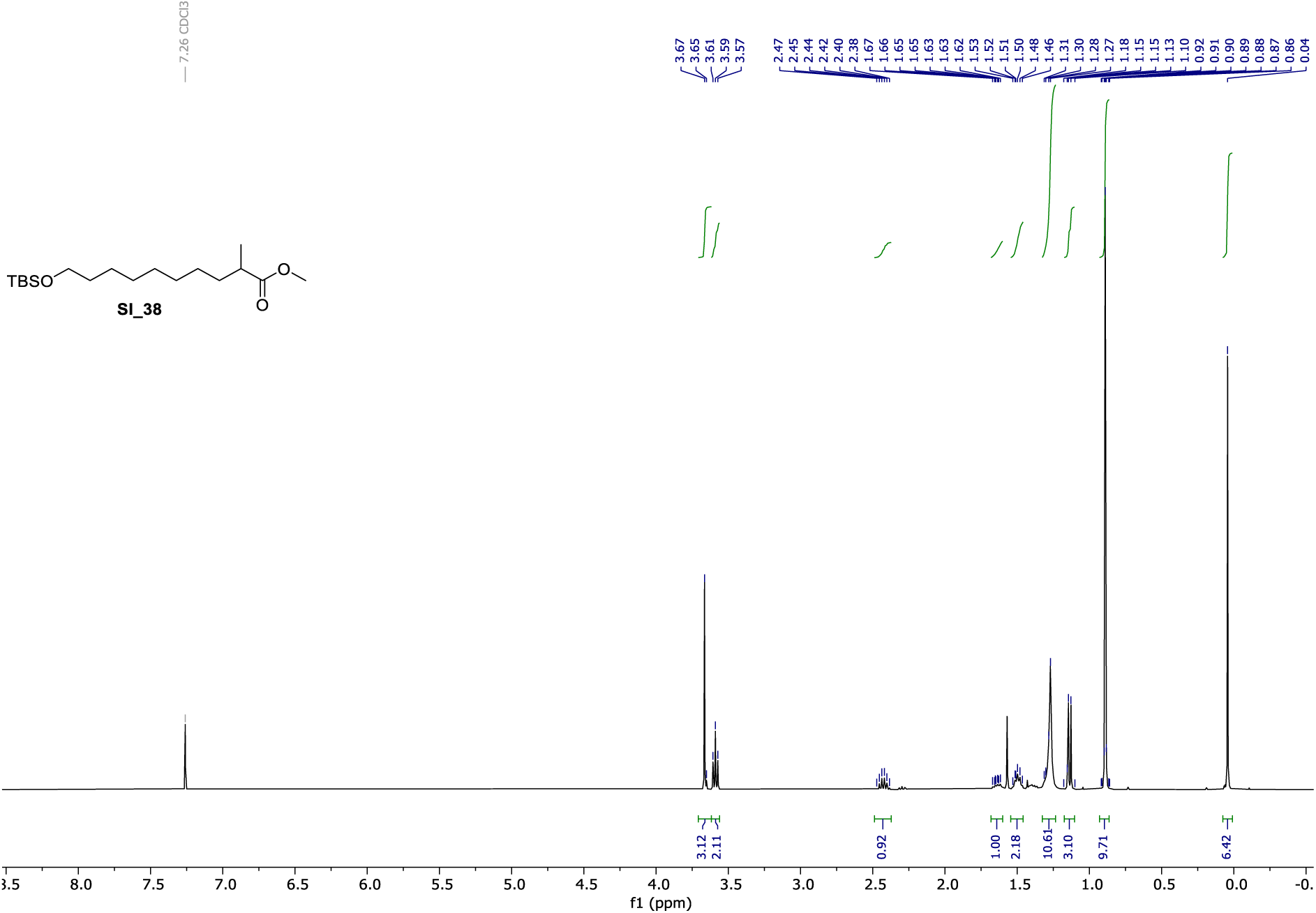

**^13^C NMR** (101 MHz, CDCl_3_) of **SI_38**.

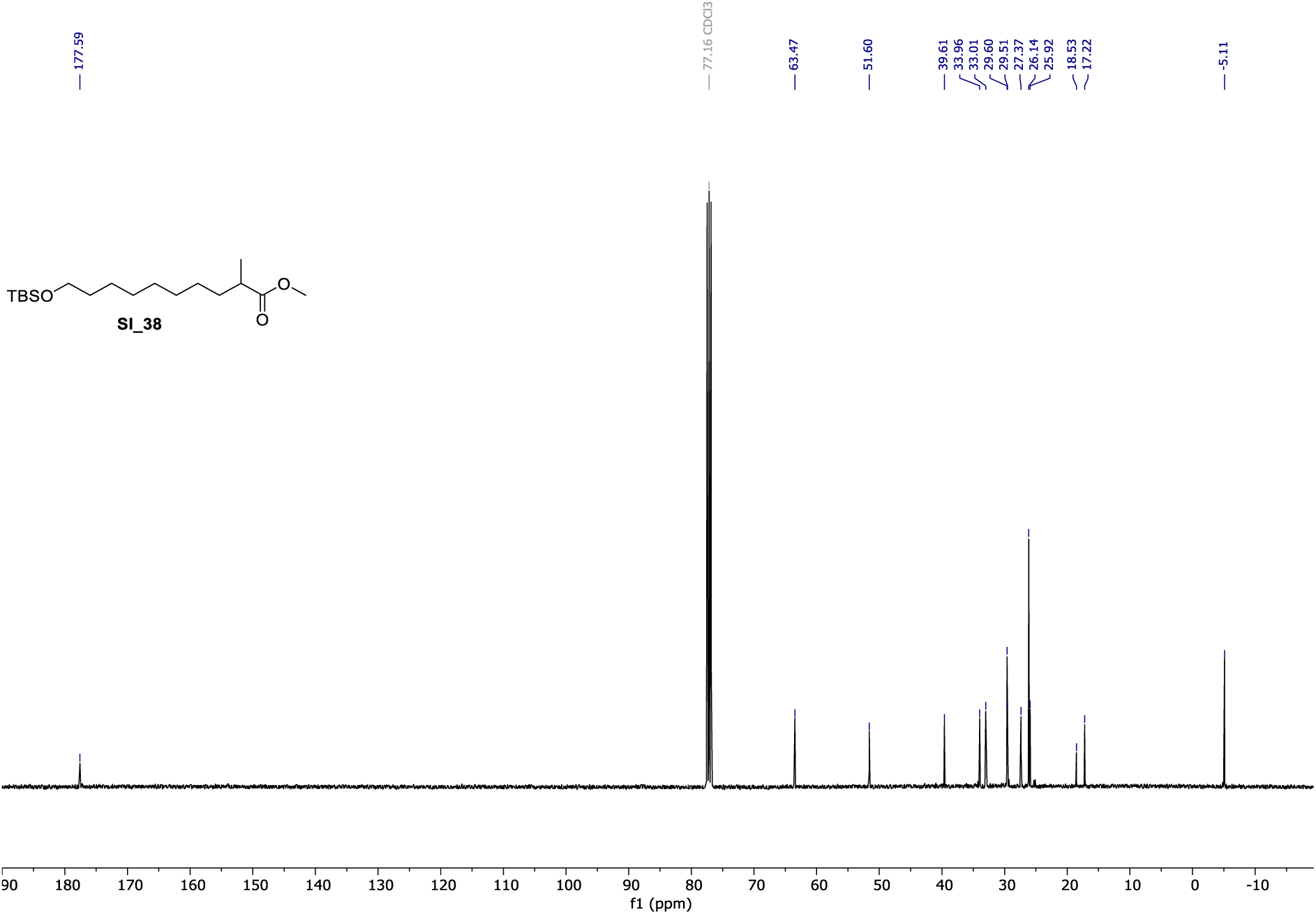

**^1^H NMR** (400 MHz, CDCl_3_) of **SI_39**.

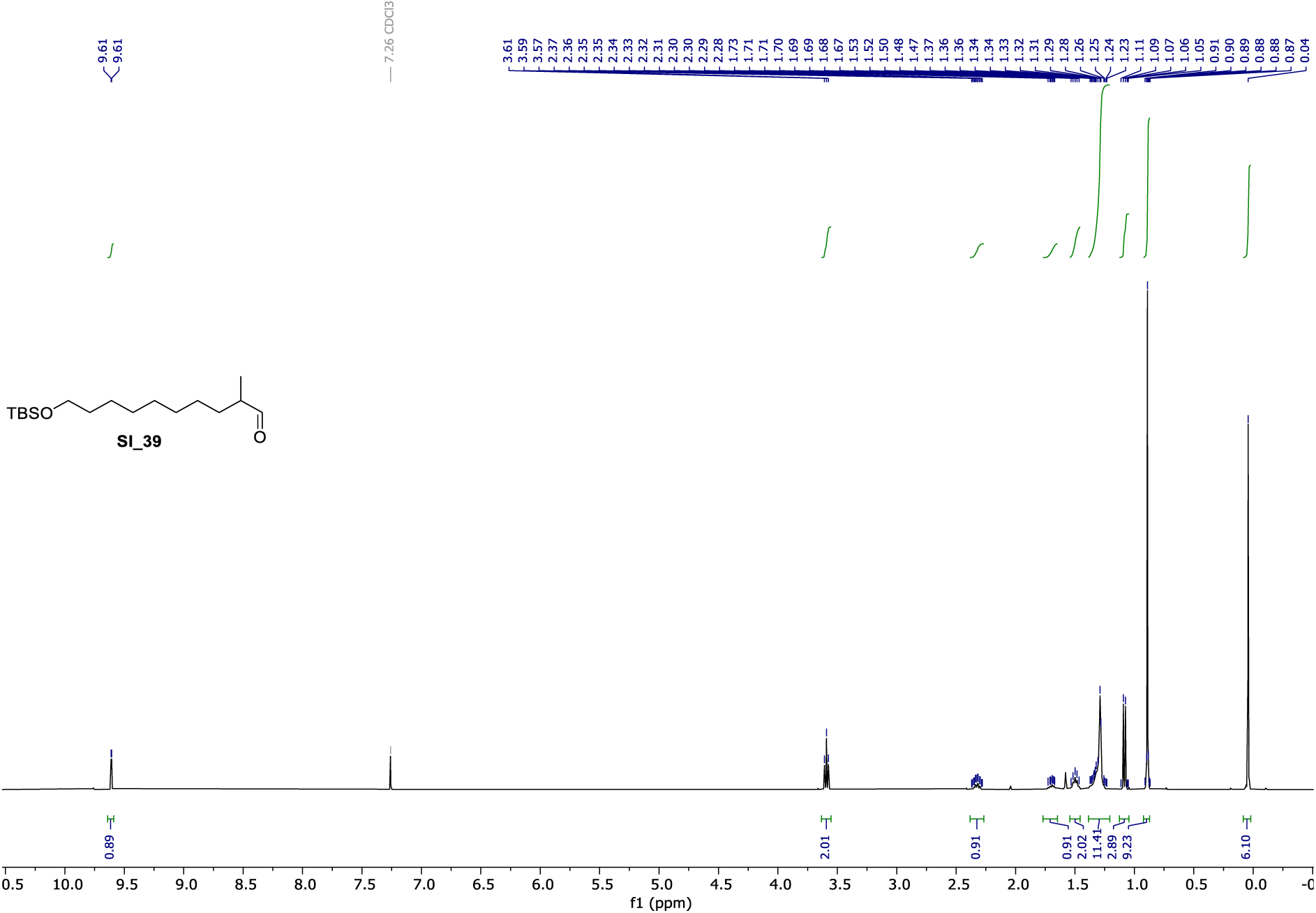

**^13^C NMR** (101 MHz, CDCl_3_) of **SI_39**.

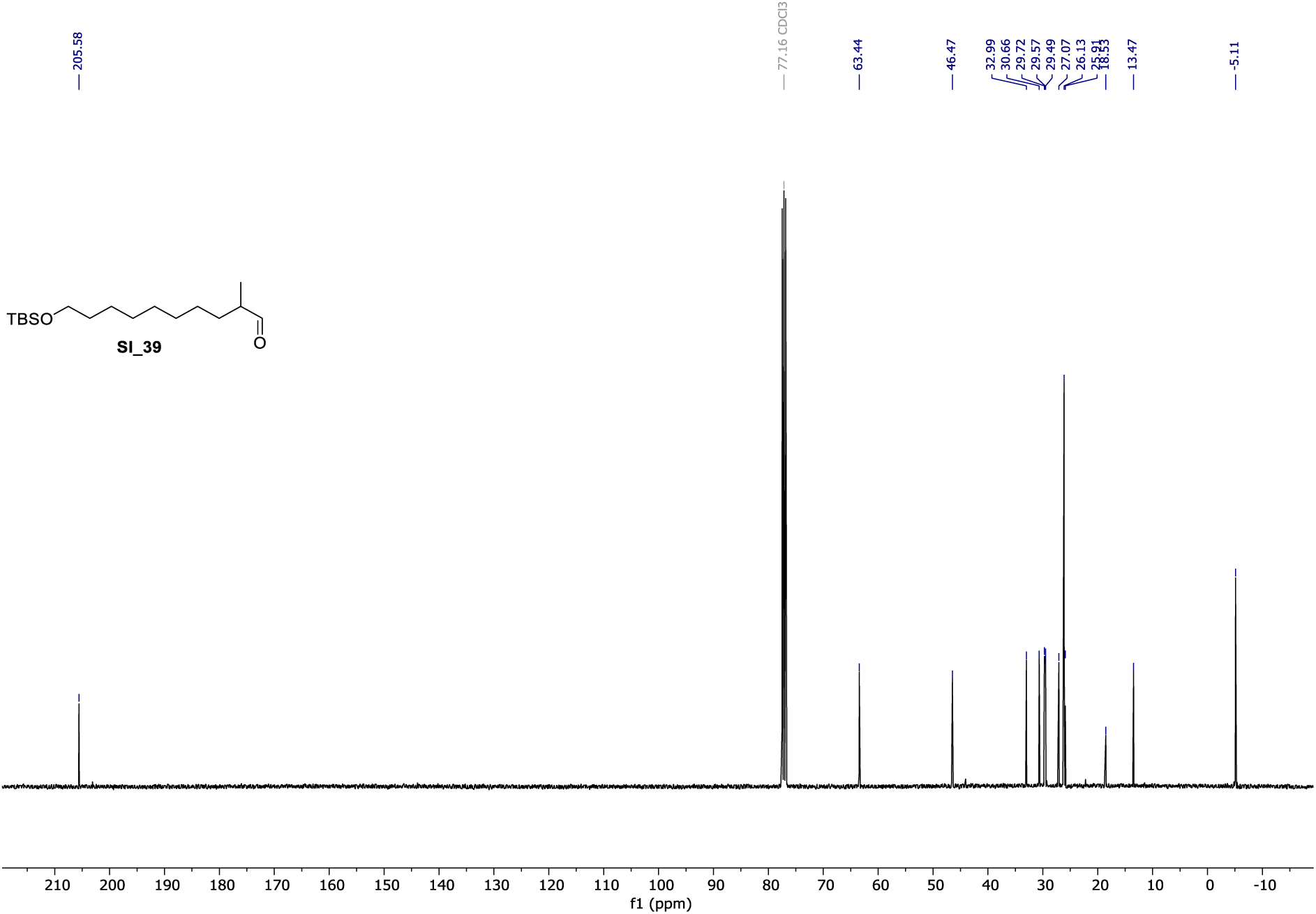

**^1^H NMR** (400 MHz, CDCl_3_) of **SI_41**.

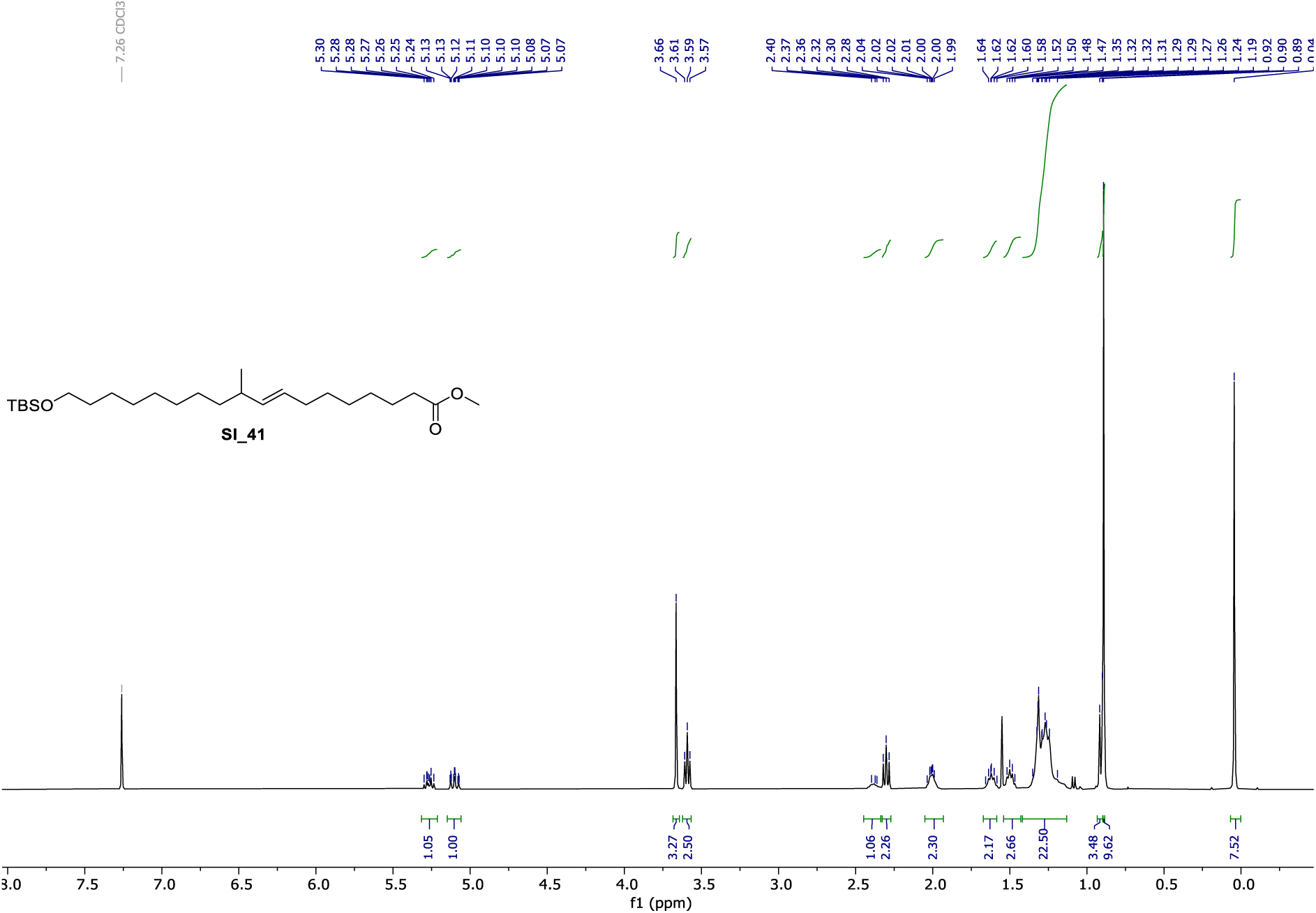

**^13^C NMR** (101 MHz, CDCl_3_) of **SI_41**.

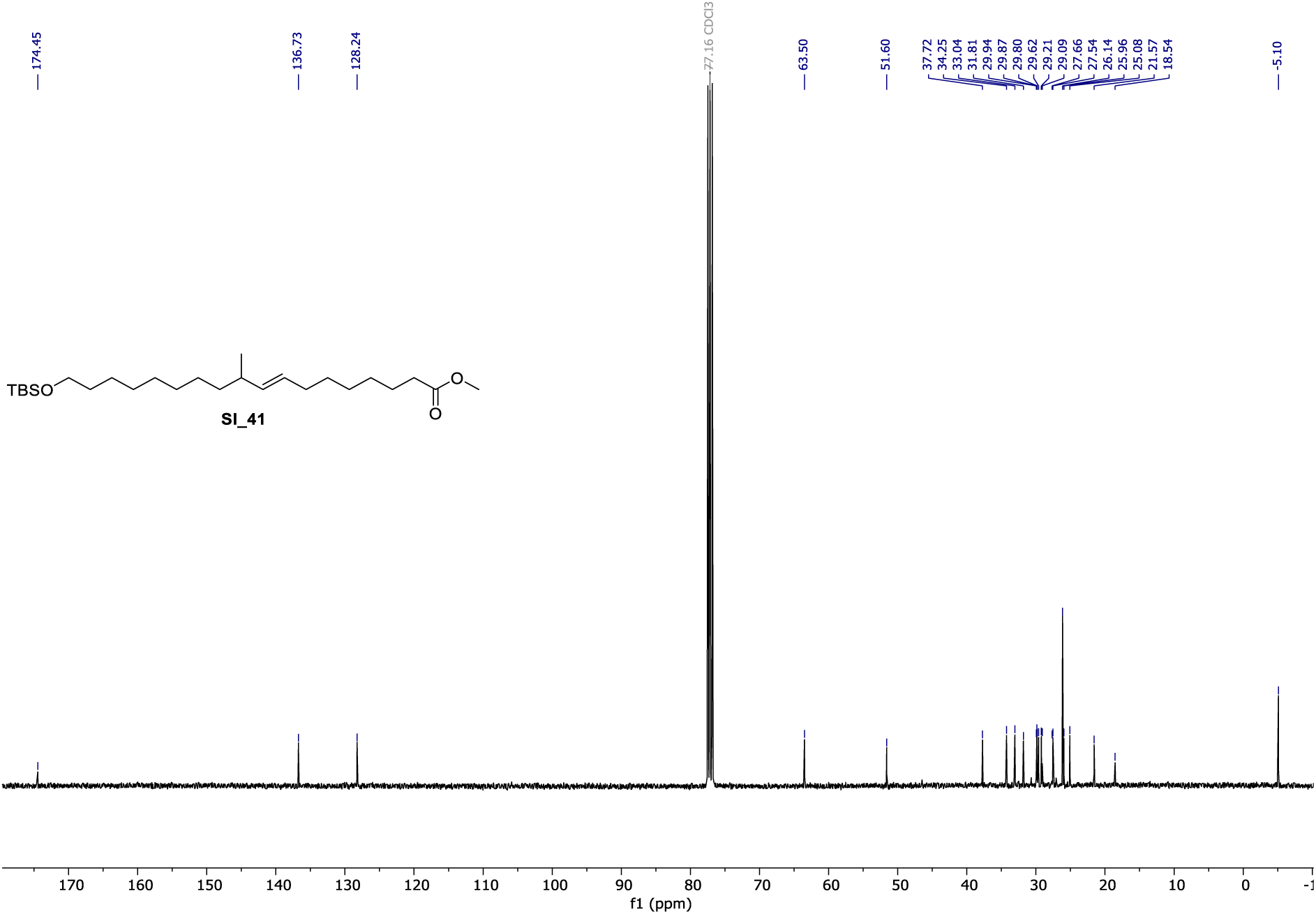

**^1^H NMR** (400 MHz, CDCl_3_) of **SI_42**.

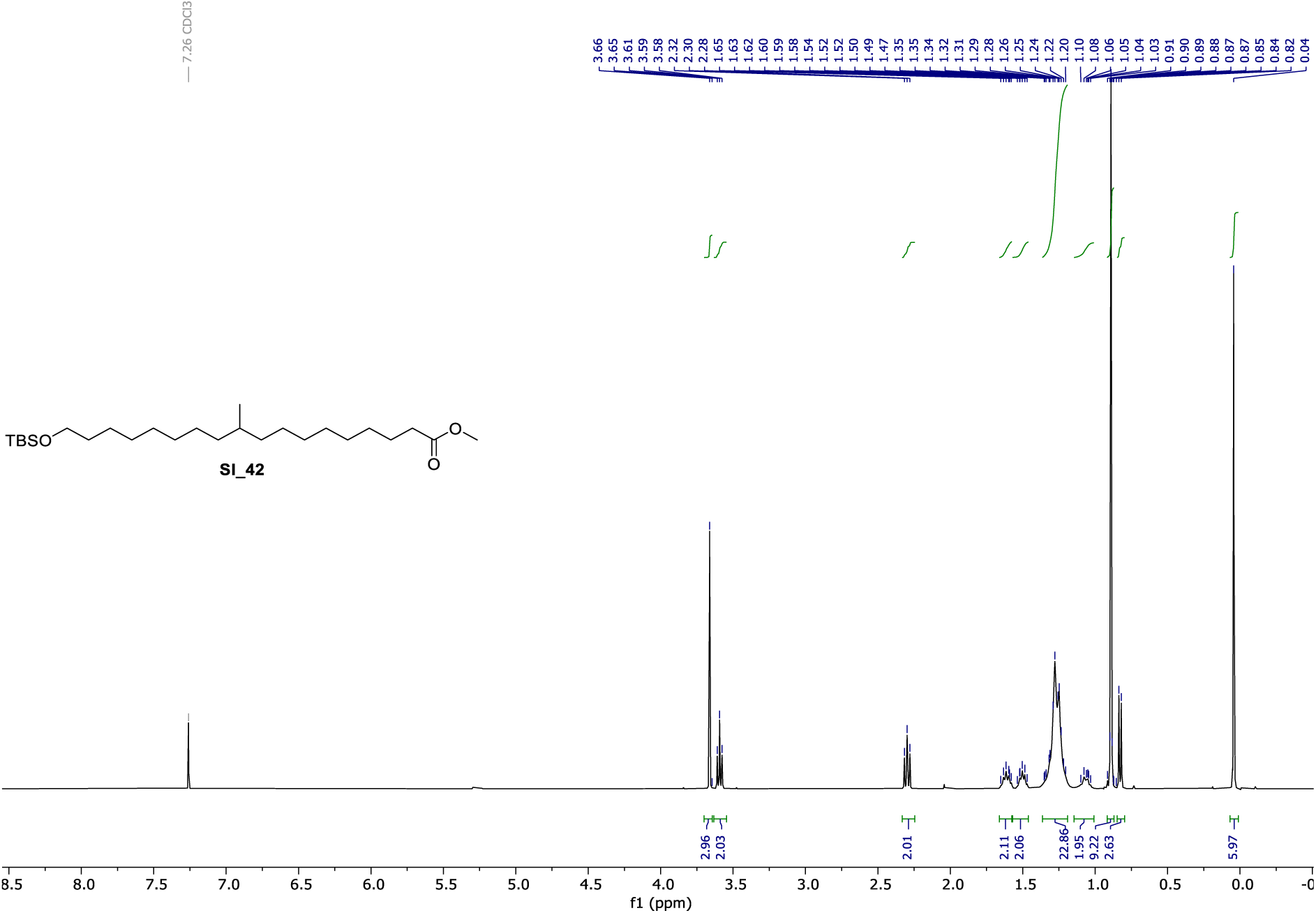

**^13^C NMR** (101 MHz, CDCl_3_) of **SI_42**.

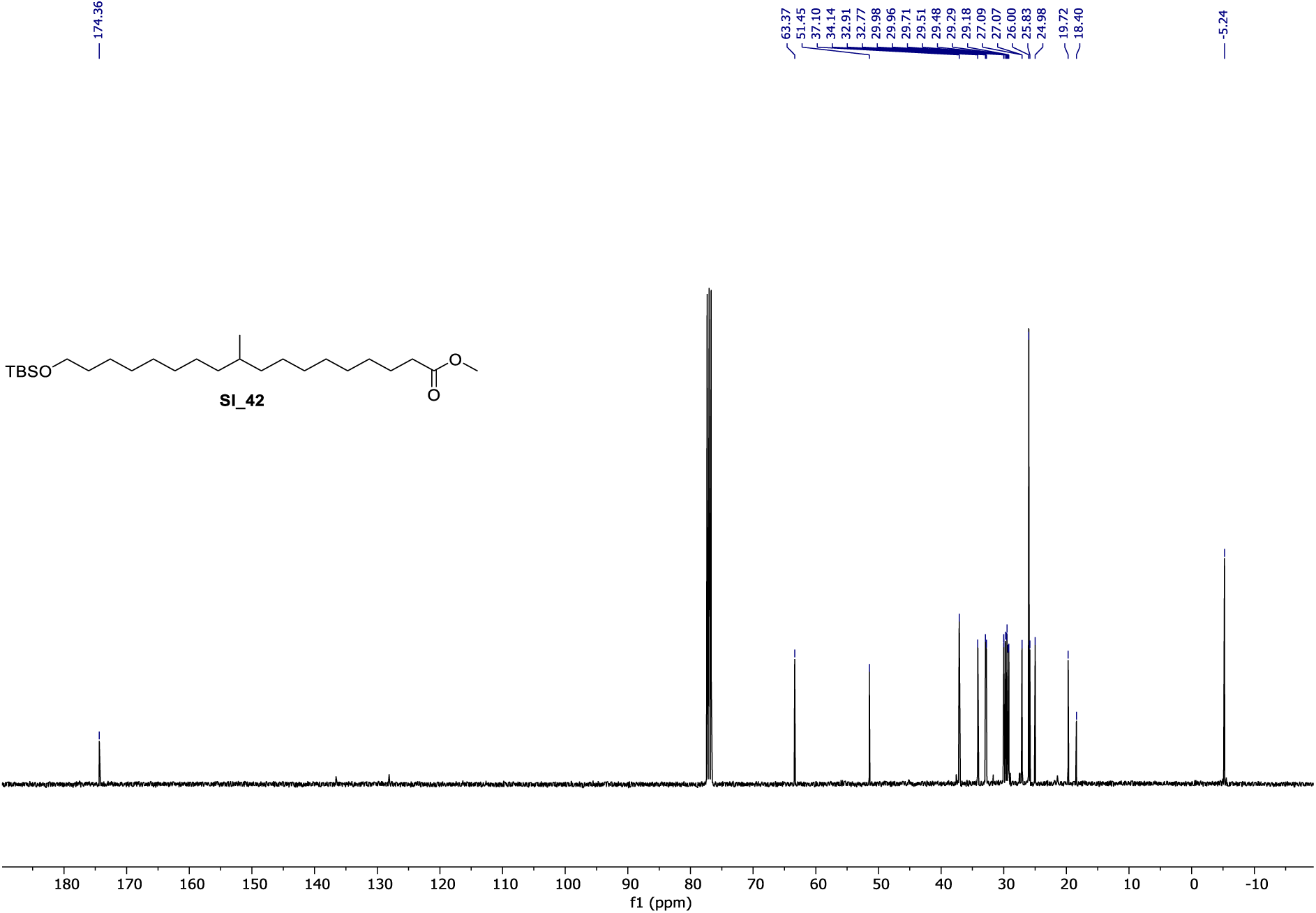

**^1^H NMR** (400 MHz, CDCl_3_) of **SI_43**.

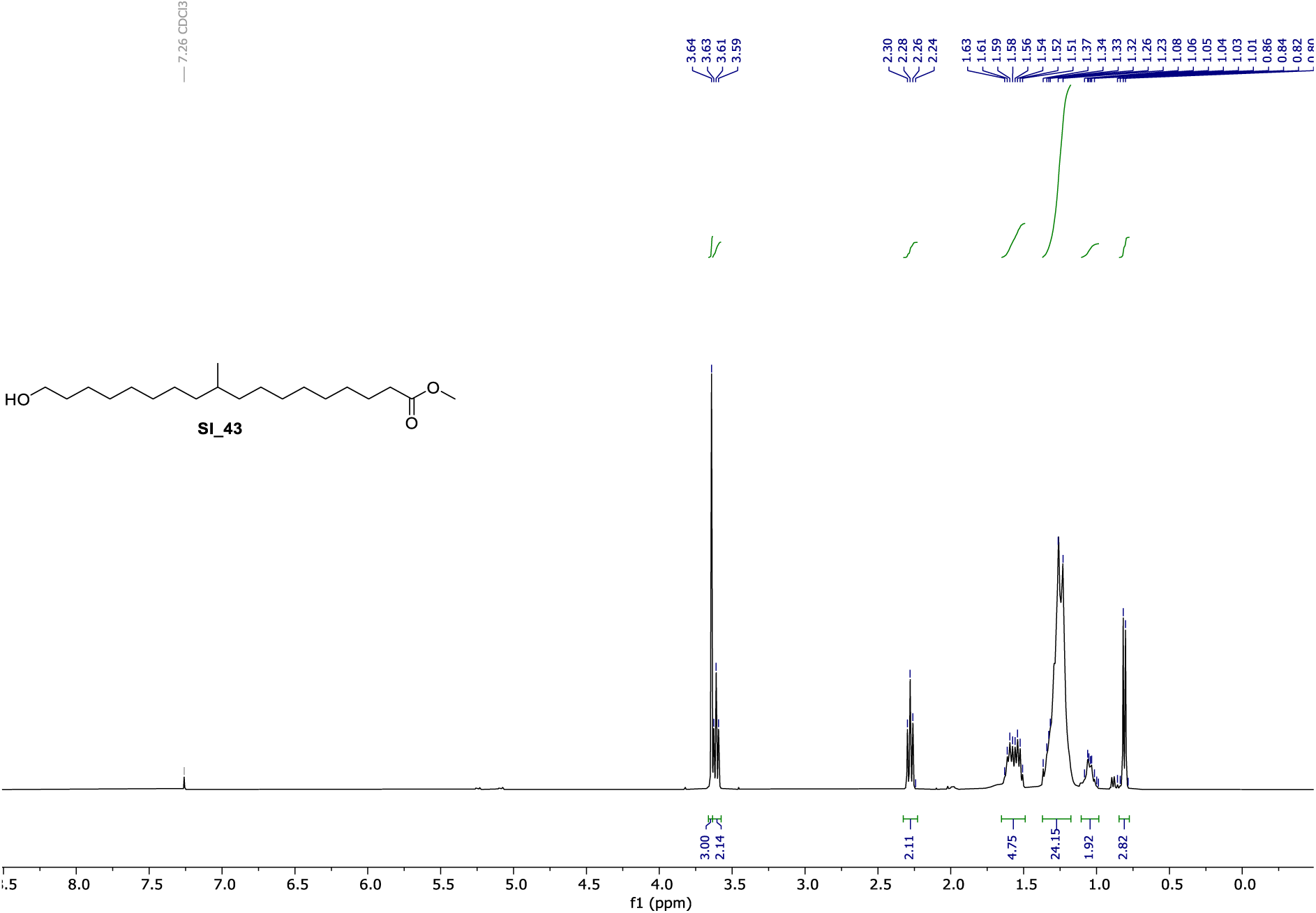

**^13^C NMR** (101 MHz, CDCl_3_) of **SI_43**.

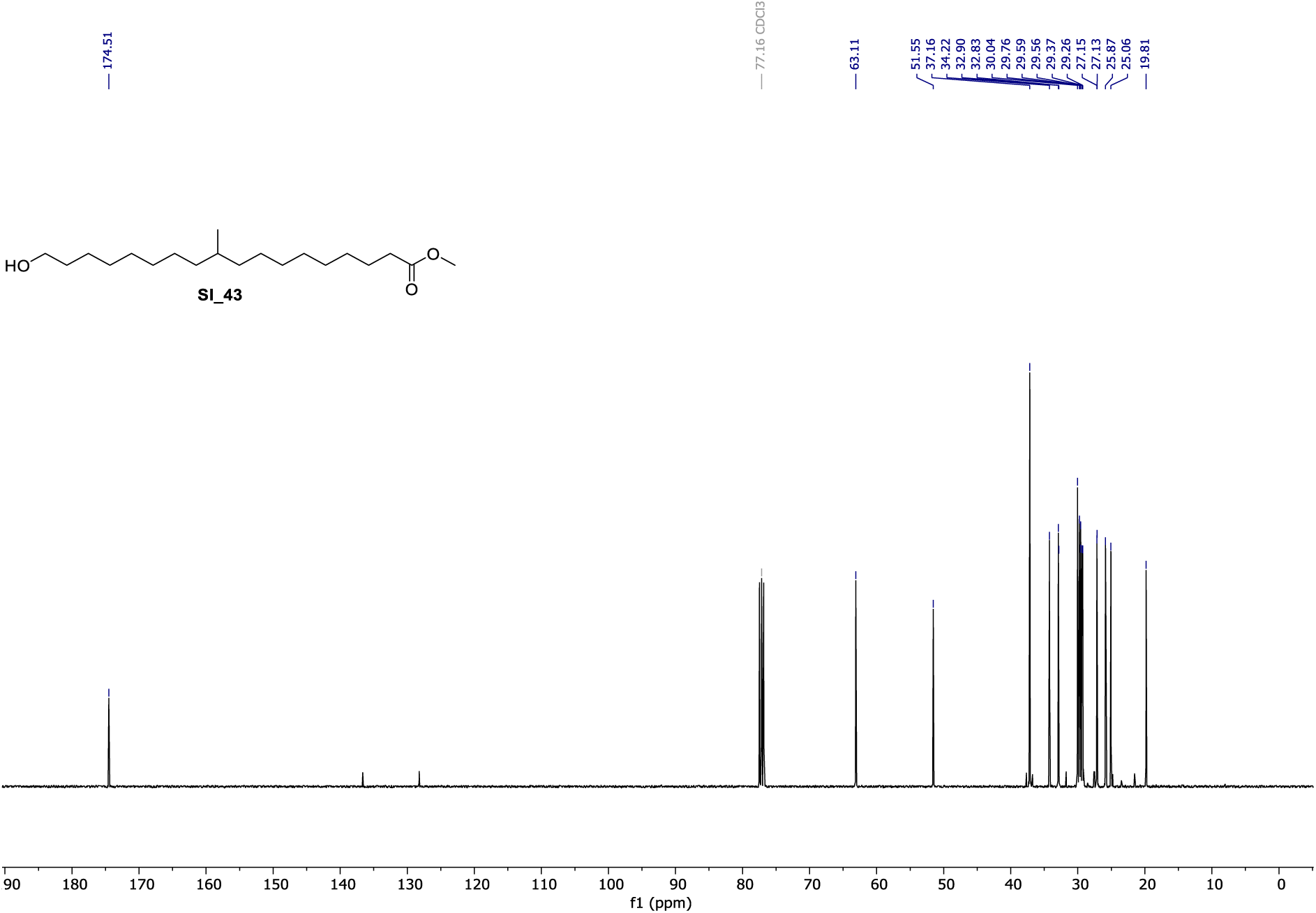

**^1^H NMR** (400 MHz, CDCl_3_) of **SI_44**.

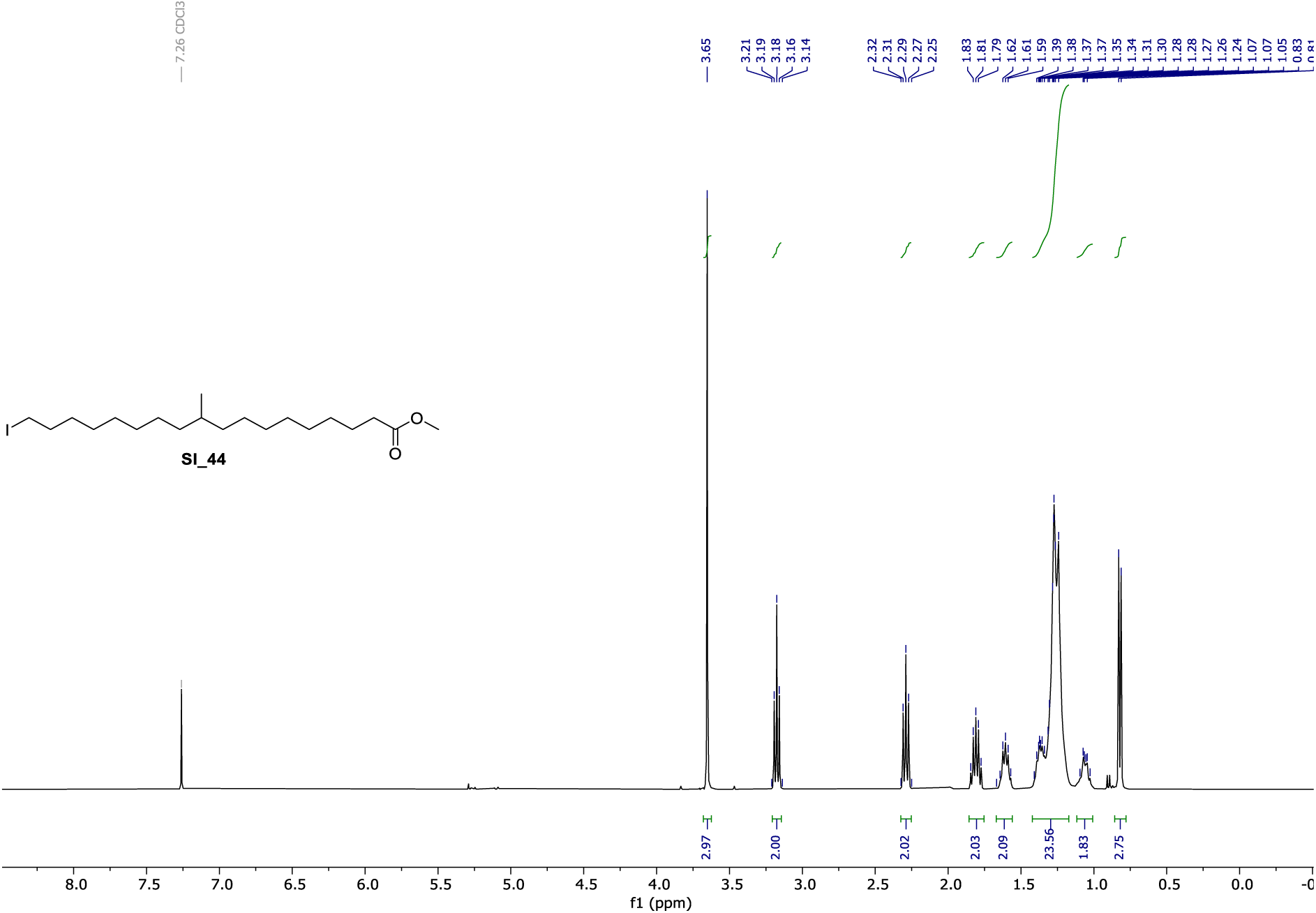

**^13^C NMR** (101 MHz, CDCl_3_) of **SI_44**.

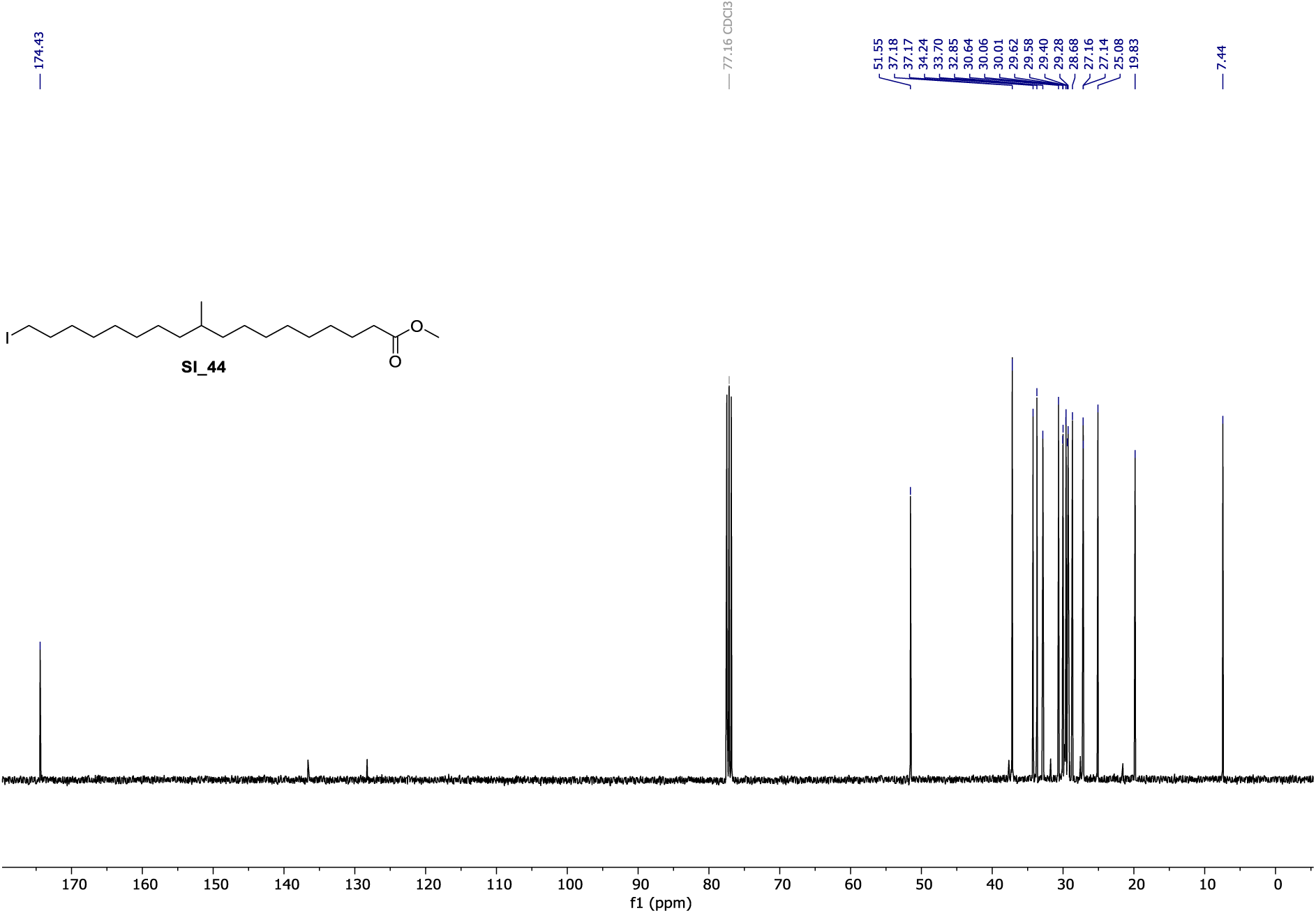

**^1^H NMR** (400 MHz, CDCl_3_) of **cl-10-MeSA**.

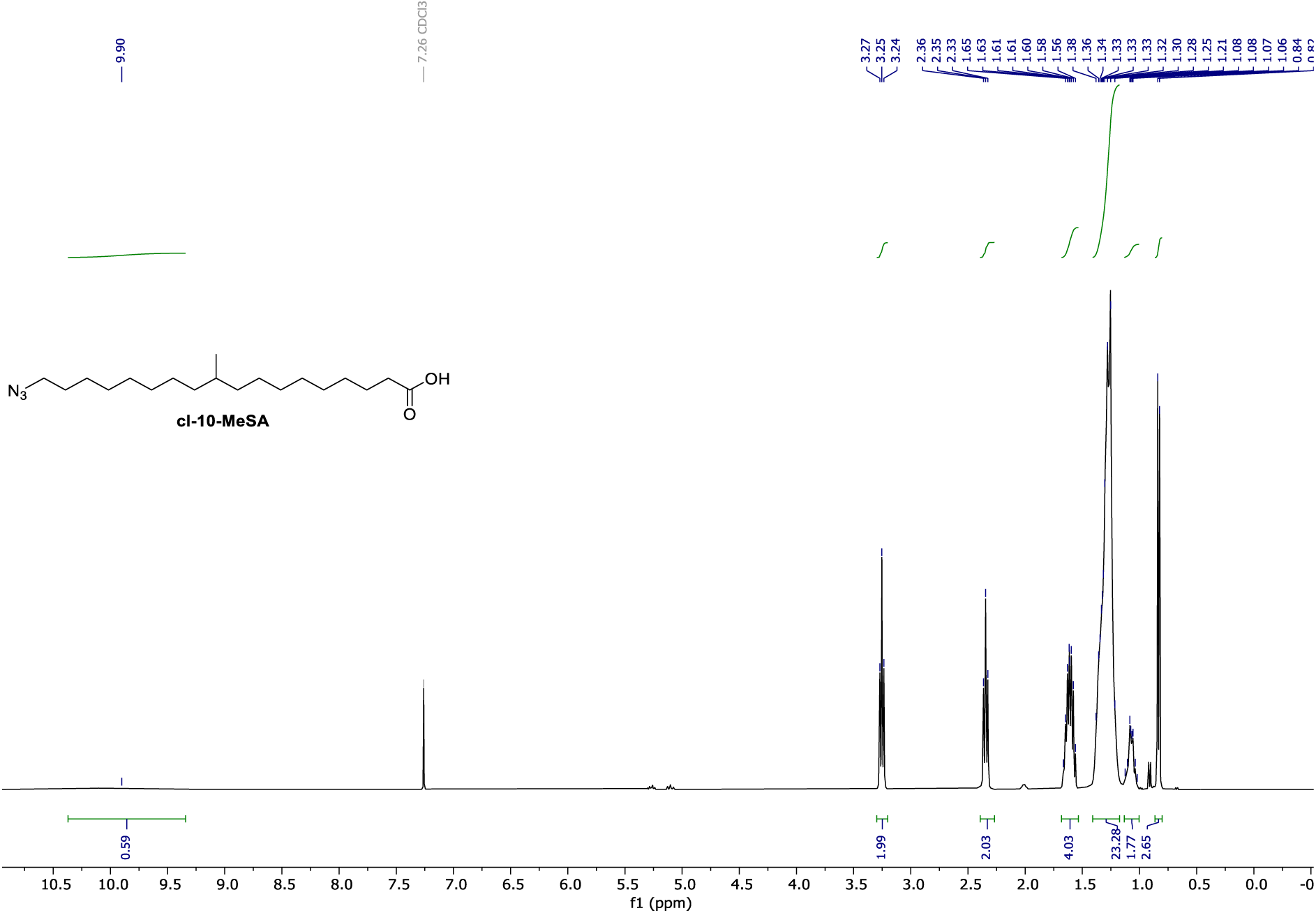

**^13^C NMR** (101 MHz, CDCl_3_) of **cl-10-MeSA**.

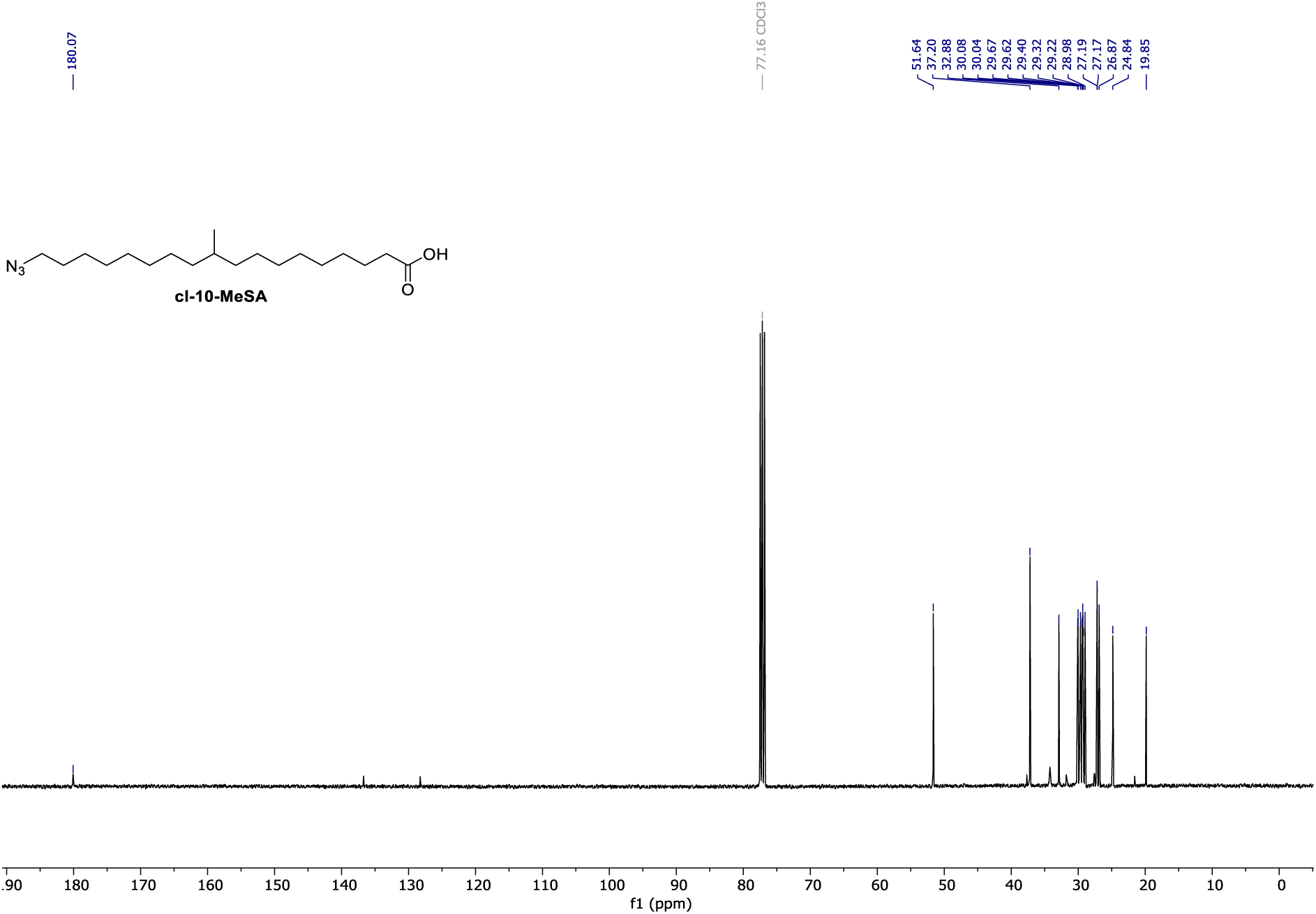

**^1^H NMR** (400 MHz, CDCl_3_) of **SI_45**.

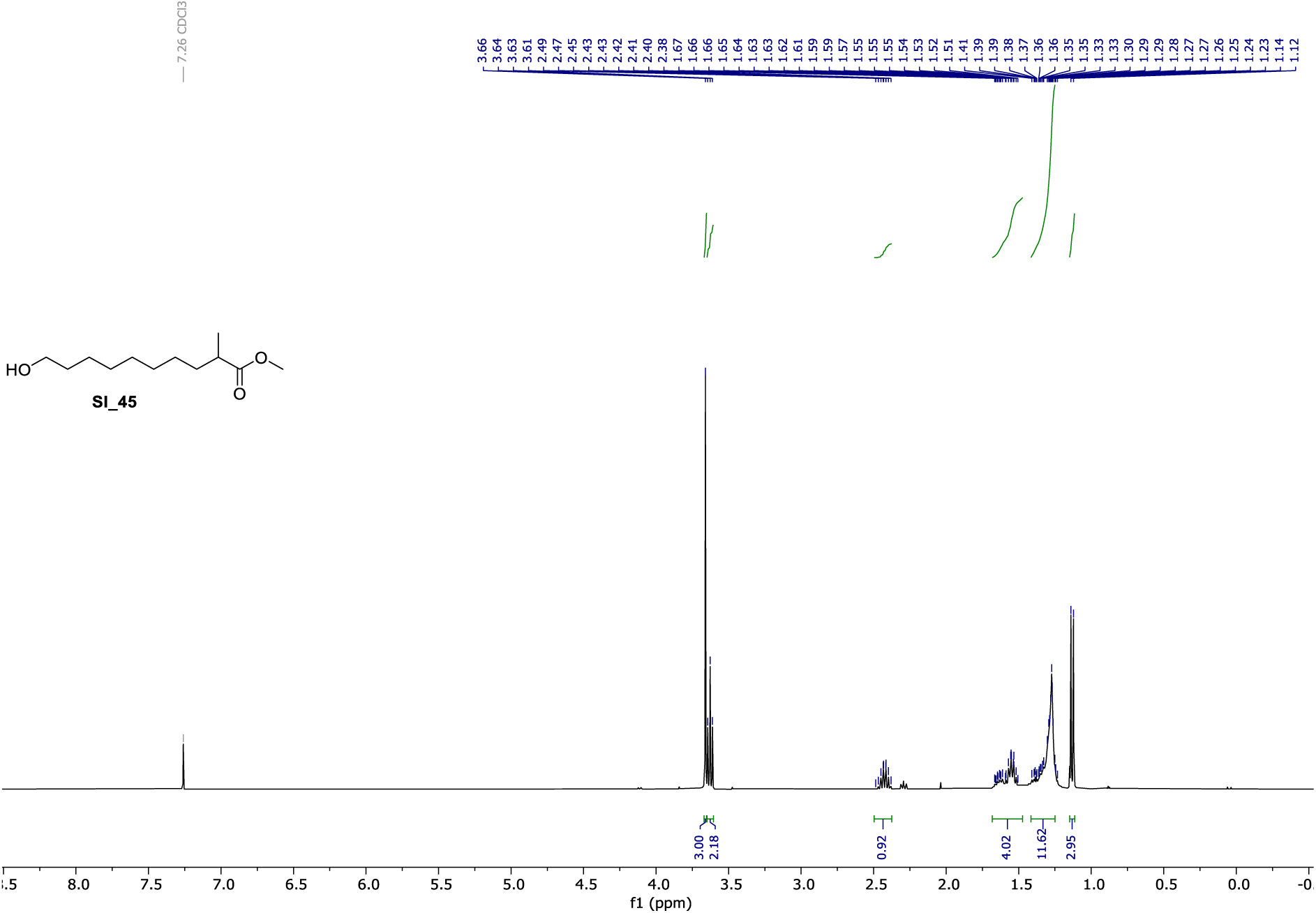

**^13^C NMR** (101 MHz, CDCl_3_) of **SI_45**.

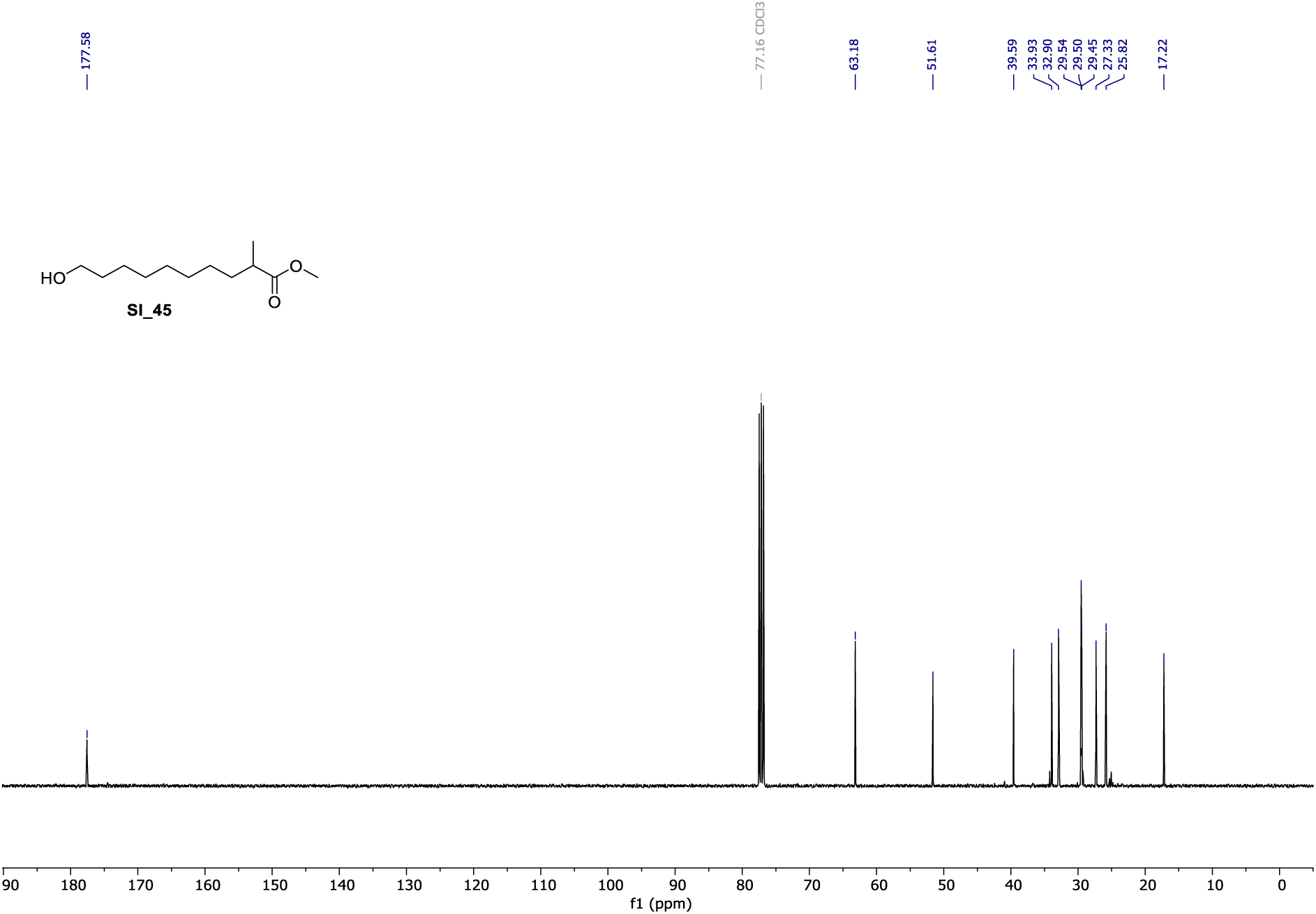

**^1^H NMR** (400 MHz, CDCl_3_) of **SI_46**.

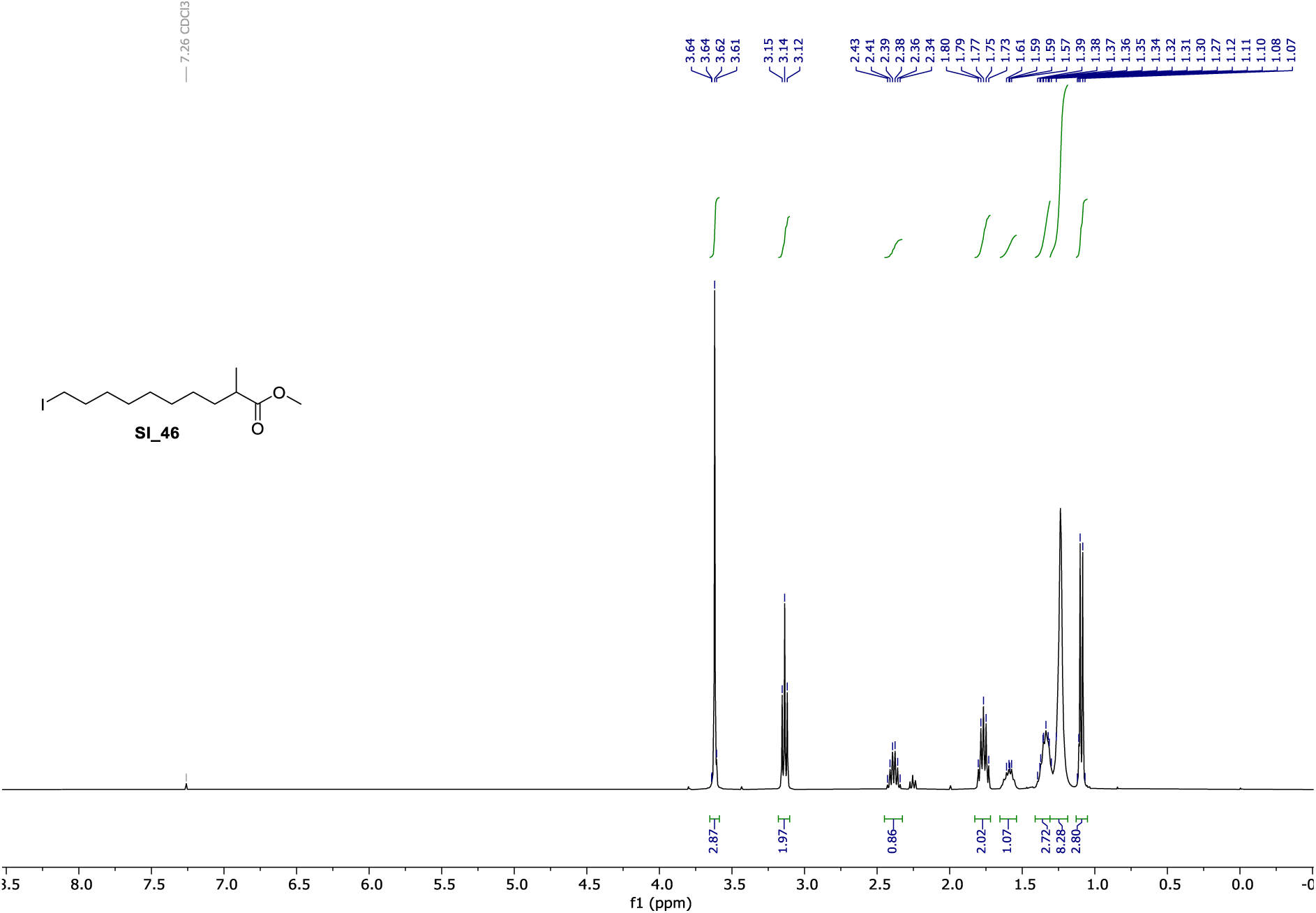

**^13^C NMR** (101 MHz, CDCl_3_) of **SI_46**.

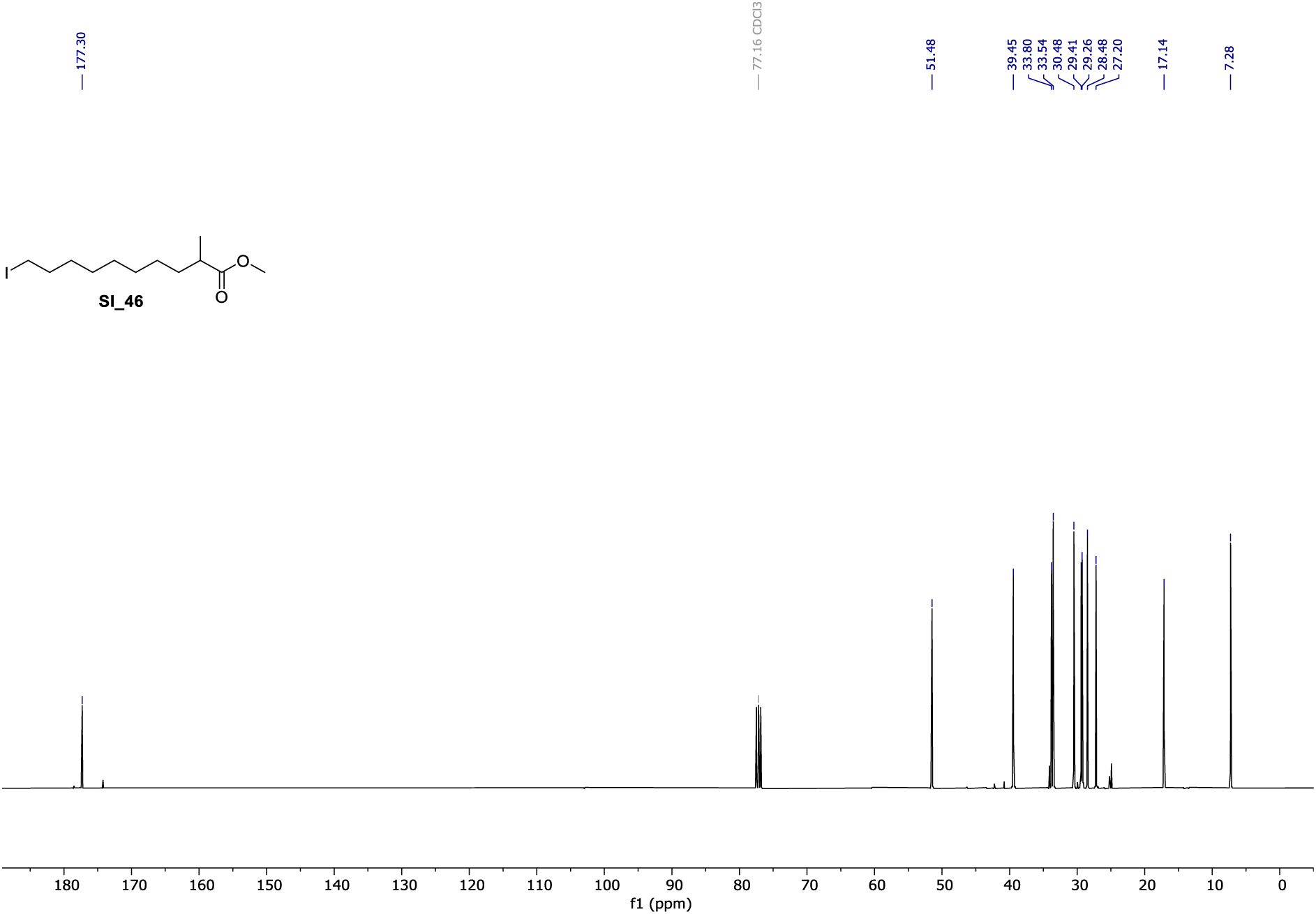

**^1^H NMR** (400 MHz, CDCl_3_) of **SI_48**.

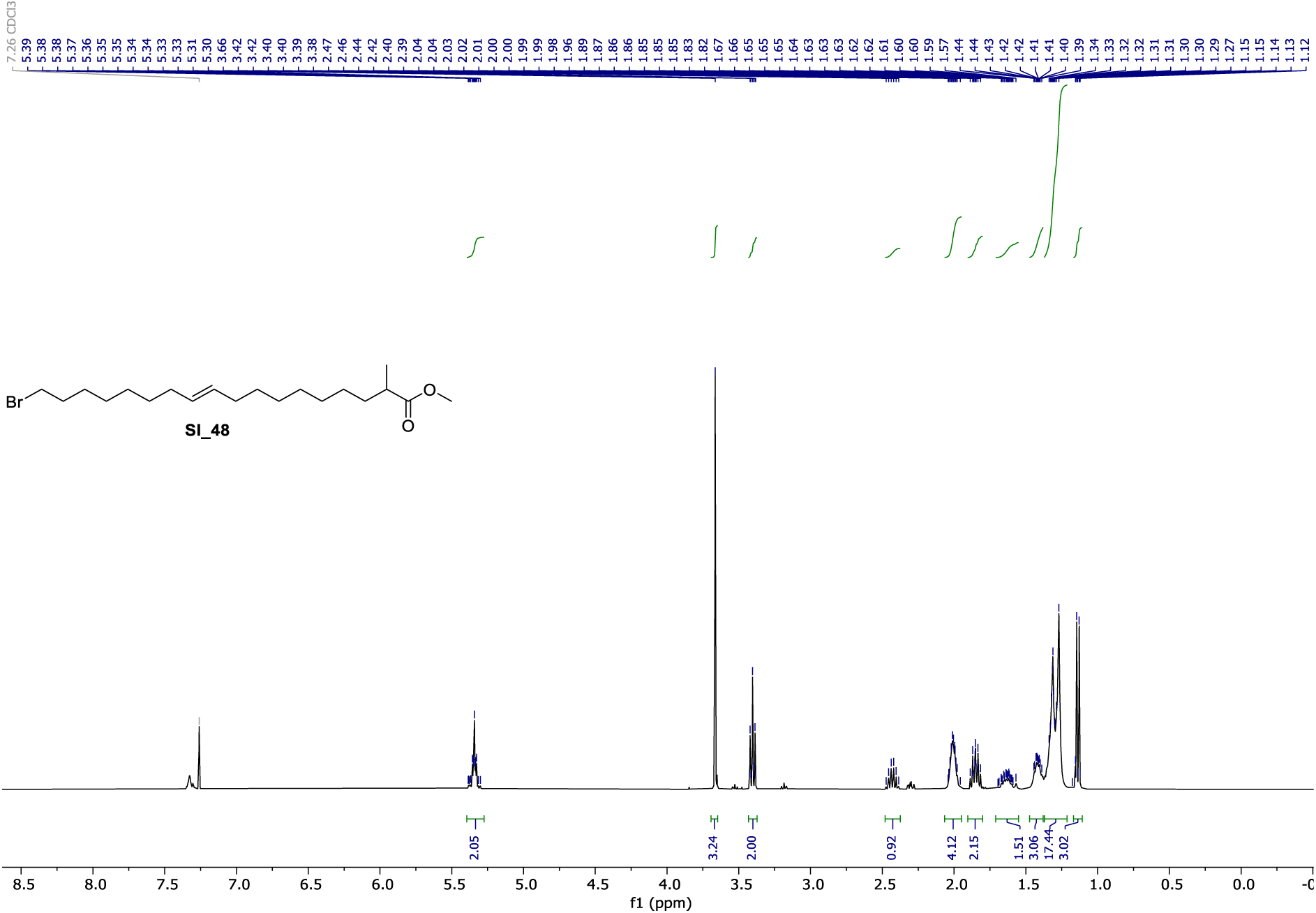

**^13^C NMR** (101 MHz, CDCl_3_) of **SI_48**.

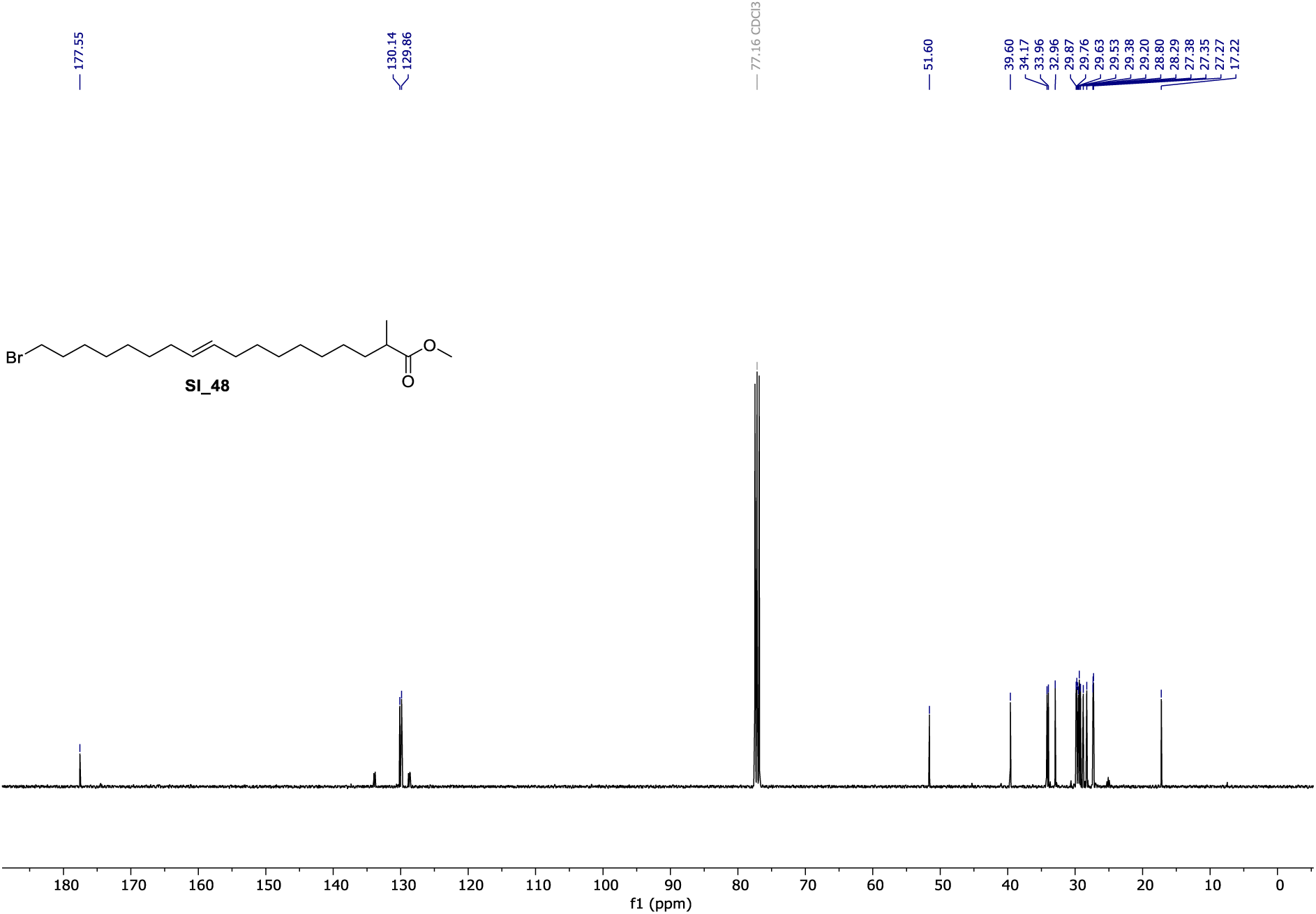

**^1^H NMR** (400 MHz, CDCl_3_) of **SI_49**.

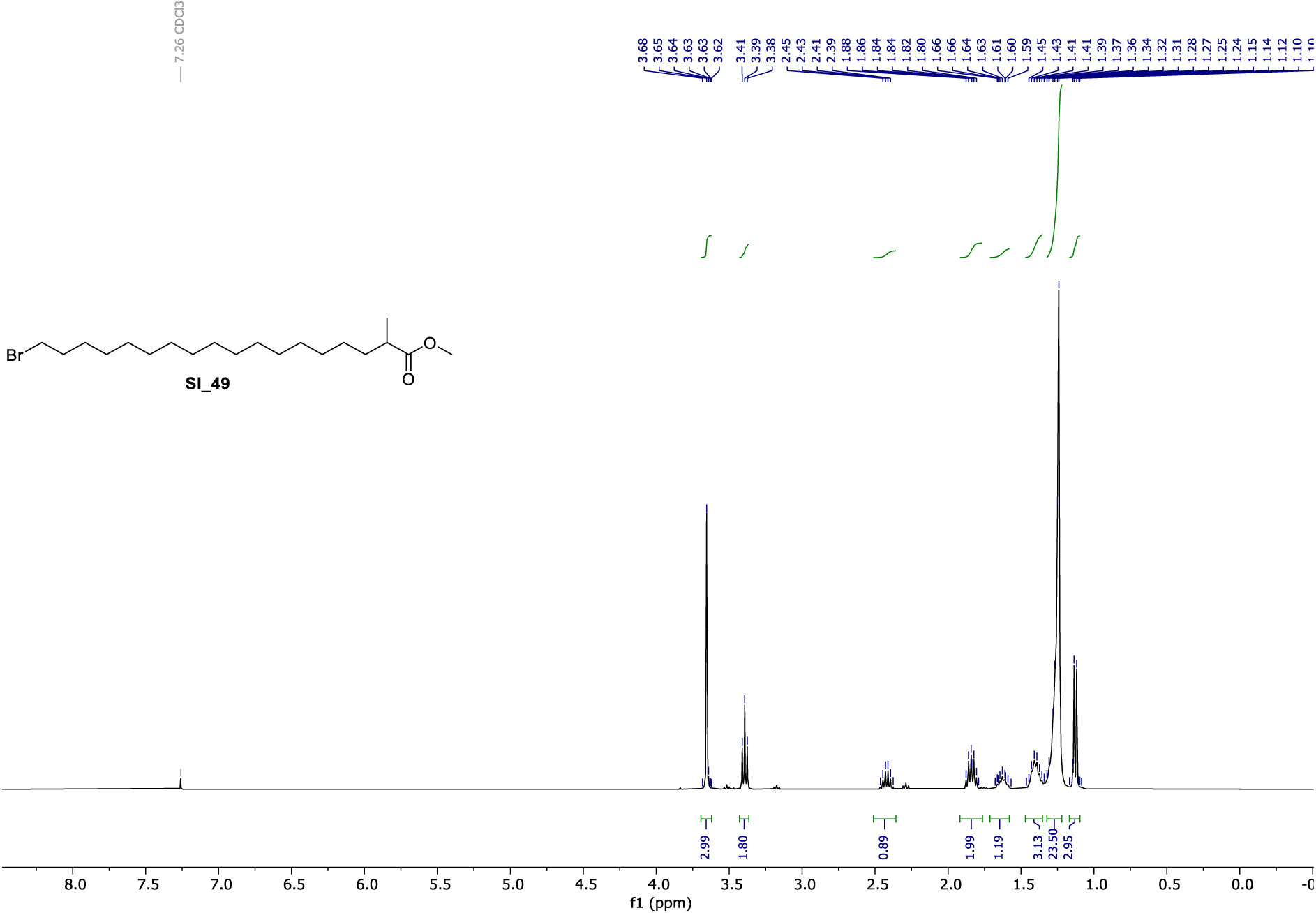

**^13^C NMR** (101 MHz, CDCl_3_) of **SI_49**.

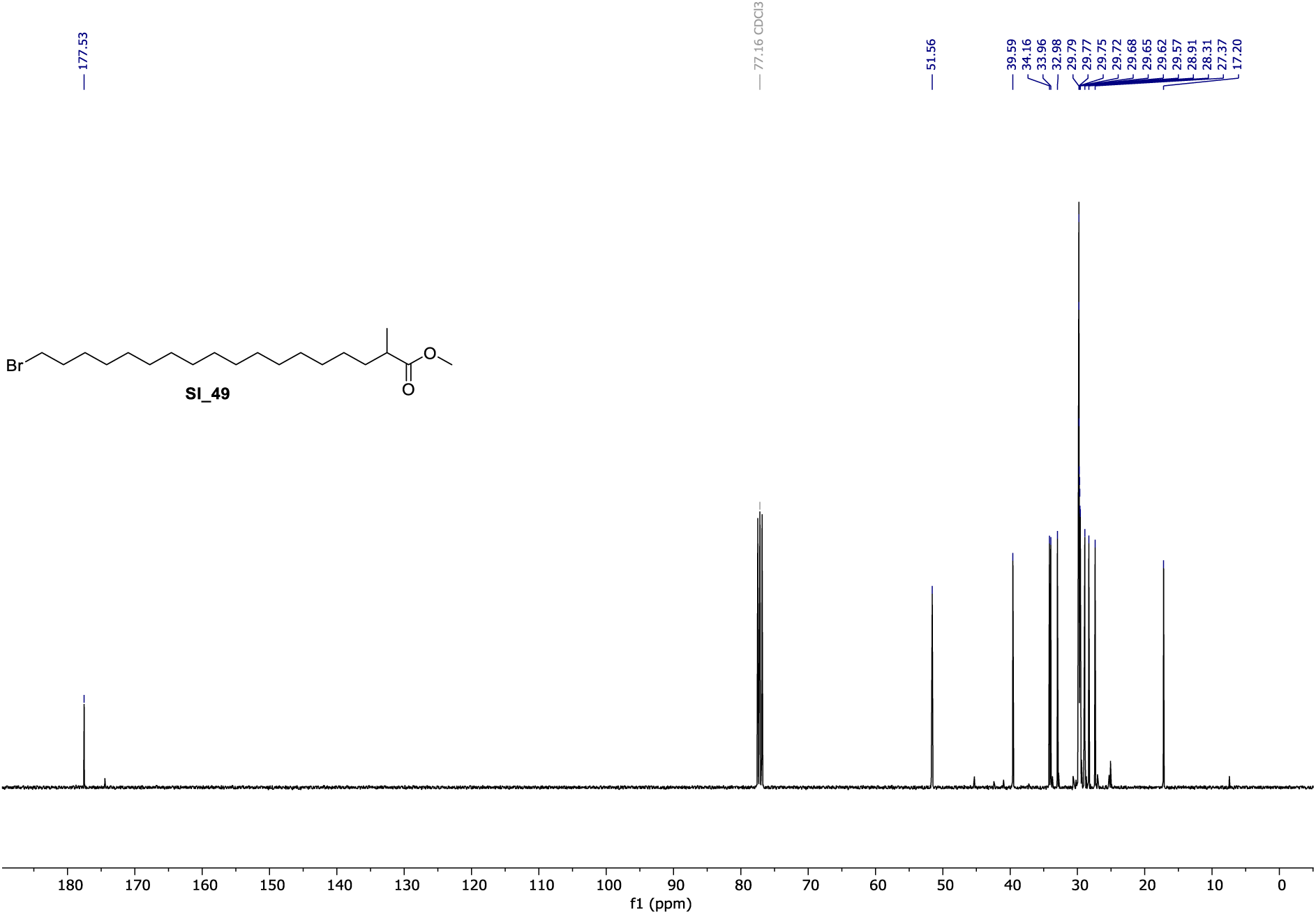

**^1^H NMR** (400 MHz, CDCl_3_) of **cl-2-MeSA**.

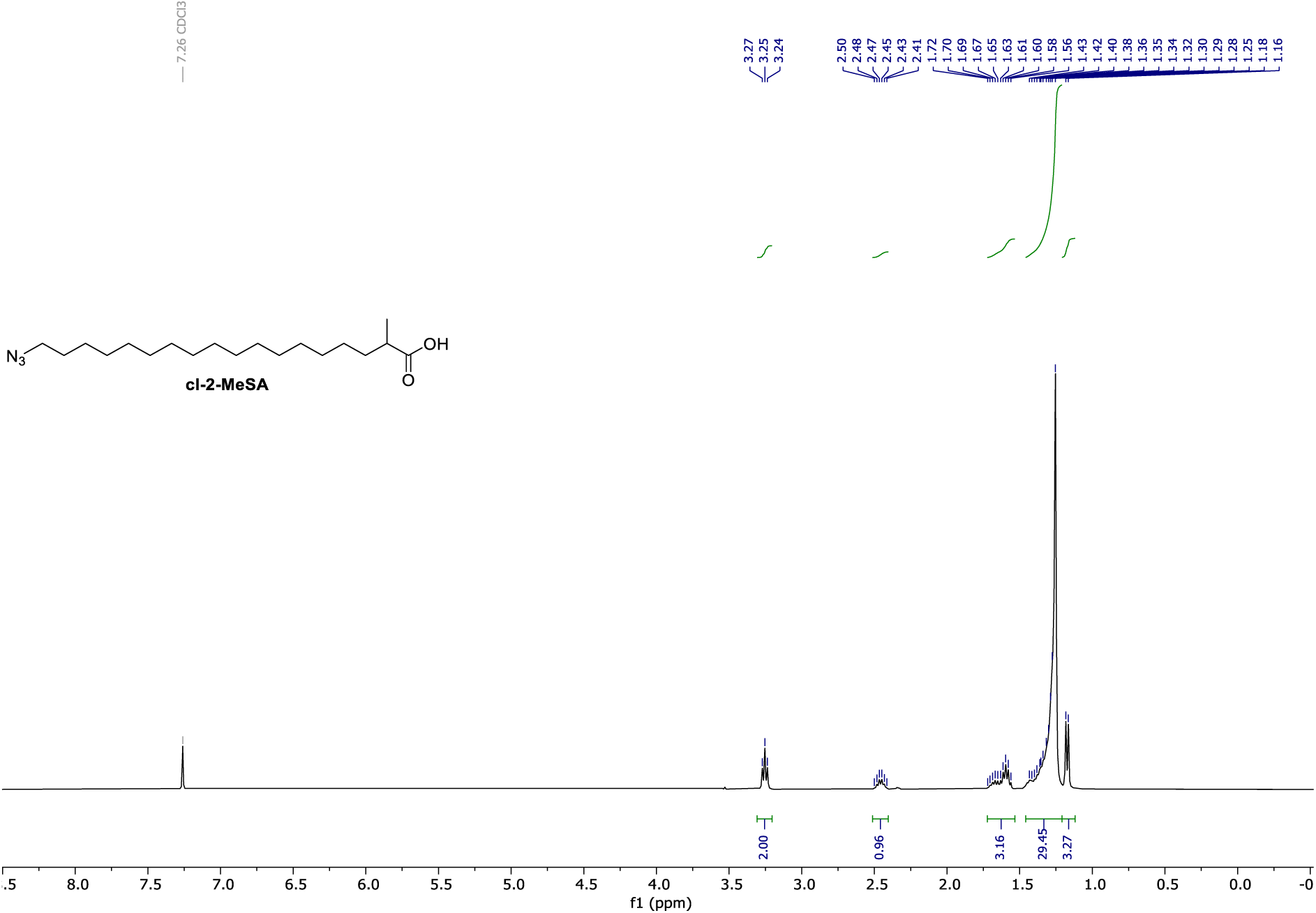

**^13^C NMR** (101 MHz, CDCl_3_) of **cl-2-MeSA**.

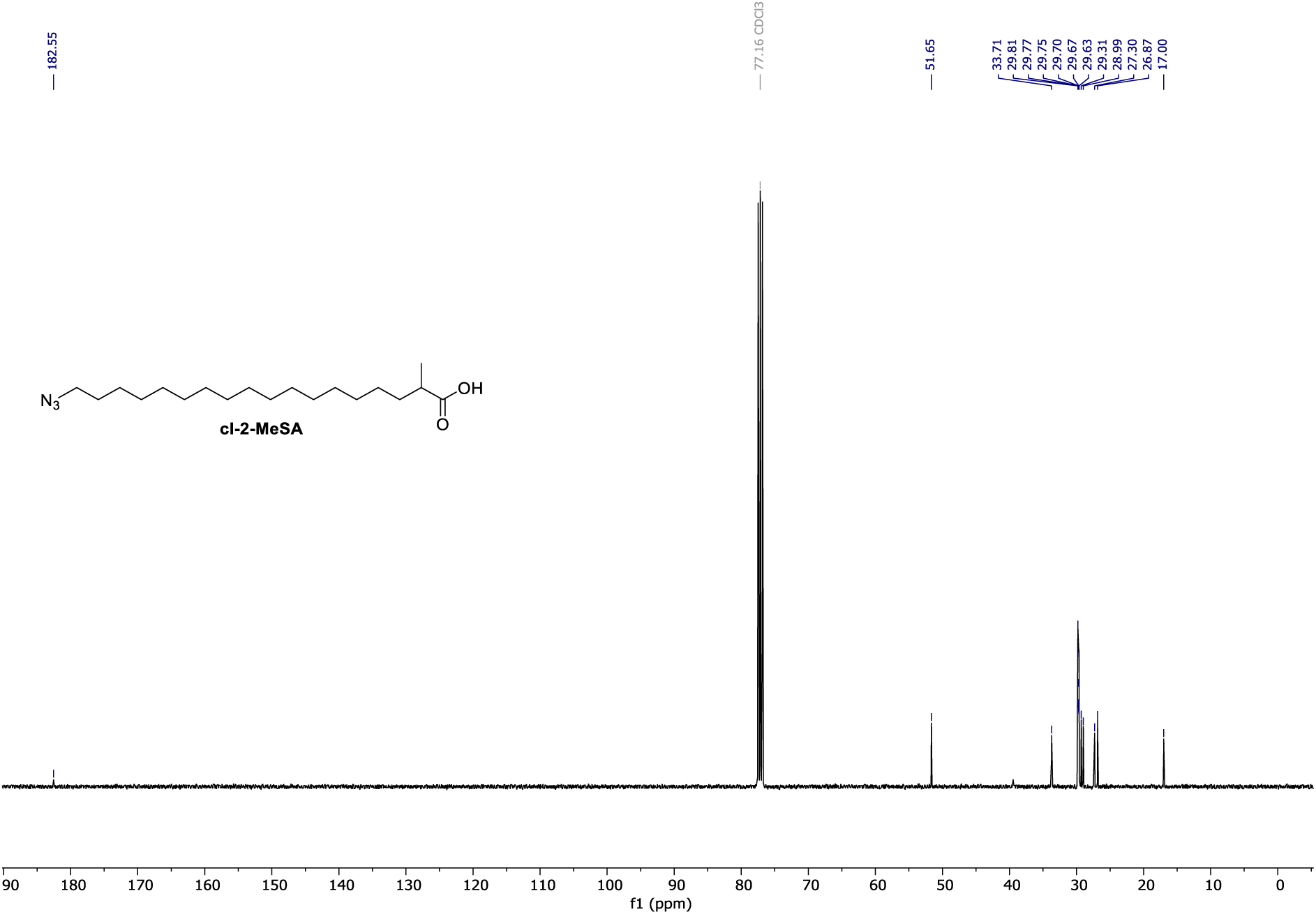

